# Rationally Engineered, Chemically Stable Tunicamycin Analogues Decouple DPAGT1 Inhibition from Non-Selective Toxicity

**DOI:** 10.64898/2026.07.28.741246

**Authors:** Katsuhiko Mitachi, Antonio Sánchez-Ruiz, David Mingle, Iván Cheng-Sánchez, Jacob M. Kirsh, Francisco Sarabia, William M. Clemons, Michio Kurosu

**Author notes:** **Corresponding Author Michio Kurosu** - Department of Pharmaceutical Sciences, College of Pharmacy, University of Tennessee Health Science Center, 881 Madison Avenue, Memphis, TN 38163, USA.

## Abstract

Tunicamycins are potent inhibitors of dolichyl-phosphate *N*-acetylglucosamine phosphotransferase (DPAGT1) but are unsuitable for therapeutic development due to non-selective cytotoxicity, acid-labile glycosidic linkages, and poor physicochemical properties. Although prior structural modifications reduced the promiscuous toxicity of tunicamycins, the intrinsic 11′-β-1″-α trehalose-type glycosidic linkage remains chemically unstable, limiting biological durability. Here, we report a rationally engineered scaffold-stabilization strategy in which the acid-labile linkage is replaced with a chemically robust cyclitol framework, enabling the concise synthesis of chemically stable and water-soluble tunicamycin analogues in only 12 synthetic steps. From this platform, TM-Cy-TBPA (**4**) was identified as a lead DPAGT1 inhibitor that potently suppresses the proliferation of breast cancer cells by inducing G_2_-phase arrest followed by apoptosis, while exhibiting minimal cytotoxicity toward nontransformed cells. The compound shows improved solubility, and favorable pharmacokinetic exposure. These results establish tunicamycin cyclitol analogues as a structurally distinct class of selective DPAGT1-targeted anticancer agents and demonstrate that stabilization of the glycosidic linkage is an effective strategy for enhancing pharmacological selectivity, improving *in vivo* performance, and simplifying the synthetic route.

## INTRODUCTION

A number of selective MraY (translocase I) inhibitors from natural sources have been reported as leads for new antibacterial agents.^1,2,3,4^ The most promising advances come from nucleoside-based inhibitors, which are grouped into six classes: muraymycins, caprazamycins, capuramycins, liposidomycins, mureidomycins and tunicamycins. ^5, 6, 7^ Tunicamycins are relatively weak MraY/MurX inhibitors (MurX is the mycobacterial homologue) and display moderate antibacterial activity. ^8, 9^ However, tunicamycins are the best-studied molecules for disruption of *N*-glycosylation in mammalian cells via inhibition of dolichyl-phosphate *N*-acetylglucosamine-phosphotransferase 1 (DPAGT1), which triggers ER-stress-mediated apoptosis. ^10, 11, 12^ Because tunicamycins inhibit proliferation of both cancerous and noncancerous cells with poor selectivity, the cytotoxicity of nucleoside antibiotics has often been attributed to DPAGT1 interaction, despite the lack of sufficient biochemical evidence supporting this assumption.^13^ The recent identification of muraymycin A1, a nucleoside antibiotic that safely targets DPAGT1 and shows selective antiproliferative activity in DPAGT1-dependent solid tumors, overturns this assumption and redefines the therapeutic potential of this class.^6^ Previous nucleoside-based anticancer agents generally act as broadly cytotoxic compounds, targeting rapidly proliferating cells indiscriminately and consequently causing damage to nontransformed tissues. As a result, these agents exhibit a narrow therapeutic window and are frequently associated with dose-limiting toxicities, particularly in highly proliferative normal tissues (*e.g*., the bone marrow, gastrointestinal epithelium, and hair follicles).^14,15,16^ This lack of selectivity underscores a fundamental limitation of conventional nucleoside analogues and highlights the need for therapeutic strategies that achieve tumor-specific targeting while minimizing systemic toxicity. One promising approach is the development of nucleoside-based agents that act through mechanisms distinct from the inhibition of DNA polymerization, thereby improving both selectivity and therapeutic efficacy. In this context, tunicamycins and muraymycin A1 are structurally distinct nucleoside natural products that provide unique molecular scaffolds for the design of selective DPAGT1 inhibitors.^9^ Among the DPAGT1-targeting nucleoside natural products, tunicamycins, represented by the predominant congener tunicamycin V (TM-V; Figure 1), are structurally less complex than muraymycin A1. This reduced structural complexity makes TM-V an attractive and synthetically tractable scaffold for the rational development of safer, DPAGT1-targeted therapeutics.^8^

**Figure 1.**
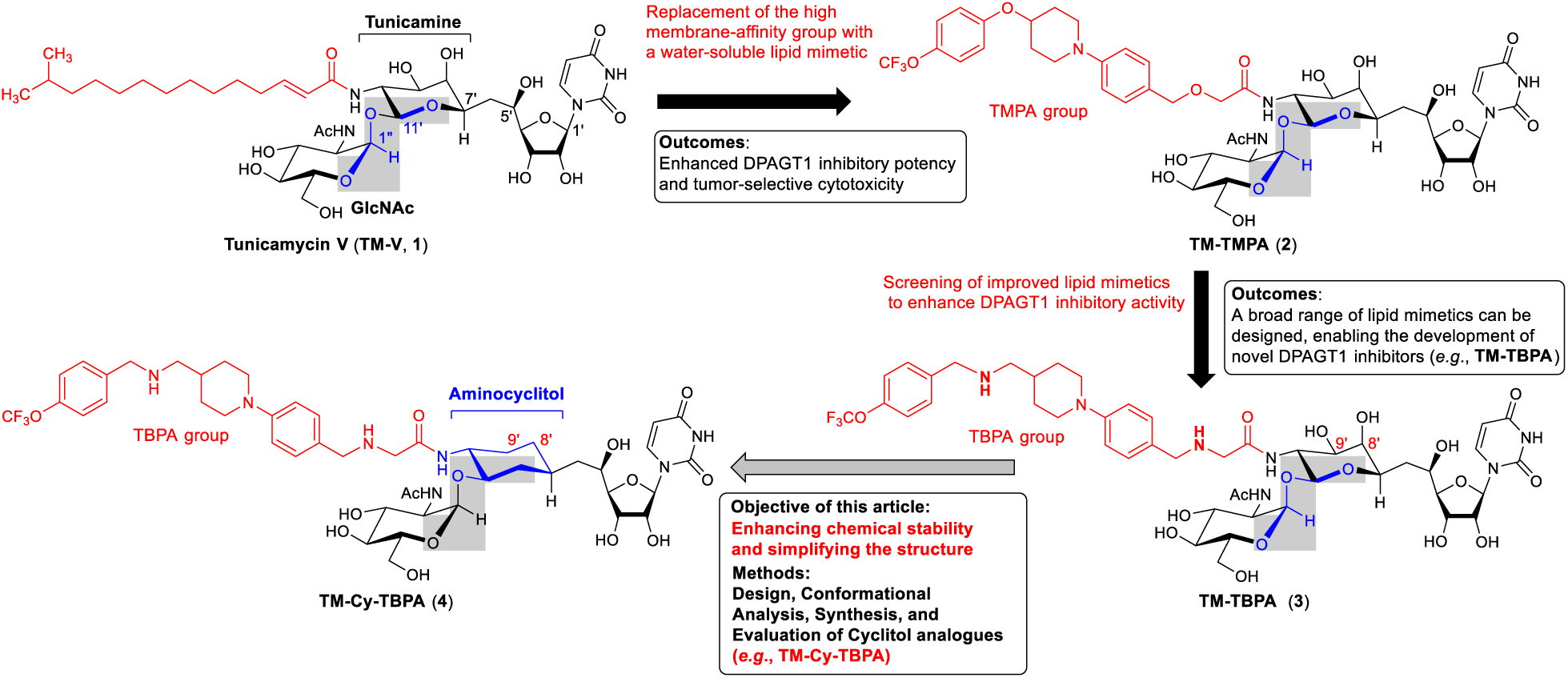
Design of chemically and metabolically stable tunicamycin V (TM-V)-based DPAGT1 Inhibitors towards *in vivo* applications.

We reported a concise total synthesis of TM-V, and our synthetic strategy is amenable to generating diverse TM-V analogues incorporating pharmacologically unique disaccharide and hydrophobic (lipid) groups.^8,17^ Our initial objective was to develop an improved tunicamycin analogue as a DPAGT1 inhibitor with significantly reduced nonspecific cytotoxicity. We found that replacing the native C15-fatty acid (C15:1-*iso*) chain of TM-V with a water-soluble lipid mimetic, [((((trifluoromethoxy)phenoxy)piperidin-1-yl)phenyl)methoxymethyl (TMPA)], enhances both DPAGT1 inhibitory potency and selective cytotoxicity (**1**→**2** in Figure 1).^8,13^ Expanding our evaluation of selectively cytotoxic DPAGT1 inhibitors with antimetastatic activity, we identified 18 novel tunicamycin analogues (see Supporting Information) with activity comparable to or exceeding that of TM-TMPA (**2**). TM-TBPA (**3**), a tunicamycin analogue incorporating the [(4-(trifluoromethoxy)benzylpiperidin-4-yl-methylamino-benzylamino)acetamide (TBPA)] lipid mimetic, emerged as a leading candidate. It exhibited a 1.2-fold enhancement in DPAGT1 inhibitory potency (IC₅₀ = 0.10 µM) and more than 5-fold greater antiproliferative and antimetastatic activity (EC₅₀ = 0.29–0.55 µM) across a panel of breast cancer cell lines. TM-TBPA (**3**) also demonstrated robust antimigration activity in triple-negative breast cancer models and induced endoplasmic reticulum (ER) stress–mediated responses, ultimately leading to apoptosis. Although we have continued to exploit the natural core scaffold to identify water-soluble lipid mimetics for the discovery of improved DPAGT1 inhibitors with antimetastatic and apoptosis-inducing activity, TM-V natural core analogues exhibit relatively short biological half-lives in pharmacokinetic studies (*e.g*., *t*_½_ = 2.6 h for **3** in CD-1 mice). In addition, the acid lability of the 11′-β-1″-α-glycosidic linkage within the disaccharide moiety makes formulation, particularly salt formation, highly challenging for *in vivo* studies. To identify pharamcologically effective DPAGT1 inhibitors of TM-V analogues, we have intruduced the cyclitol (TM-Cy) core structure which aims at improving chemical stability and PK properties (**3**→**4** in Figure 1). This study presents the rational design of novel TM-V–derived DPAGT1 inhibitors with enhanced chemical stability through an integrated strategy combining conformational analyses in aqueous and hydrophobic environments, rigorously stereocontrolled synthesis, and comprehensive physicochemical and biological characterization. Collectively, these studies establish a foundational structure–stability and pharmacological design framework for the future structure–activity relationship (SAR) optimization of selective DPAGT1 inhibitors based on the tunicamycin structure, providing a strong basis for advancing these compounds toward *in vivo* proof-of-pharmacological-concept studies and subsequent preclinical development.

## Results and Discusions

### 1. Rational Design of Tunicamycin-Derived Cyclitol Pharmacophores

The tunicamycin cyclitol analogue, TM-Cy-TBPA (**4**) is a C8’,9’-dideoxy derivative of TM-TBPA (**3**) that incorporates an aminocyclohexanol core in place of the native aminotetrahydro-2*H*-pyran-triol (tunicamine) moiety, therefore replacing the acid-labile glycosidic linkage with a chemically robust, non-trehalose-type scaffold (Figure 1). To assess whether compound **4** retains productive interactions with DPAGT1 comparable to those of TM-V, conformational analyses and molecular docking studies were performed to characterize and compare the binding modes of TM-V and TM-Cy-TBPA within the DPAGT1 active site.

#### 1.1. Comparative Analysis of the Conformational Landscapes and Thermodynamic Unfolding Penalties of TM-Cy-TBPA and TM-V

To validate the molecular design strategy (Figure 1) prior to extensive synthetic efforts, conformational and thermodynamic analyses were conducted to assess whether the designed analogue **4** exhibits favorable binding characteristics toward DPAGT1. These studies were undertaken to reduce the risk of advancing structurally attractive but conformationally unfavorable candidates that could compromise target engagement. ^18^ Specifically, we investigated the energetic accessibility of the extended, bioactive conformations of TM-V (**1**) and TM-Cy-TBPA (**4**) by comparing their relative energies with those of the corresponding compact, folded conformations. The calculations were carried out in solvent environments with contrasting dielectric properties to establish a thermodynamic basis for interpreting the subsequent biochemical and biological data. In addition, comparative analyses of the conformational landscapes and thermodynamic unfolding penalties of TM-Cy-TBPA and TM-V were performed to determine whether the engineered cyclitol scaffold and lipid-mimetic modifications preserved the bioactive conformational features required for productive interaction with the DPAGT1 active site. Such analyses provide important insight into the energetic accessibility of binding-competent conformations and help rationalize the observed biochemical and biological activities of the designed analogues.

Initial conformational ensembles for TM-V (**1**) and TM-Cy-TBPA (**4**) were generated using GOAT/GFN2-xTB implementation in ORCA 6.1.1 with ALPB implicit aqueous solvation and truncated to a 3.0 kcal/mol relative free-energy window, capturing >99.4% of the thermodynamically relevant conformational population at room temperature.^19,20^ Boltzmann weighting of the GFN2-xTB relative free energies at 25 °C was used to estimate relative conformer populations within each ensemble. Geometry optimizations and frequency calculations were subsequently performed at the PBE0-D4/def2-SVP level with CPCM (conductor-like polarizable continuum model) implicit solvent model (water), yielding qRRHO-corrected thermochemical Gibbs energies using Grimme’s quasi-rigid rotor harmonic oscillator approach,^21^ which replaces the standard harmonic oscillator treatment of low-frequency vibrational modes with a damped free-rotor interpolation to avoid artificial entropy inflation. Single-point electronic energies were then calculated at the higher PBE0-D4/def2-TZVPP level using the SMD solvation model. Final Gibbs free energies (ΔG_extended_ or ΔG_folded_) were obtained by adding the PBE0-D4/def2-TZVPP electronic energies (E_elec_) and the qRRHO thermochemical corrections (G_corr_) from the PBE0-D4/def2-SVP step (Table 1).

**Table 1.** Thermodynamic components and final unfolding free-energy penalties (ΔG_unfold_) in aqueous and lipophilic environments.ᵃ.

| Solvent | Compound | $E_{\text{elec, folded}}$<br>(hartree) | $E_{\text{elec, extended}}$<br>(hartree) | $G_{\text{corr, folded}}$<br>(hartree) | $G_{\text{corr, extended}}$<br>(hartree) | $\Delta G_{\text{unfold}}$<br>(kcal/mol) |
| --- | --- | --- | --- | --- | --- | --- |
| Water | TM-V (**1**) | -2906.053722 | -2906.035883 | 0.934514 | 0.928772 | +7.59 |
|  | TM-Cy-TBPA (**4**) | -3564.041218 | -3564.023898 | 1.061344 | 1.057062 | +8.18 |
| 1-Octanol | TM-V (**1**) | -2906.056093 | -2906.042168 | 0.933530 | 0.927700 | +5.08 |
|  | TM-Cy-TBPA (**4**) | -3564.023290 | -3564.014531 | 1.059694 | 1.055073 | +2.60 |

<sup>a</sup> The single-point electronic energies ( $E_{\text{elec}}$ , PBE0-D4/def2-TZVPP/SMD) are added to the qRRHO thermal corrections to the Gibbs free energy ( $G_{\text{corr}}$ , PBE0-D4/def2-SVP/CPCM) to yield the final absolute Gibbs free energies used to calculate the unfolding penalty ( $\Delta G_{\text{unfold}}$ ).

The studied conformers included the folded global minima of TM-V (**1**) and TM-Cy-TBPA (**4**), as well as their corresponding extended (unfolded) conformations derived from DPAGT1 docking poses in water (Figure 2). For the extended conformers, geometric restraints were applied to the aliphatic tail during geometry optimization to prevent artificial hydrophobic collapse under implicit solvent conditions. The resulting Gibbs free energies were used to evaluate the thermodynamic penalty associated with conformational unfolding (ΔG_unfold_ = G_extended_ - G_folded_) in both aqueous and lower-dielectric lipophilic environments (1-octanol). TM-V (**1**) populates a broad, quasi-continuous ensemble of folded conformers, with a high density of nearly isoenergetic microstates within 1.0 kcal/mol of the global minimum. Structural clustering (RMSD <0.5 Å) revealed that 85% of the pruned ensemble concentrates into just five clusters, with the dominant cluster alone accounting for 67% of all conformers, all adopting closely related elongated, folded geometries. TM-Cy-TBPA (**4**), by contrast, displays a markedly broader geometric footprint, with 85% of its pruned population dispersed across 35 distinct structural clusters spanning both elongated and oblate molecular shapes. However, this geometric diversity coexists with a sharply skewed thermodynamic distribution, in which over 50% of the total conformational population concentrates into just two specific geometries, separated from the remaining clusters by substantial energetic gaps. This apparent discrepancy (high geometric diversity but low thermodynamic population diversity) is a consequence of the intense Coulombic repulsion between the two protonated ammonium centers on the TBPA lipid mimetic in the folded state, while the conformational search identifies numerous geometrically distinct folded topologies, the internal charge repulsion renders all but a small number of them highly penalized energetically, confining the molecule into highly strained, specific local minima. The conformational population distributions, together with structural clustering and shape analyses for both compounds, are provided in the Supporting Information. TM-V (**1**) exhibited a clear thermodynamic preference for the compact folded resting state, with a calculated free-energy penalty of ΔG_unfold_ = +7.59 kcal/mol in aqueous solution to access the extended, bioactive conformation (Table 1). This barrier is driven by the classical hydrophobic effect;^22^ the C15:1-*iso* aliphatic chain spontaneously packs against the polar molecular core to minimize the energetically unfavorable exposure of the hydrophobic surface to the aqueous environment. This unfolding penalty constitutes a thermodynamic obstacle to productive DPAGT1 engagement, as structural evidence from crystallographic, cryo-EM, and computational analyses collectively establishes that binding within the DPAGT1 active site requires the ligand to adopt a fully extended conformation, enabling insertion of the aliphatic chain into the hydrophobic cleft of the enzyme.^23,24^ According to classical binding thermodynamics, the energetic cost of accessing this bioactive extended state must be compensated by favorable ligand–protein interactions at the binding pocket.^25,26^ In aqueous media, TM-Cy-TBPA (**4**) displays a modestly higher unfolding penalty (ΔG_unfold_ = +8.18 kcal/mol, Table 1). This elevated barrier in bulk water may act as a protective mechanism; the high aqueous dielectric (ε ≈ 80), which effectively screens the internal Coulombic repulsion between the two protonated amines, locking the molecule into a compact, soluble resting state. However, a markedly different conformational landscape emerges upon transition to a lower-dielectric lipophilic environment. In 1-octanol, which was employed here as a validated computational surrogate for the hydrophobic interior of a buried protein binding pocket,^27^ TM-V (**1**) undergoes only a modest reduction in its unfolding penalty ΔG_unfold_ = +5.08 kcal/mol). TM-Cy-TBPA (**4**), by contrast, undergoes a dramatic thermodynamic reversal: its unfolding penalty collapses to ΔG_unfold_ = +2.60 kcal/mol (Table 1), a reduction of approximately 5.6 kcal/mol relative to its behavior in bulk water and a 2.5 kcal/mol advantage over TM-V in the same lipophilic environment. This pronounced and selective solvent-dependence reveals that the energetic accessibility of the extended conformation of TM-Cy-TBPA is fundamentally controlled by environmental electrostatics, allowing it to be substantially stabilized in the low-dielectric environment of the enzyme active site relative to bulk aqueous solvent. These results revealed that TM-V (**1**) and TM-Cy-TBPA (**4**) achieved productive DPAGT1 engagement through fundamentally distinct thermodynamic mechanisms. TM-V is amphipathic, with its hydrophobic lipid tail sequestered through intramolecular folding in aqueous environments. Consequently, the energetic penalty associated with adopting the extended conformation remained moderate and only modestly dependent on solvent polarity (+7.59 kcal/mol in water and +5.08 kcal/mol in 1-octanol). TM-Cy-TBPA engages DPAGT1 through a distinct binding mechanism governed by electrostatic interactions. In bulk water, the extended conformation is disfavored because the aqueous dielectric shields the internal charge repulsion, resulting in an unfolding penalty marginally exceeding that of TM-V (+8.18 kcal/mol). Upon transition to the low-dielectric environment of the DPAGT1 hydrophobic cleft, however, this dielectric shielding is lost,^28^ and the unshielded Coulombic repulsion between the two cationic centers is magnified, heavily destabilizing the folded resting state and triggering a spontaneous Coulombic unfolding of the molecule. ^29^ This structural transition is governed by the thermodynamics of hydrophobic polyelectrolytes, where unshielded electrostatic repulsion forcefully overcomes the tendency for compact hydrophobic collapse.^30^ This reduces the unfolding penalty to +2.60 kcal/mol, well within the conventional energetic window of approximately 3–5 kcal/mol generally tolerated for productive ligand binding.^20^ TM-Cy-TBPA therefore does not achieve conformational competence through pre-organization in solution; rather, it is thermodynamically activated by the lipophilic environment of the active site itself. The molecule exploits the decrease in dielectric constant within the binding pocket to electrostatically trigger a conformational transition that renders the bioactive extended conformation accessible. The contrasting solvent-dependent thermodynamic profiles of TM-V (**1**) and TM-Cy-TBPA (**4**) carry direct mechanistic implications for DPAGT1 inhibition. For TM-V, the unfolding penalty in lipophilic environments (+5.08 kcal/mol) modestly exceeds the conventional 3–5 kcal/mol threshold associated with efficient ligand recognition,^20^ and must be offset during binding by the enthalpic stabilization provided by deep insertion of the C15:1-*iso* aliphatic chain into the hydrophobic cleft and extensive van der Waals contacts throughout the nucleoside recognition region, together with the favorable entropic contribution arising from the release of ordered water molecules from the buried active site, a hallmark of the hydrophobic effect.^31,32^ For TM-Cy-TBPA (**4**), the unfolding penalty in the lipophilic environment of the active site (+2.60 kcal/mol) falls comfortably within this energetic window, such that the bioactive extended conformation becomes thermodynamically accessible without requiring exceptional compensatory interactions. By effectively lowering its unfolding penalty via the dielectric trigger, TM-Cy-TBPA retains a greater fraction of the thermodynamic driving force for binding than the parent natural product. Once the terminal trifluoromethoxy (OCF₃) group anchors the lipid tail within the hydrophobic cleft, the conserved thermodynamic binding energy can be fully redirected toward productive target engagement. This provides a purely thermodynamic rationale for the potent inhibition of DPAGT1 by TM-Cy-TBPA, without invoking auxiliary binding interactions beyond those identified by the docking studies.

**Figure 2.**
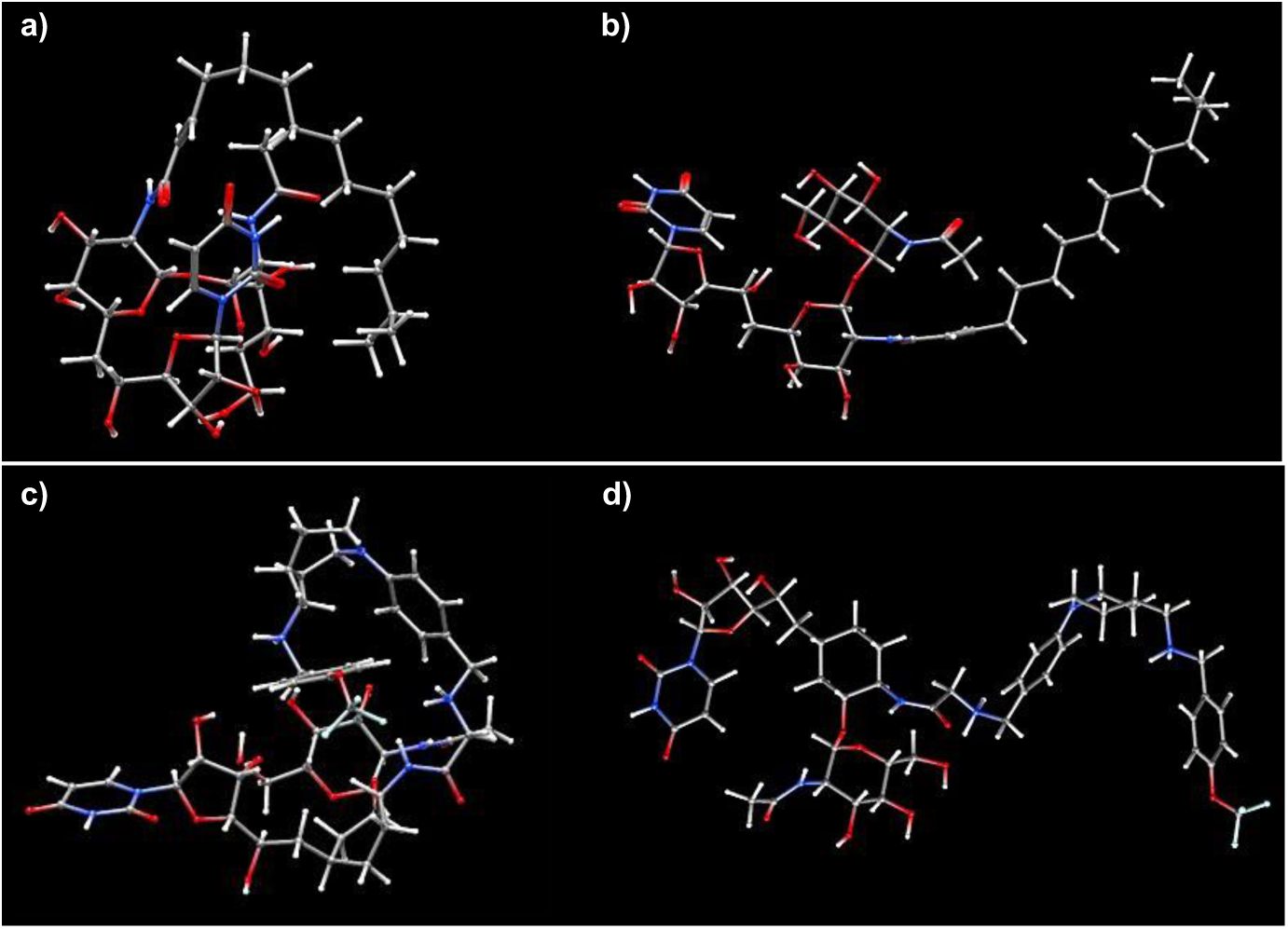
Conformational analysis of the folded and extended states of TM-V (**1**) and TM-Cy-TBPA (**4**). **Structures of the compounds studied: a**) folded TM-V in water, **b**) extended TM-V in 1-octanol, **c**) folded TM-Cy-TBPA in water, and **d**) extended TM-Cy-TBPA in 1-octanol.

### 1.2. Structural Insights into Cyclitol Analogue Engagement within the DPAGT1 Active Site

Docking calculations were performed using AutoDock Vina (v1.2.7).^33^ High-resolution structure of human DPAGT1 was retrieved from the Protein Data Bank (PDB IDs: 6BW5).^26^ Ligands were geometry optimized using the GFN2-xTB method ^34^ implemented in ORCA v6.1.1^21,22^ using the conductor-like polarizable continuum model (CPCM) with water as the solvent. The protonation states of all ligands at physiological pH (7.4) were calculated using the Kirkwood–Westheimer electrostatic model incorporating Born desolvation penalties.^35^ After assignment of the appropriate protonation states, ligand structures were converted directly to PDBQT format using Meeko software, ^36^ which assigned Gasteiger partial charges and rotatable bond definitions. No conformer generation was performed, as AutoDock Vina was configured to conduct a flexible docking search during ligand placement. The crystal structure of DPAGT1 was retrieved from the RCSB Protein Data Bank and prepared using an automated Python workflow based on PDBFixer implemented in the OpenMM software package.^37^ Crystallographic water molecules, co-crystallized solvent molecules, and non-protein heteroatoms were removed from the structure. Incomplete side-chain heavy atoms were identified, reconstructed, and repaired using PDBFixer. Explicit hydrogen atoms were added according to the predicted protonation states of titratable amino acid residues at physiological pH (7.4). The docking search box was centered on the ligand-binding pocket defined by the crystal structure, with dimensions of 62 × 25 × 25 Å to ensure complete coverage of the binding site. Docking simulations were performed with an exhaustiveness setting of 64 using the Vinardo empirical scoring function to estimate ligand binding affinities. The resulting docking poses were analyzed by visual inspection and hydrogen-bond interaction analysis (distance cutoff < 3.5 Å). Predicted ligand–DPAGT1 complexes were subsequently compared with the experimentally determined DPAGT1–ligand complex (PDB ID: 6BW5) to assess the consistency of the predicted binding modes.

Docking analysis predicted that TM-V (**1**) establishes an extensive network of polar interactions with Gln44, Leu46, Glu56, Asn119, Lys125, Asn185, and Arg303, while also forming a π–π stacking interaction with Phe249 (Figure 3B). These contacts indicate that, although explicit hydrogen bonds were not predicted during docking, the interacting residues are positioned at favorable distances and orientations to form hydrogen bonds upon minor conformational adjustments of the binding site. Notably, these interactions are consistent with those observed in the DPAGT1 crystal structure (PDB ID: 6BW5), supporting the validity of the predicted binding mode. In comparison, TM-Cy-TBPA (**4**) was predicted to interact with Gln44, Leu46, Asn185, Glu194, Arg301, and Arg303. In addition to the conserved π–π stacking interaction with Phe249 observed for TM-V (**1**), TM-Cy-TBPA (**4**) exhibited a putative π–cation interaction^38^ between Trp122 and the protonated ammonium group adjacent to the aminocyclitol moiety of the TBPA tail (Figure 3A). This additional interaction may further stabilize TM-Cy-TBPA (**4**) within the hydrophobic binding cleft of DPAGT1. Overall, the docking results demonstrate a high degree of agreement with the experimentally determined DPAGT1 structure (PDB ID: 6BW5), supporting the robustness and reproducibility of the predicted binding mode. Using the 6BW5 structure as the receptor, the lowest-energy docking poses of TM-V (**1**) and TM-Cy-TBPA (**4**) yielded docking scores of −7.752 and −6.539 kcal/mol, respectively. These results suggest that replacement of the tunicamine core with the cyclitol scaffold maintains favorable interactions with DPAGT1 and does not substantially compromise binding affinity. Although TM-Cy-TBPA (**4**) lacks the C8′,C9′-dihydroxy functionality and therefore cannot form the C9′-hydroxy hydrogen bond with Lys125 observed for TM-V (**1**), the loss of this polar interaction is compensated by enhanced hydrophobic complementarity between the TBPA lipid-mimetic moiety and the hydrophobic cleft of DPAGT1. This observation is consistent with the thermodynamic analyses described in Section 1.1, which demonstrated that the extended conformation of TM-Cy-TBPA is preferentially stabilized in low-dielectric, lipophilic environments. Despite extensive remodeling of the tunicamine scaffold, TM-Cy-TBPA retains a highly favorable binding architecture through a distinct network of complementary hydrophobic and polar interactions. These findings suggest that the cyclitol scaffold retains productive DPAGT1 recognition through a binding mode that is mechanistically distinct yet structurally comparable to that of the parent natural product.

**Figure 3.**
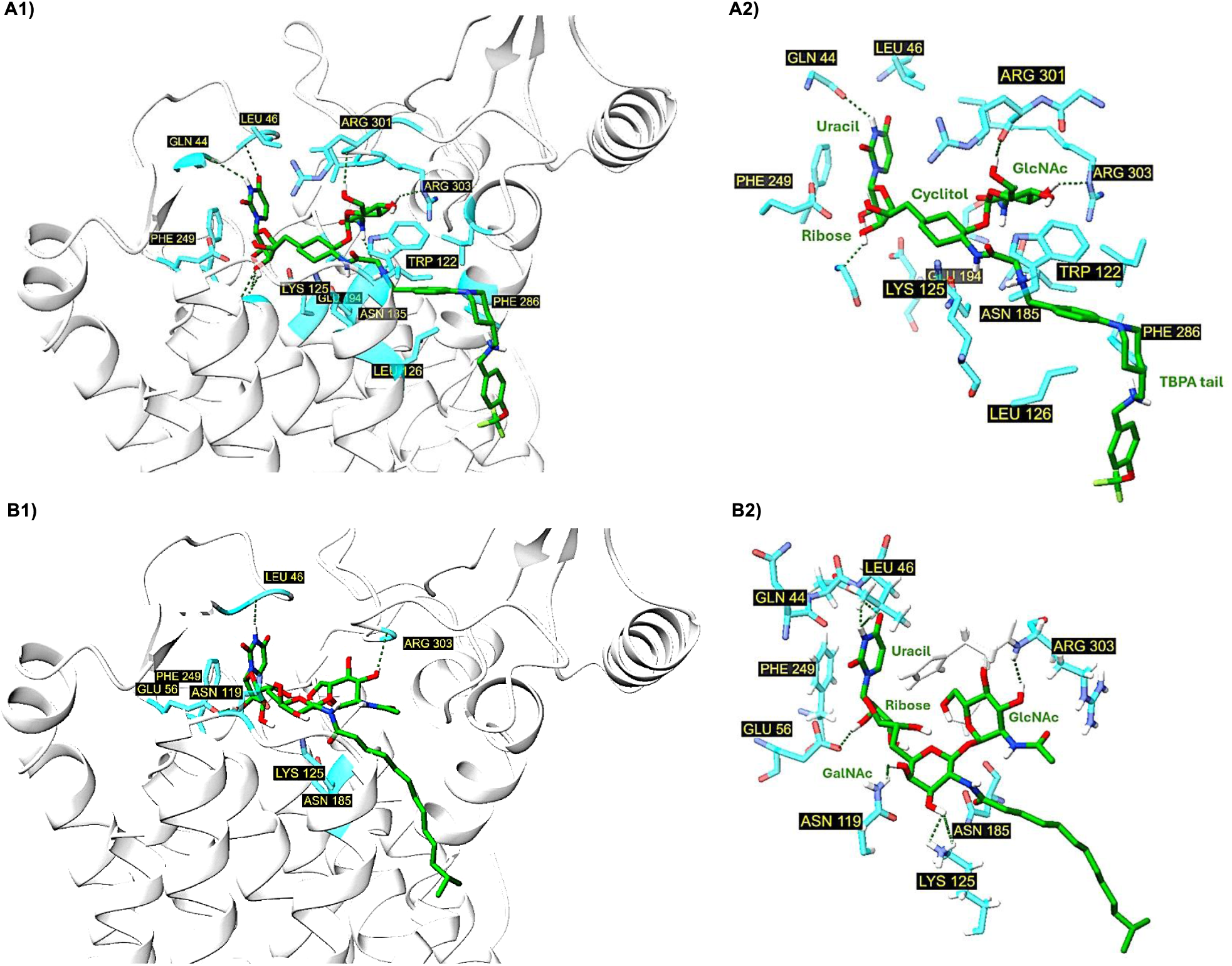
Predicted binding mode of TM-Cy-TBPA (**4**) in DPAGT1 (PDB ID: 6BW5) from Molecular Docking in reference to DPAGT1-TM-V structure. **A**: DPAGT1-TM-Cy-TBPA interactions.; **B**: DPAGT1-TM-V interactions. These figures show 3D ligand–protein interaction views, emphasizing key contacts at distances of ≤2.0 Å.

Notably, the docking studies provide strong structural support for our design strategy by demonstrating that the cyclitol scaffold serves as a conformationally competent surrogate for the native tunicamine core. The redesigned scaffold maintains the spatial organization of the uridine, GlcNAc, and lipid-mimetic pharmacophores required for productive engagement of the DPAGT1 catalytic domain. This result is particularly significant because the acid-labile tunicamine framework was successfully replaced with a chemically robust, non-1,1′-glycosidic cyclitol scaffold while largely preserving the predicted binding mode and target affinity. Furthermore, consistent with our previous observation that incorporation of an ionizable lipid-mimetic moiety into TM-TBPA (**2**) increased DPAGT1 inhibitory potency by approximately one order of magnitude relative to TM-V (**1**), the present computational studies (Figures 2 and Figure 3) further support the design hypothesis that TM-Cy-TBPA (**4**) and related cyclitol analogues retain the essential molecular recognition features required for high-affinity DPAGT1 binding while offering substantially improved chemical stability and enhanced drug-like properties.

### 2. Stereocontrolled Synthesis of TM-Cy-TBPA (4)

The synthetic strategy for the cyclitol analogue TM-Cy-TBPA (**4**) is deliberately derived from the validated route to tunicamycin V (TM-V, **1**), ensuring both strategic continuity and synthetic reliability.^8,13,19^ This platform is designed to enable modular access to a broader series of cyclitol analogues, as delineated in Figure 4. Central to this approach, a Büchner–Curtius–Schlotterbeck (BCS) reaction between *N*-nitrosoacetamide **7** and BFPM ((4,4’-bisfluorophenyl)methoxymethyl)-protected uridyl aldehyde **6** is expected to efficiently deliver the key ketone intermediate **5**, therefor establishing the carbon framework in a single convergent transformation. Following our pioneering application of the BCS reaction in natural tunicamycin synthesis, the broad utility and robustness of this methodology were subsequently established through its successful adoption in the synthesis of a complex natural product by other groups.^39,40^ The intermediate **5** synthesized by our optimized BCS reaction condition is then advanced to TM-Cy-TBPA (**4**) through a concise five-step sequence comprising asymmetric Meerwein–Ponndorf–Verley reduction, reductive acetylation, dephthaloylation–acylation, and global deprotection, providing a streamlined and stereochemically controlled route to the target molecule. The *N*-nitrosoacetamide-containing pseudo-disaccharide **7** is strategically assembled via α-glycosylation of intermediate **9**, which is readily derived from commercially available (*R*)-cyclohex-3-en-1-ylmethanol (**10**) through a sequence of highly stereoselective transformations. Collectively, this synthetic strategy integrates convergent bond construction with highly selective stereochemical control, establishing a robust and versatile platform for the synthesis of tunicamycin cyclitol analogues.

**Figure 4.**
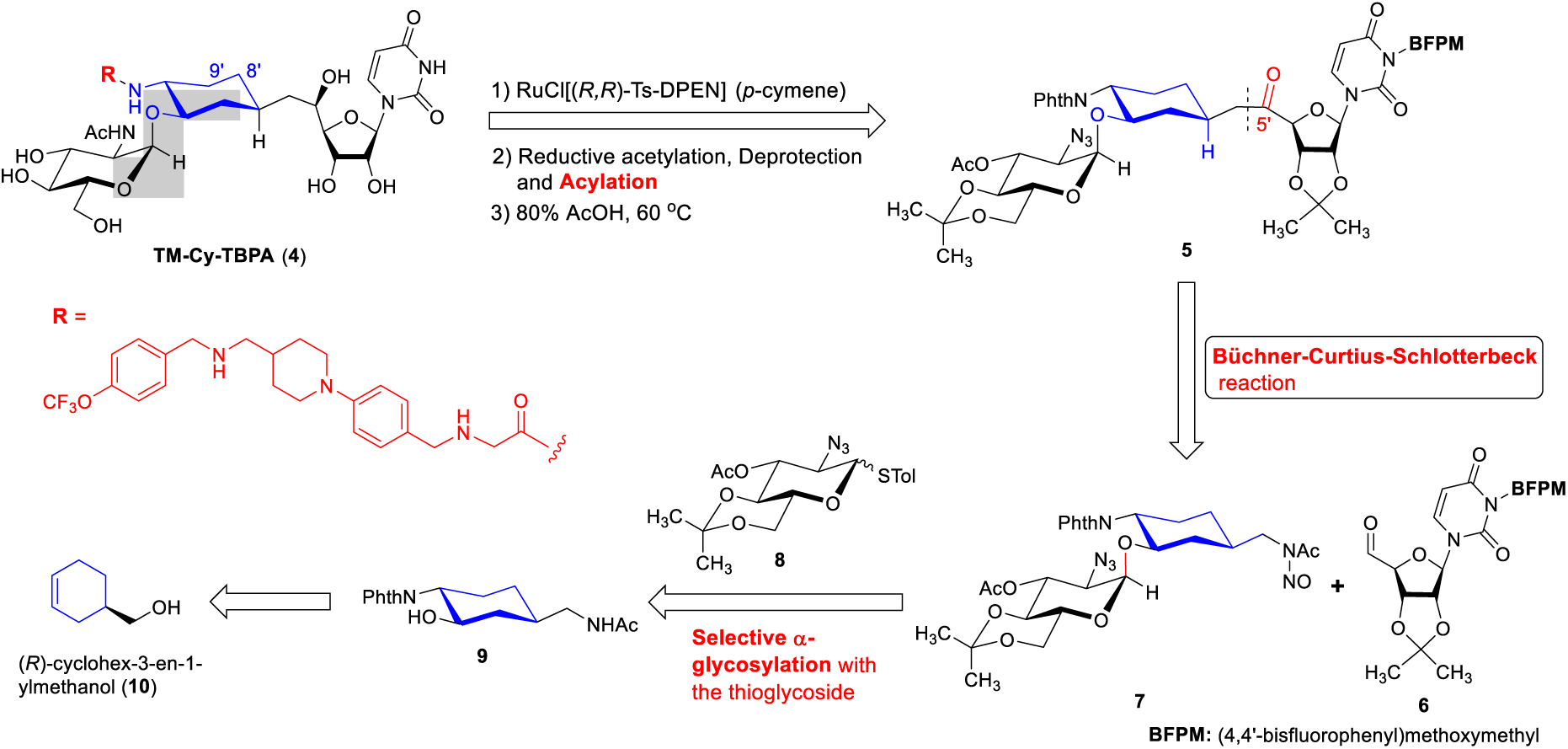
Synthetic strategy for the preparation of tunicamycin cyclitol analogues from starting material **10** via the Büchner–Curtius–Schlotterbeck (BCS) reaction.

(*R*)-Cyclohex-3-en-1-ylmethanol (**10**) is an inexpensive, commercially available starting material accessible from numerous chemical vendors. To the best of our knowledge, the stereoselective conversion of **10** into the key intermediate **9** is reported herein for the first time. A critical challenge in the synthesis of **9** is the highly stereoselective opening of epoxide **13** to furnish the desired azido-alcohol **14** exclusively. Acetamide derivative **12** was synthesized from **10** in three steps, including tosylation, azide substitution, and reductive acetylation in 86% overall yield. Subsequent epoxidation of **12** with mCPBA afforded the desired 3*R*,4*S*-epoxide **13** in 99% yield without detectable formation of the corresponding 3*S*,4*R*-diastereomer. Notably, treatment of epoxide **13** with NaN_3_/NH_4_Cl provided azido-alcohol **14** as a single diastereomer in 90% yield.

Initial stereochemical assignment of **14** as the 3*R*,4*R*-isomer was based on NOESY analysis (see compound **14** in B-2 in Scheme 1B). To unequivocally establish the stereochemistry of **14**, the undesired 3*S*,4*S*-azido-alcohol **15** was independently synthesized from epoxide **16** (Scheme 1B; see Supporting Information). Ring opening of epoxide **16** with NaN_3_/NH_4_Cl generated a separable 1:1 mixture of diastereomers **17** and **18** in 99% combined yield (49.5% each). Subsequent Ns and Boc deprotections followed by acetylation converted **17** and **18** into azido-alcohols **14** and **15**, respectively, in 75% overall yield over three steps. The stereochemistry at C3 of **14** was conclusively determined using the advanced Mosher ester method.^41^ Analysis of the Δδ(*S*−*R*) values obtained from the corresponding Mosher esters revealed positive Δδ values for the C2- and C3-protons, thereby supporting the 3*R* absolute configuration. Additional NOESY experiments further corroborated the stereochemical assignment. Compound **14** exhibited strong NOE correlations between the diaxial C3- and C5-protons, as well as among the axial C2-, C4-, and C6-protons. In contrast, these characteristic diaxial correlations were absent in **15**, which instead displayed equatorial correlations between the C3- and C4-protons, consistent with the 3*S*,4*S*-configuration. Collectively, these studies unambiguously confirmed the stereochemistry of **14** and provided mechanistic insight into the highly stereoselective opening of epoxide **13** by azide anion. As illustrated in the hypothetical transition-state model for the transformation of **13** to **14** (Scheme A-1), intramolecular hydrogen bonding between the epoxide oxygen and the NHAc amide proton likely weakens the C3−O bond, facilitating regio- and stereoselective nucleophilic attack at C3 in an SN_2_-like fashion. Azido-alcohol **14** was subsequently converted into *N*-phthaloyl-4-amino-3-hydroxycyclohexyl)methyl)acetamide **9** through hydrogenation followed by phthaloylation in 83% overall yield. The synthetic route established herein enables efficient preparation of the cyclitol glycosyl acceptor possessing the natural configurations at the C1, C3, and C4 positions in 64% overall yield from **10**. Importantly, our recent inventory survey revealed that (*R*)-*N*-(cyclohex-3-en-1-ylmethyl)acetamide (**12**, CAS No. 196703-47-6) is commercially available from suppliers such as Parchem. Thus, intermediate **9** can be synthesized from commercially available **12** in only four steps with an overall yield of 74%, further enhancing the practicality and scalability of this synthetic strategy. The synthesis of TM-Cy-TBPA (**4**) is summarized in Scheme 2. α-Glycosylation of cyclitol acceptor **9** with thioglycoside **8** under conditions (NBS, AgBF_4_) previously optimized for the synthesis of TM-V (**1**) proceeded smoothly to furnish glycoside **19** in 91% yield. The α-configuration of the glycosidic linkage was assigned on the basis of the coupling constant of H1″ [δ 4.87 (d, *J* = 3.9 Hz)] observed in the ^1^H NMR spectrum. Nitrosylation of **7** using NaNO_2_/Ac_2_O in AcOH at 0 °C afforded mono-*N*-nitrosoacetamide **7** in 91% yield. We next applied a one-pot Büchner–Curtius–Schlotterbeck (BCS) reaction developed in our laboratory.^19^ Treatment of **7** with aldehyde **6** under the optimized BCS conditions furnished ketone **5** in 90% yield. Enantioselective Meerwein–Ponndorf–Verley reduction of ketone **5** employing RuCl[(*R,R*)-Ts-DPEN](*p*-cymene) in *^i^*PrOH/HCO_2_H/Et_3_N^42^ delivered the desired 5′*R*-alcohol **20** in 92% yield with excellent stereocontrol. Subsequent reductive acetylation followed by concomitant dephthaloylation/deacetylation converted **20** into amino-alcohol **21** in 88% overall yield over two steps. Installation of the water-soluble lipid mimetic moiety **22** was accomplished using EDCI, NMM, and HOSu in DMF to provide the corresponding amide intermediate in 90% yield. Final global deprotection of the BFPM, Boc, and ketal protecting groups furnished TM-Cy-TBPA (**4**) in 85% yield. The final product was purified by reverse-phase HPLC prior to biochemical evaluation to ensure analytical purity suitable for biological studies.

**Scheme 1.**
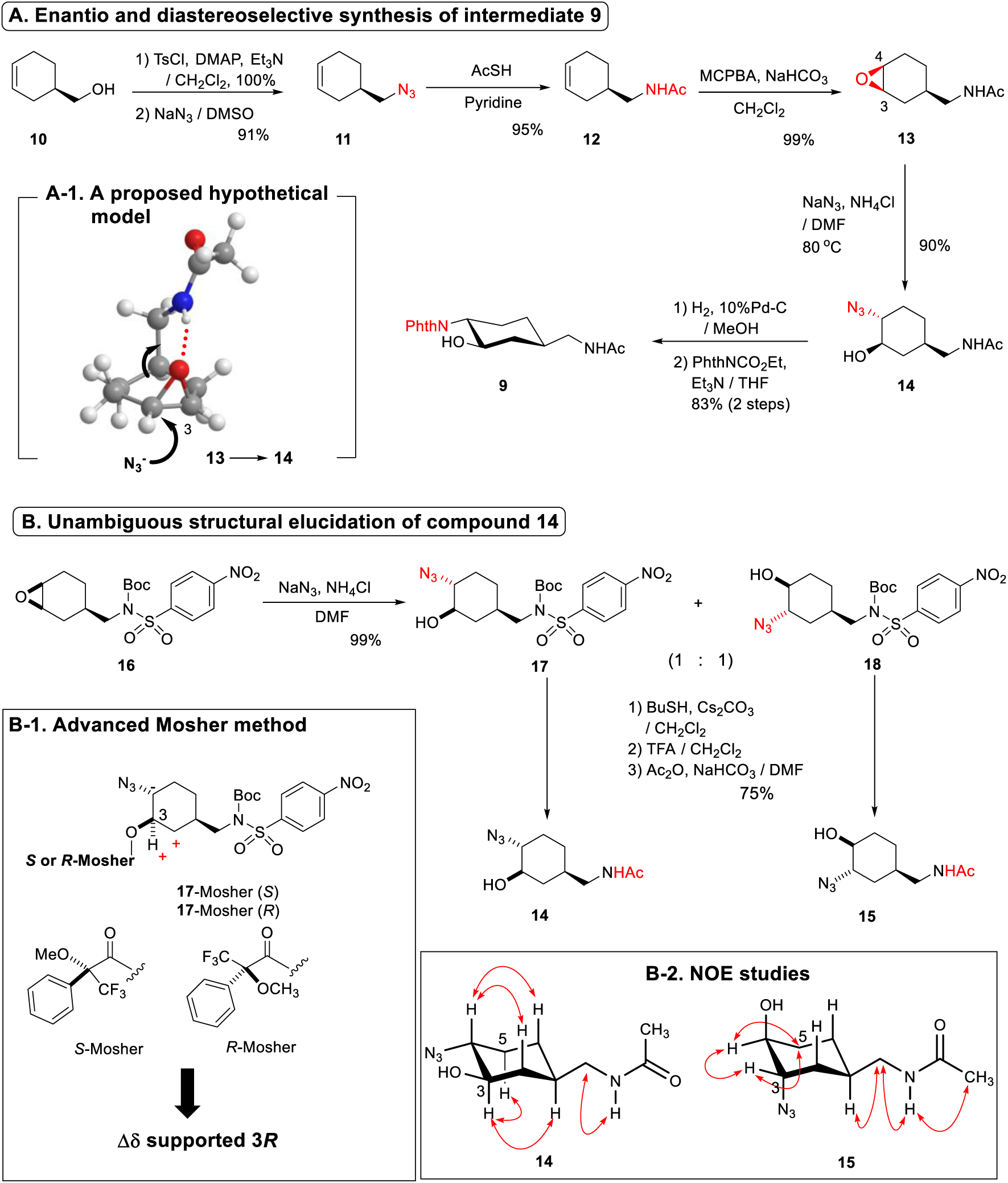
Stereoselective synthesis and structural elucidation of (1*R,*3*R,*4*R*)-*N*-phthaloyl-4-amino-3-hydroxycyclohexylmethyl acetamide (**9**).

**Scheme 2.**
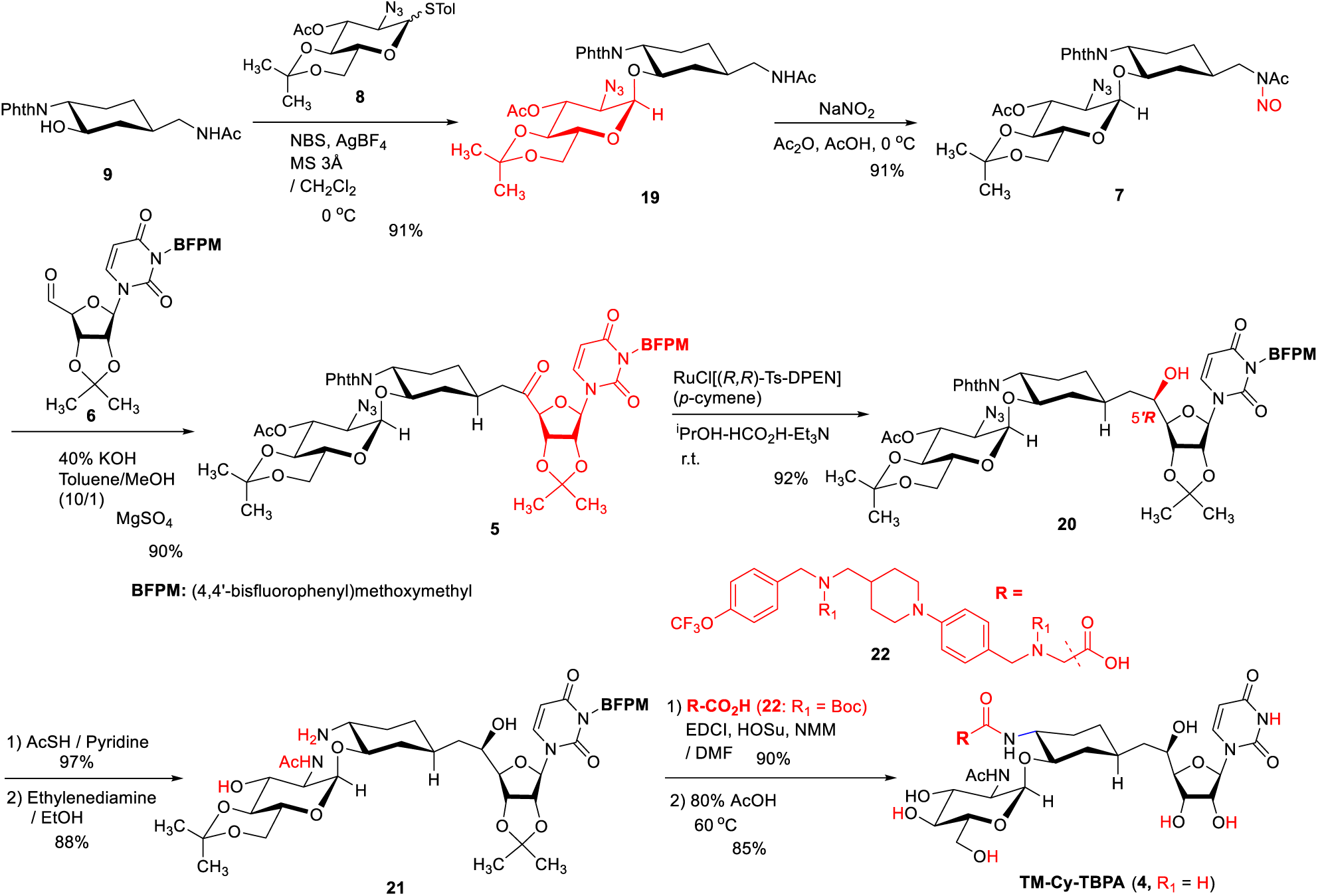
Synthesis of TM-Cy-TBPA analogue **4** through α-glycosylation and a Büchner–Curtius–Schlotterbeck (BCS) reaction.

### 3. Chemical Stability and Biochemical Properties of the Cyclitol Analogue TM-Cy-TBPA (4)

#### 3.1. Physicochemical Properties: Water Solubility and Acid Stability

Water solubility and chemical stability of cyclitol analogue **4** were initially evaluated. Compound **4** exhibited an aqueous solubility of 7.8 mg/mL, which is approximately 1.5-fold lower than that of the parent compound TM-TBPA (**3**, 12.2 mg/mL) (Supporting Information). Despite removal of the C8′,9′-diol functionality, compound **4** retained favorable aqueous solubility and readily dissolved in saline similarly to TM-TBPA. Notably, compared with TM-V (**1**, <0.2 mg/mL), compound **4** displayed markedly improved water solubility. TM-V and TM-TBPA exhibited limited stability under acidic conditions, with half-lives of approximately 1 h in 1 M HCl and 0.5 h in 2 M HCl, respectively (Figure 5). In contrast, cyclitol analogue **4** demonstrated substantially improved acid stability, remaining largely intact under both conditions with a half-life exceeding 6 h. The enhanced chemical stability of **4** is likely attributable to replacement of the acid-labile 11′-β-1″-α trehalose-type glycosidic linkage framework with the conformationally constrained cyclitol scaffold. Importantly, this improved resistance to acid-mediated degradation enabled formulation of compound **4** as its hydrochloride salt without detectable decomposition during preparation and storage. Furthermore, compound **4** readily formed a homogeneous saline solution at 2.0 mg/mL, providing a formulation suitable for subsequent biochemical, pharmacological, and pharmacokinetic evaluations. The markedly improved aqueous solubility and acid stability of compound **4** represent significant pharmaceutical advantages over TM-V and related tunicamycin analogues containing the native tunicamycin core structure.

**Figure 5.**
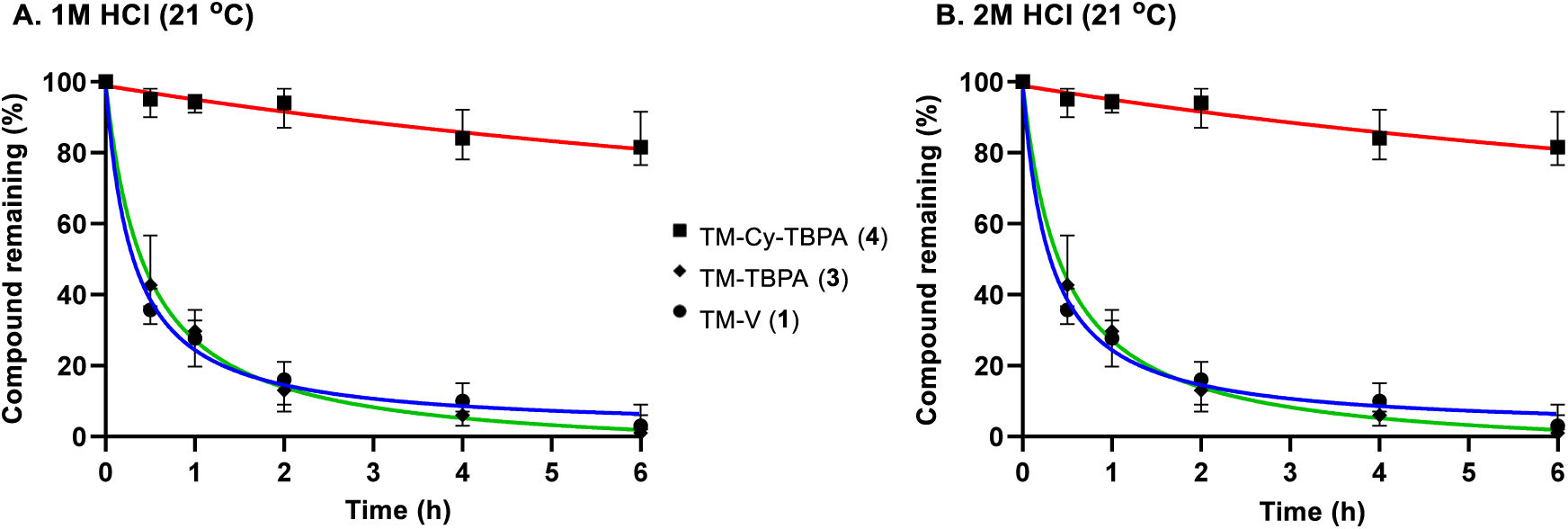
Enhanced acid stability of TM-Cy-TBPA (**4**) relative to TM-TBPA (**3**) and TM-V (**1**).^a^ ^a^ Residual amounts of the compounds were quantified by reverse-phase HPLC (see Supporting Information)

#### 3.2. Evaluation of DPAGT1 Inhibitory Activity

Conventional assays employing UDP-[^14^C]GlcNAc and natural dolichyl phosphate (Dol-P) are impractical in our laboratory setting, as separation of the radiolabeled product from excess isotope-labeled substrate is labor-intensive and poorly suited for accurate quantitative analysis. A luminescence-based assay format utilizing a commercially available UMP detection kit provides a convenient alternative for monitoring DPAGT1 enzymatic turnover. In this system, coupling enzymes convert UMP generated by the DPAGT1 reaction into ATP through sequential phosphorylation reactions, and the resulting ATP is quantified by luciferase/luciferin-mediated bioluminescence. ^43^ Although luminescence-based UMP detection assays are operationally convenient, we observed false-negative results for several DPAGT1 inhibitors developed in this study. These findings suggest that certain compounds interfered with components of the coupled detection system in a manner that masked genuine DPAGT1 inhibition rather than artificially enhancing inhibitory activity. Because the assay relies on multiple downstream enzymatic steps to generate an ATP-dependent luminescent signal, interference with coupling enzymes, nucleotide conversion, ATP generation, or luciferase activity can compromise signal formation independently of DPAGT1 catalysis. Consequently, active DPAGT1 inhibitors may appear inactive or substantially less potent than their true inhibitory activities would indicate. Therefore, DPAGT1 inhibition was further validated using an orthogonal HPLC-based assay established in our laboratory, in which formation of the glucosamine-C_6_-FITC product was directly quantified (Figure 6).^44^

**Figure 6.**
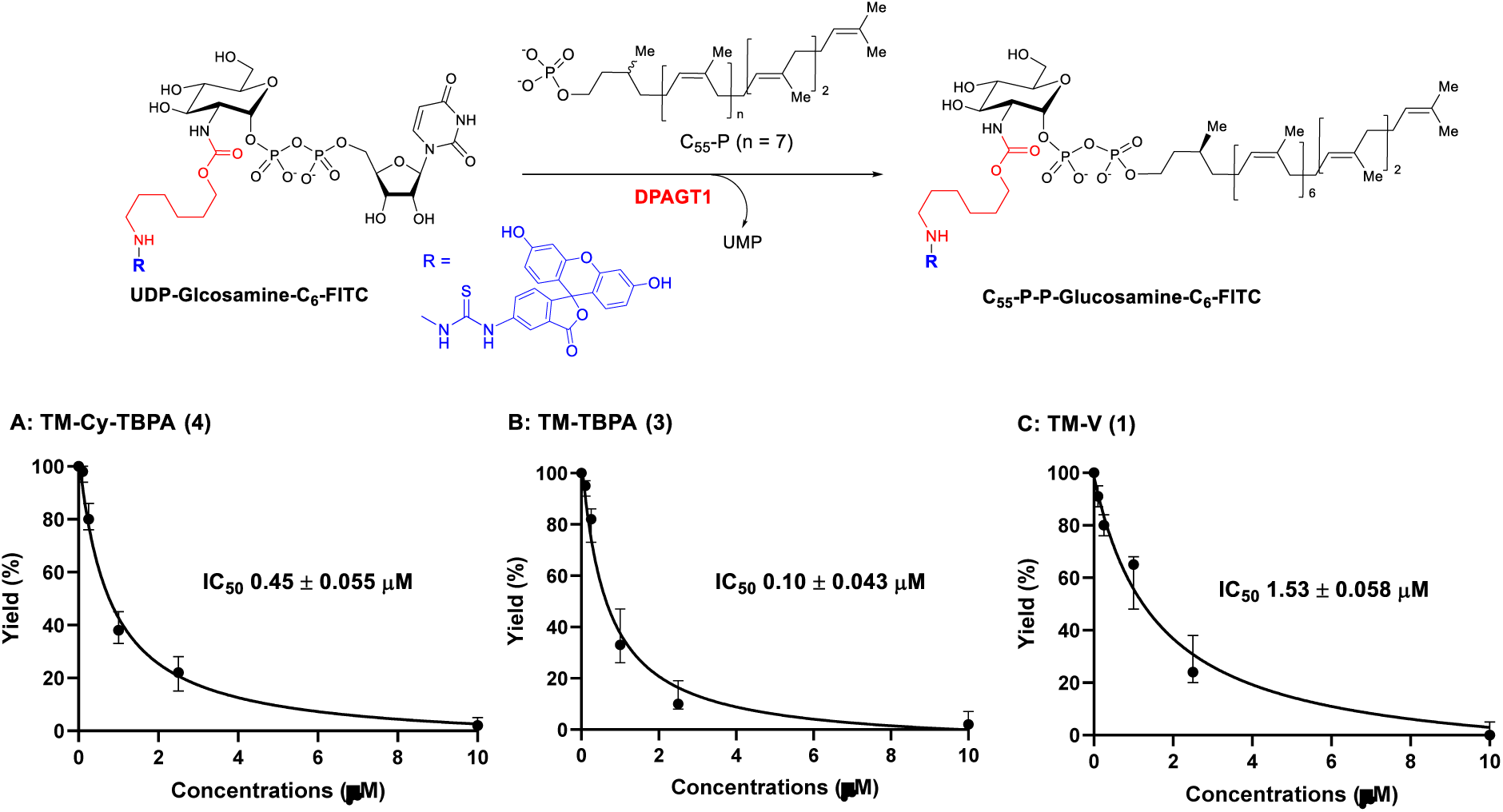
Dose–response (IC_50_) curves of TM-Cy-TBPA (**4**) compared with TM-TBPA (**3**) and TM-V (**1**).^a^ ^a^ The generated C_55_-P-P-GlcN-C_6_-FITC at each inhibitor concentration was quantified using HPLC analysis.

Under optimized reaction conditions, the DPAGT1 inhibitory activity of TM-Cy-TBPA (**4**) was systematically evaluated in comparison with TM-TBPA (**3**) and the parental natural product TM-V (**1**). Dose–response (IC_50_) curves derived from the DPAGT1 enzyme inhibition assays are summarized in Figure 6. The cyclitol analogue **4** maintained potent inhibitory activity toward DPAGT1 and exhibited inhibitory potency comparable to that of TM-TBPA (**3**), being only approximately 0.22-fold less active. Importantly, compound **4** still demonstrated approximately 3.4-fold greater inhibitory activity than TM-V (**1**), indicating that replacement of the native tunicamine core (the central amino sugar framework in Figure 1) with a cyclitol framework is well tolerated for DPAGT1 recognition and inhibition. These findings provide important mechanistic and medicinal chemistry insights into the structural requirements for selective DPAGT1 inhibition. Especially, TM-Cy-TBPA (**4**) lacks the native tunicamine framework characteristic of tunicamycin analogues and instead incorporates a conformationally constrained cyclitol surrogate in place of the tunicamine moiety. The retention of strong DPAGT1 inhibitory activity despite replacement of this structurally complex and chemically labile portion provides compelling evidence that the native tunicamine core is not an absolute structural requirement for potent DPAGT1 inhibition. Rather, these results suggest that productive DPAGT1 binding can be maintained through retention of the overall spatial arrangement of critical pharmacophoric elements using alternative scaffold architectures. This finding substantially expands the accessible chemical space for the development of tunicamycin-inspired inhibitors and provides a new structural template for the design of chemically stable DPAGT1-targeted agents. Together, these findings establish a rational framework for the further optimization of chemically and metabolically stable DPAGT1 inhibitors targeting *N*-glycan biosynthesis.

It is also noteworthy that TM-Cy-TBPA (**4**) did not exhibit detectable inhibitory activity against the bacterial lipid phosphotransferases MraY and WecA, despite the known susceptibility of these enzymes to structurally related nucleoside natural products (*e.g*., muraymycins).^3,6^ Consistent with this selectivity profile, compound **4** did not display antibacterial activity against TM-V–susceptible bacterial strains, including *Mycobacterium smegmatis*, *Bacillus cereus*, and *Bacillus subtilis*. Furthermore, no antifungal activity was observed against representative fungal pathogens, including *Candida albicans*, *Cryptococcus neoformans*, and *Aspergillus fumigatus* (see Supporting Information). These observations suggest that replacement of the native tunicamine moiety substantially altered biological specificity, resulting in selective inhibition of mammalian DPAGT1 while minimizing off-target activities against microbial phosphotransferases. Such selectivity may represent a significant therapeutic advantage for the development of DPAGT1-targeted anticancer agents by reducing undesirable antimicrobial effects and potentially improving tolerability.

**Table 1.** Cell viability of breast cancer cells following treatment with TM-Cy-TBPA (4) in comparison with TM-TBPA (3) and TM-V (1).^a^.

| Compound<sup>[b]</sup> | EC<sub>50</sub> (μM) |  |  |  |  |  |  |
| --- | --- | --- | --- | --- | --- | --- | --- |
|  | MCF-7<sup>[c]</sup> | BT-474<sup>[d]</sup> | BT-474 Clone 5<sup>[e]</sup> | SKBr-3<sup>[f]</sup> | MDA-MB-468<sup>[g]</sup> | MDA-MB-231<sup>[h]</sup> | MDA-MB-453<sup>[i]</sup> |
| **TM-Cy-TBPA (4)** | **0.69 ± 0.008** | **0.48 ± 0.033** | **0.54 ± 0.047** | **0.63 ± 0.057** | **0.65 ± 0.025** | **0.69 ± 0.055** | **0.58 ± 0.055** |
| **TM-TBPA (3)** | 0.35 ± 0.043 | 0.29 ± 0.016 | 0.30 ± 0.017 | 0.35 ± 0.082 | 0.43 ± 0.085 | 0.55 ± 0.094 | 0.45 ± 0.059 |
| **TM-V (1)** | 0.22 ± 0.020 | 0.15 ± 0.012 | 0.14 ± 0.045 | 0.10 ± 0.008 | 0.15 ± 0.012 | 0.17 ± 0.027 | 0.16 ± 0.009 |

<sup>a</sup> The IC<sub>50</sub> value was determined via WST-1 assays (see Supporting Information). [b] A synthetic product (>98%). [c] Metastatic breast adenocarcinoma (ER<sup>+</sup>/PR<sup>+</sup>/HER2<sup>-</sup>). [d] Human breast carcinoma (ER<sup>+</sup>/PR<sup>+</sup>/HER2<sup>+</sup>). [e] A subclone of BT-474 ductal carcinoma of the breast (ER<sup>+</sup>/PR<sup>+</sup>/HER2<sup>++</sup>). [f] Human breast adenocarcinoma cell (ER<sup>-</sup>/PR<sup>-</sup>/HER2<sup>+</sup>). [g] Human breast adenocarcinoma, a triple-negative breast cancer (TNBC) (ER<sup>-</sup>/PR<sup>-</sup>/HER2<sup>-</sup>). [h] Metastatic breast adenocarcinoma, a triple-negative breast cancer (TNBC) (ER<sup>-</sup>/PR<sup>-</sup>/HER2<sup>-</sup>). [i] Human breast carcinoma cell (ER<sup>-</sup>/PR<sup>-</sup>/HER2<sup>+</sup>). \**p* < 0.05 (*n* = 3).

#### 3.3. Antiproliferative Activity

The antiproliferative activities of DPAGT1 inhibitors have been reported previously, demonstrating potent anticancer activity across multiple DPAGT1-overexpressing cancer cell lines, with EC₅₀ values ranging from 0.15 to 1.25 μM.^6,9,10^ In the present study, the antiproliferative activity of TM-Cy-TBPA (**4**) was evaluated against a panel of breast cancer cell lines, and compared with those of TM-TBPA (**3**) and TM-V (**1**) (Table 1). TM-Cy-TBPA (**4**) exhibited antiproliferative activity against a panel of breast cancer cell lines, including the triple-negative subtypes MDA-MB-231 and MDA-MB-468, with EC₅₀ values ranging from 0.48 to 0.69 μM. TM-TBPA (**3**) demonstrated greater antiproliferative potency across all cancer cell lines tested relative to TM-Cy-TBPA (**4**), consistent with its stronger DPAGT1 inhibitory activity (Figure 6). We concluded that the cytotoxicity of TM-V (**1**) involves mechanisms beyond DPAGT1 inhibition, as it exhibited toxicity toward all mammalian cells at relatively similar concentrations.^8,13^ Consistent with this observation, TM-V (**1**) displayed strong cytotoxic effects against all breast cancer cell lines tested (IC₅₀ = 0.10–0.22 μM), despite possessing 15-fold and 3.4-fold weaker DPAGT1 inhibitory activity than TM-TBPA (**3**) and TM-Cy-TBPA (**4**), respectively. These data clearly demonstrate that the cyclitol core structure lacking the 8’- and 9’-hydroxy groups serves as a chemically stable tunicamine surrogate and exhibits antiproliferative activity that correlates with its DPAGT1 inhibitory activity.

#### 3.4. Evaluation of Off-Target Activities Associated with Membrane Disruption

The most critical consideration in the development of tunicamycin analogues for *in vivo* applications is ensuring that the designed molecules lack membrane-disrupting activity, enhancing the therapeutic index through reduced nonspecific toxicity. We previously demonstrated that TM-V (**1**) induces red blood cell (RBC) lysis in both a concentration- and time-dependent manner, with an IC₅₀ of approximately 5–10 μM after 24 h of incubation.^13^ To further assess TM-V-induced morphological alterations and membrane damage, we examined RBC morphology by a conventional microscopy following treatment with TM-Cy-TBPA and compared the results with those obtained for TM-V (Figure 7). Consistent with the pronounced hemolytic activity of TM-V, eosin staining revealed concentration-dependent morphological disruption of erythrocytes. At 1.0 μM, approximately 20% of RBCs had already lost their characteristic biconcave morphology and exhibited marked structural deformation. Eosin staining became diffuse and delocalized throughout the cells, accompanied by the appearance of numerous optically clear spherical regions within the cytoplasm, suggesting early membrane destabilization and redistribution of intracellular contents. In many cells, a dense dark inclusion remained localized near one pole of the erythrocyte. Upon increasing the TM-V concentration to 2.5 μM, these dark inclusions were no longer detectable, while essentially all erythrocytes displayed severe morphological distortion. At 10 μM, the erythrocytes had completely lost their native architecture, exhibiting a uniformly dense eosinophilic appearance throughout the cell interior, consistent with extensive membrane disruption and terminal structural collapse. These morphological changes closely paralleled the concentration-dependent hemolysis observed in the quantitative assay, providing direct microscopic evidence that TM-V rapidly compromises erythrocyte membrane integrity rather than eliciting a specific intracellular biochemical response (Figure 7B and 7C).

**Figure 7.**
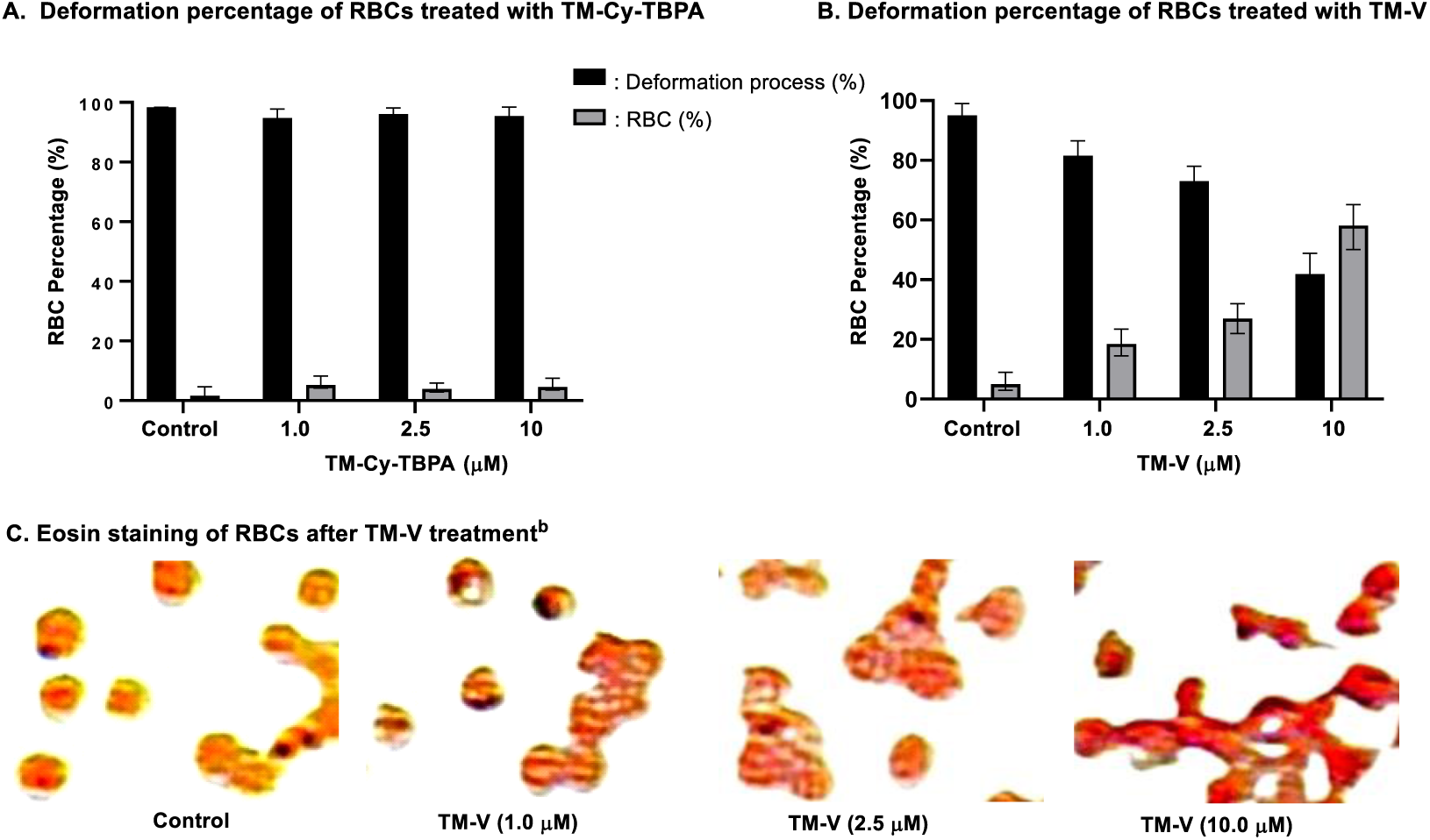
Morphological assessment of red blood cells (RBCs) in response to TM-V (**1**) and TM-Cy-TBPA (**4**).^a^ ^a^ Sheep blood cells (MP Biomedicals) are applied (24 h). Error bars represent the standard deviation of the mean in the dose response curve. The results were analyzed using a two-way ANOVA with post-hoc Dunnett’s test.; ^b^ Images were acquired at ×100 magnification (an EVOS microscope) and digitally enlarged by 480% for clarity. *P* <0.05.

The progression from localized structural abnormalities to complete loss of cellular architecture further supports membrane disruption as the primary mechanism underlying the acute cytotoxicity of TM-V. In sharp contrast, TM-Cy-TBPA (**4**) did not induce detectable heme leakage from erythrocytes or appreciable deformation of RBC morphology, even at concentrations of 10 μM or higher (Figure 6A and Supporting Information). These results indicate that cyclitol- and TBPA-based structural modification effectively suppresses the membrane-disruptive liability observed with TM-V.

To determine whether the absence of membrane-disruptive activity observed in erythrocytes extended to nucleated cells, we evaluated plasma membrane integrity in MCF-7 and MDA-MB-231 breast cancer cells following treatment with TM-Cy-TBPA and compared the results with TM-V (Figure 8 and Figure 9).

**Figure 8.**
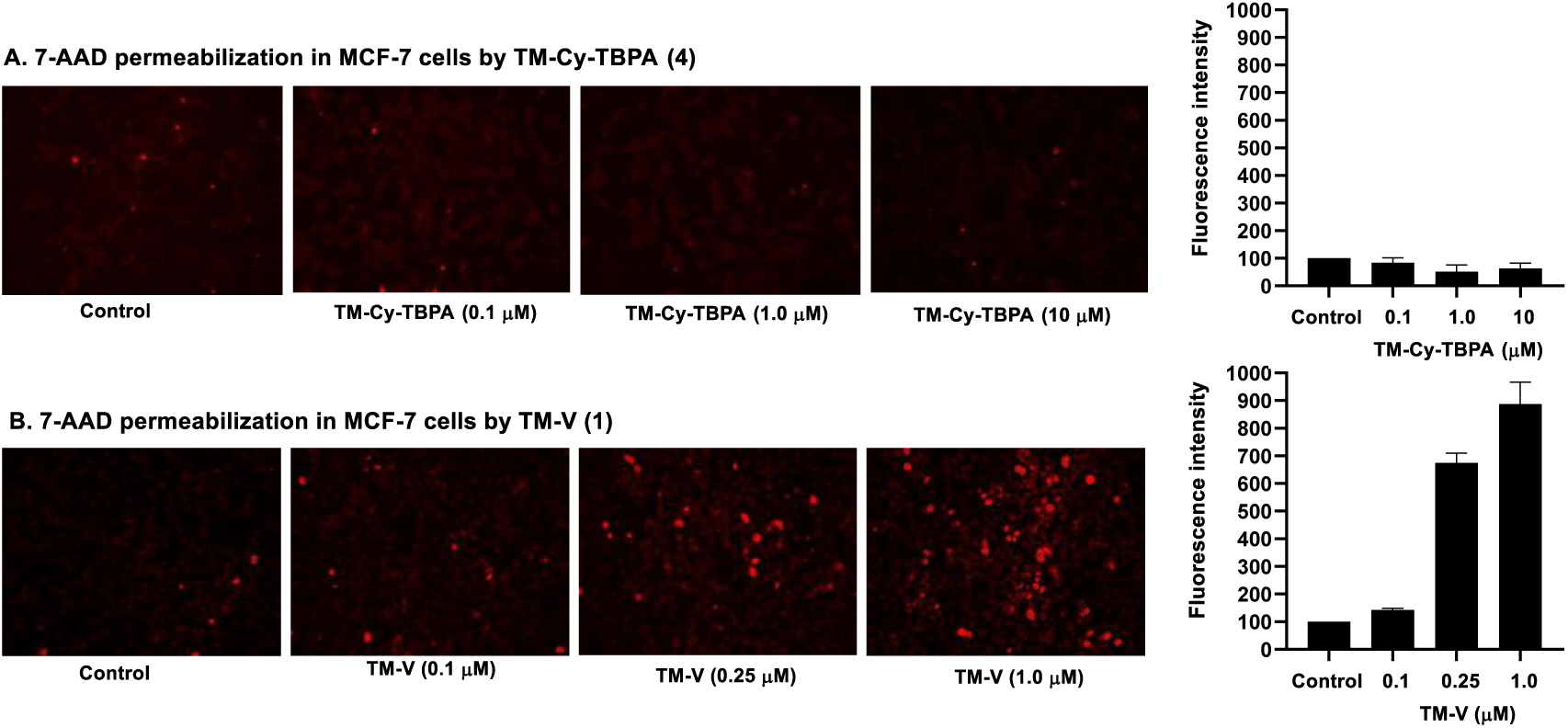
A membrane-permeation assay using 7-aminoactinomycin D (7-AAD) in the presence of TM-V(**1**) or TM-Cy-TBPA (**4**)^a^ ^a^ 7-AAD is a membrane-impermeable fluorescent DNA intercalator. TM-V induces loss of membrane integrity at 0.25 µM after 6 h. The ratio of culture medium to BD 7-AAD staining solution (BD B559925) was 20:1. *P* <0.05 versus the untreated control.

**Figure 9.**
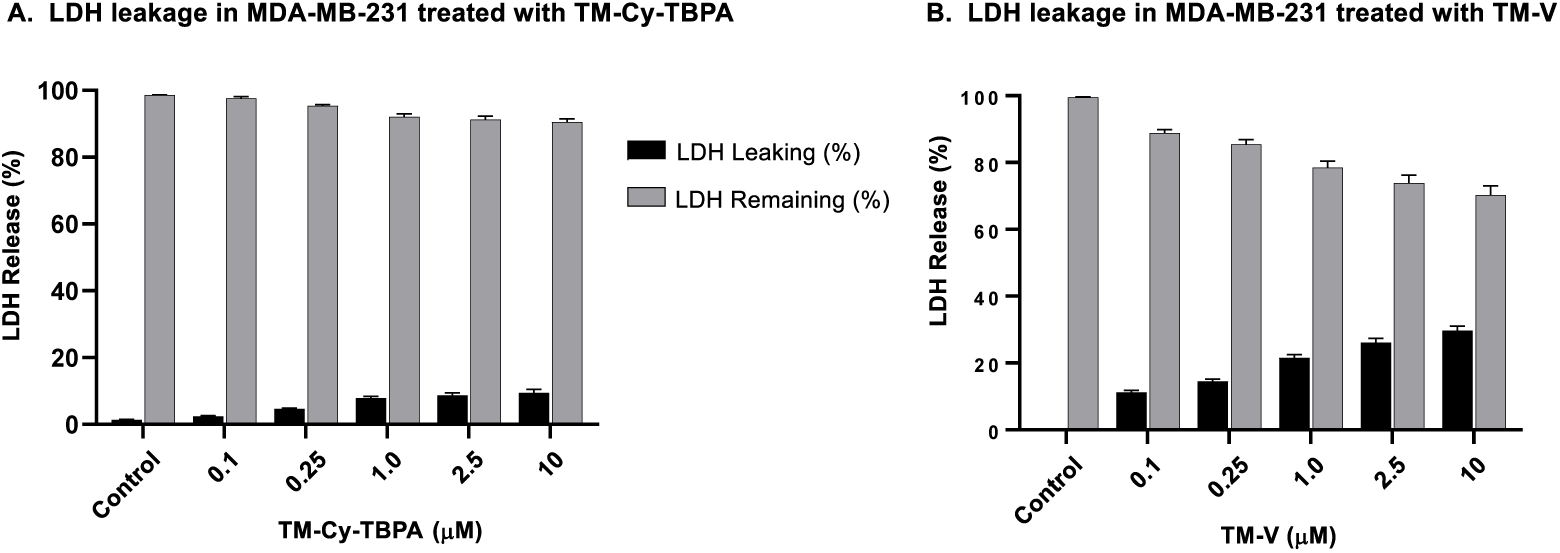
LDH release and membrane permeability assays of TM-Cy-TBPA (**4**) compared with TM-V (**1**).^a^ ^a^ Membrane permeability was evaluated using the WST-8 assay by quantifying WST-8 formazan formation (OD460) in a coupling format (Tribioscience). Cells were treated for 6 h. Lactate dehydrogenase (LDH) release was normalized to complete cell lysis induced by 1% Triton X-100, which was defined as 100% release. Released and remaining LDH values were determined independently by fluorescence and therefore may not sum to exactly 100% because of normal assay variability. Data are presented as mean ± SD (*n* = 3 independent experiments). *P* < 0.05 versus the untreated control.

A membrane-disruption assay using the cell-impermeable fluorescent dye 7-AAD (7-aminoactinomycin D) to visualize loss of plasma membrane integrity has routinely performed in our laboratory.^13^ During the 6 h assay, MCF-7 human breast cancer cells retained normal morphology and viability when exposed to TM-Cy-TBPA at concentrations 10 μM (Figure 8A). In contrast, beginning at 0.25 μM, TM-V (**1**) treatment produced a clear and concentration-dependent increase in intracellular 7-AAD fluorescence, indicating compromised membrane integrity and dye entry into the nucleus (Figure 8B). The onset and intensity of 7-AAD influx closely parallel the cytotoxicity profile of TM-V against MCF-7 (IC_50_ values in Table 1), suggesting that membrane perturbation contributes to its cytotoxic activity. By comparison, TM-Cy-TBPA did not induce detectable 7-AAD influx.

We further confirmed the preservation of plasma membrane integrity in MDA-MB-231 cells following treatment with TM-Cy-TBPA using a lactate dehydrogenase (LDH) leakage assay (Figure 9). Extracellular (released) and intracellular (remaining) LDH activities were measured independently by fluorescence. Because these measurements are performed in separate samples and are influenced by background fluorescence and pipetting variability, the calculated percentages may not sum exactly to 100%. Small deviations from 100% were considered to reflect normal experimental variability. TM-V induced >10% LDH release at a concentration of 0.1 μM, with leakage increasing in a concentration-dependent manner to approximately 30% after 6 h at 10 μM. In sharp contrast, TM-Cy-TBPA caused only minimal LDH leakage (<5.5%) even at 10 μM, a level close to the experimental background and indicative of negligible plasma membrane damage. These results demonstrate that, unlike TM-V, TM-Cy-TBPA does not compromise plasma membrane integrity, as demonstrated by its minimal hemolytic activity toward red blood cells and negligible 7-AAD uptake in cellular permeability assays. These findings indicate that TM-Cy-TBPA retains membrane integrity and does not induce nonspecific membrane disruption, consistent with its selective mechanism of action.

#### 3.5. Evaluation of the Cytotoxicity of TM-Cy-TBPA (4) against Normal Mammalian Cells

One of the major cytotoxic mechanisms of TM-V (**1**) is disruption of mammalian cell membranes, resulting in rapid and nonspecific cellular toxicity. As described above, TM-Cy-TBPA (**4**) did not exhibit appreciable membrane-disruptive activity at concentrations of 10 μM or higher in three independent assays, indicating that replacement of the native tunicamine core with the cyclitol scaffold, together with incorporation of the lipid mimetic moiety, substantially reduces this undesirable property. These findings suggested that the antiproliferative activity of TM-Cy-TBPA (**4**) may arise predominantly from selective DPAGT1 inhibition rather than from nonspecific membrane damage. To further evaluate its safety profile, the cytotoxicity of TM-Cy-TBPA (**4**) was examined in comparison with TM-TBPA (**3**) and TM-V (**1**) using a panel of nontransformed (non-cancerous) mammalian cell lines. Consistent with observations reported for muraymycin A1 and its simplified analogues,^5,6^ a major objective of this study was to determine whether tunicamycin analogues can effectively dissociate selective DPAGT1 inhibition from off-target, nonspecific cytotoxicity. Cytotoxicity assays were therefore conducted at concentrations up to 100 μM to evaluate the therapeutic window and selectivity of the cyclitol analogue (Table 2). TM-TBPA (**3**) exhibited no detectable toxicity toward the healthy cell lines listed in Table 2 at concentrations up to 100 μM. Similarly, TM-Cy-TBPA (**4**) showed no detectable toxicity against any of the normal cell lines tested for TM-TBPA (**3**). In contrast, TM-V (**1**) exhibited pronounced cytotoxicity toward all normal cell lines in Table 2, with toxic effects observed at concentrations ranging from 0.25 to 0.53 μM. The marked difference in toxicity profiles between TM-V (**1**) and the engineered analogues strongly suggests that structural modification of the tunicamycin scaffold can effectively dissociate DPAGT1 inhibition from the severe nonselective cytotoxicity associated with the parent natural product. These findings are consistent with the membrane integrity assays described above, in which TM-Cy-TBPA (**4**) did not exhibit detectable membrane-disruptive activity even at substantially higher concentrations. The absence of measurable toxicity toward healthy mammalian cells, together with its previously demonstrated antiproliferative activity against cancer cell lines, indicates that TM-Cy-TBPA (**4**) possesses a substantially improved therapeutic index relative to TM-V (**1**).

**Table 2.** Cell viability of nontransformed (non-cancerous) cell lines following treatment with TM-Cy-TBPA (4) and TM-TBPA (3) in comparison with TM-V (1).^a^.

| Compound | IC<sub>50</sub> (μM) |  |  |  |  |  |  |
| --- | --- | --- | --- | --- | --- | --- | --- |
|  | MCF-10A<sup>b</sup> | Vero<sup>c</sup> | COS-7<sup>d</sup> | BRL-3A<sup>e</sup> | PBMC<sup>f</sup> | AC16<sup>g</sup> | HEK<sup>h</sup> |
| **TM-Cy-TBPA (4)** | >100 | >100 | >100 | >100 | >100 | >100 | >100 |
| **TM-TBPA (3)** | >100 | >100 | >100 | >100 | >100 | >100 | >100 |
| **TM-V (1)** | 0.25 ± 0.003 | 0.45 ± 0.005 | 0.27 ± 0.012 | 0.36 ± 0.019 | 0.25 ± 0.004 | 0.26 ± 0.019 | 0.53 ± 0.039 |

<sup>a</sup> The IC<sub>50</sub> value was determined via WST-1 assays.; <sup>b</sup> Non-tumorigenic breast epithelial cell.; <sup>c</sup> Kidney epithelial cells from African green monkey.; <sup>d</sup> Fibroblast-like cells derived from monkey kidney tissue.; <sup>e</sup> Fibroblast-like cells isolated from the liver of a rat.; <sup>f</sup> Primary peripheral blood mononuclear cells.; <sup>g</sup> A human cardiomyocyte cell line.; <sup>h</sup> Human Epidermal Keratinocytes. \**p* < 0.05 (*n* = 3).

#### 3.6. Mechanistic Analysis of Cell Cycle Arrest and Apoptosis Induction in Breast Cancer Cells Treated with TM-Cy-TBPA (4)

Previous studies from our laboratory demonstrated that selective DPAGT1 inhibitors induce G_2_/M cell-cycle arrest in breast cancer cells, followed by apoptosis triggered by persistent endoplasmic reticulum (ER) stress and activation of the unfolded protein response (UPR).^6,13^ This antiproliferative mechanism is fundamentally distinct from that of TM-V, which induces G_0_/G_1_ arrest and triggers apoptosis together with other regulated cell death pathways, including necroptosis and pyroptosis.^10,45,46^ When ER stress persists and adaptive UPR signaling fails to restore ER proteostasis, the intrinsic mitochondrial apoptotic pathway is activated.^47,48^ This process is characterized by mitochondrial outer membrane permeabilization, release of cytochrome *c* into the cytosol, activation of executioner caspases, and ultimately irreversible apoptotic cell death. p53 and caspase-3 are key mediators that couple sustained ER stress to activation and execution of the intrinsic mitochondrial apoptotic program.^46, 49, 50^ In our previous studies using MDA-MB-231 cells, mechanistic interpretation of p53 signaling was limited because these cells constitutively overexpress a gain-of-function mutant p53 that lacks the canonical tumor-suppressive functions of wild-type p53. ^51, 52^ Consequently, MDA-MB-231 cells are not an ideal model for defining a p53-dependent mechanism linking G_2_-phase arrest to mitochondrial apoptosis. Furthermore, conventional flow cytometric analysis of cell-cycle distribution alone cannot distinguish G_2_-phase arrest from mitotic (M) arrest and therefore provides insufficient mechanistic insight into G_2_/M checkpoint regulation or the selective anticancer activity of DPAGT1 inhibitors.^53,54^ These limitations underscore the need for complementary analyses of cell-cycle regulatory proteins together with ER stress- and apoptosis-associated signaling pathways to establish the molecular mechanism underlying DPAGT1 inhibitor-induced cancer cell death.^55,56^

Western blot analyses of cell-cycle regulatory proteins (Cyclin B1, CDK1/CDK2, phospho-CDC2/CDK1 (Tyr15), Wee1, p53, p21, and phospho-histone H3) in SKBr-3 cells following treatment with TM-Cy-TBPA (**4**) are summarized in Figure 10. Consistent with G₂/M cell-cycle arrest, Cyclin B1 (the essential regulatory partner of CDK1 that drives the G_2_/M transition) accumulated in a concentration-dependent manner. Similar to MDA-MB-231 cells, CDK1 (CDC2) (cyclin-dependent kinase 1, the principal kinase that controls the G_2_/M transition) was highly expressed in SKBr-3 cells,^57^ and its total expression level remained unchanged following TM-Cy-TBPA treatment, whereas phosphorylation of CDK1 (CDC2) at Tyr15 increased. TM-Cy-TBPA also increased Wee1 (a G_2_/M checkpoint kinase that negatively regulates CDK1 through inhibitory phosphorylation at Tyr15) expression, even at a concentration of 0.1 μM. Collectively, these findings indicate that TM-Cy-TBPA maintains CDK1 in its inactive (Tyr15-phosphorylated state), therefore preventing activation of the Cyclin B1–CDK1 complex and blocking entry into mitosis (M phase). The basal p53 expression level in SKBr-3 cells was substantially lower (approximately 7.2-fold lower) than that in MDA-MB-231 cells. TM-Cy-TBPA induced a concentration-dependent increase in p53 expression, which was accompanied by a parallel increase in p21 expression, consistent with activation of the p53–p21 signaling pathway.^58,59^ Phospho-histone H3, a well-established marker of mitotic entry,^60^ was markedly decreased in a concentration-dependent manner. This finding indicates that TM-Cy-TBPA arrests cells in the G_2_ phase rather than the mitotic (M) phase of the cell cycle. Consistent with these findings, TM-Cy-TBPA also induced G2-phase arrest in MDA-MB-231 cells through a p53-independent mechanism, demonstrating that this cell-cycle arrest is conserved across both breast cancer cell lines despite the presence of different p53 mutations.^61,62^ (see Supporting Information).

**Figure 10.**
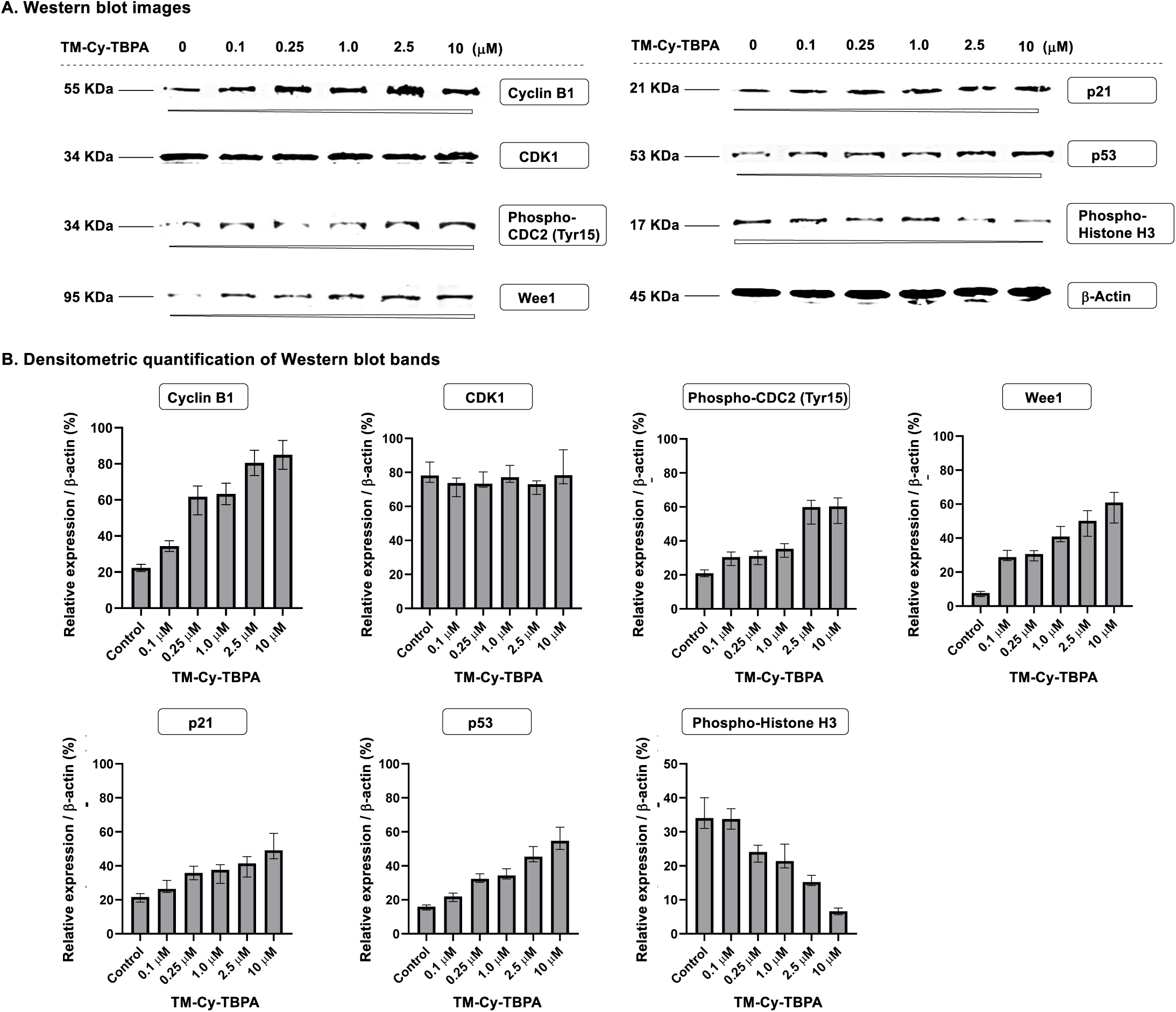
Western blot analysis of cell-cycle regulatory proteins in SKBr-3 cells following treatment with TM-Cy-TBPA (**4**). β-Actin was used as the protein loading control. Wee1 (D10D2) Rabbit Monoclonal Antibody (Cell Signaling Technology, Cat. #13084), Cyclin B1 (D5C10) Rabbit Monoclonal Antibody (Cell Signaling Technology, Cat. #12231), p53 (7F5) Rabbit Monoclonal Antibody (Cell Signaling Technology, Cat. #2527), Anti-CDK1 antibody [1/Cdk1/Cdc2] (abcam, Cat. #ab280964), p21 Waf1/Cip1 (12D1) Rabbit Monoclonal Antibody (Cell Signaling Technology, Cat. #2947), Phospho-Histone H3 (Ser10) (D2C8) Rabbit Monoclonal Antibody (Cell Signaling Technology, Cat. #3377), Phospho-cdc2 (Tyr15) (10A11) Rabbit Monoclonal Antibody (Cell Signaling Technology, Cat. #4539), and beta-Actin (E4D9Z) Mouse Monoclonal Antibody (Cell Signaling Technology, Cat. #58169) were used as primary antibody. IRDye 680RD Goat anti-Rabbit IgG Secondary Antibody (LI-COR Biotech, Cat. #926-68071) and IRDye 800CW Goat anti-Mouse IgG Secondary Antibody (LI-COR Biotech, Cat. #926-32210) were used as secondary antibody. LI-COR Biotech ODYSSEY DLX was used to scan the probe signal. The data was quantified with ImageJ1.54d (NIH) processing software. (*p*<0.05, *n* = 3,)

To elucidate the mechanism by which TM-Cy-TBPA-induced G_2_-phase arrest progresses to apoptotic cell death, we examined key molecular markers of ER stress, the unfolded protein response (UPR), and the intrinsic mitochondrial apoptotic pathway in SKBr-3 cells (Figure 11), including BiP/GRP78, XBP1s, CHOP, Bcl-2, cytochrome *c* release, cleaved caspase-3/7 (c-caspase-3/7), and cleaved PARP (c-PARP).^63,64^ TM-Cy-TBPA elicited a robust ER stress response, as evidenced by the upregulation of BiP/GRP78 (the ER chaperone that maintains protein folding homeostasis).^13^ The elevation of XBP1s (spliced X-box binding protein 1) indicates activation of the IRE1α branch of the UPR, whereas CHOP induction reflects activation of the pro-apoptotic PERK–ATF4 signaling pathway.^65,66,67^ Persistent ER stress subsequently triggered intrinsic mitochondrial apoptosis, as demonstrated by downregulation of the anti-apoptotic protein Bcl-2, promoting mitochondrial outer membrane permeabilization and the release of cytochrome *c* into the cytosol.^68^ This mitochondrial dysfunction activated the executioner caspase cascade, as evidenced by increased levels of cleaved caspase-3/7 and cleaved PARP (a DNA repair enzyme that is cleaved by activated caspase-3/7),^69,70^ confirming that TM-Cy-TBPA induces intrinsic apoptosis downstream of DPAGT1 inhibition. Similar results were obtained in MDA-MB-231 cells, demonstrating that the same mechanistic pathway operates in both breast cancer cell lines (see Supporting Information).

**Figure 11.**
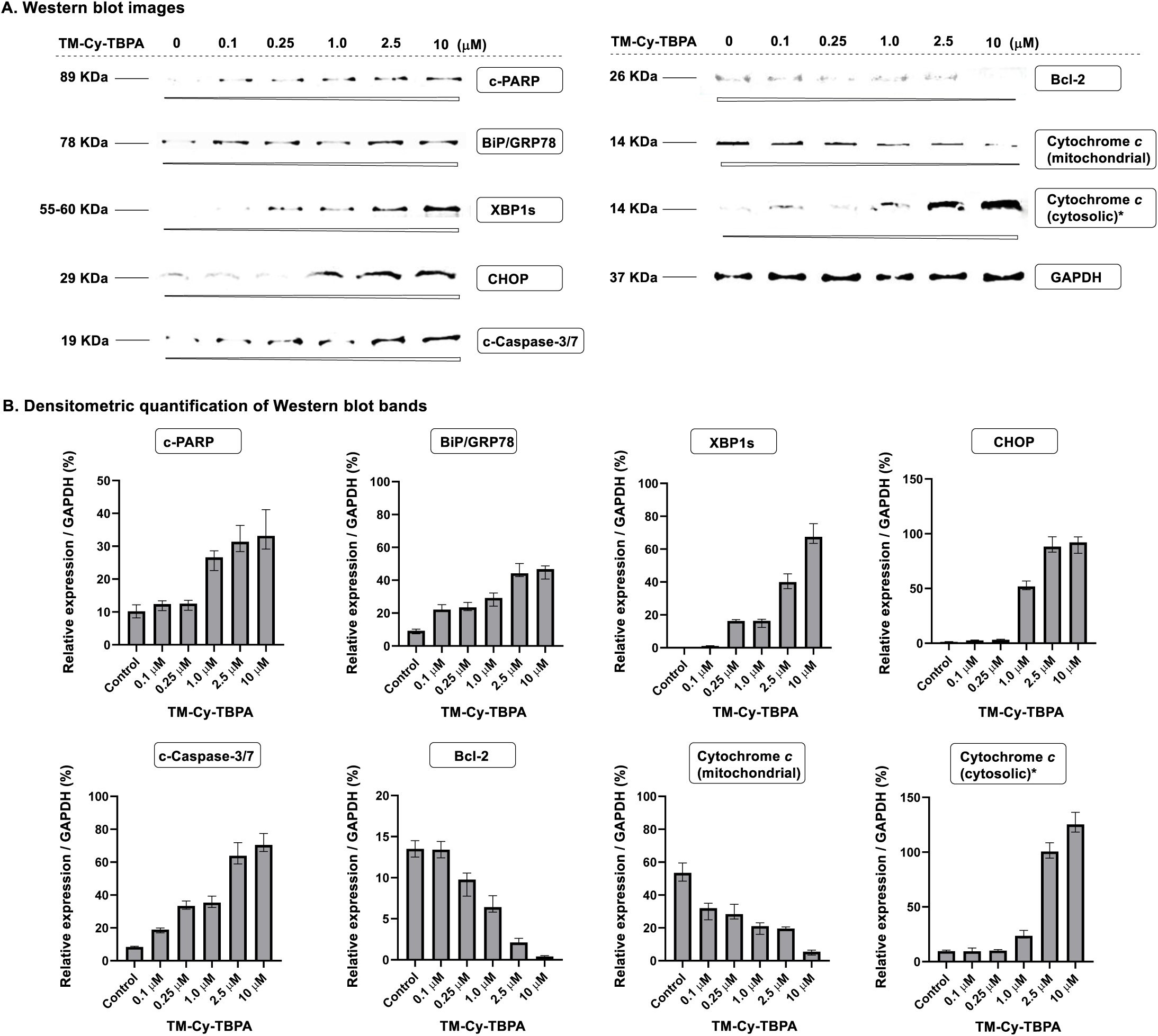
Western blot analysis of apoptosis regulatory proteins in SKBr-3 cells following treatment with TM-Cy-TBPA (**4**). GAPDH was used as the protein loading control. *To improve detection of treatment effects at lower concentrations of compound **4**, the amount of total protein loaded for Western blot analysis was increased four-fold. Cleaved PARP (Asp214) (D64E10) Rabbit Monoclonal Antibody (Cell Signaling Technology, Cat. #5625), CHOP (L63F7) Mouse Monoclonal Antibody (Cell Signaling Technoloty, Cat. #2895), BiP (C50B12) Rabbit Monoclonal Antibody (Cell Signaling Technoloty, Cat. #3177), XBP-1s (E9V3E) Rabbit Monoclonal Antibody (Cell Signaling Technoloty, Cat. #40435), Bcl-2 Recombinant Rabbit Monoclonal Antibody (JE10-17) (Invitrogen, Cat. #MA5-36172), Cleaved Caspase-3 (Asp175) Antibody (Cell Signaling Technoloty, Cat. #9661), Anti-Cytochrome c mouse mAb (Cytochrome c Releasing Apoptosis Assay Kit, BioVision, Cat. #K257-100), and GAPDH (D4C6R) Mouse Monoclonal Antibody (Cell Signaling Technoloty, Cat. #97166) were used as primary antibody. IRDye 680RD Goat anti-Rabbit IgG Secondary Antibody (LI-COR Biotech, Cat. #926-68071) and IRDye 800CW Goat anti-Mouse IgG Secondary Antibody (LI-COR Biotech, Cat. #926-32210) were used as secondary antibody. LI-COR Biotech ODYSSEY DLX was used to scan the probe signal. The data was quantified with ImageJ1.54d (NIH) processing software. (*p*<0.05, *n* = 3)

### 4. Pharmacokinetic Evaluation of the Cyclitol Analogue TM-Cy-TBPA (4) in Comparison with TM-TBPA (3)

Replacement of the native tunicamine core with the cyclitol scaffold markedly improved the chemical stability of TM-Cy-TBPA, consistent with elimination of the acid-labile glycosidic linkage present in conventional tunicamycin analogues. This scaffold-stabilization strategy was designed to enhance the pharmacological suitability of DPAGT1 inhibitors by improving molecular robustness under physiologically relevant conditions while preserving the key structural features required for productive target engagement. To establish an initial proof-of-pharmacological concept for tunicamycin-derived DPAGT1 inhibitors as anticancer agents *in vivo*, the pharmacokinetic (PK) properties of TM-Cy-TBPA were evaluated and directly compared with those of TM-TBPA. Particular emphasis was placed on determining whether incorporation of the cyclitol framework could retain systemic exposure and overall drug-like behavior without compromising the favorable biological activity. These studies were intended to assess the translational feasibility of the cyclitol analogue as a chemically stable and pharmacologically tractable scaffold for further preclinical development.

Both compounds were administered to CD-1 mice (n = 3) either intravenously (i.v., 10 mg/kg) or orally (p.o., 30 mg/kg). Following a single administration, plasma concentrations were quantified at multiple time points using LC–MS analysis. Pharmacokinetic parameters, including the area under the plasma concentration–time curve (*AUC*), apparent volume of distribution (*V*_z_), systemic clearance (*CL*), and terminal elimination half-life (*t*_1/2_) were determined using noncompartmental analysis.^71,72^

Following oral administration, oralbioavailability (*F*) was calculated, and the oral exposure-to-potency index (*AUC_inf_*_(p.o.)_ / EC_50_) was determined to assess systemic exposure relative to antiproliferative activity.^73^ To our surprise, despite the unfavorable **chemical stability associated with the native tunicamine core structure (Figure 5), TM-**TBPA (**3**) nevertheless exhibited a promising pharmacokinetic profile in mice. TM-Cy-TBPA (**4**) also displayed promising pharmacokinetic characteristics despite achieving relatively comparable systemic exposure levels (Table 3). TM-TBPA demonstrated moderately higher systemic exposure than TM-Cy-TBPA, with an *AUC* of 28.7 versus 21.0, respectively. In parallel, TM-TBPA showed lower systemic clearance (*CL* = 5.8) and a substantially smaller apparent volume of distribution (*V*_z_ = 1.3) relative to TM-Cy-TBPA (*CL* = 8.6; *V*_z_ = 4.5). The lower *V*_z_ of TM-TBPA suggests that the compound remains more confined to the vascular/plasma compartment, whereas TM-Cy-TBPA distributes more extensively into peripheral tissues.^69^ Notably, despite its lower *AUC* and higher clearance, TM-Cy-TBPA exhibited a substantially prolonged terminal elimination half-life (*t*_1/2_ = 6.3 h) compared with TM-TBPA (*t*_1/2_ = 2.5 h). This extended half-life is consistent with the markedly larger volume of distribution observed for TM-Cy-TBPA, suggesting sustained tissue distribution and slower terminal redistribution/elimination kinetics. Following oral administration, TM-TBPA demonstrated superior oral exposure, with higher *AUC*, bioavailability, and *AUC_inf_*_(p.o.)_ / EC₅₀ values than TM-Cy-TBPA. *In vitro* Caco-2 permeability assays demonstrated that both TM-TBPA and TM-Cy-TBPA exhibited minimal transporter-mediated efflux liabilities, with efflux ratios of 0.6 and 1.1, respectively. In mouse plasma, TM-Cy-TBPA displayed higher plasma protein binding than TM-TBPA (87.3% vs. 72.3%), whereas the opposite trend was observed in human plasma, where TM-Cy-TBPA exhibited slightly lower protein binding (68.5% vs. 74.3%). In mouse pharmacokinetic studies, both compounds achieved promising systemic exposure following i.v. administration. However, despite the more favorable *in vitro* permeability profile of TM-Cy-TBPA, oral PK studies revealed lower oral bioavailability for TM-Cy-TBPA relative to TM-TBPA (0.86% vs. 1.86%) (see Supporting Information). These observations underscore the complexity of predicting the pharmacokinetic behavior of tunicamycin analogues solely from conventional *in vitro* ADME parameters. ^74^ Nevertheless, TM-Cy-TBPA achieved measurable systemic exposure following oral administration while maintaining a favorable relationship between systemic exposure and antiproliferative potency. Together with its prolonged terminal half-life following intravenous administration, these findings establish TM-Cy-TBPA as a promising lead scaffold for further pharmacokinetic and medicinal chemistry optimization.

**Table 3.** Comparative pharmacokinetic (PK) profiles of TM-Cy-TBPA (4) and TM-TBPA (3).

| Compound | Intravenous (i.v.) administration<sup>a</sup> |  |  |  | Oral (p.o.) administration<sup>b</sup> |  |  |
| --- | --- | --- | --- | --- | --- | --- | --- |
| | *AUC<sub>inf</sub>*<br>($\mu\text{g}\cdot\text{hr}/\text{mL}$)<sup>c</sup> | *V<sub>z</sub>*<br>(L/kg)<sup>d</sup> | *CL*<br>(mL/kg·min)<sup>e</sup> | *t<sub>1/2</sub>*<br>(h)<sup>f</sup> | *AUC<sub>inf</sub>*<br>($\mu\text{g}\cdot\text{hr}/\text{mL}$) | *F*<br>(%)<sup>g</sup> | *AUC<sub>inf</sub>*<br>(p.o.) / *EC<sub>50</sub>*<sup>h</sup> |
| TM-Cy-TBPA (**4**) | 21.01 | 4.54 | 8.60 | 6.30 | 0.71 | 0.86 | 1.16 |
| TM-TBPA (**3**) | 28.65 | 1.30 | 5.82 | 2.52 | 2.75 | 1.86 | 7.07 |

<sup>a</sup> Dose (i.v.) = 10 mg/kg/mouse.; <sup>b</sup> Dose (p.o.) = 30 mg/kg/mouse.; <sup>c</sup> *AUC<sub>inf</sub>* = Area under the plasma concentration–time curve extrapolated to infinity.; <sup>d</sup> *V<sub>z</sub>* = The apparent volume of distribution during the terminal phase.; <sup>e</sup> *CL* = Clearance.; <sup>f</sup> *t<sub>1/2</sub>* = Terminal elimination half-life.; <sup>g</sup> *F* = bioavailability.; <sup>h</sup> *AUC<sub>inf</sub>*(p.o.) / *EC<sub>50</sub>* = Exposure-to-potency index.

Our previously reported muraymycin A1 analogue, APPB (aminouridyl phenoxypiperidinbenzylbutanamide), demonstrated potent antitumor efficacy in both xenograft and orthotopic mouse models and exhibited favorable pharmacokinetic properties (AUC*inf* = 27.10 μg·h/mL, *V*_z_ = 2.8, CL = 6.2, and *t*_1/2_ = 6.2 h).^9,10^ Although TM-Cy-TBPA displayed moderately higher systemic clearance than APPB (8.6 vs. 6.2), its substantially larger apparent volume of distribution (*V*_z_) resulted in a longer terminal half-life while maintaining comparable systemic exposure (Table 3). These pharmacokinetic characteristics indicate that TM-Cy-TBPA undergoes extensive tissue distribution and prolonged systemic residence, properties that may facilitate sustained DPAGT1 target engagement and durable antitumor activity *in vivo*.

## CONCLUSIONS

The tunicamycin cyclitol analogue TM-Cy-TBPA was rationally designed through extensive computational studies, including conformational landscape analysis and molecular docking. These studies established that the (*1R,2R,5R*)-2-amino-5-alkylcyclohexan-1-ol scaffold serves as an effective tunicamine surrogate that replaces the central carbohydrate scaffold of tunicamycins by conformationally, stabilizing the bioactive conformation required for DPAGT1 binding and recognition. In addition, the water-soluble TBPA lipid mimetic, which contains two cationic centers, was designed to reduce the energetic barrier (ΔΔG) associated with the transition from the folded conformation favored in aqueous solution to the extended conformation required for binding within the dolichyl phosphate (Dol-P)-binding cleft of DPAGT1, therefore facilitating productive target engagement. The rationally designed TM-Cy-TBPA (**4**) was synthesized stereoselectively in an efficient 12-step sequence from commercially available starting material **12** using a key Büchner–Curtius–Schlotterbeck (BCS) reaction, providing an overall yield of 34%. TM-Cy-TBPA was comprehensively evaluated through chemical and metabolic stability studies, DPAGT1 enzyme inhibition assays, membrane-disruption assays, mammalian cell-based antiproliferative assays, mechanistic studies of cell-cycle arrest and apoptosis, and pharmacokinetic evaluation in CD-1 mice. TM-Cy-TBPA retained potent DPAGT1 inhibitory activity (IC₅₀ = 0.45 μM) while exhibiting markedly improved acid stability (*t*_1/2_ > 6 h) and a substantially prolonged biological half-life compared with TM-TBPA (**3**) (6.3 vs. 2.5 h). Moreover, incorporation of the water-soluble TBPA lipid mimetic effectively eliminated the membrane-disrupting properties associated with TM-V (**1**), as demonstrated by i) the absence of red blood cell (RBC) morphological changes and (ii) minimal membrane permeabilization and cytotoxicity in 7-AAD uptake and LDH release assays. Importantly, TM-Cy-TBPA exhibited no detectable cytotoxicity toward nontransformed mammalian cell lines (CC₅₀ ≥ 100 μM) while maintaining potent antiproliferative activity against a panel of breast cancer cell lines, including triple-negative breast cancer (TNBC), with EC₅₀ values of 0.4–0.8 μM. Consequently, TM-Cy-TBPA achieved exceptional selectivity indices (SIs) of >125–250, representing more than a 100-fold improvement over TM-V (SIs = 1.1–2.2). Collectively, these findings demonstrate that rational cyclitol and lipid-mimetic engineering transformed TM-V into a chemically stable, metabolically robust, and highly cancer cell-selective DPAGT1 inhibitor without sacrificing inhibitory activity. Here, we define the molecular mechanism of TM-Cy-TBPA-induced cell-cycle arrest in SKBr-3 and MDA-MB-231 breast cancer cells, which exhibit distinct p53 expression levels, with MDA-MB-231 expressing substantially higher levels of mutant p53 than SKBr-3. By evaluating key regulators of the G_2_/M checkpoint, we demonstrate that TM-Cy-TBPA arrests cells in the G_2_ phase rather than the mitotic (M) phase of the cell cycle. Subsequent analysis of apoptotic signaling revealed that prolonged G_2_-phase arrest is followed by activation of the mitochondria-mediated intrinsic apoptotic pathway. In preliminary pharmacokinetic studies, TM-Cy-TBPA exhibited favorable pharmacokinetic properties, including a prolonged systemic half-life and extensive tissue distribution, that compare favorably with those of our proprietary DPAGT1 inhibitor APPB, a muraymycin A1-derived lead compound with demonstrated *in vivo* antitumor efficacy. These findings establish TM-Cy-TBPA as a promising next-generation DPAGT1 inhibitor for *in vivo* evaluation. The antitumor efficacy of TM-Cy-TBPA and its advanced analogues, the cryo-EM structures of their complexes with DPAGT1, and the molecular mechanisms by which these tunicamycin cyclitol analogues selectively disrupt cancer-associated *N*-glycosylation while sparing normal cellular viability will be reported in future studies.

## Acknowledgments

This work was supported by the National Institutes of Health (NIH) under Grants R01GM114611 and R01 CA308127. NMR data were acquired using instrumentation supported by the NIH Shared Instrumentation Grant Program. Flow cytometry and cell-sorting instrumentation were supported by NIH Grant 1S10OD032329-01A1. M.K. gratefully acknowledges the U.S.–Egypt Fulbright Program for supporting his scholarly appointment at the College of Pharmacy, King Salman International University, Egypt. M.K. also thanks Mr. Bradley Morrison (Anviron, CA, USA) for valuable discussions on breast cancer drug discovery. F.S. gratefully acknowledges financial support from the Ministerio de Ciencia, Innovación y Universidades (Spain) under Grant PID2023-150577OB-I00.

## EXPERIMENTAL SECTION

### Chemistry. General Information

Unless otherwise noted, all reagents and solvents were purchased from commercial suppliers and used without further purification. THF, CH₂Cl₂, and DMF were purified using an Innovative Technology Pure-Solve solvent purification system. Reactions were performed under an argon atmosphere and monitored by TLC (silica gel 60 F254, 0.25 mm; EMD), LC–MS (Shimadzu LCMS-2020; solvent A, 0.1% formic acid in water; solvent B, acetonitrile), or both. TLC spots were visualized under UV light (254 nm) or by staining with ceric ammonium molybdate, anisaldehyde, phosphomolybdic acid, or ninhydrin, followed by heating. Flash chromatography was performed using silica gel (SiliCycle Purasil, 60 Å, 230–400 mesh). ^1^H and ^13^C NMR spectra were recorded at 400 and 100 MHz, respectively. Chemical shifts (δ) are reported in ppm relative to residual solvent signals (CDCl_3_: δH = 7.26 ppm, δC = 77.16 ppm; CD_3_CN: δH = 1.94 ppm, δC = 1.32ppm; CD_3_OD: δH =3.31 ppm, δC =49.00 ppm; DMSO-*d*_6_: δH = 2.50 ppm, δC = 39.52 ppm; D_2_O: δH = 4.79 ppm), and coupling constants (*J*) are reported in Hz. Signal multiplicities are designated as s, d, dd, t, q, quin, hept, m, and br. IR spectra were recorded on a PerkinElmer FT1600 spectrometer. Analytical HPLC was performed on a Shimadzu LC-20AD system. High-resolution mass spectra (HRMS) were acquired on a Waters Xevo G2-S QTOF instrument using ESI/APCI ionization.

### General Information for Biochemical Assays

All biochemical and cell-based assays were performed according to the manufacturers’ instructions or the procedures described below. Test compounds were dissolved in dimethyl sulfoxide (DMSO) or saline to prepare stock solutions and diluted with the appropriate assay medium immediately before use. The final DMSO concentration did not exceed 0.5% (v/v) and had no measurable effect on assay performance. Unless otherwise stated, experiments were performed in triplicate and repeated independently at least three times. Data are expressed as the mean ± standard deviation (SD).

### Comparative Analysis of the Conformational Landscapes and Thermodynamic Unfolding Penalties

The conformational landscapes and thermodynamic unfolding penalties of neutral TM-V and diprotonated TM-Cy-TBPA were investigated. The diprotonated state of TM-Cy-TBPA was selected because docking-based protonation analysis identified it as the most stable species at physiological pH. Since productive binding within the DPAGT1 active site requires adoption of a fully extended conformation, this analysis was designed to quantify the energetic penalty associated with unfolding each compound from its preferred aqueous resting state. All calculations were performed using the ORCA 6.1.1 program package. The conformational search was conducted with the GOAT algorithm using the semiempirical GFN2-xTB method coupled with the ALPB implicit aqueous solvation model. The resulting ensembles were pruned using a 3.0 kcal/mol energy cutoff, thereby retaining conformers representing >99.4% of the thermodynamically relevant population (see Supporting Information).

### Molecular Docking

The binding modes between the molecules and the target protein were investigated using AutoDock Vina (v1.2.7). Ligands were optimized using the GFN2-xTB method37 implemented in ORCA 6.1.121,22 using aqueous implicit solvation. The crystal structure of the target protein DPAGT1 was obtained from the PDB database (PDB ID: 6BW5). Protonation states at physiological pH (7.4) were calculated using the Kirkwood–Westheimer electrostatics with Born desolvation penalties. Protonation states at physiological pH (7.4) were calculated using the Kirkwood–Westheimer electrostatics with Born desolvation penalties. Once properly ionized, the ligand structures were directly converted to pdbqt format unsing the Meeko software, assigning Gasteiger partial charges and rotatable bond topologies. No conformer generation was conducted, as Vina was configured to run a flexible docking search. The 3D crystal structure of the protein target was retrieved from the RCSB Protein Data Bank and prepared using an automated Python workflow built upon PDBFixer found in OpenMM.40 Crystallographic water molecules, co-crystallized solvents, and non-protein heteroatoms were removed. The docking search box was centered on the target binding pocket, obtained from the PDB structure, with dimensions of 62 x 25 x 25 Å to ensure full coverage of the binding site. Simulations were executed with an exhaustiveness setting of 64, using the Vinardo empirical scoring function to evaluate binding affinity. Post-docking pose visualization and hydrogen-bonding interaction analysis (<3.5 Å) were performed to evaluate binding modes, and the obtained complexes were compared with that from 6BW5 structure.

### Synthesis of TM-Cy-TBPA (4)

#### (*R*)-4-(Azidomethyl)cyclohex-1-ene (11)

To a stirred solution of **10** (5.5 g, 49.1 mmol) and Et_3_N (20.4 mL, 147 mmol) in CH_2_Cl_2_ (50 mL) was added TsCl (13.9 g, 98.1 mmol), DMAP (1.2 g, 9.8 mmol) at 0°C. After being stirred for 1h, the reaction mixture was quenched with sat. aq. NaHCO_3_, extracted with EtOAc, and washed with brine. The combined organic extracts were dried over Na_2_SO_4_ and concentrated *in vacuo*. The crude mixture was passed through a silica gel pad (hexanes/EtOAc = 3/1). To a stirred solution of the crude product (13.0 g, 48.8 mmol) in DMF (48 mL) was added NaN_3_ (15.0 g, 24.4 mmol). After being stirred at for 2 h, the reaction was quenched with sat. aq. NaHCO_3_ and extracted with EtOAc. The combined organic extracts were dried over Na_2_SO_4_ and concentrated *in vacuo*. The crude mixture was purified by silica gel column chromatography (hexanes/EtOAc = 3/1) to afford **11** (6.1 g, 91% for 2 steps): ^1^H NMR (400 MHz, CDCl_3_) δ 5.71 – 5.61 (m, 2H), 3.21 (dd, *J* = 6.7, 2.1 Hz, 2H), 2.18 – 2.04 (m, 3H), 1.92 – 1.70 (m, 3H), 1.37 – 1.22 (m, 1H).

#### (*R*)-*N*-(Cyclohex-3-en-1-ylmethyl)acetamide (12)

To a stirred solution of **11** (4.0 g, 29.2 mmol) in pyridine (16 mL) was added AcSH (8 mL). After being stirred at r.t. for 15 h under the dark condition, the reaction mixture was concentrated *in vacuo*. The crude product was purified by silica gel column chromatography (hexanes/EtOAc = 2/1) to afford **12** (4.2 g, 95%):^1^H NMR (400 MHz, CDCl_3_) δ 6.20 (s, 1H), 5.68 – 5.53 (m, 2H), 3.13 (t, *J* = 6.3 Hz, 2H), 2.02 (dd, *J* = 9.1, 4.5 Hz, 4H), 1.96 (s, 3H), 1.79 – 1.59 (m, 3H), 1.26 – 1.17 (m, 1H).

#### *N*-(((1*R,*3*R,*6*S*)-7-Oxabicyclo[4.1.0]heptan-3-yl)methyl)acetamide (13)

To a solution of **12** (1.0 g, 6.53 mmol) and Na_2_CO_3_ (2.0 g, 19.6 mmol) in CH_2_Cl_2_ (13 mL) was added MCPBA (2.3 g, 13.1 mmol). After being stirred at r.t. for 1 h, the reaction mixture was filtered and evaporated *in vacuo*. The crude mixture was purified by silica gel column chromatography (hexanes/EtOAc = 1/2 - CHCl_3_/MeOH = 98/2) to afford **13** (1.0 g, 99%): [α]^21^ + 0.19 (*c* = 0.5, MeOH); IR (thin film) ν_max_ = 3319, 1630 cm^-1^; ^1^H NMR (400 MHz, CDCl_3_) δ 5.52 (d, *J* = 40.0 Hz, 1H), 3.20 – 3.02 (m, 5H), 2.20 – 2.11 (m, 1H), 2.07 – 1.99 (m, 1H), 1.97 (d, *J* = 1.2 Hz, 3H), 1.89 – 1.77 (m, 1H), 1.77 – 1.57 (m, 2H), 1.54 – 1.30 (m, 3H), 1.14 (qd, *J* = 12.3, 4.6 Hz, 1H), 0.97 (qd, *J* = 11.3, 6.7 Hz, 1H); (ESI^+^) *m/z* calcd for C_9_H_15_NO_2_ [M+H] 169.1103, found: 169.1187.

#### *N*-(((1*R*,3*R*,4*R*)-4-Azido-3-hydroxycyclohexyl)methyl)acetamide (14)

To a stirred solution of **13** (1.0 g, 6.5 mmol) and NH_4_Cl (3.5 g, 65 mmol) in DMF (10 mL) was added NaN_3_ (2.1 g, 32 mmol) at 80 °C. After being stirred for 2 h, the reaction mixture was quenched with H_2_O, extracted with AcOEt, and washed with brine. The combined organic extracts were dried over Na_2_SO_4_ and concentrated *in vacuo*. The crude mixture was purified by silica gel column chromatography (hexanes/EtOAc = 50/50 - 20/80 - 0/100) to afford **14** (1.2 g, 90%): [α]^21^ + 0.14 (*c* = 0.5, MeOH); IR (thin film) ν_max_ = 2934, 2099, 1729, 1588, 1534, 1352, 1285, 1154, 1087, 739 cm^-1^; ^1^H NMR (400 MHz, CDCl_3_) δ 5.53 (s, 1H), 3.61 (td, *J* = 9.0, 5.1 Hz, 2H), 3.22 (dh, *J* = 26.5, 6.7 Hz, 2H), 2.00 (s, 3H), 1.92 – 1.76 (m, 5H), 1.60 (dt, *J* = 13.3, 5.7 Hz, 2H), 1.51 (q, *J* = 5.7 Hz, 2H); HRMS (ESI^+^) *m/z* calcd for C_9_H_16_N_4_O_2_ [M+H] 212.1273, found: 213.1273.

#### *N*-(((1*R,*3*R,*4*R*)-4-(1,3-Dioxoisoindolin-2-yl)-3-hydroxycyclohexyl)methyl)acetamide (9)

To a stirred solution of **14** (1.0 g, 4.7 mmol) in MeOH (9.4 mL) was added 10% Pd/C (0.2 g). After being stirred at r.t. for 4 h under H_2_ atmosphere, the reaction solution was filtered through a pad of celite and evaporated *in vacuo*. To a stirred solution of the crude mixture and Et_3_N (1.97 mL, 14.1 mmol) in THF (9.4 mL) was added PhthCO_2_Et (1.2 g, 5.65 mmol). After being stirred at r.t. for 9 h, the reaction was quenched with brine, and extracted with CHCl_3_. The combined organic extracts were dried over Na_2_SO_4_ and evaporated *in vacuo*. The crude mixture was purified by silica gel column chromatography (hexanes/EtOAc = 50/50 - 20/80 - 0/100) to afford **9** (1.2 g, 83 % for 2 steps): [α]^21^ 0.54 (*c* = 0.09, MeOH); IR (thin film) ν_max_ = 3351, 2100, 1634, 650 cm^-1^; ^1^H NMR (400 MHz, CDCl_3_) δ 7.82 (dd, *J* = 5.4, 3.0 Hz, 2H), 7.71 (dd, *J* = 5.5, 3.1 Hz, 2H), 5.72 (s, 1H), 4.35 (td, *J* = 10.6, 4.7 Hz, 1H), 4.20 – 4.10 (m, 1H), 3.63 – 3.54 (m, 1H), 3.28 – 3.20 (m, 1H), 2.41 (td, *J* = 13.4, 5.1 Hz, 2H), 2.20 – 2.09 (m, 2H), 2.05 (s, 1H), 2.01 (s, 4H), 1.80 – 1.53 (m, 13H); HRMS (ESI^+^) *m/z* calcd for C_47_H_64_F_3_N_7_O_14_ [M+H] 316.3570, found: 316.5588.

#### (4a*R,*6*S,*7*R,*8*R,*8a*S*)-6-(((1*R,*2*R,*5*R*)-5-(Acetamidomethyl)-2-(1,3-dioxoisoindolin-2-yl)cyclohexyl)oxy)-7-azido-2,2-dimethylhexahydropyrano[3,2-*d*][1,3]dioxin-8-yl acetate (19)

To a stirred suspension of **9** (0.5 g, 1.6 mmol), **8** (1.2 g, 3.2 mmol), and MS3Å (1.0 g) in CH_2_Cl_2_ (5 mL) were added AgBF_4_ (0.3 g, 1.6 mmol) and NBS (0.7 g, 3.2 mmol) at 0 °C. After being stirred for 16 h, the reaction mixture was added Et_3_N (1.0 mL). The reaction was quenched with sat. aq. NaHCO_3_ and passed through a pad of celite. The solution was extracted with EtOAc, washed with brine, and dried over Na_2_SO_4_. The combined organic extracts were evaporated *in vacuo*. The crude mixture was purified by silica gel column chromatography (hexanes/EtOAc = 30/70 - 20/80 - 10/90) to afford **19** (0.8 g, 91%): [α]^21^ - 0.07 (*c* = 0.60, MeOH); IR (thin film) ν_max_ = 2931, 2107, 1708, 1380, 1225, 1035 cm^-1^; ^1^H NMR (400 MHz, CDCl_3_) δ 7.82 (dt, *J* = 7.1, 3.5 Hz, 2H), 7.68 (dd, *J* = 5.4, 3.0 Hz, 2H), 5.80 (s, 1H), 5.21 (t, *J* = 9.8 Hz, 1H), 4.87 (d, *J* = 3.9 Hz, 1H), 4.44 (td, *J* = 10.5, 4.5 Hz, 1H), 4.32 – 4.21 (m, 1H), 4.12 (q, *J* = 7.1 Hz, 1H), 3.87 – 3.75 (m, 2H), 3.71 – 3.50 (m, 4H), 3.20 (ddd, *J* = 24.6, 9.3, 4.5 Hz, 2H), 2.26 – 2.15 (m, 2H), 2.00 (d, *J* = 5.6 Hz, 6H), 1.81 – 1.65 (m, 7H), 1.41 (s, 4H), 1.35 (s, 4H), 1.31 – 1.23 (m, 3H); ^13^C NMR (100 MHz, CDCl_3_) δ 169.47, 168.80, 133.80, 131.97, 123.04, 99.93, 99.78, 79.43, 72.30, 69.95, 63.78, 62.19, 62.02, 50.54, 40.42, 32.46, 30.97, 29.14, 28.95, 25.60, 23.36, 20.78, 18.92; HRMS (ESI^+^) *m/z* calcd for C_28_H_35_N_5_O_9_ [M+H] 585.6140, found: 585.6139.

#### (4a*R,*6*S,*7*R,*8*R,*8a*S*)-7-Azido-6-(((1*R,*2*R,*5*R*)-2-(1,3-dioxoisoindolin-2-yl)-5-((*N*-nitroso acetamido)methyl)cyclohexyl)oxy)-2,2-dimethylhexahydropyrano[3,2-*d*][1,3]dioxin-8-yl acetate (7)

To a stirred solution of **19** (0.8 g, 1.4 mmol) in a 4:1 mixture of Ac_2_O and AcOH (10 mL) was added NaNO_2_ (2.3 g, 35.0 mmol) at 0 °C. After being stirred for 2 h, the reaction was quenched with ice-cold water, and extracted with CHCl_3_. The combined organic extracts were dried over Na_2_SO_4_ and evaporated *in vacuo*. The crude mixture was purified by silica gel column chromatography (hexanes/EtOAc = 3/1) to afford **7** (0.8 g, 91%): [α]^21^ - 0.09 (*c* = 0.11, MeOH); IR (thin film) ν_max_ = 2108, 1709, 1380 cm^-1^; ^1^H NMR (400 MHz, CDCl_3_) δ 7.83 (dd, *J* = 5.4, 3.0 Hz, 2H), 7.67 (dd, *J* = 5.6, 3.0 Hz, 2H), 5.23 (t, *J* = 9.8 Hz, 1H), 4.81 (d, *J* = 3.8 Hz, 1H), 4.36 (pd, *J* = 10.6, 4.2 Hz, 2H), 4.09 (ddd, *J* = 19.4, 13.8, 8.0 Hz, 2H), 3.91 – 3.78 (m, 3H), 3.71 – 3.62 (m, 1H), 3.55 (t, *J* = 9.5 Hz, 1H), 3.20 (dd, *J* = 10.2, 3.9 Hz, 1H), 2.80 (s, 3H), 2.25 (td, *J* = 12.8, 5.3 Hz, 1H), 1.99 (s, 3H), 1.92 – 1.80 (m, 1H), 1.66 – 1.51 (m, 4H), 1.42 (s, 4H), 1.38 (s, 3H); ^13^C NMR (100 MHz, CDCl_3_) δ 169.47, 168.80, 133.80, 131.97, 123.04, 99.93, 99.78, 79.43, 72.30, 70.22, 63.78, 62.19, 62.02, 50.54, 40.42, 32.46, 30.84, 28.95, 25.60, 23.36, 20.78, 18.92; HRMS (ESI^+^) *m/z* calcd for C_28_H_34_N_6_O_10_ [M+H] 614.2336, found: 614.2336.

#### (4a*R,*6*S,*7*R,*8*R,*8a*S*)-7-Acetamido-6-(((1*R,*2*R,*5*R*)-5-(2-((3a*S,*4*S,*6*R,*6a*R*)-6-(3-((bis(4-fluorophenyl)methoxy)methyl)-2,4-dioxo-3,4-dihydropyrimidin-1(2*H*)-yl)-2,2-dimethyltetrahydrofuro[3,4-*d*][1,3]dioxol-4-yl)-2-oxoethyl)-2-(1,3-dioxoisoindolin-2-yl)cyclohexyl)oxy)-2,2-dimethylhexahydropyrano[3,2-*d*][1,3]dioxin-8-yl acetate (5)

To a solution of **7** (0.8 g, 1.3 mmol) in a 10:1 mixture of toluene and MeOH (11 mL) was added 40% aq. KOH (1.8 mL, 12.9 mmol) at 0 °C. After being stirred for 30 min, MgSO_4_ (1.5 g) was added to the reaction mixture. After being stirred for 5 min at 0 °C, **6** (1.0 g, 1.9 mmol) was added. After being stirred at r.t. for 12 h, the reaction solution was filtered and evaporated *in vacuo*. The crude product was purified by silica gel column chromatography (hexanes/EtOAc = 50/50) to afford **5** (1.2 g, 90 %): [α]^21^ 0.08 (*c* = 0.11, MeOH); IR (thin film) ν_max_ = 3322, 2942, 2832, 1446, 1022, 822, 725, 694, 682, 666 cm^-1^; ^1^H NMR (400 MHz, CDCl_3_) δ 7.74 – 7.68 (m, 2H), 7.58 (dd, *J* = 5.4, 3.4 Hz, 2H), 7.26 (s, 1H), 7.19 – 7.13 (m, 4H), 6.89 (td, *J* = 8.6, 4.5 Hz, 4H), 5.69 (d, *J* = 7.9 Hz, 1H), 5.58 (s, 1H), 5.49 (s, 1H), 5.33 (d, *J* = 9.8 Hz, 1H), 5.29 – 5.22 (m, 2H), 5.13 (t, *J* = 9.8 Hz, 1H), 4.87 (d, *J* = 6.4 Hz, 1H), 4.73 (d, *J* = 3.9 Hz, 1H), 4.52 (d, *J* = 2.4 Hz, 1H), 4.27 (dd, *J* = 10.7, 4.7 Hz, 1H), 4.23 – 4.11 (m, 1H), 3.75 – 3.63 (m, 2H), 3.57 (t, *J* = 11.8 Hz, 1H), 3.46 (t, *J* = 9.2 Hz, 1H), 3.08 (dd, *J* = 10.1, 3.9 Hz, 1H), 2.84 (dd, *J* = 16.9, 8.3 Hz, 1H), 2.46 – 2.27 (m, 2H), 2.14 (td, *J* = 14.7, 6.1 Hz, 1H), 1.92 (s, 3H), 1.83 (d, *J* = 12.7 Hz, 1H), 1.66 – 1.57 (m, 3H), 1.56 – 1.48 (m, 6H), 1.45 (d, *J* = 13.9 Hz, 1H), 1.38 – 1.28 (m, 10H); ^13^C NMR (100 MHz, CDCl_3_) δ 205.90, 169.51, 168.68, 163.40, 162.35, 160.95, 151.03, 142.21, 137.48, 133.73, 131.96, 128.70, 122.98, 115.33, 115.11, 114.07, 102.33, 99.89, 98.38, 94.18, 84.65, 82.81, 81.85, 79.40, 72.30, 69.94, 69.45, 63.84, 62.18, 61.96, 60.42, 50.64, 39.26, 32.66, 29.27, 28.95, 28.30, 26.73, 25.04, 21.08, 20.78, 18.92, 14.21; HRMS (ESI^+^) *m/z* calcd for C_54_H_58_F_2_N_4_O_16_ [M+H] 1056.3816, found: 1056.3856.

#### (2*S,*3*R,*4*R,*5*S,*6*R*)-3-Azido-2-(((1*R,*2*R,*5*S*)-5-((*R*)-2-((3a*R,*4*R,*6*R,*6a*R*)-6-(3-((bis(4-fluorophenyl)methoxy)methyl)-2,4-dioxo-3,4-dihydropyrimidin-1(2*H*)-yl)-2,2-Dimethyltetrahydrofuro[3,4-*d*][1,3]dioxol-4-yl)-2-hydroxyethyl)-2-(1,3-dioxoisoindolin-2-yl)cyclohexyl)oxy)-5-hydroxy-6-(hydroxymethyl)tetrahydro-2*H*-pyran-4-yl acetate methane (20)

To a stirred solution of **5** (1.2 g, 1.1 mmol) in a 1:3:6 mixture of HCO_2_H:Et_3_N:^i^PrOH (10 mL) was added RuCl[(*R*,*R*)-Ts-DPEN](*p*-cymene) (72 mg, 0.1 mmol). After being stirred at r.t. for 17 h, the reaction was quenched with H_2_O, and extracted with CHCl_3_. The combined organic extracts were dried over Na_2_SO_4_ and evaporated *in vacuo*. The crude mixture was purified by silica gel column chromatography (hexanes/EtOAc = 50/50) to afford **20** (1.1 g, 92%): [α]^21^ 0.27 (*c* = 0.5, MeOH); IR (thin film) = ν_max_ 3322, 2942, 2832, 1446, 1022, 725, 694, 682, 666 cm^-1^; ^1^H NMR (400 MHz, CDCl_3_) δ 7.82 (dt, *J* = 7.2, 3.5 Hz, 2H), 7.68 (dd, *J* = 5.5, 3.1 Hz, 2H), 7.29 (dd, *J* = 9.0, 4.6 Hz, 3H), 7.18 (d, *J* = 8.1 Hz, 1H), 6.98 (td, *J* = 8.7, 3.2 Hz, 4H), 5.76 – 5.66 (m, 2H), 5.50 (s, 2H), 5.42 (d, *J* = 3.2 Hz, 1H), 5.23 (t, *J* = 9.8 Hz, 1H), 5.03 (dd, *J* = 6.8, 3.6 Hz, 1H), 4.96 (dd, *J* = 6.8, 3.1 Hz, 1H), 4.84 (d, *J* = 3.8 Hz, 1H), 4.49 – 4.29 (m, 2H), 4.17 – 4.07 (m, 2H), 3.95 (d, *J* = 9.8 Hz, 1H), 3.88 – 3.75 (m, 2H), 3.73 – 3.62 (m, 1H), 3.55 (t, *J* = 9.4 Hz, 1H), 3.30 (s, 1H), 3.19 (dd, *J* = 10.3, 3.8 Hz, 1H), 2.39 (td, *J* = 12.7, 5.0 Hz, 1H), 2.27 (s, 1H), 2.00 (s, 4H), 1.85 – 1.65 (m, 6H), 1.62 (s, 10H), 1.41 (d, *J* = 4.8 Hz, 6H), 1.36 (s, 3H); ^13^C NMR (101 MHz, CDCl3) δ 169.46, 168.87, 163.41, 162.16, 160.96, 151.06, 141.77, 137.51, 133.75, 132.02, 128.60, 128.46, 123.00, 115.37, 115.16, 114.82, 102.39, 99.90, 99.79, 97.03, 89.58, 82.94, 82.17, 79.75, 78.41, 72.33, 70.00, 69.20, 63.81, 62.23, 62.06, 50.79, 28.97, 27.31, 25.25, 20.79, 18.91; HRMS (ESI^+^) *m/z* calcd for C_52_H_56_F_2_N_6_O_15_ [M+H] 1042.3772, found: 1042.3782.

#### (4a*R,*6*S,*7*R,*8*R,*8a*S*)-7-acetamido-6-(((1*R,*2*R,*5*S*)-5-((*R*)-2-((3a*R,*4*R,*6*R,*6a*R*)-6-(3-((bis(4-fluorophenyl)methoxy)methyl)-2,4-dioxo-3,4-dihydropyrimidin-1(2*H*)-yl)-2,2-dimethyltetrahydrofuro[3,4-*d*][1,3]dioxol-4-yl)-2-hydroxyethyl)-2-(1,3-dioxoisoindolin-2-yl)cyclohexyl)oxy)-2,2-dimethylhexahydropyrano[3,2-*d*][1,3]dioxin-8-yl acetate

To a stirred solution of **20** (1.0 g, 1.0 mmol) in pyridine (2 mL) was added AcSH (2 mL). After being stirred at r.t. for 4 days under the dark condition, the reaction mixture was concentrated *in vacuo*. The crude product was purified by silica gel column chromatography (hexanes/EtOAc = 50/50 - 30/70) to afford the corresponding acetamide (1.0 g, 97%): [α]^21^ 0.03 (*c* = 0.05, MeOH); IR (thin film) = ν_max_ 3322, 2942, 2832, 1446, 1022, 1079, 822, 725, 694, 682, 666 cm^-1^; ^1^H NMR (400 MHz, CDCl_3_) δ 7.80 (dd, *J* = 5.4, 3.4 Hz, 2H), 7.73 (dd, *J* = 5.6, 2.9 Hz, 2H), 7.28 (q, *J* = 4.5 Hz, 3H), 7.15 (d, *J* = 8.2 Hz, 1H), 6.98 (t, *J* = 8.6 Hz, 4H), 5.69 (q, *J* = 4.6 Hz, 2H), 5.53 – 5.47 (m, 2H), 5.41 (d, *J* = 10.0 Hz, 1H), 5.39 – 5.34 (m, 1H), 5.09 – 4.95 (m, 3H), 4.63 (dd, *J* = 11.1, 4.1 Hz, 1H), 4.52 (q, *J* = 6.1 Hz, 1H), 4.19 – 4.02 (m, 3H), 3.92 (d, *J* = 9.2 Hz, 1H), 3.86 (dd, *J* = 10.1, 4.7 Hz, 1H), 3.79 – 3.63 (m, 3H), 2.43 – 2.24 (m, 2H), 1.98 (s, 3H), 1.78 (s, 6H), 1.62 (s, 6H), 1.60 (s, 4H), 1.55 (s, 3H), 1.41 (d, *J* = 5.6 Hz, 6H), 1.36 (s, 3H), 1.26 (d, *J* = 11.9 Hz, 3H); HRMS (ESI^+^) *m/z* calcd for C_54_H_60_F_2_N_4_O_16_ [M+H] 1058.3972, found:1058.3982.

#### *N*-((4a*R*,6*S*,7*R*,8*R*,8a*S*)-6-(((1*R*,2*R*,5*S*)-2-Amino-5-(2-((2*R*,5*R*)-5-(3-((bis(4-fluorophenyl)methoxy)methyl)-2,4-dioxo-3,4-dihydropyrimidin-1(2*H*)-yl)-3,4-dihydroxytetrahydrofuran-2-yl)-2-hydroxyethyl)cyclohexyl)oxy)-8-hydroxy-2,2-dimethylhexahydropyrano[3,2-*d*][1,3]dioxin-7-yl)acetamide (21)

To a stirred solution of the acetamide (1.0 g, 0.9 mmol) in EtOH (9 mL) was added ethylenediamine (1 mL). After being stirred at 60 °C for 5 h, the reaction mixture was concentrated *in vacuo*. The crude mixture was purified by a short path silica gel column chromatography (CHCl_3_/MeOH/ Et_3_N = 90/10/0.1) to afford **21** (0.7 g, 88%): ^1^H NMR (400 MHz, MeOD) δ 7.67 (d, *J* = 8.2 Hz, 1H), 7.33 (dd, *J* = 8.7, 5.4 Hz, 5H), 7.03 (t, *J* = 8.7 Hz, 5H), 5.83 (t, *J* = 3.2 Hz, 1H), 5.73 (s, 1H), 5.67 (t, *J* = 7.2 Hz, 1H), 5.57 – 5.50 (m, 2H), 4.79 (ddd, *J* = 12.2, 6.7, 3.0 Hz, 1H), 3.97 (dd, *J* = 10.5, 3.9 Hz, 2H), 3.85 (q, *J* = 4.0 Hz, 2H), 3.80 – 3.67 (m, 5H), 3.58 (dd, *J* = 17.9, 8.9 Hz, 2H), 3.51 (s, 1H), 3.35 (d, *J* = 1.0 Hz, 1H), 3.32 (s, 3H), 3.05 (q, *J* = 7.3 Hz, 3H), 2.00 (s, 3H), 1.57 (s, 3H), 1.52 (s, 4H), 1.39 (d, *J* = 2.8 Hz, 6H); HRMS (ESI^+^) *m/z* calcd for C_44_H_58_F_2_N_4_O_13_ [M+H] 887.3812, found: 887.3845.

#### *tert*-Butyl (2-(((1*R,*2*R,*4*S*)-2-(((4a*R,*6*S,*7*R,*8*R,*8a*S*)-7-acetamido-8-hydroxy-2,2-dimethylhexahydropyrano[3,2-*d*][1,3]dioxin-6-yl)oxy)-4-((*R*)-2-((3a*R,*4*R,*6*R,*6a*R*)-6-(3-((bis(4-fluorophenyl)methoxy)methyl)-2,4-dioxo-3,4-dihydropyrimidin-1(2*H*)-yl)-2,2-dimethyltetrahydrofuro[3,4-*d*][1,3]dioxol-4-yl)-2-hydroxyethyl)cyclohexyl)amino)-2-oxoethyl)(4-(4-(((*tert*-butoxycarbonyl)(4-(trifluoromethoxy)benzyl)amino)methyl)piperidin-1-yl)benzyl)carbamate

To a stirred solution of **21** (50 mg, 56 µmol), **22** (74 mg, 0.11 mmol), HOSu (13 mg, 0.11 mmol) and NMM (50 µL, 0.45 mmol) in DMF (0.5 mL) was added EDCI (32 mg, 0.17 mmol). After being stirred at r.t. for 5 h, the reaction was quenched with sat. aq. NaHCO_3_ and extracted with CHCl_3_. The combined organic extracts were dried over Na_2_SO_4_ and concentrated *in vacuo*. The crude product was purified by short path silica gel column chromatography (hexanes/EtOAc = 50/50 - CHCl_3_/MeOH = 95/5) to afford the coupling product (77 mg, 90%): [α]^21^ -0.03 (*c* = 0.90, MeOH); IR (thin film) ν_max_ = 3279, 1637, 1015 cm^-1^; ^1^H NMR (400 MHz, CD_3_OD) δ 7.67 (d, *J* = 8.2 Hz, 1H), 7.40 – 7.28 (m, 7H), 7.25 (d, *J* = 8.8 Hz, 3H), 7.15 (d, *J* = 8.6 Hz, 2H), 7.05 – 6.97 (m, 5H), 6.93 (d, *J* = 8.6 Hz, 2H), 5.86 (d, *J* = 2.8 Hz, 1H), 5.72 (s, 1H), 5.66 (d, *J* = 8.1 Hz, 1H), 5.51 (s, 2H), 4.78 (d, *J* = 4.4 Hz, 1H), 4.50 (s, 3H), 4.45 – 4.27 (m, 2H), 3.96 (d, *J* = 10.5 Hz, 2H), 3.90 – 3.78 (m, 4H), 3.75 (d, *J* = 10.0 Hz, 3H), 3.68 (q, *J* = 8.3 Hz, 5H), 3.59 (t, *J* = 9.2 Hz, 1H), 3.49 (s, 1H), 3.18 (d, *J* = 16.5 Hz, 2H), 2.61 (s, 2H), 2.00 (s, 3H), 1.73 (d, *J* = 13.0 Hz, 5H), 1.58 (s, 5H), 1.51 (s, 8H), 1.47 (d, *J* = 12.7 Hz, 15H), 1.40 (d, *J* = 7.5 Hz, 15H); ^13^C NMR (100 MHz, CD_3_OD) δ 172.20, 169.93, 163.35, 163.06, 160.92, 156.07, 151.40, 150.79, 141.27, 138.09, 128.38, 120.83, 116.71, 114.78, 114.56, 114.09, 101.02, 99.58, 92.62, 89.19, 84.06, 81.77, 79.62, 74.74, 69.78, 68.55, 67.98, 64.10, 61.88, 54.52, 29.53, 28.16, 27.33, 26.26, 25.89, 24.19, 21.37, 18.02; HRMS (ESI^+^) *m/z* calcd for C_77_H_98_F_5_N_7_O_19_ [M+H] 1519.6838, found:1519.6887.

#### *N*-((1*R,*2*R,*4*S*)-2-(((2*S,*3*R,*4*R,*5*S,*6*R*)-3-Acetamido-4,5-dihydroxy-6-(hydroxymethyl)tetrahydro-2*H*-pyran-2-yl)oxy)-4-((2*R*)-2-((2*R,*5*R*)-5-(2,4-dioxo-3,4-dihydropyrimidin-1(2*H*)-yl)-3,4-dihydroxytetrahydrofuran-2-yl)-2-hydroxyethyl)cyclohexyl)-2-((4-(4-(((4-(trifluoromethoxy)benzyl)amino)methyl)piperidin-1-yl)benzyl)amino)acetamide (TM-Cy-TBPA (4))

A solution of the compound synthesized above (30 mg, 20 µmol) in a 4:1 mixture of AcOH and H_2_O (1 mL) was stirred at 60 °C for 5 h. The reaction mixture was concentrated *in vacuo*. The crude product was purified by short path silica gel column chromatography (CHCl_3_/MeOH = 90/10 - CHCl_3_/MeOH - CHCl_3_/MeOH/H_2_O/NH_4_OH = 56/42/7/3) The resulted mixture was purified by reverse-phase HPLC [column: HYPERSIL GOLD^TM^ (C18, 12 µm, 175 Å, 250 x 10 mm), solvents: 60:40 MeOH:H_2_O, flow rate: 3.0 mL/min, UV: 254 nm] to afford TM-Cy-TBPA (**4**, 17 mg, 85%): [α]^21^ + 0.74 (*c* = 0.50, MeOH); IR (thin film) ν_max_ = 3351, 2100, 1634, 650.0 cm^-1^; ^1^H NMR (400 MHz, MeOD) δ 7.91 (d, *J* = 8.2 Hz, 1H), 7.50 (d, *J* = 8.8 Hz, 2H), 7.27 (d, *J* = 8.3 Hz, 2H), 7.21 (d, *J* = 8.7 Hz, 2H), 6.96 (d, *J* = 8.7 Hz, 2H), 5.90 (d, *J* = 5.6 Hz, 1H), 5.71 (d, *J* = 8.1 Hz, 1H), 4.65 (s, 4H), 4.21 (d, *J* = 4.2 Hz, 2H), 3.93 – 3.80 (m, 8H), 3.74 – 3.61 (m, 9H), 3.35 (s, 1H), 3.24 (s, 2H), 2.67 (t, *J* = 11.4 Hz, 2H), 2.59 (d, *J* = 6.8 Hz, 2H), 2.06 (s, 2H), 2.00 (s, 3H), 1.91 – 1.81 (m, 4H), 1.77 (s, 1H), 1.67 (s, 3H), 1.63 – 1.41 (m, 6H), 1.37 (s, 5H), 1.30 (s, 7H); ^13^C NMR (100 MHz, MeOD) δ 172.20, 171.93, 164.76, 151.18, 148.31, 141.73, 137.84, 130.03, 129.09, 120.68, 116.84, 101.55, 97.16, 88.74, 88.23, 73.85, 72.88, 71.18, 71.00, 69.17, 68.37, 61.38, 54.20, 54.09, 52.38, 52.15, 50.48, 49.95, 35.21, 29.95, 29.40, 28.08, 26.75, 25.66, 21.41; HRMS (ESI^+^) *m/z* calcd for C_47_H_64_F_3_N_7_O_14_ [M+H] 1008.4463, found:1008.4448.

### Biochemical Assay Procedures

#### Water solubility of TM-Cy-TBPA (4) and TM-TBPA (3)

A suspension of **3** or **4** in H_2_O (50 μL) was stirred for 24 h, and the precipitate was separated by centrifugation at 10,000 x g for 5 min. The upper solution (1 μL) was analyzed via reverse-phase HPLC [column: Kinetex (1.7 μm XB-C18, 100 Å, 150 x 2.10 mm), solvents: 60:40 MeOH : H_2_O, flow rate: 0.5 mL/min, UV: 254 nm]. The area of the peak for **3** or **4** was quantified. The concentrations were determined via the HPLC intensity-concentration curves (see Supporting Information).

#### DPAGT1 expression and purification

Human DPAGT1 was transiently expressed in suspension Expi293 cells for 36 h. Cells were harvested and lysed by passage through a 26-gauge needle (10 times), and membrane proteins were extracted in buffer containing 1% n-dodecyl-β-D-maltoside (DDM). DPAGT1 was purified by HA affinity chromatography using HA-agarose resin, followed by size-exclusion chromatography on a Superdex 200 column.^5,7,8^

#### DPAGT1 assays

DPAGT1 assay substrate, UDP-Glucosamine-C_6_-FITC was chemically synthesized according to the reported procedures. UDP-glucosamine-C_6_-FITC (2 mM stock, 0.56 μL), MgCl₂ (0.5 M, 4.0 μL), β-mercaptoethanol (50 mM, 5.0 μL), CHAPS (20%, 2.5 μL), C55-dolichyl phosphate (2 mM, 1.7 μL), inhibitor (0–50 μg/mL in 50 mM Tris buffer, pH 8.0), and Tris buffer were combined in a 1.5 mL microcentrifuge tube to a final volume of 40 μL. The reaction was initiated by the addition of purified DPAGT1 (10 μL; final reaction volume, 50 μL) and incubated for 3 h. Reactions were quenched with n-butanol (150 μL), vortexed, and centrifuged at 25,000 × g for 5 min. The upper organic phase was collected, and a 30 μL aliquot was analyzed by reverse-phase HPLC (Kinetex C8, 5 μm, 100 Å, 150 × 4.6 mm; gradient elution from 85:15 to 100:0 MeOH/0.05 M NH₄HCO₃; flow rate, 0.5 mL min⁻¹; detection at 485 nm). The peak area corresponding to C55-PP-glucosamine-C_6_-FITC was quantified, and IC₅₀ values were determined by nonlinear regression of the percentage inhibition versus inhibitor concentration.^42^

#### Cytotoxicity assay

Cells (5 × 10^4^ cells/well, 195 μL) were seeded into 96-well plates and treated with 5 μL of inhibitor solution at the indicated concentrations. Following incubation for 48–96 h at 37 °C in a humidified atmosphere containing 5% CO₂, WST-1 cell proliferation reagent (10 μL) was added to each well, and the plates were incubated for an additional 3 h. Cell viability was determined by measuring the conversion of WST-1 to its soluble formazan product. Absorbance was measured at 440 nm using a BioTek Synergy HT microplate reader. EC₅₀ and CC_50_ values were calculated by nonlinear regression analysis of cell viability versus inhibitor concentration. All cell lines used in this study were obtained from ATCC.

#### Eosin staining of red blood cells

A drop of blood was placed on a clean glass slide and spread into a thin smear, which was allowed to air-dry. The smear was fixed by briefly immersing the slide in absolute methanol, followed by staining with 0.5% eosin in ethanol for 30 min. Excess stain was removed by gently rinsing the slide with water. After air-drying, images were acquired using an AMG EVOS XL transmitted-light microscope.

#### Membrane disruption assay using 7-AAD

A confluent cell monolayer was established in Nunc™ Lab-Tek™ II CC2™ 8-well chamber slides (Thermo Scientific, Cat. No. 154941PK). Cells were incubated for 6 h at 37 °C in a humidified atmosphere containing 5% CO₂ with complete culture medium supplemented with 7-AAD staining solution (1:20 dilution; BD Biosciences, Cat. No. B559925) and either TM-V (0–1.0 μM) or TM-Cy-TBPA (0–10 μM). Following treatment, the culture medium was removed, and the cells were washed twice with PBS before fixation with 4% paraformaldehyde in PBS for 30 min at 4 °C. The fixed cells were washed twice with PBS containing 0.2% Tween-20, mounted with glass coverslips, and examined by fluorescence microscopy. Fluorescence intensity was quantified using ImageJ software (v1.54d; National Institutes of Health).

#### LDH release and membrane permeability assay

Lactate dehydrogenase (LDH) release was measured using the LDH Cytotoxicity Colorimetric Assay Kit (TriboScience, Cat. No. TBS2002) according to the manufacturer’s instructions. Cells (1–2 × 10⁴ cells/well, 98 μL) were seeded into 96-well plates and allowed to attach. Test compounds (2.0 μL) were then added to each well to achieve the desired final concentrations. All treatments were performed in duplicate, and cell-free wells containing culture medium and the corresponding drug concentrations were included as blank controls. After incubation with the compounds for 6 h at 37 °C in a humidified atmosphere containing 5% CO₂, either 10 μL of cell lysis buffer (maximum LDH release control) or PBS (spontaneous LDH release) was added to each well, followed by incubation for an additional 10–20 min at 37 °C with gentle shaking (50–100 rpm). Subsequently, 50 μL of the supernatant from each well was transferred to a new 96-well plate, mixed with 50 μL of LDH working solution, and incubated for 1 h at 37 °C with gentle shaking in the dark.

#### SDS–PAGE and Western blot analysis

A confluent cell monolayer was established in 6-well plates and incubated with complete culture medium containing the test inhibitor (0–10 μM) for 48 h at 37 °C in a humidified atmosphere with 5% CO₂. Following treatment, the culture medium was removed, and the cells were washed with PBS before lysis in Pierce RIPA buffer (Thermo Scientific, Cat. No. 89901) supplemented with 1× Pierce Protease and Phosphatase Inhibitor Cocktail (Thermo Scientific, Cat. No. 88668). Cell lysates were clarified by centrifugation at 25,000 × g for 30 min at 4 °C, and the supernatants were collected. Protein concentrations were determined using the Quick Start™ Bradford Protein Assay (Bio-Rad, Cat. No. 500-0205). Equal amounts of protein (50 μg per sample) were separated by SDS–PAGE on 10% polyacrylamide gels and transferred for Western blot analysis. Precision Plus Protein™ Dual Color Standards (Bio-Rad, Cat. No. 161-0374) were used as molecular weight markers. Membranes were probed with primary antibodies (see Table 10 and 11). Horseradish peroxidase (HRP)-conjugated anti-rabbit IgG (Cell Signaling Technology, Cat. No. 7074) or anti-mouse IgG (Cell Signaling Technology, Cat. No. 7076) secondary antibodies were used for detection. Immunoreactive bands were visualized using Clarity™ Western ECL Substrate (Bio-Rad, Cat. No. 170-5060) and exposed to Classic Blue BX X-ray film (MidSci, Ref. No. 6045983). Band intensities were quantified using ImageJ software (v1.54d; National Institutes of Health).

#### Cytochrome *c* release assay

Cytochrome *c* release was evaluated using the Abcam Cytochrome *c* Release Assay Kit (Fisher Scientific, Cat. No. 50-220-4699) according to the manufacturer’s instructions. Confluent cells in 6-well plates were treated with inhibitor (0–10 μM) for 48 h at 37 °C under 5% CO₂. Cytosolic and mitochondrial fractions were prepared using the kit extraction buffers and differential centrifugation. Protein concentrations were determined using the Quick Start™ Bradford Protein Assay (Bio-Rad, Cat. No. 500-0205). Equal amounts of protein (10 μg) from each fraction were analyzed by 12% SDS–PAGE followed by Western blotting. Immunoreactive bands were visualized using Clarity™ Western ECL Substrate (Bio-Rad, Cat. No. 170-5060) and Classic Blue BX X-ray film (MidSci, Ref. No. 6045983). Band intensities were quantified using ImageJ software (v1.54d; National Institutes of Health).

#### Pharmacokinetic studies

All *in vivo* pharmacokinetic studies in CD-1 mice were performed by BioDuro (Irvine, CA) as a contract research organization (CRO). Animal dosing, sample collection, bioanalysis, and noncompartmental pharmacokinetic analyses were conducted according to BioDuro’s established protocols and institutional guidelines for the ethical use of laboratory animals.

## ASSOCIATED CONTENT

### Supporting Information

This material is available free of charge via the Internet at http://pubs.acs.org. The Supporting Information includes experimental procedures; additional biological assay data; copies of ^1^H and ^13^C NMR spectra for all new compounds; analytical HPLC chromatograms demonstrating the purity of new compounds; high-resolution mass spectrometry (HRMS) data; and supplementary Figures and Tables supporting the biological and pharmacokinetic studies.

## NOTES

The authors declare no competing financial interest.

## ABBREVIATIONS

DPAGT1: dolichylphosphate *N*-acetyl glucosamine phosphotrans-ferase 1;
TM-V: tunicamycin V;
TM-Cy-TBPA: a tunicamycin cyclitol analogue;
MraY: phospho-MurNAc-pentapeptide translocase (translocase I);
MurX: mycobacterial translocase I;
WecA: UDP-*N*-acetylglucosamine:undecaprenyl-phosphate *N*-acetylglucosamine-1-phosphate transferase;
IC_50_: half-maximal inhibitory concentration;
EC₅₀: half-maximal effective concentration;
CC₅₀: half-maximal cytotoxic concentration;
SI: selectivity index;
ER: endoplasmic reticulum;
TMPA: ((((trifluoromethoxy)phenoxy)piperidin-1-yl)phenyl)methoxymethyl;
TBPA: 4-(trifluoromethoxy)benzylpiperidin-4-yl-methylamino-benzylamino)acetamide;
CD-1 mice: outbred mouse strain (Charles River Laboratories);
TM-Cy: tunicamycin cyclitol;
SAR: structure–activity relationship;
GOAT: gradient optimization of atomic topologies;
GFN2-xTB: geometry, frequency, noncovalent, eXtended tight-binding (second generation);
ORCA: *ORCA* quantum chemistry program package (version 6.1.1);
ALPB: analytical linearized poisson–boltzmann (implicit solvation model);
PBE0: Perdew–Burke–Ernzerhof hybrid exchange–correlation functional;
def2-SVP: split-valence polarized basis set (Ahlrichs def2 family);
CPCM: conductor-like polarizable continuum model;
qRRHO: quasi-rigid rotor harmonic oscillator;
def2-TZVPP: Ahlrichs triple-ζ valence basis set with two sets of polarization functions;
RMSD: root-mean-square deviation;
C15:1-*iso*: 15 carbon atoms:1 carbon double bond: *iso* = methyl branching at the second-to-last carbon atom;
cryo-EM: cryo-electron microscopy;
GOAT: global optimizer algorithm;
XTB: extended tight-binding program;
LigPrep: Schrödinger LLC, New York, NY, USA;
BCS: Büchner–Curtius–Schlotterbeck;
BFPM: 4,4’-bisfluorophenyl)methoxymethyl;
mCPBA: *m*-chloroperoxybenzoic acid;
NOESY: nuclear overhauser effect spectroscopy;
Ns: 4-nitrobenzenesulfonyl;
Boc: *tert*-butyloxycarbonyl;
SN_2_: bimolecular nucleophilic substitution;
NBS: *N*-bromosuccinimide;
EDCI: (1-ethyl-3-(3-dimethylaminopropyl)carbodiimide;
HOSu: *N*-hydroxysuccinimide;
NMM: *N*-methylmorpholine;
DMF: dimethylformamide;
Phth: phthaloyl;
Ac: acetyl;
Dol-P: dolichyl phosphate;
UMP: uridine 5′-monophosphate;
ATP: adenosine 5′-triphosphate;
HPLC: high-performance liquid chromatography;
FITC: fluorescein isothiocyanate;
MDA-MB-231: human triple-negative breast cancer cells (ATCC HTB-26);
MDA-MB-468: human triple-negative breast cancer cells (ATCC HTB-132);
SKBr-3: human HER2-positive breast cancer cells (ATCC HTB-30);
RBC: red blood cell;
ANOVA: one-way analysis of variance;
MCF-7: human breast adenocarcinoma cells (ATCC HTB-22);
7-AAD: 7-aminoactinomycin D;
LDH: lactate dehydrogenase;
UPR: unfolded protein response;
p53: p53 protein;
p21: cyclin-dependent kinase inhibitor 1A;
G_2_: gap 2 phase;
M: mitosis;
G1: gap 1 phase;
G_0_: a cellular state outside of the replicative cell cycle;
CDK1: cyclin-dependent kinase 1;
CDK2: cyclin-dependent kinase 2;
CDC2: cell division cycle protein 2 (cyclin-dependent kinase 1);
BiP/GRP78: binding immunoglobulin protein/glucose-regulated protein 78;
XBP1s: spliced X-box binding protein 1;
CHOP: CCAAT/enhancer-binding protein homologous protein;
Bcl-2: B-cell lymphoma 2, IRE1α, inositol-requiring enzyme 1 alpha;
PERK: protein kinase R-like endoplasmic reticulum kinase;
ATF4: activating transcription factor 4;
PARP: poly(ADP-ribose) polymerase;
PK: pharmacokinetic;
p.o: oral administration;
AUC: area under the concentration–time curve, V_z_, volume of distribution during the terminal phase;
CL: systemic (total body) clearance;
APPB: aminouridyl phenoxypiperidinbenzylbutanamide: a muraymycin A1 analogue.

