## Supplemental Files for "Rationally Engineered, Chemically Stable Tunicamycin Analogues Decouple DPAGT1 Inhibition from Non-Selective Toxicity"

for

**Table of Contents**

1. General S2
2. Experimental procedure S3-S44
3. References S45
4. Copies of $^{1}H$ and $^{13}C$ NMR spectra S46-S79

### 1. General

All chemicals were purchased from commercial sources and used without further purification unless otherwise noted. THF, $CH_2Cl_2$, and DMF were purified via Innovative Technology's Pure-Solve System. All reactions were performed under an Argon atmosphere. All stirring was performed with an internal magnetic stirrer. Reactions were monitored by TLC using 0.25 mm coated commercial silica gel plates (EMD, Silica Gel $60F_{254}$). TLC spots were visualized by UV light at 254 nm, or stained with ceric ammonium molybdate or anisaldehyde or phosphomolybdic acid hydrate or ninhydrin solutions by heating on a hot plate. Reactions were also monitored by using SHIMADZU LCMS-2020 with solvents: A: 0.1% formic acid in water, B: acetonitrile. Flash chromatography was performed with SiliCycle silica gel (Purasil 60 Å, 230-400 Mesh). Proton magnetic resonance ($^{1}$H-NMR) spectral data were recorded on 400 MHz instruments. Carbon magnetic resonance ($^{13}$C-NMR) spectral data were recorded on 100 MHz instruments. For all NMR spectra, chemical shifts (δH, δC) were quoted in parts per million (ppm), and *J* values were quoted in Hz. $^{1}$H and $^{13}$C NMR spectra were calibrated with residual undeuterated solvent ($CDCl_3$: δH = 7.26 ppm, δC = 77.16 ppm; $CD_3CN$: δH = 1.94 ppm, δC = 1.32ppm; $CD_3OD$: δH =3.31 ppm, δC =49.00 ppm; DMSO-$d_6$: δH = 2.50 ppm, δC = 39.52 ppm; $D_2O$: δH = 4.79 ppm) as an internal reference. The following abbreviations were used to designate the multiplicities: s = singlet, d = doublet, dd = double doublets, t = triplet, q = quartet, quin = quintet, hept = heptet, m = multiplet, br = broad. Infrared (IR) spectra were recorded on a Perkin-Elmer FT1600 spectrometer. HPLC analyses were performed with a Shimadzu LC-20AD HPLC system. HR-MS data were obtained from a Waters Xevo G2-S QTOF measured by ESI/APCI Quadrupole TOF.

**Table S1:** Eighteen novel tunicamycin analogues with enhanced activity relative to tunicamycin V (TM-V).

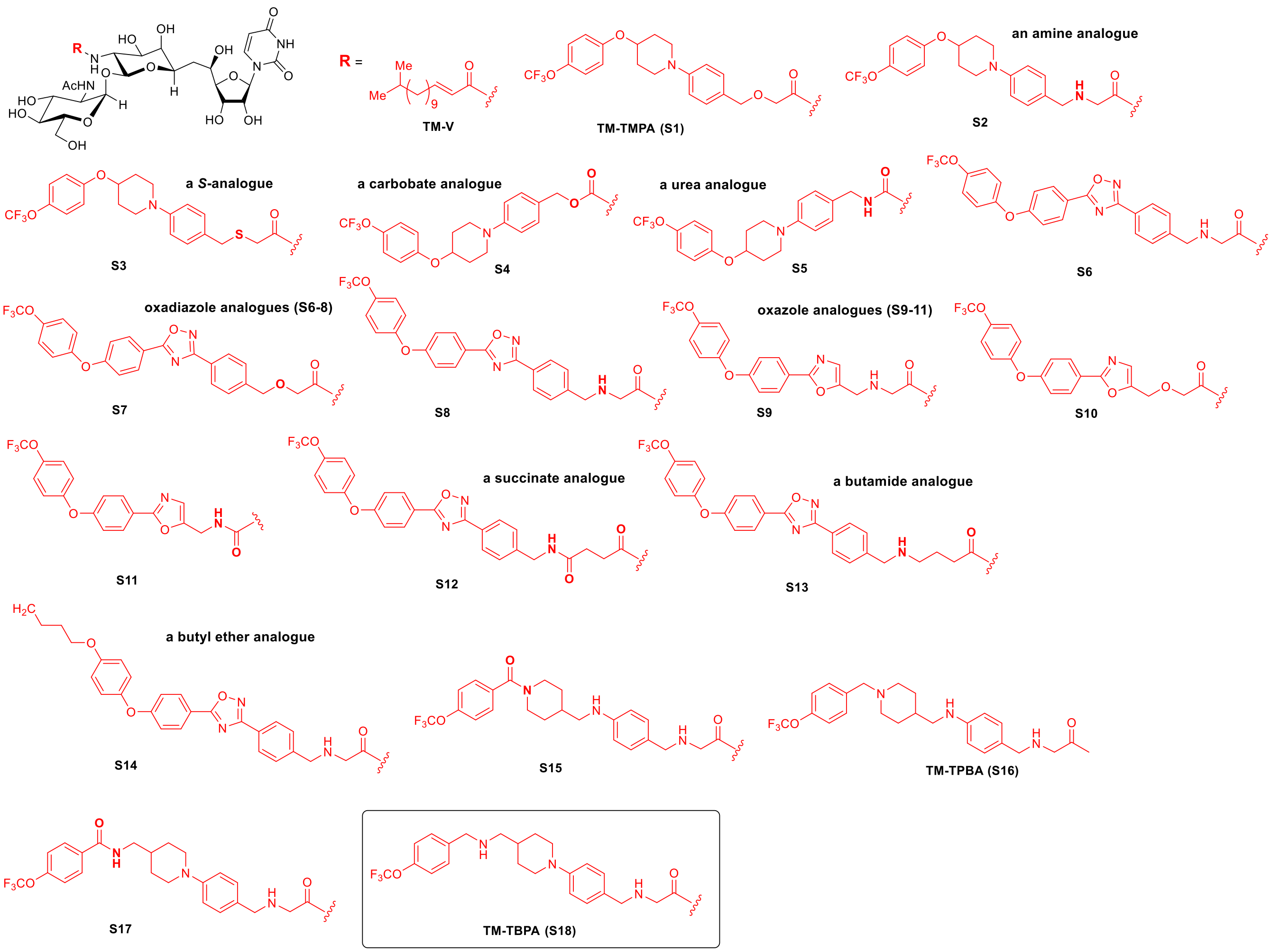


| | TM-V | | | S1 | | | S2 | | | S3 | | | S4 | | |
|---|---|---|---|---|---|---|---|---|---|---|---|---|---|---|---|
| concentrations (μM) | 0.1 | 1.0 | 5.0 | 0.1 | 1.0 | 5.0 | 0.1 | 1.0 | 5.0 | 0.1 | 1.0 | 5.0 | 0.1 | 1.0 | 5.0 |
| enzymatic assays (DPAGT1) | − | + | + | − | + | +++ | − | ++ | ++ | − | + | ++ | − | + | ++ |
| scratch assays | − | + | cell losses | + | + | ++ | − | ++ | +++ | − | + | +++ | − | + | ++ |
| cardiolipin expression inhibition (NAO) | − | + | cell losses | − | + | ++ | − | ++ | +++ | − | − | ++ | − | + | ++ |

| | S5 | | | S6 | | | S7 | | | S8 | | | S9 | | |
|---|---|---|---|---|---|---|---|---|---|---|---|---|---|---|---|
| concentrations (μM) | 0.1 | 1.0 | 5.0 | 0.1 | 1.0 | 5.0 | 0.1 | 1.0 | 5.0 | 0.1 | 1.0 | 5.0 | 0.1 | 1.0 | 5.0 |
| enzymatic assays (DPAGT1) | − | + | ++ | − | + | ++ | − | − | ++ | − | − | + | − | + | ++ |
| scratch assays | − | + | + | − | + | +++ | − | − | ++ | − | − | ++ | − | ++ | +++ |
| cardiolipin expression inhibition (NAO) | − | + | ++ | + | ++ | +++ | − | + | ++ | − | − | + | − | + | ++ |

| | S10 | | | S11 | | | S12 | | | S13 | | | S14 | | |
|---|---|---|---|---|---|---|---|---|---|---|---|---|---|---|---|
| concentrations (μM) | 0.1 | 1.0 | 5.0 | 0.1 | 1.0 | 5.0 | 0.1 | 1.0 | 5.0 | 0.1 | 1.0 | 5.0 | 0.1 | 1.0 | 5.0 |
| enzymatic assays (DPAGT1) | − | + | ++ | − | + | ++ | − | + | +++ | − | − | + | − | + | ++ |

| | | | | | | | | | | | | | | | |
|---|---|---|---|---|---|---|---|---|---|---|---|---|---|---|---|
| scratch assays | − | − | ++ | − | + | +++ | − | + | +++ | − | − | + | − | − | ++ |
| cardiolipin expression inhibition (NAO) | − | + | ++ | − | + | +++ | − | ++ | +++ | − | − | ++ | − | + | ++ |

| | **S15** | | | **S16** | | | **S17** | | | **S18 (3)** | | |
|---|---|---|---|---|---|---|---|---|---|---|---|---|
| concentrations (μM) | 0.1 | 1.0 | 5.0 | 0.1 | 1.0 | 5.0 | 0.1 | 1.0 | 5.0 | 0.1 | 1.0 | 5.0 |
| enzymatic assays (DPAGT1) | + | + | ++ | + | ++ | +++ | + | + | ++ | + | ++ | +++ |
| scratch assays | − | + | +++ | − | ++ | +++ | − | + | +++ | − | ++ | +++ |
| cardiolipin expression inhibition (NAO) | − | + | +++ | + | ++ | +++ | − | + | +++ | + | ++ | +++ |

Inhibition is categorized semi-quantitatively as follows: −: no inhibition.; +: 5–30% inhibition.; ++: 30–70% inhibition.; +++: ≥70% inhibition. Values represent percent inhibition measured at the indicated concentration and time point.

Data for **3** (**S18**)

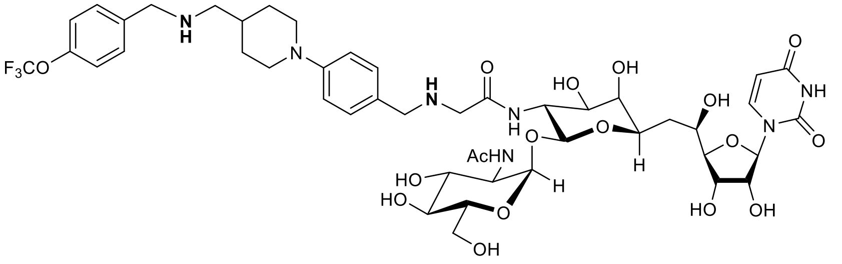

**TM-TBPA** (3)

$^{1}$H NMR (400 MHz, MeOD) δ 7.90 (d, *J* = 7.9 Hz, 1H), 7.79 – 7.74 (m, 2H), 7.72 – 7.67 (m, 4H), 7.41 (d, *J* = 7.8 Hz, 2H), 5.87 (d, *J* = 4.4 Hz, 1H), 5.71 (d, *J* = 7.9 Hz, 1H), 5.40 (s, 1H), 5.07 (d, *J* = 3.5 Hz, 1H), 4.32 (d, *J* = 6.6 Hz, 4H), 4.21 (d, *J* = 3.5 Hz, 2H), 3.94 – 3.85 (m, 2H), 3.85 – 3.74 (m, 6H), 3.73 – 3.56 (m, 6H), 3.17 – 3.12 (m, 2H), 2.37 – 2.29 (m, 1H), 2.22 – 2.15 (m, 2H), 2.09 – 2.04 (m, 1H), 2.00 (s, 3H), 1.95 – 1.88 (m, 2H), 1.80 (d, *J* = 14.0 Hz, 1H), 1.72 – 1.65 (m, 1H), 1.61 – 1.53 (m, 2H), 1.50 – 1.41 (m, 2H); $^{13}$C NMR (100 MHz, MeOD) δ 175.93, 173.58, 164.77, 164.54, 162.34, 152.20, 142.73, 139.51, 129.75 (2C), 129.67 (2C), 116.16 (2C), 115.94 (2C), 115.44, 102.30, 100.96, 99.49, 94.16, 90.68, 85.54, 83.21, 80.95, 76.34, 76.10, 71.20, 70.02, 69.55, 65.45, 63.21, 55.87, 37.04, 32.54, 29.53, 27.58, 26.79, 25.54, 23.44, 22.74, 19.38, 14.33; HRMS (ESI+) *m/z* calcd for $C_{46}H_{63}F_3N_7O_{17}$ [M + H] 1042.4233, found: 1042.4251.

HPLC analysis for TM-TBPA (**3**)

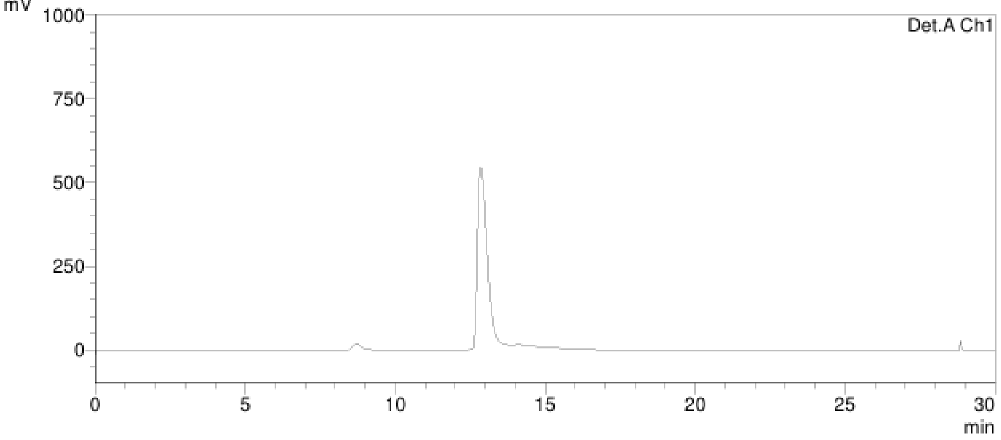

HPLC conditions for TM-TBPA (**3**)
solvent: 60:40 MeOH/$H_2O$
UV: 254 nm
flow rate: 0.5 ml/ min
column: Kinetex 1.7 μm XB-C18, 100 Å, 150 x 2.10 mm

**Comparative Analysis of the Conformational Landscapes and Thermodynamic Unfolding Penalties of TM-Cy-TBPA and TM-V**

The conformational landscapes and thermodynamic unfolding penalties of neutral TM-V and diprotonated TM-Cy-TBPA were investigated. The diprotonated state of TM-Cy-TBPA was selected because docking-based protonation analysis identified it as the most stable species at physiological pH. Since productive binding within the DPAGT1 active site requires adoption of a fully extended conformation, this analysis was designed to quantify the energetic penalty associated with unfolding each compound from its preferred aqueous resting state.

**Computational Methods**

All calculations were performed using the ORCA 6.1.1 program package.[1,2] The conformational search was conducted with the GOAT algorithm using the semiempirical GFN2-xTB method[3] coupled with the ALPB implicit aqueous solvation model.[4] The resulting ensembles were pruned using a 3.0 kcal/mol energy cutoff, thereby retaining conformers representing >99.4% of the thermodynamically relevant population.

The global minima geometries were subsequently optimized using the PBE0 functional in conjunction with the def2-SVP basis set and Grimme's D4 dispersion correction.[5] Solvent effects during optimization were implicitly incorporated using the Conductor-like Polarizable Continuum Model (CPCM) for water. Vibrational frequency calculations were performed directly at the PBE0-D4/def2-SVP level on the optimized structures to confirm they were true minima (zero imaginary frequencies). Thermochemical corrections were computed utilizing Grimme's quasi-RRHO scheme[6] applying a cutoff frequency of 100 $cm^{-1}$.

To refine the energetic values, single-point electronic energy calculations were performed on the optimized geometries using the extended def2-TZVPP basis set at the PBE0-D4 level, employing the SMD implicit solvation model for water. Final Gibbs free energies were obtained by adding the qRRHO thermochemical corrections to the SMD-solvated def2-TZVPP electronic energies.

**GOAT Optimization and Conformational Ensemble Analysis**

The structures of TM-V and the cyclitol analogue were subjected to GOAT optimization and conformational sampling using the GFN2-xTB semiempirical method with ALPB implicit aqueous solvation. The resulting conformational ensembles were subsequently pruned using a 3.0 kcal/mol energy cutoff, corresponding to retention of conformers encompassing approximately 99.4% of the thermodynamically accessible population at room temperature. The geometric distributions of these pruned ensembles were then analyzed to characterize the dominant solution-phase conformations and overall molecular shapes adopted in aqueous media.

**Figure S1.** Population distribution of conformers within the pruned (<3.0 kcal/mol) conformational ensemble generated from GOAT calculations for TM-V.

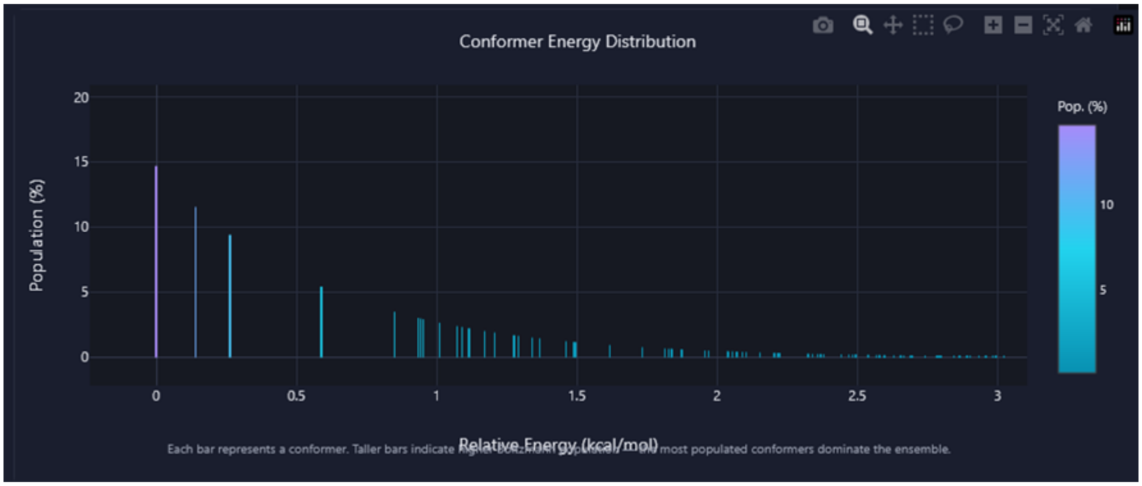


Figure S1 illustrates the conformational population distribution of TM-V as a function of relative conformer energy. The global minimum accounted for approximately 15% of the total ensemble population, followed by an exponential decay in conformer populations with increasing energy. The higher-energy region of the landscape (>1.0 kcal/mol) displayed a dense continuum of accessible states, consistent with substantial conformational flexibility and extensive low-energy structural diversity.

**Figure S2.** Shape analysis of the pruned conformational ensemble (<3.0 kcal/mol) generated for TM-V following GOAT conformational sampling.

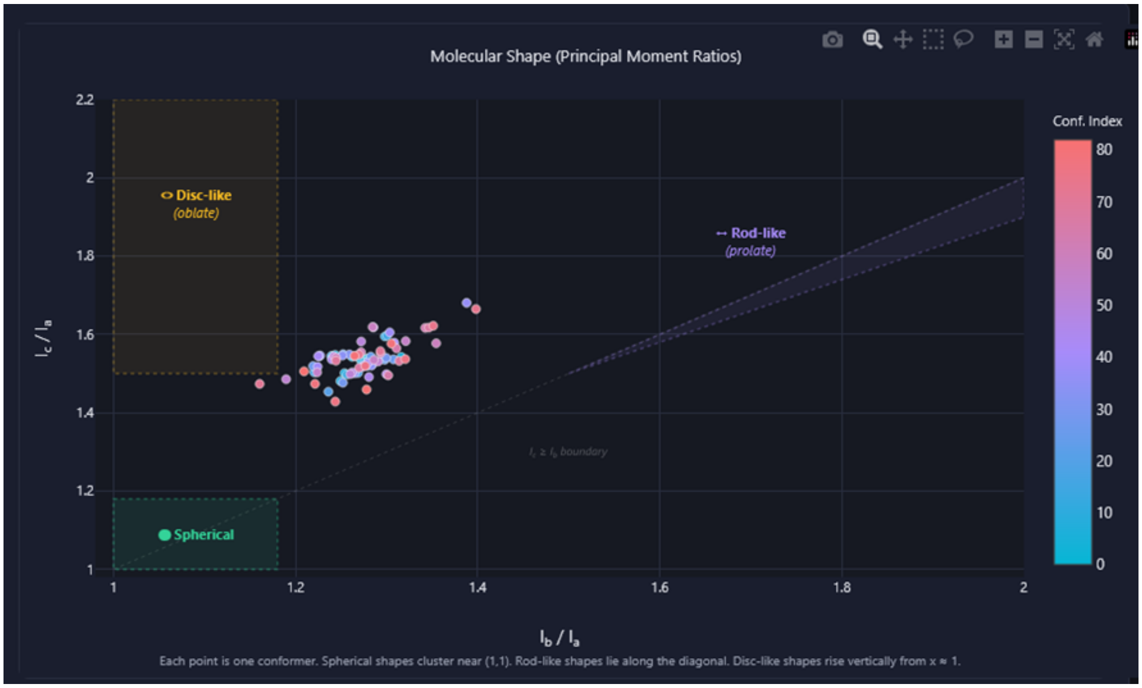


Shape analysis of the pruned TM-V conformational ensemble revealed predominantly elongated and irregular molecular geometries that clustered within an RMSD threshold of <0.5 Å (**Figure S2**). Approximately 85% of the conformers within the ensemble were distributed among the first five major clusters. The relative populations and superimposed structures of the first 10 conformers within each cluster are presented in **Table S2**.

**Table S2.** Population distribution of the major conformational clusters comprising 85% of the pruned TM-V ensemble, together with superimposed structures of the first 10 conformers within each cluster.

| Cluster ID | Number of conformes in the cluster | Superimposed conformational structures |
|---|---|---|
| 1 | 55 (67%) | 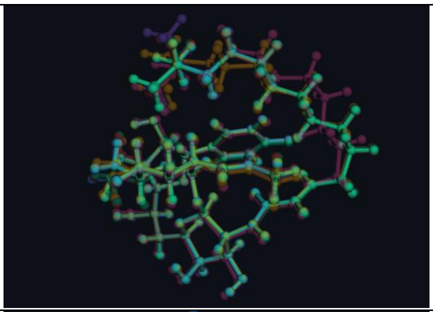 |
| 2 | 6 (7%) | 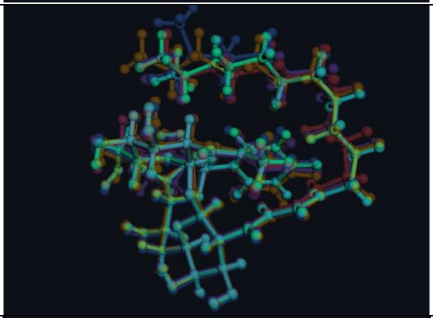 |
| 3 | 5 (6%) | 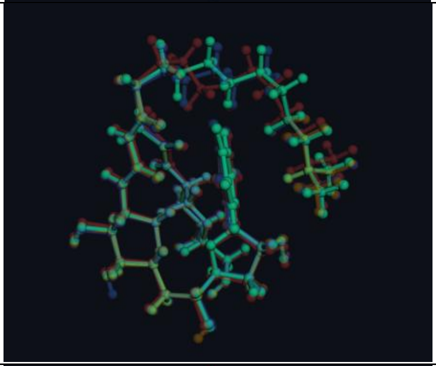 |
| 4 | 3 (4%) | 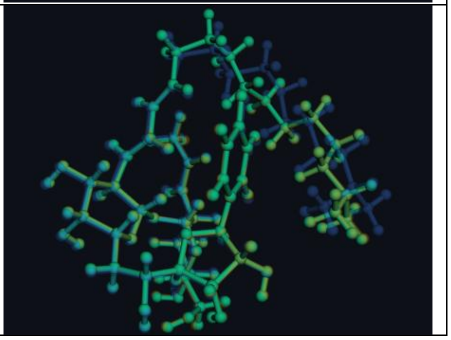 |

| | | |
|---|---|---|
| 5 | 2 (2%) | 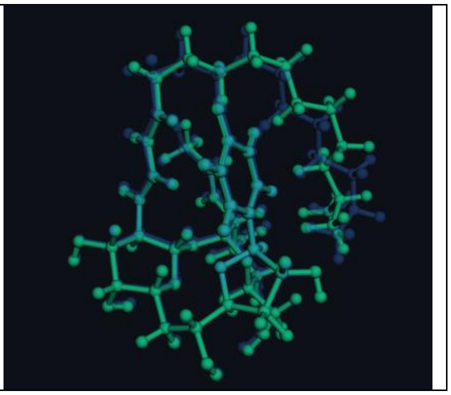 |

**Figure S3.** Population distribution of conformers within the pruned conformational ensemble of TM-Cy-TBPA.

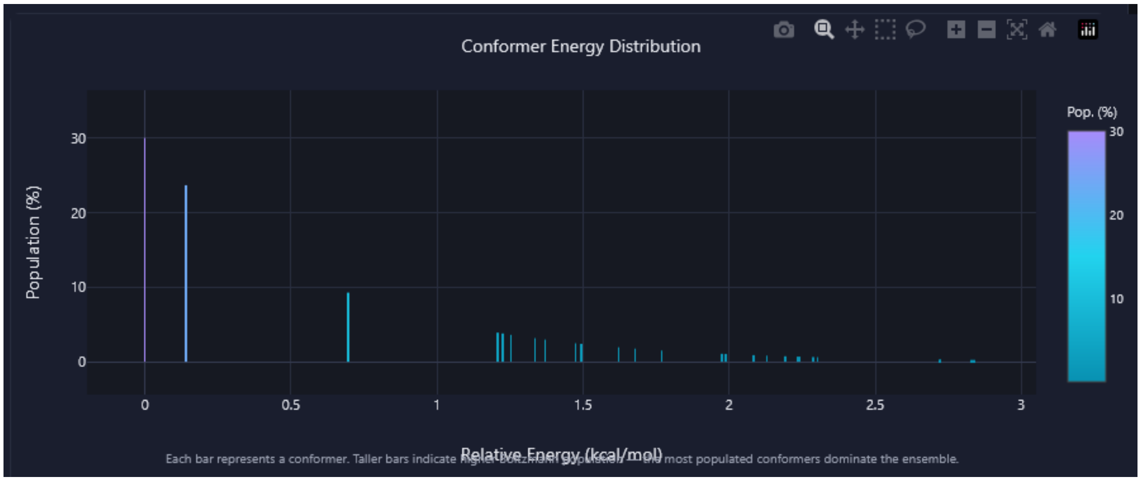


An analogous conformational analysis was performed for TM-Cy-TBPA, which likewise exhibited an exponential decay in conformer population with increasing relative energy. However, in contrast to TM-V, the ensemble displayed a markedly stronger population bias toward the first three dominant geometries, indicating a substantially more restricted conformational landscape (**Figure S3**).

**Figure S4**. Shape analysis of the pruned conformational ensemble (<3.0 kcal/mol) generated for the cyclitol analogue following GOAT conformational sampling.

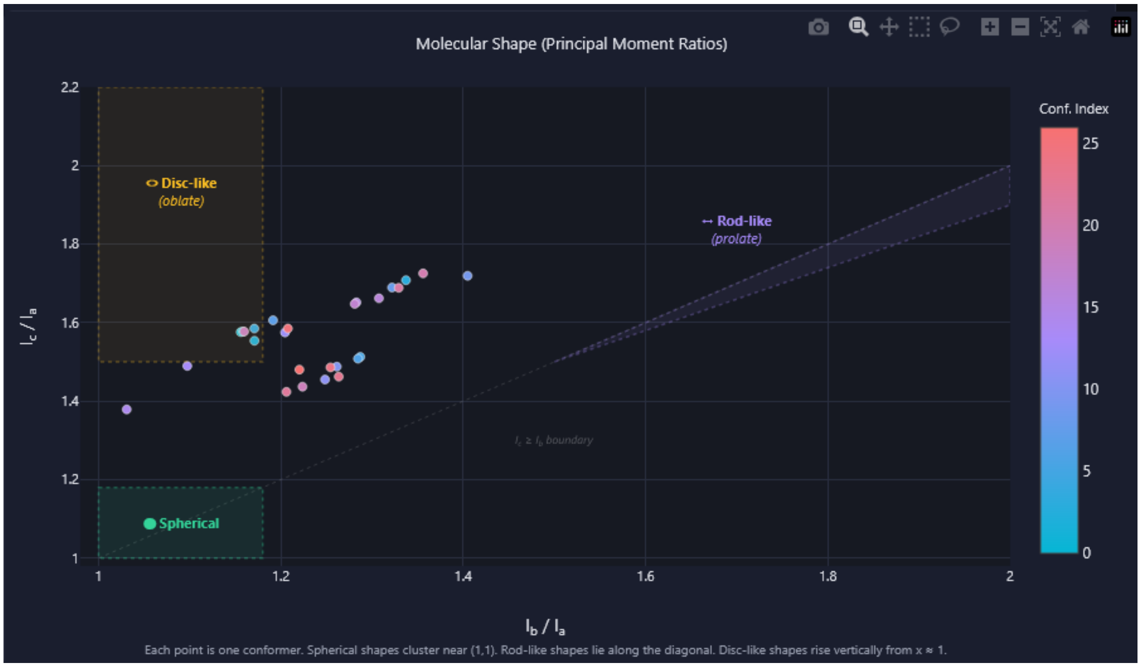


Shape analysis of the pruned conformational ensemble of the cyclitol analogue revealed a broader structural dispersion than observed for TM-V. Although a subset of conformers adopted relatively oblate geometries, the majority exhibited irregular elongated shapes characteristic of partially extended conformations. This increased diversity of molecular shapes was also reflected in the clustering behavior of the ensemble, where 85% of the pruned conformational population was distributed across 35 distinct clusters, each exhibiting substantially smaller individual populations relative to those observed for the TM-V ensemble (**Table S3**).

**Table S3**. Population distribution of the major conformational clusters comprising 85% of the pruned ensemble population of TM-Cy-TBPA, together with superimposed structures of the first 10 conformers within each cluster.

| Cluster ID | Number of conformers in the cluster | Superimposed conformational structures |
|---|---|---|
| 1 | 5 (19%) | 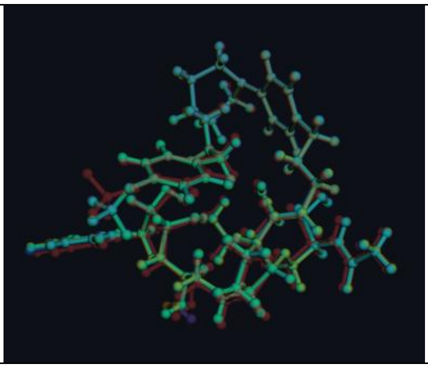 |

| | | |
|---|---|---|
| 2 | 5 (19%) | 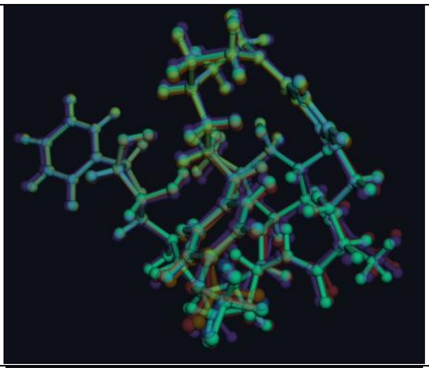 |
| 3 | 4 (15%) | 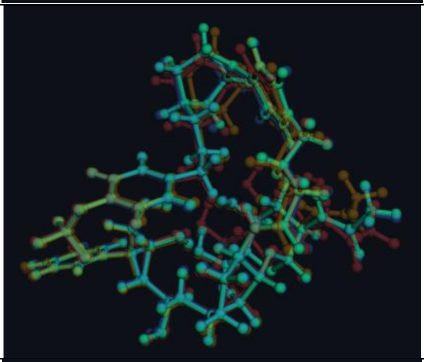 |
| 4 | 2 (8%) | 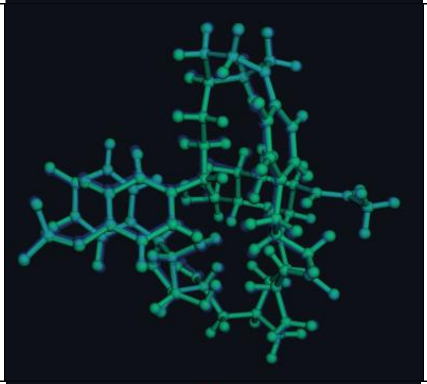 |
| 5 | 2 (8%) | 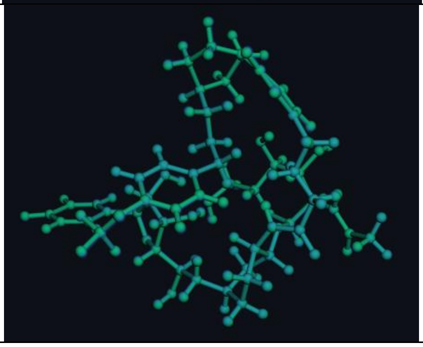 |

| | | |
|---|---|---|
| 6 | 2 (8%) | 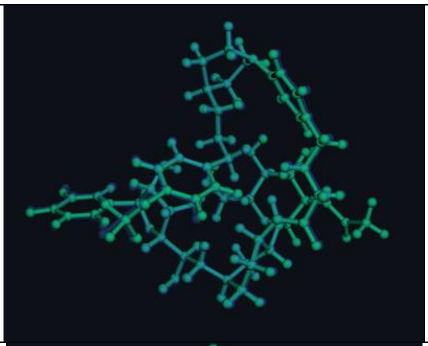 |
| 7 | 1 (4%) | 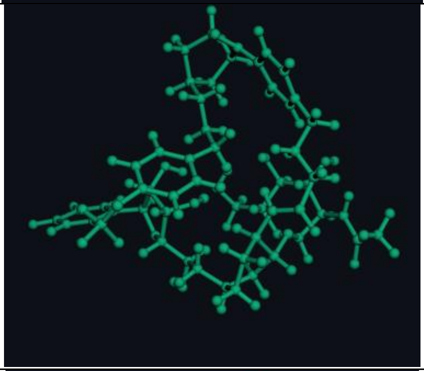 |
| 8 | 1 (4%) | 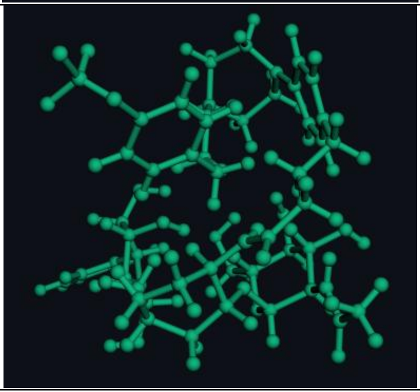 |

**Conformational Ensembles and Geometric Restriction**

Both compounds preferentially adopted compact folded conformations in bulk aqueous media, though driven by entirely distinct physicochemical forces. For TM-V, the folded state minimized the energetically unfavorable hydrophobic exposure of its lipidic side chain. For TM-Cy-TBPA, the folded state was stabilized because the high dielectric constant of water efficiently shielded the intense Coulombic repulsion between its two protonated amines, permitting a compact resting state.

- **TM-V:**
  TM-V exhibited a classical Boltzmann conformational distribution characterized by a dense continuum of accessible, near-isoenergetic folded states, showing a high density of accessible states particularly in the higher energy landscape (>1.0 kcal/mol above the global minimum). The neutral lipidic tail retained substantial configurational freedom, enabling extensive dihedral rotation and conformational sliding within the folded ensemble. This produced a highly dynamic energy landscape with considerable conformational entropy.
- **TM-Cy-TBPA:**
  In contrast, TM-Cy-TBPA displayed a markedly skewed conformational distribution in which >50% of the thermodynamic population was confined to only two dominant geometries separated by pronounced

energetic gaps. The strong Coulombic repulsion between the two protonated amines functioned as an "electrostatic straitjacket," severely restricting conformational flexibility. This intramolecular electrostatic tension heavily destabilized intermediate folded states, substantially reduced configurational entropy, and trapped the molecule within a limited number of highly strained local minima.

**Thermodynamics of Unfolding ($\Delta G_{unfold}$)**

- **TM-V:**
The conformational behavior of TM-V was governed by its neutral, amphipathic nature. Unfolding the folded aqueous conformer incurred in a thermodynamic penalty ($\Delta G_{unfold}$ = +7.59 kcal/mol), driven primarily by classical hydrophobic effects: the C15:1-iso aliphatic chain spontaneously packed against the polar molecular core to minimize energetically unfavorable exposure to the aqueous environment. When evaluated in a low-dielectric lipophilic environment (1-octanol), which simulates the dielectric conditions of the membrane-bound DPAGT1 active site, the unfolding penalty underwent only a modest reduction to +5.08 kcal/mol. This indicated that the energetic cost of extending the hydrophobic tail remains a substantial thermodynamic barrier that is relatively insensitive to solvent polarity drops.

- **TM-Cy-TBPA:**
The thermodynamic unfolding profile of the dicationic analogue was fundamentally dictated by environmental electrostatics. In bulk water ($\varepsilon \approx 80$), the high dielectric constant effectively shielded the internal Coulombic repulsion between the two protonated amines. Consequently, the molecule remained locked in a compact, stable resting state with a slightly higher unfolding penalty than TM-V ($\Delta G_{unfold}$ =+8.18 kcal/mol). However, a dramatic thermodynamic reversal occurred upon transition to the low-dielectric lipophilic environment of 1-octanol ($\varepsilon \approx 10$). The loss of aqueous dielectric shielding magnified the unshielded Coulombic repulsion between the two cationic centers. This electrostatic destabilization forcefully disrupted the folded resting state, causing the unfolding penalty to collapse to just +2.60 kcal/mol. This structural transition represented a spontaneous Coulombic unfolding of the molecule. TM-Cy-TBPA therefore functions via a "dielectric trigger", exploiting the lipidic environment of the DPAGT1 active site to spontaneously render the bioactive extended conformation thermodynamically accessible without requiring a heavy compensatory binding penalty.

**TM-V (Folded conformer, global minimum, DFT optimized, water):**
Charge: 0 | Multiplicity: 1 | Imaginary frequency: ---
$E_{elec}$(def2-TZVPP): -2906.053722 a.u.
Thermal correction to Gibbs Free Energy (quasi-RRHO) (def2-SVP): 0.934514 a.u.
Absolute Gibbs Free Energy (TZVPP): -2905.119208 a.u.

| | | | |
|---|---|---|---|
| O | 0.710307 | 2.100657 | -2.482632 |
| C | 1.146349 | 1.182485 | -3.181817 |
| N | 0.341645 | 0.240234 | -3.748606 |
| C | -1.111338 | 0.243194 | -3.684729 |
| C | -1.729346 | 1.448821 | -4.444080 |
| O | -1.459099 | 1.365395 | -5.824335 |
| C | -3.241536 | 1.572939 | -4.175581 |
| O | -3.970533 | 0.575158 | -4.870043 |
| C | -3.468175 | 1.532696 | -2.663289 |
| C | -4.912063 | 1.730911 | -2.206251 |
| C | -5.103597 | 2.366182 | -0.815753 |
| O | -6.478798 | 2.385503 | -0.484960 |
| C | -4.273204 | 1.712119 | 0.306976 |
| C | -4.646607 | 2.138491 | 1.750761 |
| O | -5.145455 | 1.082855 | 2.545777 |
| C | -3.304428 | 2.679190 | 2.338018 |

| | | | |
|---|---|---|---|
| O | -3.124847 | 2.353500 | 3.682653 |
| C | -2.268341 | 2.051581 | 1.385065 |
| N | -0.995655 | 2.745950 | 1.240354 |
| C | -0.961834 | 3.992073 | 0.656179 |
| C | 0.190327 | 4.631242 | 0.343843 |
| C | 1.465768 | 3.986744 | 0.605190 |
| N | 1.335000 | 2.745019 | 1.235673 |
| C | 0.181713 | 2.067907 | 1.559514 |
| O | 0.196742 | 0.965911 | 2.080698 |
| O | 2.571601 | 4.429025 | 0.331011 |
| O | -2.901863 | 2.068039 | 0.129017 |
| O | -3.027793 | 0.250600 | -2.198563 |
| C | -1.625399 | 0.110371 | -2.232525 |
| O | -1.305298 | -1.173698 | -1.781042 |
| C | -0.872412 | -1.334126 | -0.447334 |
| C | 0.046629 | -2.567907 | -0.410000 |
| N | 1.131028 | -2.429427 | -1.353369 |
| C | 2.409783 | -2.795562 | -1.081945 |
| C | 3.398756 | -2.585604 | -2.209155 |
| O | 2.759619 | -3.241590 | 0.009569 |
| C | -0.778393 | -3.849901 | -0.620356 |
| O | 0.001313 | -5.026661 | -0.570367 |
| C | -1.917879 | -3.900239 | 0.406015 |
| O | -2.767152 | -4.999545 | 0.164896 |
| C | -2.759848 | -2.615577 | 0.350812 |
| C | -3.772425 | -2.573426 | 1.492601 |
| O | -4.711777 | -1.517778 | 1.400999 |
| O | -1.926353 | -1.462027 | 0.466817 |
| C | 2.596056 | 1.013765 | -3.475628 |
| C | 3.502678 | 1.897473 | -3.026196 |
| C | 4.966510 | 1.876731 | -3.330630 |
| C | 5.906737 | 2.147211 | -2.143860 |
| C | 6.025902 | 1.028727 | -1.100798 |
| C | 4.871647 | 0.904831 | -0.101422 |
| C | 5.120962 | -0.174015 | 0.955213 |
| C | 4.028083 | -0.279974 | 2.019346 |
| C | 4.293296 | -1.376972 | 3.055020 |
| C | 3.250605 | -1.480151 | 4.175327 |
| C | 1.865712 | -1.939883 | 3.710136 |
| C | 0.828455 | -2.158994 | 4.825990 |
| C | -0.442648 | -2.794757 | 4.252635 |
| C | 0.492354 | -0.869775 | 5.585472 |
| H | 0.775772 | -0.445838 | -4.360847 |
| H | -1.444951 | -0.678014 | -4.201328 |
| H | -1.249242 | 2.374260 | -4.073863 |
| H | -2.099067 | 0.729316 | -6.189631 |
| H | -3.580355 | 2.549031 | -4.575138 |
| H | -3.918990 | -0.247394 | -4.353487 |
| H | -2.832792 | 2.329288 | -2.223072 |
| H | -5.443776 | 2.373714 | -2.931788 |
| H | -5.423251 | 0.747116 | -2.225453 |

| | | | |
|---|---|---|---|
| H | -4.790249 | 3.429558 | -0.861617 |
| H | -6.813484 | 1.473872 | -0.503802 |
| H | -4.370679 | 0.608100 | 0.238320 |
| H | -5.411095 | 2.935536 | 1.748889 |
| H | -4.876313 | 0.212501 | 2.179517 |
| H | -3.275364 | 3.782106 | 2.223486 |
| H | -3.741618 | 1.612328 | 3.850047 |
| H | -2.031748 | 1.023462 | 1.721517 |
| H | -1.938655 | 4.437886 | 0.438025 |
| H | 0.178910 | 5.618836 | -0.124742 |
| H | 2.203145 | 2.250237 | 1.454193 |
| H | -1.161922 | 0.879835 | -1.581328 |
| H | -0.309527 | -0.438633 | -0.120688 |
| H | 0.493735 | -2.603866 | 0.602071 |
| H | 0.895128 | -2.106329 | -2.289931 |
| H | 4.063229 | -3.463343 | -2.272482 |
| H | 2.919266 | -2.416837 | -3.187040 |
| H | 4.029177 | -1.708892 | -1.973197 |
| H | -1.228329 | -3.832563 | -1.633787 |
| H | 0.521325 | -5.020874 | 0.250850 |
| H | -1.454393 | -3.968654 | 1.419885 |
| H | -2.198772 | -5.783715 | 0.088923 |
| H | -3.302138 | -2.596896 | -0.620480 |
| H | -4.284130 | -3.554076 | 1.535818 |
| H | -3.228606 | -2.436939 | 2.445988 |
| H | -5.346767 | -1.710588 | 0.693010 |
| H | 2.907572 | 0.155961 | -4.088902 |
| H | 3.135281 | 2.740109 | -2.418455 |
| H | 5.240852 | 0.927558 | -3.830709 |
| H | 5.143052 | 2.678458 | -4.079485 |
| H | 5.607578 | 3.092525 | -1.646040 |
| H | 6.912161 | 2.337584 | -2.564526 |
| H | 6.169509 | 0.058476 | -1.621032 |
| H | 6.959104 | 1.194375 | -0.525537 |
| H | 4.715969 | 1.884166 | 0.397446 |
| H | 3.922515 | 0.685644 | -0.630179 |
| H | 6.095387 | 0.019257 | 1.451000 |
| H | 5.232630 | -1.156562 | 0.451292 |
| H | 3.054940 | -0.454538 | 1.519410 |
| H | 3.927764 | 0.695372 | 2.541419 |
| H | 4.371772 | -2.354458 | 2.535296 |
| H | 5.288862 | -1.199538 | 3.510057 |
| H | 3.622276 | -2.189381 | 4.942923 |
| H | 3.177368 | -0.498163 | 4.685227 |
| H | 1.448888 | -1.212208 | 2.985985 |
| H | 1.982881 | -2.889959 | 3.149267 |
| H | 1.266010 | -2.875353 | 5.555077 |
| H | -0.910495 | -2.130840 | 3.500380 |
| H | -0.225076 | -3.757251 | 3.753267 |
| H | -1.194134 | -2.988167 | 5.040553 |
| H | 1.381864 | -0.420269 | 6.062019 |

| | | | |
|---|---|---|---|
| H | -0.246479 | -1.061419 | 6.385685 |
| H | 0.056993 | -0.110854 | 4.906898 |

**TM-V (Extended conformer, DFT optimized, water):**
Charge: 0 | Multiplicity: 1 | Imaginary frequency: ---
$E_{elec}$(def2-TZVPP): -2906.035883 a.u.
Thermal correction to Gibbs Free Energy (quasi-RRHO) (def2-SVP): 0.928772 a.u.
Absolute Gibbs Free Energy (TZVPP): -2905.107111 a.u.

| | | | |
|---|---|---|---|
| C | -1.206670 | 0.427672 | 4.609061 |
| C | -1.499708 | -0.432300 | 3.427448 |
| C | -2.805518 | -0.458846 | 1.298187 |
| C | -2.666204 | -2.164895 | -1.032887 |
| C | -1.827810 | -2.292445 | -3.445162 |
| C | -2.244217 | -2.560961 | -4.896940 |
| C | -2.137141 | -3.991935 | -6.815515 |
| C | -2.435048 | -4.031400 | -5.298884 |
| C | 2.350514 | -4.035295 | -5.934595 |
| O | -0.902088 | -1.486923 | 3.200058 |
| N | -2.450022 | 0.073462 | 2.598703 |
| C | -3.506367 | -1.828208 | 1.338613 |
| O | -4.678337 | -1.702545 | 2.124105 |
| C | -3.876294 | -2.272295 | -0.094962 |
| O | -4.936567 | -1.477916 | -0.582901 |
| C | -3.016715 | -2.448388 | -2.489318 |
| O | -0.758079 | -3.139653 | -3.068642 |
| O | -1.212590 | -2.078134 | -5.768633 |
| C | -1.080194 | -2.875046 | -6.913358 |
| O | -3.283170 | -3.582642 | -7.519943 |
| O | -3.749122 | -4.462844 | -5.025715 |
| N | 0.308918 | -3.378672 | -7.003698 |
| C | 1.069006 | -3.593077 | -5.882960 |
| C | 2.973523 | -4.303046 | -7.217372 |
| O | 4.114519 | -4.698747 | -7.404365 |
| N | 2.119738 | -4.061784 | -8.301851 |
| C | 0.814105 | -3.621045 | -8.280058 |
| O | 0.155475 | -3.457784 | -9.292537 |
| O | -2.085018 | -0.864796 | -0.957210 |
| C | -1.608113 | -0.544337 | 0.325801 |
| O | -0.977902 | 0.699521 | 0.234478 |
| C | 0.411301 | 0.740610 | 0.480400 |
| O | 1.156310 | 0.035309 | -0.472982 |
| C | 1.059446 | 0.507548 | -1.818726 |
| C | 1.939371 | -0.417025 | -2.655783 |
| O | 1.619568 | -1.784206 | -2.464889 |
| C | 1.495814 | 1.978128 | -1.895023 |
| O | 1.385849 | 2.505447 | -3.202526 |
| C | 0.720879 | 2.839894 | -0.883322 |
| O | 1.235304 | 4.150155 | -0.834655 |
| C | 0.819257 | 2.223554 | 0.519732 |
| N | 0.057369 | 2.954217 | 1.505457 |
| C | 0.606704 | 3.535111 | 2.604047 |

| | | | |
|---|---|---|---|
| O | 1.808843 | 3.483190 | 2.857041 |
| C | -0.371394 | 4.235350 | 3.523442 |
| H | -1.841307 | 1.309520 | 4.780141 |
| H | -3.511832 | 0.264651 | 0.849914 |
| H | -1.917720 | -2.916298 | -0.693231 |
| H | -1.501468 | -1.232727 | -3.414988 |
| H | -3.186333 | -2.009284 | -5.103488 |
| H | -1.743625 | -4.958265 | -7.196331 |
| H | -1.685464 | -4.658703 | -4.777937 |
| H | 2.927810 | -4.186763 | -5.018560 |
| H | -2.878223 | 0.958604 | 2.856857 |
| H | -2.813060 | -2.573606 | 1.777909 |
| H | -4.945576 | -2.586348 | 2.421079 |
| H | -4.171829 | -3.346966 | -0.053925 |
| H | -5.555885 | -1.373286 | 0.160910 |
| H | -3.826224 | -1.766378 | -2.807477 |
| H | -3.421902 | -3.475875 | -2.551903 |
| H | 0.027450 | -2.592902 | -2.851243 |
| H | -1.251926 | -2.286105 | -7.832442 |
| H | -4.045039 | -3.943067 | -7.028713 |
| H | -3.743542 | -5.425218 | -4.903942 |
| H | 0.572906 | -3.372706 | -4.931169 |
| H | 2.502544 | -4.239710 | -9.233017 |
| H | -0.890223 | -1.322037 | 0.667645 |
| H | 0.648119 | 0.255040 | 1.448037 |
| H | 0.006014 | 0.421468 | -2.163950 |
| H | 3.006864 | -0.234180 | -2.409048 |
| H | 1.797979 | -0.185005 | -3.725327 |
| H | 1.749164 | -1.984816 | -1.522320 |
| H | 2.571768 | 2.041936 | -1.630198 |
| H | 0.457146 | 2.445903 | -3.484657 |
| H | -0.351190 | 2.847055 | -1.193997 |
| H | 1.238973 | 4.488736 | -1.744776 |
| H | 1.875839 | 2.261351 | 0.842600 |
| H | -0.948710 | 3.022982 | 1.365507 |
| C | -0.156645 | 0.183189 | 5.408844 |
| H | -1.362941 | 4.394431 | 3.068892 |
| C | 0.288238 | 1.057123 | 6.537460 |
| H | 0.052749 | 5.206628 | 3.828552 |
| C | 1.665927 | 1.682828 | 6.260913 |
| H | -0.498055 | 3.627855 | 4.438120 |
| C | 2.137996 | 2.636541 | 7.358323 |
| C | 3.473250 | 3.308809 | 7.034115 |
| C | 3.961111 | 4.274222 | 8.115247 |
| C | 5.294393 | 4.946400 | 7.783390 |
| C | 5.788108 | 5.912178 | 8.861706 |
| C | 7.124212 | 6.579513 | 8.527234 |
| C | 7.602030 | 7.546335 | 9.614743 |
| H | 7.668099 | 6.996226 | 10.576358 |
| C | 8.949818 | 8.254143 | 9.365959 |
| H | 0.460073 | -0.702835 | 5.187391 |

| | | | |
|---|---|---|---|
| H | 0.346970 | 0.454454 | 7.467326 |
| H | -0.458377 | 1.854537 | 6.717116 |
| H | 1.627673 | 2.226782 | 5.296710 |
| H | 2.413717 | 0.875335 | 6.122728 |
| H | 2.220148 | 2.087309 | 8.319081 |
| H | 1.365808 | 3.416730 | 7.522256 |
| H | 3.380726 | 3.851697 | 6.070253 |
| H | 4.244549 | 2.528827 | 6.864133 |
| H | 4.054229 | 3.729868 | 9.078123 |
| H | 3.189638 | 5.053877 | 8.286341 |
| H | 5.200053 | 5.489921 | 6.820078 |
| H | 6.064516 | 4.165711 | 7.610645 |
| H | 5.880174 | 5.369277 | 9.825663 |
| H | 5.019447 | 6.694734 | 9.033100 |
| H | 7.025520 | 7.114252 | 7.560952 |
| H | 7.885601 | 5.791337 | 8.357967 |
| H | 6.822653 | 8.321316 | 9.769067 |
| C | 10.131240 | 7.277641 | 9.316019 |
| C | 8.923701 | 9.160392 | 8.129091 |
| H | 9.111912 | 8.911841 | 10.245732 |
| H | 8.825053 | 8.578025 | 7.193589 |
| H | 8.080138 | 9.874751 | 8.168091 |
| H | 9.856226 | 9.749256 | 8.049150 |
| H | 10.079152 | 6.610391 | 8.434964 |
| H | 11.092510 | 7.820956 | 9.256160 |
| H | 10.165430 | 6.637511 | 10.217415 |

**TM-Cy-TBPA (Folded conformer, global minimum, DFT optimized, water):**
Charge: +2 | Multiplicity: 1 | Imaginary frequency: ---
$E_{elec}$(def2-TZVPP): -3564.041218 a.u.
Thermal correction to Gibbs Free Energy (quasi-RRHO) (def2-SVP): 1.061344 a.u.
Absolute Gibbs Free Energy (TZVPP): -3562.979874 a.u.

| | | | |
|---|---|---|---|
| H | 5.392072 | -4.578515 | 7.639685 |
| C | 4.466777 | -4.182473 | 8.101201 |
| C | 5.240851 | -2.125918 | 9.437126 |
| C | 4.868255 | -1.971506 | 6.948432 |
| C | 5.893228 | -1.773662 | 8.087365 |
| C | 3.867577 | -3.139655 | 7.149041 |
| C | 4.793970 | -3.605613 | 9.488195 |
| H | 4.347643 | -1.478386 | 9.510458 |
| H | 4.305576 | -1.030533 | 6.844338 |
| H | 6.776604 | -2.423841 | 7.921898 |
| H | 3.717496 | -3.627267 | 6.167772 |
| H | 5.593269 | -4.208938 | 9.952130 |
| H | 3.785412 | -5.047431 | 8.206670 |
| H | 5.417348 | -2.104039 | 5.999691 |
| H | 3.918800 | -3.675798 | 10.160365 |
| C | 2.478601 | -2.648800 | 7.606003 |
| H | 2.567522 | -2.051229 | 8.531072 |
| H | 1.847079 | -3.523104 | 7.869367 |
| C | 1.738930 | -1.827583 | 6.540691 |

| | | | |
|---|---|---|---|
| H | 2.407582 | -1.036520 | 6.150039 |
| O | 1.402137 | -2.610902 | 5.407914 |
| H | 0.861250 | -3.358894 | 5.709542 |
| C | 0.502149 | -1.125633 | 7.109320 |
| C | 0.717857 | -0.157128 | 8.280814 |
| C | -0.555421 | 0.708876 | 8.174238 |
| C | -0.746195 | 0.792309 | 6.634083 |
| H | -0.230436 | -1.896387 | 7.435806 |
| H | 0.747254 | -0.697427 | 9.243767 |
| H | -1.396918 | 0.148731 | 8.617338 |
| H | -0.275298 | 1.725139 | 6.262095 |
| O | -0.100188 | -0.319708 | 6.076753 |
| O | 1.891733 | 0.626245 | 8.230417 |
| H | 2.024967 | 1.095299 | 7.376312 |
| O | -0.491465 | 1.950959 | 8.815051 |
| H | 0.409680 | 2.299948 | 8.698382 |
| N | -2.142985 | 0.883919 | 6.179913 |
| C | -2.540817 | 1.979313 | 5.449881 |
| H | -1.757588 | 2.724025 | 5.264359 |
| C | -3.801530 | 2.150465 | 4.985238 |
| H | -4.074537 | 3.039220 | 4.410252 |
| C | -4.807360 | 1.134979 | 5.241186 |
| N | -4.312917 | 0.049874 | 5.975677 |
| H | -4.976248 | -0.699882 | 6.184728 |
| C | -3.040055 | -0.147756 | 6.467178 |
| O | -5.973467 | 1.160323 | 4.880181 |
| O | -2.735880 | -1.141243 | 7.102419 |
| O | 6.346454 | -0.418474 | 8.193873 |
| N | 6.042098 | -1.743141 | 10.599079 |
| H | 5.557452 | -1.162883 | 11.278697 |
| C | 7.298373 | -2.163372 | 10.852930 |
| O | 7.906180 | -2.992900 | 10.189792 |
| C | 8.021041 | -1.524279 | 12.048403 |
| H | 7.946419 | -2.174242 | 12.937575 |
| H | 9.083026 | -1.448578 | 11.765222 |
| N | 7.542481 | -0.159619 | 12.417919 |
| H | 6.727845 | -0.218730 | 13.048778 |
| C | 8.567354 | 0.749703 | 13.073239 |
| H | 8.840804 | 0.281424 | 14.032861 |
| H | 9.445546 | 0.744822 | 12.406430 |
| C | 7.976876 | 2.117275 | 13.236833 |
| C | 6.580191 | 4.605990 | 13.426756 |
| C | 7.904018 | 2.997458 | 12.144141 |
| C | 7.385483 | 2.525951 | 14.440944 |
| C | 6.721703 | 3.744126 | 14.546846 |
| C | 7.225654 | 4.207699 | 12.226196 |
| H | 8.500207 | 0.430512 | 9.358881 |
| C | 10.136285 | 1.256485 | 8.431868 |
| C | 11.000238 | 0.794404 | 9.587560 |
| H | 11.085020 | 1.615949 | 10.323295 |
| N | 5.838129 | 5.763526 | 13.508187 |

| | | | |
|---|---|---|---|
| C | 5.702339 | 6.680376 | 12.385876 |
| H | 5.619535 | 7.702749 | 12.801520 |
| H | 6.618300 | 6.687053 | 11.774110 |
| C | 4.784553 | 5.926361 | 14.511387 |
| H | 5.092794 | 6.628212 | 15.314279 |
| H | 4.602250 | 4.953901 | 14.999508 |
| C | 4.456164 | 6.362113 | 11.537558 |
| H | 4.712511 | 5.625792 | 10.750326 |
| H | 4.141744 | 7.276503 | 11.002732 |
| C | 3.474661 | 6.413732 | 13.857166 |
| H | 3.457372 | 7.516982 | 13.791239 |
| H | 2.623462 | 6.131556 | 14.503190 |
| C | 3.299333 | 5.847678 | 12.437397 |
| H | 2.334521 | 6.220292 | 12.041537 |
| C | 3.215258 | 4.316852 | 12.480553 |
| H | 4.198501 | 3.856686 | 12.672486 |
| H | 2.518233 | 3.974814 | 13.263790 |
| N | 2.725247 | 3.745104 | 11.184916 |
| N | 8.815982 | 0.943023 | 8.539756 |
| C | 3.087748 | 2.305026 | 10.935694 |
| H | 2.661994 | 2.039052 | 9.953195 |
| H | 4.187689 | 2.274133 | 10.861700 |
| C | 2.589591 | 1.372027 | 12.007178 |
| C | 1.674288 | -0.379215 | 13.972623 |
| C | 3.468081 | 0.874109 | 12.980282 |
| C | 1.241722 | 0.980524 | 12.037707 |
| C | 0.775929 | 0.109458 | 13.022319 |
| C | 3.017405 | -0.003628 | 13.967713 |
| H | 12.013308 | 0.567003 | 9.219087 |
| H | 10.588089 | -0.088026 | 10.105063 |
| O | 10.597442 | 1.883996 | 7.483473 |
| O | 6.048239 | 0.643167 | 6.126470 |
| O | 1.202481 | -1.209609 | 14.992131 |
| C | 1.207880 | -2.547959 | 14.809364 |
| F | 0.710195 | -3.103196 | 15.907922 |
| F | 0.460814 | -2.929135 | 13.763669 |
| F | 2.440185 | -3.035932 | 14.607208 |
| C | 5.113690 | 1.586247 | 6.639352 |
| C | 6.968448 | 0.174791 | 7.090065 |
| C | 6.917317 | 2.501895 | 8.134333 |
| C | 7.815871 | 1.362351 | 7.589945 |
| C | 5.854285 | 2.860062 | 7.082100 |
| H | 6.393883 | 2.132390 | 9.042126 |
| H | 8.361505 | 1.751754 | 6.709241 |
| H | 6.368479 | 3.283005 | 6.192409 |
| H | 4.594018 | 1.159511 | 7.524214 |
| H | 7.613029 | -0.554700 | 6.556919 |
| C | 4.048870 | 1.824356 | 5.564466 |
| H | 3.806070 | 0.851776 | 5.098771 |
| H | 4.443918 | 2.484309 | 4.762869 |
| O | 2.846166 | 2.342383 | 6.108723 |

| | | | |
|---|---|---|---|
| H | 3.072854 | 3.172761 | 6.565460 |
| O | 4.949760 | 3.858546 | 7.523277 |
| H | 4.682037 | 3.653669 | 8.433527 |
| O | 7.666088 | 3.616706 | 8.560926 |
| H | 8.145280 | 3.986072 | 7.801008 |
| H | 1.703062 | 3.872275 | 11.117011 |
| H | 3.101541 | 4.295341 | 10.396812 |
| H | 7.199514 | 0.331112 | 11.576188 |
| H | 8.363751 | 2.722042 | 11.185459 |
| H | 7.172779 | 4.827166 | 11.326198 |
| H | 6.308626 | 4.028726 | 15.519476 |
| H | 7.447703 | 1.878436 | 15.325663 |
| H | 3.702343 | -0.384592 | 14.733138 |
| H | 4.523013 | 1.175653 | 12.970273 |
| H | 0.544330 | 1.355317 | 11.277633 |
| H | -0.278103 | -0.187568 | 13.053936 |

**TM-Cy-TBPA (Extended conformer, DFT optimized, water):**
Charge: +2 | Multiplicity: 1 | Imaginary frequency: ---
$E_{elec}$(def2-TZVPP): -3564.023898 a.u.
Thermal correction to Gibbs Free Energy (quasi-RRHO) (def2-SVP): 1.057062 a.u.
Absolute Gibbs Free Energy (TZVPP): -3562.966836 a.u.

| | | | |
|---|---|---|---|
| C | 26.080337 | 6.199750 | 38.543816 |
| C | 24.396705 | 4.548372 | 37.598675 |
| C | 26.804031 | 3.865234 | 37.933853 |
| C | 25.569964 | 3.710418 | 37.045852 |
| C | 27.242271 | 5.334551 | 38.036352 |
| C | 24.824536 | 6.023776 | 37.691076 |
| N | 23.222972 | 4.366213 | 36.749485 |
| H | 23.397287 | 4.129028 | 35.770693 |
| C | 21.909042 | 4.449444 | 37.048275 |
| O | 21.037737 | 4.191398 | 36.220689 |
| C | 21.536341 | 4.939800 | 38.447524 |
| N | 20.358111 | 4.236370 | 39.027424 |
| H | 20.551389 | 3.199382 | 39.090584 |
| H | 20.298591 | 4.533592 | 40.012893 |
| C | 18.995906 | 4.461480 | 38.402284 |
| C | 18.608990 | 5.910871 | 38.359386 |
| C | 17.907916 | 8.696207 | 38.299478 |
| C | 18.050200 | 6.549903 | 39.477126 |
| C | 18.791690 | 6.686843 | 37.204789 |
| C | 18.447972 | 8.034852 | 37.165645 |
| C | 17.703315 | 7.896578 | 39.455304 |
| N | 17.627099 | 10.052498 | 38.290515 |
| C | 17.569853 | 10.810883 | 37.046164 |
| C | 16.768814 | 10.663525 | 39.299441 |
| C | 16.151118 | 10.810959 | 36.461600 |
| C | 15.300618 | 10.660766 | 38.852309 |
| C | 15.152507 | 11.367844 | 37.492374 |
| C | 13.712325 | 11.443428 | 36.987310 |
| N | 13.090971 | 10.108527 | 36.701087 |

| | | | |
|---|---|---|---|
| H | 13.650666 | 9.601974 | 35.997132 |
| H | 13.129710 | 9.517204 | 37.546359 |
| C | 11.666850 | 10.194220 | 36.225276 |
| C | 11.079008 | 8.828973 | 35.982452 |
| C | 10.020383 | 6.288756 | 35.547167 |
| C | 11.202866 | 8.213940 | 34.727784 |
| C | 10.419624 | 8.147875 | 37.016338 |
| C | 9.889184 | 6.874688 | 36.806667 |
| C | 10.677457 | 6.941116 | 34.503195 |
| O | 9.542404 | 4.991667 | 35.346699 |
| C | 8.277146 | 4.813679 | 34.908196 |
| F | 7.364130 | 5.326726 | 35.744853 |
| F | 8.064118 | 5.371449 | 33.707901 |
| F | 8.069341 | 3.506167 | 34.807900 |
| C | 28.504860 | 5.480357 | 38.909182 |
| C | 29.526865 | 6.510563 | 38.409907 |
| C | 30.199376 | 6.075113 | 37.103939 |
| C | 31.250767 | 7.001879 | 36.488052 |
| C | 32.032282 | 5.999509 | 35.612757 |
| C | 32.014611 | 4.725275 | 36.505110 |
| O | 30.880122 | 4.827181 | 37.330453 |
| N | 31.957347 | 3.459379 | 35.765446 |
| C | 30.811052 | 3.120999 | 35.086919 |
| C | 30.639659 | 1.931210 | 34.463548 |
| C | 31.699973 | 0.939459 | 34.505940 |
| N | 32.832714 | 1.373618 | 35.201945 |
| C | 33.032915 | 2.572788 | 35.855246 |
| O | 31.666934 | -0.177474 | 34.009974 |
| O | 34.061166 | 2.824982 | 36.455551 |
| H | 33.611955 | 0.713969 | 35.261090 |
| O | 32.162017 | 7.576644 | 37.402163 |
| H | 31.793958 | 7.474713 | 38.305153 |
| O | 33.312878 | 6.422855 | 35.257811 |
| H | 33.681541 | 6.861707 | 36.046510 |
| O | 30.502873 | 6.781672 | 39.416929 |
| H | 30.887151 | 5.932867 | 39.697366 |
| O | 25.189296 | 2.358666 | 36.768477 |
| C | 23.188316 | 1.161856 | 38.473697 |
| C | 25.433153 | 1.323130 | 37.678673 |
| C | 23.838987 | -0.302179 | 36.534216 |
| C | 25.309187 | -0.001665 | 36.895437 |
| C | 22.975407 | -0.195553 | 37.795646 |
| O | 24.563550 | 1.339223 | 38.787708 |
| O | 21.595455 | -0.313899 | 37.500252 |
| H | 21.446698 | -1.171866 | 37.070195 |
| O | 23.674094 | -1.549616 | 35.894060 |
| H | 24.031735 | -2.246428 | 36.469568 |
| N | 26.170838 | 0.001389 | 35.737717 |
| H | 26.068302 | 0.778369 | 35.087817 |
| C | 27.012041 | -1.018927 | 35.422673 |
| C | 27.755431 | -0.863908 | 34.111663 |

| | | | |
|---|---|---|---|
| O | 27.169523 | -2.001759 | 36.141965 |
| C | 22.391246 | 1.316898 | 39.770556 |
| O | 21.008201 | 1.517590 | 39.518623 |
| H | 20.722393 | 0.820785 | 38.895281 |
| H | 26.375738 | 7.265878 | 38.559574 |
| H | 25.857767 | 5.921754 | 39.594538 |
| H | 24.166833 | 4.171239 | 38.614077 |
| H | 26.601142 | 3.478368 | 38.952974 |
| H | 27.632175 | 3.263247 | 37.513480 |
| H | 25.817305 | 4.122967 | 36.044812 |
| H | 27.473709 | 5.670554 | 37.004005 |
| H | 23.992405 | 6.623116 | 38.104738 |
| H | 25.002188 | 6.401272 | 36.663978 |
| H | 22.353772 | 4.829720 | 39.176220 |
| H | 21.300389 | 6.018216 | 38.382408 |
| H | 18.302947 | 3.876036 | 39.030417 |
| H | 19.032608 | 4.015265 | 37.399446 |
| H | 17.868907 | 5.978001 | 40.397523 |
| H | 19.213512 | 6.224261 | 36.303586 |
| H | 18.609794 | 8.571796 | 36.227313 |
| H | 17.266003 | 8.323365 | 40.361843 |
| H | 17.873890 | 11.852235 | 37.272688 |
| H | 18.301467 | 10.435860 | 36.315138 |
| H | 16.884402 | 10.171323 | 40.276221 |
| H | 17.108104 | 11.708589 | 39.443173 |
| H | 16.125097 | 11.420120 | 35.538034 |
| H | 15.892130 | 9.771969 | 36.171294 |
| H | 14.675734 | 11.165173 | 39.613750 |
| H | 14.955590 | 9.607748 | 38.796245 |
| H | 15.420814 | 12.433426 | 37.654109 |
| H | 13.063275 | 11.932115 | 37.734128 |
| H | 13.655766 | 12.020736 | 36.048421 |
| H | 11.108660 | 10.742067 | 37.002025 |
| H | 11.678275 | 10.801542 | 35.305347 |
| H | 11.715427 | 8.735608 | 33.909689 |
| H | 10.315667 | 8.617421 | 38.002564 |
| H | 9.382385 | 6.338414 | 37.616468 |
| H | 10.781838 | 6.455945 | 33.526538 |
| H | 28.217352 | 5.770960 | 39.937623 |
| H | 29.022272 | 4.505479 | 38.996229 |
| H | 29.018199 | 7.475358 | 38.213070 |
| H | 29.405673 | 5.935254 | 36.338444 |
| H | 30.766241 | 7.789849 | 35.874529 |
| H | 31.468591 | 5.807231 | 34.678389 |
| H | 32.938958 | 4.675891 | 37.109082 |
| H | 30.016213 | 3.875322 | 35.084890 |
| H | 29.711045 | 1.701863 | 33.935019 |
| H | 22.853527 | 1.952581 | 37.770836 |
| H | 26.448913 | 1.391645 | 38.110703 |
| H | 23.480615 | 0.455840 | 35.808656 |
| H | 25.672344 | -0.805740 | 37.564049 |

| | | | |
|---|---|---|---|
| H | 23.295213 | -0.993323 | 38.507173 |
| H | 22.551954 | 0.435973 | 40.427403 |
| H | 22.763867 | 2.205765 | 40.312261 |
| H | 27.510880 | -1.724030 | 33.463449 |
| H | 28.842960 | -0.896425 | 34.298335 |
| H | 27.513152 | 0.067614 | 33.574459 |

**TM-V (Folded conformer, global minimum, DFT optimized, 1-octanol):**
Charge: 0 | Multiplicity: 1 | Imaginary frequency: ---
$E_{elec}$(def2-TZVPP): -2906.056093 a.u.
Thermal correction to Gibbs Free Energy (quasi-RRHO) (def2-SVP): 0.933530 a.u.
Absolute Gibbs Free Energy (TZVPP): -2905.122563 a.u.

| | | | |
|---|---|---|---|
| O | 0.705487 | 2.079836 | -2.461576 |
| C | 1.145367 | 1.172970 | -3.170604 |
| N | 0.343354 | 0.224076 | -3.736022 |
| C | -1.110363 | 0.231662 | -3.676972 |
| C | -1.723875 | 1.432315 | -4.448134 |
| O | -1.454380 | 1.332880 | -5.827190 |
| C | -3.236328 | 1.563411 | -4.182077 |
| O | -3.965269 | 0.560854 | -4.870055 |
| C | -3.466141 | 1.533023 | -2.670016 |
| C | -4.909447 | 1.738570 | -2.214808 |
| C | -5.098016 | 2.371679 | -0.822705 |
| O | -6.472512 | 2.396164 | -0.490865 |
| C | -4.266643 | 1.715460 | 0.298442 |
| C | -4.641483 | 2.137097 | 1.743281 |
| O | -5.145495 | 1.080569 | 2.533032 |
| C | -3.298506 | 2.672362 | 2.334737 |
| O | -3.119532 | 2.338276 | 3.676587 |
| C | -2.261908 | 2.051300 | 1.377637 |
| N | -0.989762 | 2.746623 | 1.237047 |
| C | -0.955496 | 3.997460 | 0.662406 |
| C | 0.196103 | 4.638941 | 0.354087 |
| C | 1.472625 | 3.992505 | 0.609171 |
| N | 1.341078 | 2.746326 | 1.232151 |
| C | 0.188268 | 2.065100 | 1.548546 |
| O | 0.203626 | 0.957411 | 2.056651 |
| O | 2.577559 | 4.435544 | 0.336925 |
| O | -2.896867 | 2.075665 | 0.121422 |
| O | -3.031132 | 0.251800 | -2.198794 |
| C | -1.628291 | 0.112879 | -2.225352 |
| O | -1.306478 | -1.165394 | -1.761229 |
| C | -0.888225 | -1.312782 | -0.421015 |
| C | 0.045829 | -2.533313 | -0.366360 |
| N | 1.144415 | -2.379508 | -1.292315 |
| C | 2.395295 | -2.840449 | -1.034482 |
| C | 3.417577 | -2.589203 | -2.122763 |
| O | 2.693253 | -3.397593 | 0.020871 |
| C | -0.760490 | -3.824226 | -0.584777 |
| O | 0.031643 | -4.992661 | -0.533311 |
| C | -1.901161 | -3.887480 | 0.439772 |

| | | | |
|---|---|---|---|
| O | -2.728038 | -5.004817 | 0.205642 |
| C | -2.764026 | -2.617769 | 0.366573 |
| C | -3.787624 | -2.580285 | 1.498189 |
| O | -4.733467 | -1.531218 | 1.393976 |
| O | -1.950674 | -1.450248 | 0.480534 |
| C | 2.594252 | 1.019217 | -3.474853 |
| C | 3.493860 | 1.910053 | -3.025255 |
| C | 4.956740 | 1.905217 | -3.333830 |
| C | 5.896455 | 2.170682 | -2.145312 |
| C | 6.021527 | 1.043407 | -1.112544 |
| C | 4.865244 | 0.902915 | -0.117789 |
| C | 5.113470 | -0.188659 | 0.925727 |
| C | 4.014114 | -0.313115 | 1.980970 |
| C | 4.275859 | -1.424074 | 3.002364 |
| C | 3.243595 | -1.524285 | 4.132241 |
| C | 1.847121 | -1.958773 | 3.678143 |
| C | 0.818001 | -2.162936 | 4.804300 |
| C | -0.462476 | -2.793522 | 4.246525 |
| C | 0.497501 | -0.865476 | 5.556398 |
| H | 0.774650 | -0.434253 | -4.379658 |
| H | -1.445276 | -0.693377 | -4.185990 |
| H | -1.240091 | 2.359411 | -4.086555 |
| H | -2.110818 | 0.711407 | -6.188278 |
| H | -3.572454 | 2.536737 | -4.590497 |
| H | -3.920734 | -0.254662 | -4.342016 |
| H | -2.828448 | 2.329465 | -2.232633 |
| H | -5.437262 | 2.385613 | -2.939418 |
| H | -5.425861 | 0.757508 | -2.237039 |
| H | -4.781663 | 3.434199 | -0.867802 |
| H | -6.810208 | 1.485941 | -0.505099 |
| H | -4.360943 | 0.611069 | 0.227102 |
| H | -5.403951 | 2.936129 | 1.742836 |
| H | -4.880538 | 0.210203 | 2.164683 |
| H | -3.268694 | 3.776224 | 2.227362 |
| H | -3.742718 | 1.602418 | 3.842437 |
| H | -2.025178 | 1.020917 | 1.707032 |
| H | -1.932547 | 4.444397 | 0.448020 |
| H | 0.184994 | 5.629591 | -0.107819 |
| H | 2.208957 | 2.248591 | 1.443915 |
| H | -1.169293 | 0.890210 | -1.580920 |
| H | -0.341683 | -0.408705 | -0.091298 |
| H | 0.473993 | -2.559909 | 0.654769 |
| H | 0.936049 | -1.974987 | -2.203793 |
| H | 4.100760 | -3.451942 | -2.181666 |
| H | 2.962989 | -2.410618 | -3.111046 |
| H | 4.020522 | -1.702483 | -1.854299 |
| H | -1.210767 | -3.808365 | -1.598506 |
| H | 0.656631 | -4.914597 | 0.207617 |
| H | -1.440364 | -3.937633 | 1.455508 |
| H | -2.141049 | -5.774930 | 0.126606 |
| H | -3.297186 | -2.616095 | -0.610188 |

| | | | |
|---|---|---|---|
| H | -4.292124 | -3.564842 | 1.540276 |
| H | -3.254613 | -2.435841 | 2.456365 |
| H | -5.356231 | -1.728236 | 0.676910 |
| H | 2.911428 | 0.167420 | -4.093826 |
| H | 3.119732 | 2.745472 | -2.411653 |
| H | 5.238340 | 0.963967 | -3.844870 |
| H | 5.124575 | 2.716831 | -4.073932 |
| H | 5.591586 | 3.109460 | -1.638895 |
| H | 6.900522 | 2.370820 | -2.564712 |
| H | 6.173858 | 0.078936 | -1.641138 |
| H | 6.952088 | 1.210216 | -0.533430 |
| H | 4.706078 | 1.875708 | 0.392542 |
| H | 3.917823 | 0.688802 | -0.651551 |
| H | 6.083520 | 0.002994 | 1.430568 |
| H | 5.234107 | -1.164424 | 0.410667 |
| H | 3.043644 | -0.482095 | 1.473573 |
| H | 3.909442 | 0.654408 | 2.516757 |
| H | 4.337006 | -2.396614 | 2.471378 |
| H | 5.277747 | -1.263090 | 3.449647 |
| H | 3.613105 | -2.246799 | 4.888396 |
| H | 3.190309 | -0.546728 | 4.653102 |
| H | 1.435297 | -1.221866 | 2.960505 |
| H | 1.944167 | -2.908653 | 3.113294 |
| H | 1.256141 | -2.877439 | 5.534907 |
| H | -0.927312 | -2.134873 | 3.487815 |
| H | -0.256419 | -3.764942 | 3.759658 |
| H | -1.211661 | -2.969412 | 5.040747 |
| H | 1.395147 | -0.416544 | 6.017962 |
| H | -0.232192 | -1.045999 | 6.367513 |
| H | 0.057950 | -0.110573 | 4.876201 |

**TM-V (Extended conformer, DFT optimized, 1-octanol):**
Charge: 0 | Multiplicity: 1 | Imaginary frequency: ---
$E_{elec}$(def2-TZVPP): -2906.042168 a.u.
Thermal correction to Gibbs Free Energy (quasi-RRHO) (def2-SVP): 0.927700 a.u.
Absolute Gibbs Free Energy (TZVPP): -2905.114468 a.u.

| | | | |
|---|---|---|---|
| C | -0.079122 | 2.296436 | 3.468467 |
| C | -0.071540 | 1.113069 | 2.562850 |
| C | -1.595150 | -0.505084 | 1.420316 |
| C | -1.149203 | -2.632588 | -0.487718 |
| C | -1.371127 | -3.265972 | -2.957257 |
| C | -1.981255 | -4.283776 | -3.929649 |
| C | -1.652443 | -6.104715 | -5.449031 |
| C | -1.291341 | -5.653972 | -4.014761 |
| C | 1.711374 | -3.485871 | -7.107055 |
| O | 0.959888 | 0.649444 | 2.071017 |
| N | -1.304681 | 0.604901 | 2.304778 |
| C | -1.099319 | -1.869170 | 1.933716 |
| O | -1.665771 | -2.096502 | 3.211723 |
| C | -1.533558 | -2.985133 | 0.956479 |
| O | -2.927913 | -3.183292 | 1.050011 |

| | | | |
|---|---|---|---|
| C | -1.704796 | -3.623899 | -1.504050 |
| O | 0.028045 | -3.169920 | -3.147390 |
| O | -1.905492 | -3.756576 | -5.260287 |
| C | -1.699313 | -4.765682 | -6.210874 |
| O | -2.939192 | -6.669962 | -5.464761 |
| O | -1.797696 | -6.535251 | -3.037164 |
| N | -0.466886 | -4.476291 | -6.977101 |
| C | 0.575937 | -3.777630 | -6.424634 |
| C | 1.861921 | -3.918883 | -8.484410 |
| O | 2.825951 | -3.723712 | -9.208047 |
| N | 0.748884 | -4.626631 | -8.959704 |
| C | -0.412775 | -4.944943 | -8.289453 |
| O | -1.316089 | -5.583953 | -8.799256 |
| O | -1.614079 | -1.329585 | -0.831250 |
| C | -1.072870 | -0.314911 | -0.021042 |
| O | -1.510180 | 0.906746 | -0.534194 |
| C | -0.530098 | 1.764158 | -1.080463 |
| O | 0.064403 | 1.245816 | -2.237658 |
| C | -0.811533 | 1.015088 | -3.342766 |
| C | 0.066390 | 0.459154 | -4.461856 |
| O | 0.851071 | -0.639083 | -4.030267 |
| C | -1.525189 | 2.318642 | -3.732377 |
| O | -2.434259 | 2.137834 | -4.800759 |
| C | -2.218479 | 2.960411 | -2.517903 |
| O | -2.729011 | 4.232251 | -2.840374 |
| C | -1.220064 | 3.109212 | -1.360319 |
| N | -1.819647 | 3.644588 | -0.160979 |
| C | -1.404892 | 4.794834 | 0.436120 |
| O | -0.491259 | 5.484791 | -0.002909 |
| C | -2.160131 | 5.177723 | 1.693089 |
| H | -1.040369 | 2.615540 | 3.897959 |
| H | -2.696919 | -0.557654 | 1.339124 |
| H | -0.036786 | -2.646862 | -0.545187 |
| H | -1.853248 | -2.294549 | -3.190687 |
| H | -3.047015 | -4.439818 | -3.656385 |
| H | -0.895739 | -6.792594 | -5.882727 |
| H | -0.195304 | -5.525296 | -3.916386 |
| H | 2.519061 | -2.920034 | -6.635142 |
| H | -2.100695 | 1.043941 | 2.758577 |
| H | 0.008245 | -1.848109 | 1.989014 |
| H | -1.124158 | -2.753837 | 3.675162 |
| H | -0.981273 | -3.915034 | 1.229797 |
| H | -3.141402 | -3.143031 | 1.998779 |
| H | -2.802413 | -3.683683 | -1.389251 |
| H | -1.303793 | -4.625690 | -1.260942 |
| H | 0.250257 | -2.260431 | -3.440875 |
| H | -2.520771 | -4.793386 | -6.949386 |
| H | -3.047645 | -7.120847 | -4.606935 |
| H | -1.126426 | -7.207654 | -2.843673 |
| H | 0.427297 | -3.458793 | -5.386840 |
| H | 0.800376 | -4.963403 | -9.923494 |

| | | | |
|---|---|---|---|
| H | 0.037660 | -0.370195 | -0.037805 |
| H | 0.303047 | 1.903472 | -0.363267 |
| H | -1.569329 | 0.252690 | -3.058440 |
| H | 0.719874 | 1.267178 | -4.854373 |
| H | -0.571420 | 0.111654 | -5.292688 |
| H | 1.393324 | -0.329233 | -3.284970 |
| H | -0.765630 | 3.039762 | -4.099245 |
| H | -3.104744 | 1.487686 | -4.531419 |
| H | -3.039063 | 2.276281 | -2.193211 |
| H | -3.273891 | 4.126190 | -3.637013 |
| H | -0.434821 | 3.824214 | -1.667923 |
| H | -2.561621 | 3.102812 | 0.275937 |
| C | 1.049014 | 2.975226 | 3.730080 |
| H | -2.902597 | 4.427299 | 2.010374 |
| C | 1.158758 | 4.180242 | 4.621961 |
| H | -2.675557 | 6.139217 | 1.520166 |
| C | 2.563258 | 4.793372 | 4.676459 |
| H | -1.435439 | 5.338633 | 2.510301 |
| C | 2.830445 | 5.628680 | 5.931938 |
| C | 4.315625 | 5.891047 | 6.202215 |
| C | 4.621151 | 6.359935 | 7.628237 |
| C | 6.095253 | 6.228996 | 8.015983 |
| C | 6.419261 | 6.764575 | 9.412214 |
| C | 7.848682 | 6.456177 | 9.875264 |
| C | 8.354077 | 7.397537 | 10.971041 |
| C | 9.956551 | 5.784981 | 12.140720 |
| C | 9.793291 | 7.145602 | 11.453348 |
| H | 1.974403 | 2.601359 | 3.262520 |
| H | 0.866727 | 3.868210 | 5.647043 |
| H | 0.402497 | 4.937057 | 4.330613 |
| H | 2.760918 | 5.383271 | 3.759175 |
| H | 3.297727 | 3.962128 | 4.658466 |
| H | 2.417716 | 5.076409 | 6.802013 |
| H | 2.266393 | 6.583135 | 5.893693 |
| H | 4.725347 | 6.612445 | 5.466293 |
| H | 4.866335 | 4.943084 | 6.025492 |
| H | 4.014988 | 5.763298 | 8.341280 |
| H | 4.288848 | 7.410363 | 7.762612 |
| H | 6.728741 | 6.754141 | 7.271122 |
| H | 6.386616 | 5.159388 | 7.953001 |
| H | 5.694102 | 6.359695 | 10.147812 |
| H | 6.260435 | 7.863174 | 9.414631 |
| H | 8.534880 | 6.535986 | 9.005610 |
| H | 7.906860 | 5.400103 | 10.204080 |
| H | 8.285899 | 8.438246 | 10.593197 |
| H | 10.449196 | 7.152019 | 10.555641 |
| C | 10.258887 | 8.277544 | 12.375926 |
| H | 7.670104 | 7.353350 | 11.846454 |
| H | 9.633030 | 8.328777 | 13.288185 |
| H | 11.305872 | 8.131651 | 12.699982 |
| H | 10.196821 | 9.261373 | 11.875051 |

| | | | |
|---|---|---|---|
| H | 9.306986 | 5.713759 | 13.035144 |
| H | 9.697961 | 4.943679 | 11.473248 |
| H | 10.999001 | 5.629665 | 12.474879 |

**TM-Cy-TBPA (Folded conformer, global minimum, DFT optimized, 1-octanol):**
Charge: +2 | Multiplicity: 1 | Imaginary frequency: ---
$E_{elec}$(def2-TZVPP): -3564.023290 a.u.
Thermal correction to Gibbs Free Energy (quasi-RRHO) (def2-SVP): 1.059694 a.u.
Absolute Gibbs Free Energy (TZVPP): -3562.963596 a.u.

| | | | |
|---|---|---|---|
| H | 5.113212 | -4.846115 | 7.805460 |
| C | 4.265607 | -4.343401 | 8.309583 |
| C | 5.225858 | -2.283095 | 9.501841 |
| C | 4.775072 | -2.228818 | 7.020455 |
| C | 5.841097 | -2.021973 | 8.118284 |
| C | 3.698427 | -3.298110 | 7.337598 |
| C | 4.754923 | -3.745451 | 9.642176 |
| H | 4.344621 | -1.617513 | 9.565708 |
| H | 4.282417 | -1.259971 | 6.842977 |
| H | 6.691328 | -2.713234 | 7.955602 |
| H | 3.463176 | -3.823043 | 6.393405 |
| H | 5.589060 | -4.355515 | 10.034901 |
| H | 3.517564 | -5.134203 | 8.503992 |
| H | 5.289908 | -2.483856 | 6.077203 |
| H | 3.959955 | -3.776768 | 10.410184 |
| C | 2.370935 | -2.672615 | 7.813520 |
| H | 2.534586 | -2.021506 | 8.691842 |
| H | 1.687548 | -3.477658 | 8.155803 |
| C | 1.659505 | -1.871569 | 6.712415 |
| H | 2.364696 | -1.139880 | 6.272176 |
| O | 1.270938 | -2.695762 | 5.628177 |
| H | 0.667579 | -3.377560 | 5.965532 |
| C | 0.465586 | -1.077196 | 7.247394 |
| C | 0.733466 | -0.034462 | 8.342318 |
| C | -0.515198 | 0.864601 | 8.186805 |
| C | -0.741436 | 0.830421 | 6.649112 |
| H | -0.293474 | -1.790065 | 7.638953 |
| H | 0.754579 | -0.507858 | 9.339978 |
| H | -1.363308 | 0.374786 | 8.694820 |
| H | -0.268804 | 1.725395 | 6.194431 |
| O | -0.117652 | -0.326409 | 6.165454 |
| O | 1.935565 | 0.698531 | 8.229196 |
| H | 2.077627 | 1.095946 | 7.342319 |
| O | -0.402293 | 2.154476 | 8.720406 |
| H | 0.470004 | 2.505091 | 8.472503 |
| N | -2.146487 | 0.904543 | 6.220165 |
| C | -2.544533 | 1.928975 | 5.391642 |
| H | -1.755853 | 2.641237 | 5.121527 |
| C | -3.810367 | 2.070700 | 4.933345 |
| H | -4.084450 | 2.902124 | 4.278691 |
| C | -4.824637 | 1.098040 | 5.303903 |
| N | -4.328446 | 0.081632 | 6.132104 |

| | | | |
|---|---|---|---|
| H | -4.996675 | -0.636346 | 6.421417 |
| C | -3.049851 | -0.085142 | 6.617510 |
| O | -5.995279 | 1.104740 | 4.963224 |
| O | -2.743312 | -1.017932 | 7.337985 |
| O | 6.345983 | -0.685075 | 8.160599 |
| N | 6.062192 | -1.853820 | 10.625488 |
| H | 5.540847 | -1.472488 | 11.412350 |
| C | 7.397542 | -1.991942 | 10.736796 |
| O | 8.115349 | -2.591685 | 9.945167 |
| C | 8.054996 | -1.294367 | 11.939010 |
| H | 7.946774 | -1.895413 | 12.859196 |
| H | 9.127171 | -1.200880 | 11.709899 |
| N | 7.517122 | 0.074359 | 12.210275 |
| H | 6.677569 | 0.019467 | 12.806986 |
| C | 8.489100 | 1.036507 | 12.870364 |
| H | 8.833012 | 0.547640 | 13.796721 |
| H | 9.340992 | 1.126965 | 12.175529 |
| C | 7.813008 | 2.347850 | 13.132094 |
| C | 6.352689 | 4.766488 | 13.578162 |
| C | 7.662516 | 3.304414 | 12.113598 |
| C | 7.250419 | 2.636050 | 14.385177 |
| C | 6.549691 | 3.815101 | 14.613945 |
| C | 6.953565 | 4.482516 | 12.322664 |
| H | 8.707739 | -0.513912 | 8.816477 |
| C | 10.076555 | 1.005213 | 8.711644 |
| C | 11.074605 | 0.183456 | 9.491913 |
| H | 11.300494 | 0.697464 | 10.443195 |
| N | 5.599952 | 5.898692 | 13.791886 |
| C | 5.459933 | 6.950871 | 12.797171 |
| H | 5.418113 | 7.915568 | 13.338513 |
| H | 6.359374 | 7.009389 | 12.164306 |
| C | 4.577502 | 5.952073 | 14.836822 |
| H | 4.914482 | 6.551536 | 15.708198 |
| H | 4.397282 | 4.929854 | 15.212072 |
| C | 4.182335 | 6.774899 | 11.953466 |
| H | 4.405854 | 6.162917 | 11.056882 |
| H | 3.864606 | 7.761143 | 11.569863 |
| C | 3.256685 | 6.525066 | 14.280173 |
| H | 3.246545 | 7.627854 | 14.354726 |
| H | 2.417618 | 6.168882 | 14.905168 |
| C | 3.045908 | 6.145666 | 12.803147 |
| H | 2.071676 | 6.560963 | 12.479384 |
| C | 2.972807 | 4.621182 | 12.655015 |
| H | 3.957064 | 4.153208 | 12.815572 |
| H | 2.263239 | 4.178412 | 13.373750 |
| N | 2.524675 | 4.202186 | 11.284671 |
| N | 8.871685 | 0.434429 | 8.477576 |
| C | 3.021387 | 2.852878 | 10.836641 |
| H | 2.601882 | 2.673255 | 9.832703 |
| H | 4.116319 | 2.950455 | 10.747610 |
| C | 2.665372 | 1.729019 | 11.772501 |

| | | | |
|---|---|---|---|
| C | 2.048724 | -0.418471 | 13.438959 |
| C | 3.634018 | 1.199626 | 12.637134 |
| C | 1.375262 | 1.177347 | 11.769997 |
| C | 1.059537 | 0.105189 | 12.604496 |
| C | 3.333083 | 0.125591 | 13.476096 |
| H | 12.016294 | 0.136938 | 8.917880 |
| H | 10.730755 | -0.841419 | 9.703424 |
| O | 10.349979 | 2.142994 | 8.315183 |
| O | 6.089361 | 0.352514 | 6.081629 |
| O | 1.725134 | -1.466022 | 14.300583 |
| C | 1.932362 | -2.740136 | 13.896665 |
| F | 1.525149 | -3.539897 | 14.871432 |
| F | 1.256245 | -3.042667 | 12.780396 |
| F | 3.227245 | -2.995506 | 13.648578 |
| C | 5.207563 | 1.347945 | 6.591418 |
| C | 6.986365 | -0.149857 | 7.039246 |
| C | 7.066156 | 2.161381 | 8.094956 |
| C | 7.902759 | 0.985274 | 7.539080 |
| C | 6.020756 | 2.572990 | 7.040074 |
| H | 6.511275 | 1.793991 | 8.991221 |
| H | 8.456350 | 1.361397 | 6.653805 |
| H | 6.559432 | 2.979968 | 6.159426 |
| H | 4.658159 | 0.949290 | 7.471661 |
| H | 7.587934 | -0.920665 | 6.512826 |
| C | 4.165416 | 1.640648 | 5.507989 |
| H | 3.849104 | 0.678216 | 5.065510 |
| H | 4.611911 | 2.247749 | 4.691895 |
| O | 3.005675 | 2.263100 | 6.035704 |
| H | 3.298026 | 3.087939 | 6.463619 |
| O | 5.168171 | 3.613621 | 7.492490 |
| H | 4.867506 | 3.396272 | 8.388888 |
| O | 7.853259 | 3.268051 | 8.471002 |
| H | 8.800027 | 3.055674 | 8.306829 |
| H | 1.494907 | 4.245749 | 11.227661 |
| H | 2.850849 | 4.890993 | 10.587507 |
| H | 7.197653 | 0.510560 | 11.329679 |
| H | 8.087656 | 3.125953 | 11.116117 |
| H | 6.850579 | 5.175701 | 11.482020 |
| H | 6.166191 | 4.007545 | 15.620695 |
| H | 7.374314 | 1.926565 | 15.214308 |
| H | 4.086578 | -0.280612 | 14.159988 |
| H | 4.643330 | 1.630172 | 12.656495 |
| H | 0.608955 | 1.577588 | 11.094119 |
| H | 0.052144 | -0.325475 | 12.606544 |

**TM-Cy-TBPA (Extended conformer, DFT optimized, 1-octanol):**
Charge: +2 | Multiplicity: 1 | Imaginary frequency: ---
$E_{elec}$(def2-TZVPP): -3564.014531 a.u.
Thermal correction to Gibbs Free Energy (quasi-RRHO) (def2-SVP): 1.055073 a.u.
Absolute Gibbs Free Energy (TZVPP): -3562.959458 a.u.

| | | | |
|---|---|---|---|
| C | 26.060796 | 4.695882 | 38.395752 |

| | | | |
|---|---|---|---|
| C | 24.608377 | 2.729305 | 37.669869 |
| C | 27.130359 | 2.477106 | 38.003980 |
| C | 25.907345 | 2.031652 | 37.203265 |
| C | 27.337136 | 3.994607 | 37.915789 |
| C | 24.817360 | 4.251556 | 37.620548 |
| N | 23.542004 | 2.226125 | 36.794231 |
| H | 23.784110 | 1.353540 | 36.318864 |
| C | 22.329089 | 2.667492 | 36.439049 |
| O | 21.629929 | 2.074409 | 35.612615 |
| C | 21.721317 | 3.904982 | 37.116962 |
| N | 20.350918 | 4.065104 | 36.557434 |
| H | 20.234407 | 3.179547 | 36.000595 |
| H | 19.650926 | 4.026682 | 37.310457 |
| C | 20.107375 | 5.235564 | 35.649142 |
| C | 19.078260 | 6.226049 | 36.146064 |
| C | 17.193040 | 8.235316 | 36.923236 |
| C | 18.398859 | 6.150526 | 37.363005 |
| C | 18.793517 | 7.325160 | 35.315646 |
| C | 17.881396 | 8.302513 | 35.687026 |
| C | 17.477039 | 7.129949 | 37.749414 |
| N | 16.277344 | 9.226399 | 37.280910 |
| C | 16.647025 | 10.626217 | 37.078044 |
| C | 15.419881 | 9.017884 | 38.437543 |
| C | 15.418126 | 11.521188 | 36.942426 |
| C | 14.153409 | 9.873626 | 38.348635 |
| C | 14.491476 | 11.356808 | 38.154108 |
| C | 13.270824 | 12.274762 | 38.140946 |
| N | 12.346177 | 12.040511 | 36.976116 |
| H | 12.730164 | 12.483495 | 36.128695 |
| H | 12.331605 | 11.031231 | 36.757845 |
| C | 10.906925 | 12.493659 | 37.175470 |
| C | 9.938005 | 11.366389 | 36.939759 |
| C | 8.192468 | 9.240389 | 36.494444 |
| C | 9.416932 | 11.132576 | 35.658372 |
| C | 9.572514 | 10.515452 | 37.994038 |
| C | 8.701401 | 9.447771 | 37.777382 |
| C | 8.544272 | 10.068848 | 35.427836 |
| O | 7.367119 | 8.141161 | 36.260578 |
| C | 6.026422 | 8.283447 | 36.375270 |
| F | 5.654848 | 8.667507 | 37.604053 |
| F | 5.530662 | 9.184483 | 35.516394 |
| F | 5.477302 | 7.104404 | 36.115768 |
| C | 28.587216 | 4.409117 | 38.704213 |
| C | 29.063961 | 5.853554 | 38.505310 |
| C | 29.640427 | 6.115366 | 37.107085 |
| C | 29.995791 | 7.575456 | 36.785082 |
| C | 31.171138 | 7.393029 | 35.801940 |
| C | 31.880265 | 6.153693 | 36.401717 |
| O | 30.862001 | 5.362000 | 36.973129 |
| N | 32.628572 | 5.351061 | 35.438157 |
| C | 31.957639 | 4.655226 | 34.460092 |

| | | | |
|---|---|---|---|
| C | 32.581232 | 3.883708 | 33.539227 |
| C | 34.030751 | 3.764822 | 33.560874 |
| N | 34.626537 | 4.497620 | 34.595324 |
| C | 34.020954 | 5.283280 | 35.553340 |
| O | 34.713271 | 3.110249 | 32.790693 |
| O | 34.650237 | 5.864039 | 36.417105 |
| H | 35.645458 | 4.444562 | 34.660855 |
| O | 30.427519 | 8.359817 | 37.878958 |
| H | 30.511802 | 7.782429 | 38.666970 |
| O | 31.988615 | 8.513843 | 35.676248 |
| H | 32.025139 | 8.919619 | 36.562840 |
| O | 30.029605 | 6.202723 | 39.496896 |
| H | 30.778463 | 5.587387 | 39.406513 |
| O | 25.768785 | 0.607514 | 37.129515 |
| C | 23.547401 | -1.210260 | 37.780236 |
| C | 25.638316 | -0.140992 | 38.317577 |
| C | 25.674518 | -2.416452 | 37.180966 |
| C | 26.414347 | -1.463337 | 38.137942 |
| C | 24.227882 | -2.579388 | 37.659419 |
| O | 24.295161 | -0.373171 | 38.657391 |
| O | 23.472712 | -3.370157 | 36.762988 |
| H | 23.948732 | -4.206715 | 36.630079 |
| O | 26.316376 | -3.665306 | 37.040241 |
| H | 26.442505 | -4.054826 | 37.921633 |
| N | 27.775590 | -1.212462 | 37.734463 |
| H | 27.917894 | -0.687484 | 36.874305 |
| C | 28.856733 | -1.717926 | 38.390032 |
| C | 30.201392 | -1.366263 | 37.787339 |
| O | 28.760781 | -2.392801 | 39.409399 |
| C | 22.124102 | -1.284311 | 38.344469 |
| O | 21.209348 | -1.856473 | 37.442962 |
| H | 21.596544 | -2.695305 | 37.136224 |
| H | 26.150655 | 5.793815 | 38.304963 |
| H | 25.920412 | 4.484486 | 39.475817 |
| H | 24.395035 | 2.430567 | 38.717844 |
| H | 27.028791 | 2.198594 | 39.071609 |
| H | 28.016069 | 1.940908 | 37.615188 |
| H | 26.064182 | 2.325413 | 36.145784 |
| H | 27.486339 | 4.247096 | 36.844341 |
| H | 23.950156 | 4.775413 | 38.054953 |
| H | 24.897524 | 4.569423 | 36.561634 |
| H | 21.663533 | 3.761645 | 38.209740 |
| H | 22.302994 | 4.817206 | 36.918592 |
| H | 19.794737 | 4.822718 | 34.673792 |
| H | 21.077201 | 5.736194 | 35.485544 |
| H | 18.573043 | 5.337779 | 38.081208 |
| H | 19.295924 | 7.415280 | 34.343428 |
| H | 17.681753 | 9.123241 | 34.989464 |
| H | 17.000627 | 7.026915 | 38.728799 |
| H | 17.267009 | 10.983104 | 37.934697 |
| H | 17.271813 | 10.724267 | 36.177302 |

| | | | |
|---|---|---|---|
| H | 15.123002 | 7.956273 | 38.482533 |
| H | 15.955188 | 9.245409 | 39.390278 |
| H | 15.745750 | 12.573277 | 36.843945 |
| H | 14.888598 | 11.253096 | 36.005660 |
| H | 13.556434 | 9.728609 | 39.268424 |
| H | 13.543046 | 9.493044 | 37.503308 |
| H | 15.056222 | 11.688094 | 39.051170 |
| H | 12.671436 | 12.106149 | 39.051661 |
| H | 13.559928 | 13.338963 | 38.111194 |
| H | 10.828791 | 12.881039 | 38.203317 |
| H | 10.731891 | 13.329467 | 36.480277 |
| H | 9.692470 | 11.790981 | 34.825147 |
| H | 9.968876 | 10.688054 | 39.002467 |
| H | 8.420653 | 8.777623 | 38.597244 |
| H | 8.143309 | 9.879976 | 34.426027 |
| H | 28.395501 | 4.272879 | 39.786835 |
| H | 29.422678 | 3.728352 | 38.446043 |
| H | 28.217956 | 6.552212 | 38.654375 |
| H | 28.916714 | 5.765796 | 36.341668 |
| H | 29.139886 | 8.084937 | 36.299866 |
| H | 30.777498 | 7.139043 | 34.797055 |
| H | 32.617253 | 6.474921 | 37.161822 |
| H | 30.865881 | 4.750828 | 34.478283 |
| H | 32.015578 | 3.341534 | 32.776952 |
| H | 23.485274 | -0.767751 | 36.761118 |
| H | 26.059984 | 0.406356 | 39.180888 |
| H | 25.652770 | -1.974425 | 36.163867 |
| H | 26.462036 | -1.944346 | 39.134016 |
| H | 24.249274 | -3.044391 | 38.674510 |
| H | 22.153745 | -1.821482 | 39.320527 |
| H | 21.788708 | -0.251233 | 38.558239 |
| H | 30.747516 | -0.711831 | 38.490361 |
| H | 30.131076 | -0.859082 | 36.811192 |
| H | 30.795152 | -2.289441 | 37.674209 |

**Molecular docking**

In this study, molecular docking calculations were performed to model the binding modes of tunicamycin V (TM-V) and TM-Cy-TBPA within the catalytic site of DPAGT1. Docking studies were conducted using the Maestro interface implemented in the Schrödinger 2022-3 software suite (Schrödinger, LLC, New York, NY, USA) or using Vina 1.2.7[7] employing empirical pKa values for establishing the protonation state of the ligands. The ligand structures of TM-V and TM-Cy-TBPA were obtained from the PubChem database, while the cryo-EM structure of human DPAGT1 was retrieved from the Protein Data Bank (PDB ID: 9ZNN and 6BW5).

**Figure S5**. Superimposition of TM-V and TM-Cy-TBPA within the DPAGT1 catalytic domain and clarification of the role of the C8’,9’-diol interactions of TM-V.

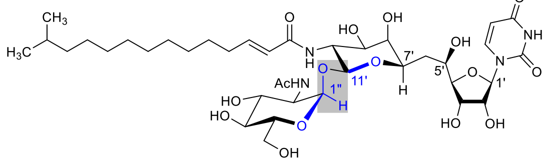

**Tunicamycin V (TM-V, 1)**

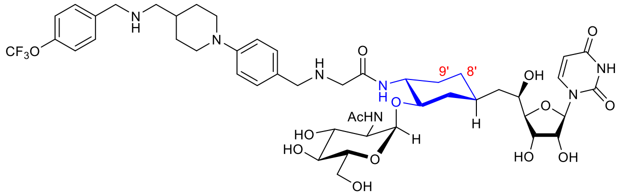

**TM-Cy-TBPA (4)**

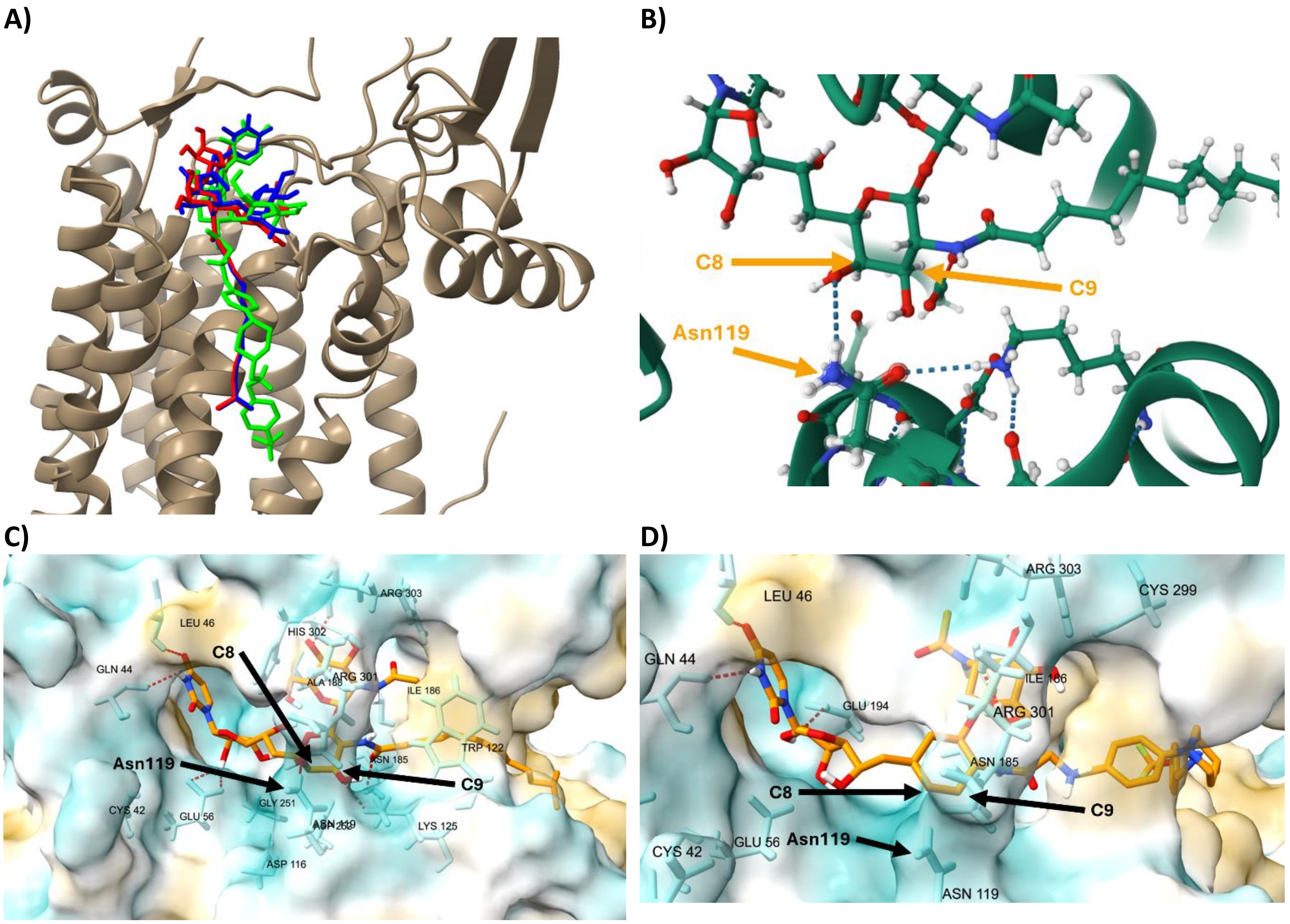


**A):** Superimposition of the crystal structure of TM-V (red) with the most stable docked poses of TM-V (blue) and TM-Cy-TBPA (green) within the binding cleft of DPAGT1.; **B):** Crystal structure of DPAGT1 (PDB ID: 6BW5) showing the hydrogen-bonding interaction between the side chain of Asn119 and the C8'-hydroxyl group of TM-V.; **C)**: Molecular docking analysis of TM-V within the catalytic domain of DPAGT1.; **D)**: Molecular docking analysis of TM-Cy-TBPA within the catalytic domain of DPAGT1. These docking experiments were run using Vina 1.2.7 using the Vinardo scoring function.

### The synthesis of compound 15 for structural characterization described in Scheme 1.

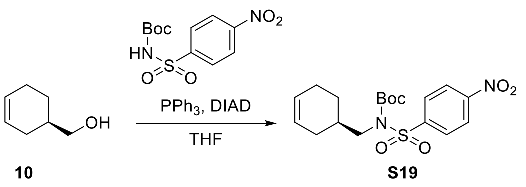

***tert*-Butyl (*R*)-(cyclohex-3-en-1-ylmethyl)((4-nitrophenyl)sulfonyl)carbamate (S19)**

To a stirred solution of $PPh_3$ (7.6 g, 29.0 mmol), *tert*-Butyl ((4-nitrophenyl)sulfonyl)carbamate (2.5 g, 22.3 mmol) in THF (22 mL) was added DIAD (29.0 mmol, 5.7 mL). After being stirred at r.t. for 1 h, the reaction mixture was evaporated, and the crude mixture was purified by silica gel column chromatography (hexanes/EtOAc/$CH_2Cl_2$ = 8/1/1) to afford **S19** (10.0 g, 95%): $[\alpha]_D^{21}$ + 1.45 (*c* = 0.6, MeOH); IR (thin film) $\nu_{max}$ = 1731, 1534, 1351, 1154 cm$^{-1}$; $^1$H NMR (400 MHz, $CDCl_3$) δ 8.39 – 8.33 (m, 2H), 8.13 – 8.07 (m, 2H), 5.74 – 5.63 (m, 2H), 3.84 – 3.70 (m, 2H), 2.20 – 2.06 (m, 4H), 1.81 (ddt, *J* = 13.0, 10.8, 5.5 Hz, 2H), 1.35 (s, 9H); $^{13}$C NMR (100 MHz, $CDCl_3$) δ 150.74, 150.24, 145.82, 129.33, 127.08, 125.37, 123.88, 85.18, 77.34, 77.23, 77.03, 76.71, 52.70, 34.37, 29.01, 27.88, 26.15, 24.74; HRMS (ESI$^+$) *m/z* calcd for $C_{18}H_{24}N_2O_6S$ [M+H] 396.1355, found: 397.1342.

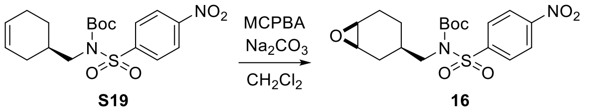

***tert*-Butyl(((1*R*,3*R*,6*S*)-7-oxabicyclo[4.1.0]heptan-3-yl)methyl)((4 nitrophenyl)sulfonyl)carbamate (16)**

To a stirred solution of **S19** (10 g, 25.2 mmol) and $Na_2CO_3$ (8.0 g, 75.0 mmol) in $CH_2Cl_2$ (300 mL) was added *m*CPBA (8.7 g, 50.5 mmol). After being stirred at r.t. for 1 h, the reaction mixture was filtered and evaporated. The crude mixture was purified by silica gel column chromatography (hexanes/EtOAc = 4/1 - 3/1) to afford **16** (10.0 g, 96%): $[\alpha]_D^{21}$ + 0.013 (*c* = 0.2, MeOH); IR (thin film) $\nu_{max}$ = 1730, 1533, 1351, 1151 cm$^{-1}$; $^1$H NMR (400 MHz, $CDCl_3$) δ 8.33 – 8.27 (m, 2H), 8.05 – 7.99 (m, 2H), 3.62 (ddd, *J* = 15.5, 7.4, 1.7 Hz, 2H), 3.18 – 3.08 (m, 2H), 2.14 (dtd, *J* = 14.3, 5.1, 2.0 Hz, 1H), 2.09 – 1.98 (m, 1H), 1.87 – 1.63 (m, 2H), 1.59 – 1.31 (m, 3H), 1.28 (d, *J* = 1.1 Hz, 9H), 1.21 – 1.09 (m, 1H); $^{13}$C NMR (100 MHz, $CDCl_3$) δ 170.39, 77.35, 77.24, 77.04, 76.72, 70.63, 62.01, 42.82, 32.79, 30.00, 27.88, 24.39, 23.35; HRMS (ESI$^+$) *m/z* calcd for $C_{18}H_{24}N_2O_7S$ [M+H] 412.1304, found: 413.1305.

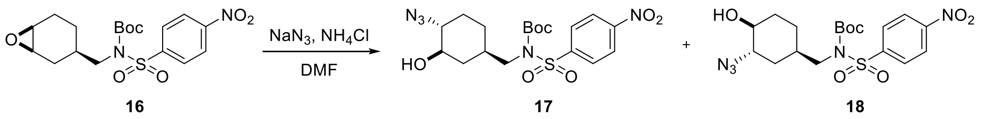

***tert*-Butyl (((1*R*,3*S*,4*S*)-3-amino-4-hydroxycyclohexyl)methyl)((4-nitrophenyl)sulfonyl)carbamate (17, 18)**

To a stirred solution of **16** (10.0 g, 24.2 mmol) and $NH_4Cl$ (12.9 g, 242 mmol) in DMF (100 mL) was added $NaN_3$ (7.8 g, 121.3 mmol). After being stirred at 80 ºC for 2 h, the reaction mixture was quenched with $H_2O$, extracted with EtOAc, and washed with brine. The combined organic extracts were dried over $Na_2SO_4$ and concentrated *in vacuo*. The crude mixture was purified by silica gel column chromatography (hexanes/EtOAc = 5/1) to afford **17, 18** (6.8 g). **17**: $[\alpha]_D^{21}$ - 0.046 (*c* = 0.2, MeOH); IR (thin film) $\nu_{max}$ = 2934, 2099, 1729, 1588, 1534, 1352, 1285, 1154, 1087, 739 cm$^{-1}$; $^1$H NMR (400 MHz, $CDCl_3$) δ 8.40 – 8.35 (m, 2H), 8.11 – 8.07 (m, 2H), 3.83 (qd, *J* = 14.5, 7.6 Hz, 2H), 3.73 – 3.60 (m, 2H), 2.29 (s, 1H), 1.97 – 1.82 (m, 2H), 1.68 – 1.54 (m, 11H), 1.35 (s, 10H); HRMS (ESI$^+$) *m/z* calcd for $C_{18}H_{27}N_3O_7S$ [M+H] 429.4880, found: 429.4886. **18**: $[\alpha]_D^{21}$ + 0.144 (*c* = 0.5, MeOH); IR (thin film) $\nu_{max}$ = 2934, 2099, 1729, 1588, 1534, 1352, 1285, 1154, 1087, 739 cm$^{-1}$; $^1$H NMR (400 MHz, $CDCl_3$) δ 8.36 (dd, *J* = 8.9, 2.2 Hz, 1H), 8.09 (dd, *J* = 8.8, 2.1 Hz, 1H), 7.87 (dd, *J* = 8.9, 2.4 Hz, 1H), 7.14 – 7.08 (m, 1H), 3.80 (dtd, *J* = 13.0, 8.6, 5.2 Hz, 3H), 3.48 (dq, *J* = 10.8, 7.0 Hz, 1H), 2.34 (s, 1H), 1.96 (dt, *J* = 7.9, 4.0 Hz, 2H), 1.83 – 1.67 (m, 3H), 1.67 – 1.47 (m, 4H), 1.35 (d, *J* = 2.3 Hz, 9H).

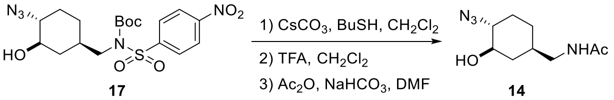


**_N_-(((1_R_,3_R_,4_R_)-4-Azido-3-hydroxycyclohexyl)methyl)acetamide (14)**

To a stirred solution of **17** (0.80 g, 1.8 mmol) and $CsCO_3$ (2.8 g, 8.8 mmol) was treated with BuSH (0.8 g, 8.8 mmol) in $CH_2Cl_2$ (3.5 mL). After being stirred at r.t. for 15 h, the reaction mixture was filtered and evaporated *in vacuo*. The solution of the crude product in a 7:3 mixture of $CH_2Cl_2$ and TFA (9 mL) was stirred at r.t. for 1h, all volatiles were evaporated *in vacuo*. To a solution of the residue and $NaHCO_3$ (0.5 g, 5.9 mmol) in DMF (2.4 mL) was added $Ac_2O$ (0.6 mL, 5.9 mmol). After being stirred at r.t. for 14 h, the reaction solution was filtered and evaporated *in vacuo*. The crude mixture was purified by silica gel column chromatography ($CHCl_3$/MeOH = 98/2) to afford **14** (0.3 g, 75% for 3 steps): $[\alpha]^{21}_{D}$ + 0.14 (*c* = 0.5, MeOH); IR (thin film) $\nu_{max}$ = 2934, 2099, 1729, 1588, 1534, 1352, 1285, 1154, 1087, 739 $cm^{-1}$; $^1H$ NMR (400 MHz, $CDCl_3$) δ 5.53 (s, 1H), 3.61 (td, *J* = 9.0, 5.1 Hz, 2H), 3.22 (dh, *J* = 26.5, 6.7 Hz, 2H), 2.00 (s, 3H), 1.92 – 1.76 (m, 5H), 1.60 (dt, *J* = 13.3, 5.7 Hz, 2H), 1.51 (q, *J* = 5.7 Hz, 2H); HRMS ($ESI^+$) *m/z* calcd for $C_9H_{16}N_4O_2$ [M+H] 212.1273, found: 212.1273.

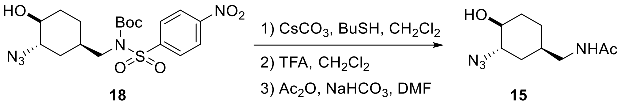


**_N_-(((1_R_,3R,4_R_)-3-Azido-4-hydroxycyclohexyl)methyl)acetamide (15)**

To a stirred solution of **18** (0.80 g, 1.8 mmol) and $CsCO_3$ (2.8 g, 8.8 mmol) was treated with BuSH (0.8 g, 8.8 mmol) in $CH_2Cl_2$ (3.5 mL). After being stirred at r.t. for 16 h, the reaction mixture was filtered and evaporated *in vacuo*. The solution of the crude product in a 7:3 mixture of $CH_2Cl_2$ and TFA (9 mL) was stirred at r.t. for 1h, all volatiles were evaporated *in vacuo*. To a solution of the residue and $NaHCO_3$ (0.5 g, 5.9 mmol) in DMF (2.4 mL) was added $Ac_2O$ (0.6 mL, 5.9 mmol). After being stirred at r.t. for 15 h, the reaction solution was filtered and evaporated *in vacuo*. The crude mixture was purified by silica gel column chromatography ($CHCl_3$/MeOH = 98/2) to afford **15** (0.3 g, 75% for 3 steps): $[\alpha]^{21}_{D}$ + 0.144 (*c* = 0.5, MeOH); IR (thin film) $\nu_{max}$ = 2934, 2099, 1729, 1588, 1534, 1352, 1285, 1154, 1087, 739.02 $cm^{-1}$; $^1H$ NMR (400 MHz, $CDCl_3$) δ 5.97 (s, 1H), 3.79 (td, *J* = 6.7, 3.7 Hz, 1H), 3.48 (td, *J* = 6.8, 3.9 Hz, 1H), 3.17 (dq, *J* = 17.1, 6.8 Hz, 2H), 2.01 (s, 3H), 1.93 (ddd, *J* = 12.5, 7.5, 3.1 Hz, 1H), 1.61 (dddd, *J* = 33.7, 17.4, 7.8, 3.9 Hz, 3H), 1.48 – 1.36 (m, 1H); HRMS ($ESI^+$) *m/z* calcd for $C_9H_{16}N_4O_2$ [M+H] 212.1273, found: 212.1283.

### Water solubility of TM-Cy-TBPA (4) and TM-TBPA (3)

**Figure S6.** Determination of aqueous solubility by HPLC analysis.

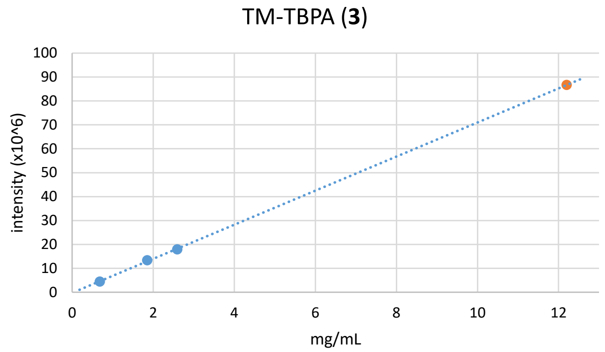


| mg/mL | 0.68 | 1.85 | 2.59 | sat. |
|---|---|---|---|---|
| intensity | 4.45628 | 13.37552 | 17.95734 | 86.71324 |

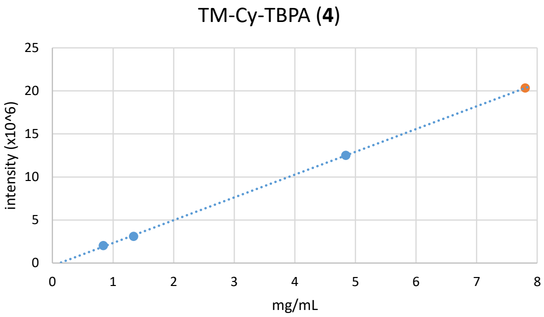


| mg/mL | 0.84 | 1.34 | 4.84 | sat. |
|---|---|---|---|---|
| intensity | 2.027465 | 3.099609 | 12.520636 | 20.239246 |

**Chemical stability under acidic conditions**

A solution of **3**, **4** or TM-V in MeOH was treated with 1M or 2M HCl solution in MeOH/$H_2O$ (1/1) at r.t. for 0.5, 1, 2, 4, 6 h. The area of the residual peak for the compounds was quantified by reverse-phase HPLC.

HPLC conditions for TM-TBPA (**3**) and TM-Cy-TBPA (**4**)
solvent: 60:40 MeOH/$H_2O$
UV: 254 nm
flow rate: 0.5 ml/ min
column: Kinetex 1.7 μm XB-C18, 100 Å, 150 x 2.10 mm

HPLC conditionss for TM-V (**1**)
solvent: 90:10 MeOH/$H_2O$
UV: 254 nm
flow rate: 0.5 ml/ min
column: Kinetex 1.7 μm XB-C18, 100 Å, 150 x 2.10 mm

**Purity of TM-Cy-TBPA (4) synthesized in Scheme 2.**

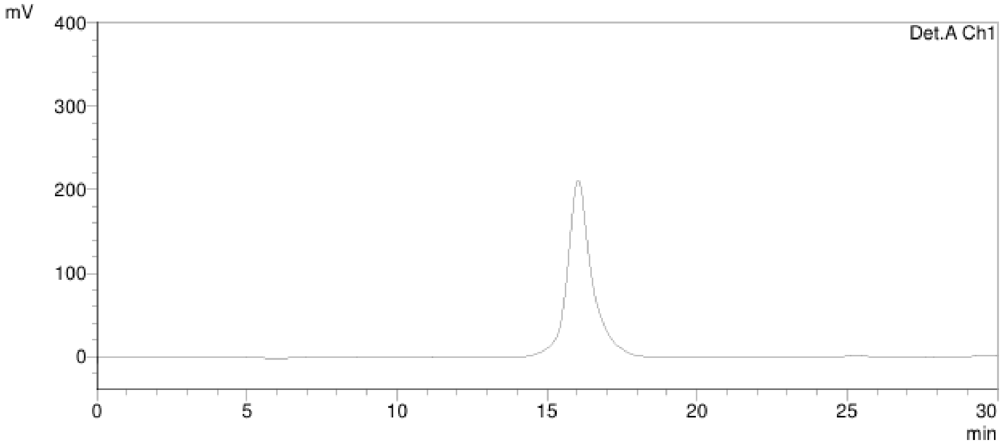


HPLC conditions
solvent: 60:40 MeOH/$H_2O$
UV: 254 nm
flow rate: 0.5 ml/ min
column: Kinetex 1.7 μm XB-C18, 100 Å, 150 x 2.10 mm

**Expression and purification of MraY**
The gene *mraY* of *Hydrogenivirga spp.* 128-5-R1-1 was cloned with an *N*-terminal $His_6$ tag into a pET22b vector. The plasmid was transformed and expressed in *E. coli* NiCo21(DE3) pLEMO competent cells. The proteins were purified using a nickel, cation exchange, and size exclusion chromatography. The final storage buffer was 20 mM HEPES pH 7.5, 100 mM NaCl, 10% glycerol, 5 mM βME, 0.15% n-decyl-β-D-maltopyranoside (DM).

**Preparation of membrane fraction P-60 containing MurX**
*M. smegmatis* (ATCC607) cells were harvested by centrifugation (4700 rpm) at 4 ºC followed by washing with 0.9% saline solution (thrice). The washed cell pellets were washed with homogenization buffer (containing 50 mM

$K_2HPO_4$, 5 mM $MgCl_2$, 5 mM 1,4-dithio-DL-threitol, and 10% glycerol, pH = 7.2) (thrice), and approximately 5 g of pellet (wet weight) was collected. The washed cell pellets were suspended in homogenization buffer and disrupted by probe sonication on ice (5 s on and 2 s off for 1 min, then cool down for 1.5 min, 5 cycles, cool down for 15 min, and then, 5 s on and 2 s off for 1 min, cool down for 1.5 min, 5 cycles). The resulting suspension was centrifuged at 4,700 x g for 15 min at 4 ºC to remove unbroken cells. The lysate was centrifuged at 25,000 x g for 20 min at 4 ºC. The supernatant was subjected to ultracentrifugation at 60,000 x g for 1 h at 4 ºC. The supernatant was discarded, and the membrane fraction containing MraY enzyme (P-60) was suspended in the Tris-HCl buffer (pH = 7.5). Total protein concentrations were approximately 8 to 10 mg/mL. Aliquots were stored in Eppendorf tubes at -80 ºC.

**MraY/MurX assays**

MraY/MurX assay substrates, Park's nucleotide-$N^{\varepsilon}$-$C_6$-dansylthiourea and neryl phosphate, were chemically synthesized according to the reported procedures.[3e,3f] Park's nucleotide-$N^{\varepsilon}$-$C_6$-dansylthiourea[23] (2 mM stock solution, 1.88 µL), $MgCl_2$ (0.5 M, 5 µL), KCl (2 M, 5 µL), Triton X-100 (0.5%, 5.63 µL), Tris buffer (pH 8.0, 50 mM), neryl phosphate (0.1 M, 2.25 µL), and inhibitor molecule (0 - 50 g/mL in Tris buffer) were placed in a 1.5 mL Eppendorf tube. To a stirred reaction mixture, MraY/MurX (10 µL) was added (total volume of reaction mixture: 50 µL adjusted with Tris buffer). The reaction mixture was incubated for 2 h at room temperature (26 ºC) and quenched with $CHCl_3$ (100 µL). Two phases were mixed via vortex and centrifuged at 25,000 xg for 10 min. The upper aqueous phase was assayed via reverse-phase HPLC. The water phase (10 µL) was injected into HPLC (solvent: $CH_3CN$/0.05 M aq. $NH_4HCO_3$ = 25:75; UV: 350 nm; flow rate: 0.5 mL/min; column: Kinetex 5µm C8, 100 A, 150 x 4.60 mm), and the area of the peak for lipid I-neryl derivative was quantified to obtain the $IC_{50}$ value. The $IC_{50}$ values were calculated from plots of the percentage product inhibition versus the inhibitor.

**Bacterial WecA Inhibitory activity**

WecA assay substrate, UDP-Glucosamine-C6-FITC was chemically synthesized according to the reported procedures. UDP-Glucosamine-C6-FITC (2 mM stock solution, 0.56 µL), $MgCl_2$ (0.5 M, 4.0 µL), β-mercaptoethanol (50 mM, 5.0 µL), CHAPS (20%, 2.5 µL), Tris buffer (pH 8.0, 50 mM), undecaprenyl phosphate (2 mM, 1.7 µL), and inhibitor molecule (10 mM stock solution in DMSO) were place in a 1.5 mL Eppendorf tube. To a stirred reaction mixture, P-60 (10.0 µL) was added (total volume of reaction mixture: 50 µL adjust with Tris buffer). The reaction mixture was incubated for 3 h and quenched with n-butanol (150 µL). Two phases were mixed via vortex and centrifuged at 25,000 xg for 5 min. The upper organic phase was assayed via reverse-phase HPLC. The organic phase (30 µL) was injected into HPLC (solvent: gradient elution of 85:15 to 95:5 MeOH/0.05 M aq. $NH_4HCO_3$; UV: 485 nm; flow rate: 0.5 ml/ min; column: Kinetex 5 µm C8, 100 Å, 150 x 4.60 mm), and the area of the peak for C55-P-P-glucosamine-$C_6$-FITC was quantified to obtain the $IC_{50}$ value. The $IC_{50}$ values were calculated from plots of the percentage product inhibition versus the inhibitor concentration.

**Bacterial strains and growth of bacteria**

*Mycobacterium tuberculosis* ($H_{37}$Rv), *Candida albicans* (18M), *Cryptococcus neoformans* (NIH9hi90), *Aspergillus fumigatus* (ASFU-2263), *Enterococcus faecium* (UAA714 NR-32065), *Staphylococcus aureus* (71080 NR-46418), *Klebsiella pneumoniae* (BWH22 NR-41899), *Candida glabrata* (DSY565 NR-51686) and *Cryptococcus gattii* (C17 NR-50431) were obtained through BEI Resources, National Institute of Allergy and Infectious Diseases (NIAID), National Institutes of Health (NIH). *Mycobacterium smegmatis* (ATCC 607), *Bacillus subtilis* (ATCC6051), *Staphylococcus aureus* (ATCC 6538), *Enterococcus faecium* (ATCC 349), *Pseudomonas aeruginosa* (ATCC 27853) and *E. coli* (ATCC 35218) were obtained from American Type Culture Collection (ATCC). A single colony of *Mycobacterium* was obtained on Difco Middlebrook 7H10 nutrient agar enriched with 10% oleic acid, albumin, dextrose, and catalase (OADC) for *M. tuberculosis* by incubating for 15 days, and with albumin, dextrose, and catalase (ADC) for *M. smegmatis* by incubating for 48 h at 37ºC in a static incubator. Seed cultures and larger cultures were obtained using Middlebrook 7H9 broth enriched with OADC (for *M. tuberculosis*) by incubating for 15 days and ADC (for *M. smegmatis*) by incubating for 48h at 37°C in a shaking incubator (200rpm) until log phase to be an optical density (OD) of 0.2-0.5. The OD was monitored at 600 and 570 nm using a 96-well microplate reader. The others were obtained on the recommended agar media.

**Antibacterial and Antifungal Activities of TM-Cy-TBPA (4) in Comparison with TM-V (1)**
Minimum inhibitory concentrations were determined by broth dilution microplate alamar blue assay or by OD measurement. All compounds were stored in DMSO (1 mg/100 µL concentration). This concentration was used as the stock solution for all MIC studies. Each compound from stock solution was placed in the first well of a sterile 96 well plate and a serial dilution was conducted with the culturing broth (total volume of 10 µL). The bacterial suspension at log phase (190 µL) was added to each well (total volume of 200 µL), and was incubated for 24 h at 37 °C. 20 µL of resazurin (0.02%) was added to each well and incubated for 4 h for *Mycobacterium spp.* (National Committee for Clinical Laboratory Standards (NCCLS) method (pink = growth, blue = no visible growth)). The OD measurements were performed for all experiments prior to colorimetric. The MIC values were determined according to the colorimetric assays using resazurin. The absorbance of each well was also measured at 570 and 600 nm via UV-Vis.

**Table S4.** MIC of TM-Cy-TBPA (**4**) against bacteria and fungus susceptable to TM-V (**1**).[a]

| Compound | $MIC_{90}$ (µg/mL)* | | |
|---|---|---|---|
| | *B. subtilis* ATCC6051[b] | *B. cereus* NRRL B-568[c] | *M. smegmatis* ATCC607[d,e]] |
| **TM-Cy-TBPA** (**4**) | >35 | >35 | >35 |
| **TM-V** (**1**) | 0.3 | 0.2 | 12.5 |

| Compound | $MIC_{50}$ (µg/mL) | | | | | |
|---|---|---|---|---|---|---|
| | *C. albicans* 18M[f] | *C. glabrata* DSY565 (NR-51686)[g] | *C. neoformans* NIH9hi90[h] | *C. gattii* C17[i] | *A. fumigatus* ASFU-2263[j] | *S. cerevisiae* BY4742[k] |
| **TM-Cy-TBPA** (**4**) | >35 | >35 | >35 | >35 | >35 | >35 |
| **TM-V** (**1**) | **<0.1** | **0.9** | **0.2** | **<0.1** | **0.9** | **1.2** |

[a] Microplate Alamar (resazurin) blue assays were applied. [b] *Bacillus subtilis* (ATCC6051). [c] *Bacillus cereus* NRRL B-568 (10876D05™). [d] *Mycobacterium smegmatis* (ATCC607). [e] A nutrient-deficient condition was applied. [f] *Candida albicans* 18M. [g] *Candida glabrata* DSY565 (NR-51686). [h] *Cryptococcus neoformans* NIH9hi90. [i] *Cryptococcus gattii* C17 [j] *Aspergillus fumigatus* ASFU-2263. [k] *Saccharomyces cerevisiae* BY4742A All bacterial and fungal strains were acquired from ATCC and EBI, respectively. *$p < 0.01$ ($n = 3$).

**Membrane disruption assays**

**Figure S7.** Eosin staining.[a]

**RBCs treated with TM-V**

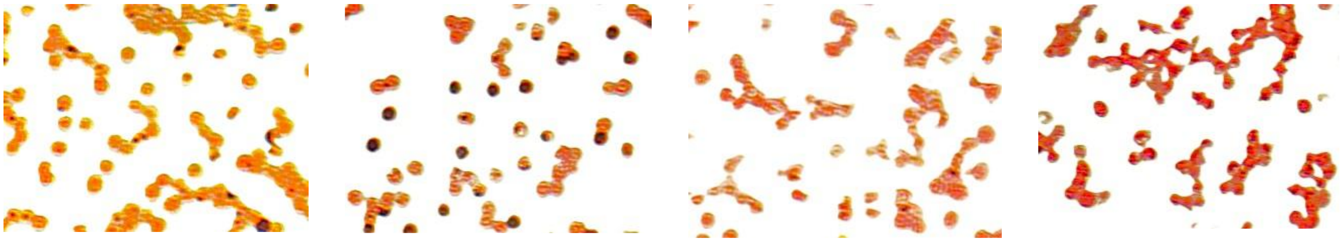

Control 1.0 μM 2.5 μM 10.0 μM

**RBCs treated with TM-Cy-TBPA**

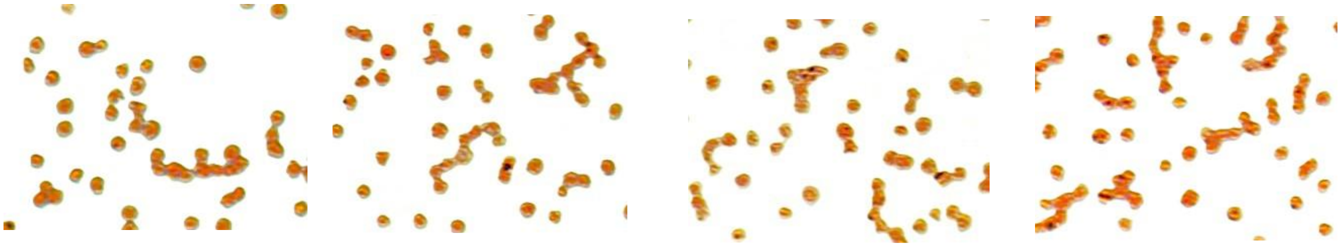

Control 1.0 μM 2.5 μM 10.0 μM

[a] Images were acquired at ×100 magnification (an EVOS microscope)

**Figure S8.** LDH release and membrane permeability assays.

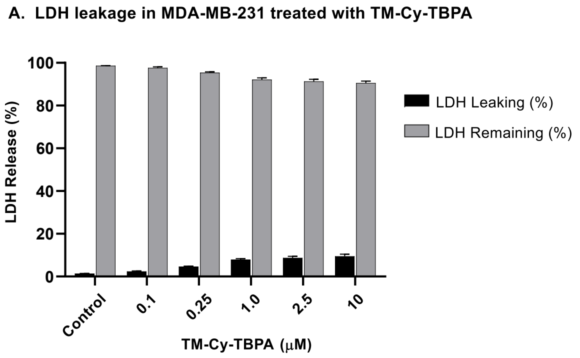


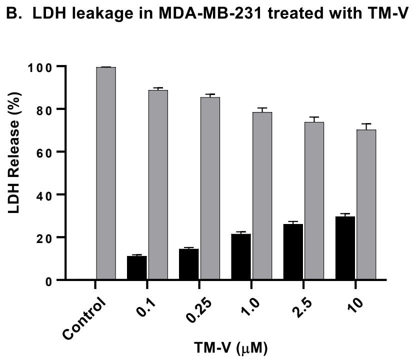


| | (%) | error (%) | (%) | error (%) |
|---|---|---|---|---|
| control | 1.85 | ±0.1 | 98.15 | ±0.2 |
| 0.1 μM | 2.45 | ±0.2 | 97.55 | ±0.4 |
| 0.25 μM | 2.74 | ±0.2 | 97.26 | ±0.3 |
| 1.0 μM | 4.45 | ±0.3 | 95.55 | ±0.5 |
| 2.5 μM | 5.25 | ±0.3 | 94.75 | ±0.6 |
| 10 μM | 5.43 | ±0.3 | 94.57 | ±0.6 |

| | (%) | error (%) | (%) | error (%) |
|---|---|---|---|---|
| control | 0.43 | ±0.02 | 99.57 | ±0.04 |
| 0.1 μM | 11.24 | ±0.55 | 88.76 | ±1.1 |
| 0.25 μM | 14.53 | ±0.7 | 85.47 | ±1.4 |
| 1.0 μM | 21.54 | ±1 | 78.46 | ±2 |
| 2.5 μM | 26.16 | ±1.2 | 73.84 | ±2.4 |
| 10 μM | 29.70 | ±1.35 | 70.30 | ±2.7 |

A. LDH leakage in MCF-7 treated with TM-Cy-TBPA

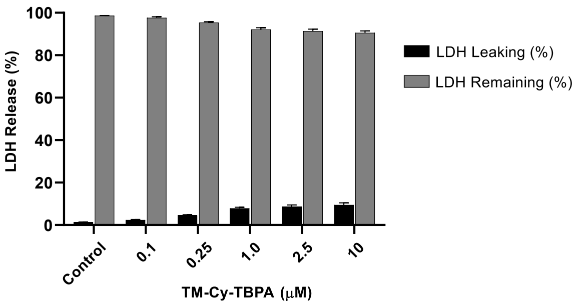


| | (%) | error (%) | (%) | error (%) |
|---|---|---|---|---|
| control | 1.37 | ±0.1 | 98.63 | ±0.1 |
| 0.1 μM | 2.36 | ±0.3 | 97.64 | ±0.5 |
| 0.25 μM | 4.61 | ±0.3 | 95.39 | ±0.4 |
| 1.0 μM | 7.84 | ±0.5 | 92.16 | ±0.8 |
| 2.5 μM | 8.75 | ±0.7 | 91.25 | ±1 |
| 10 μM | 9.48 | ±1 | 90.52 | ±0.9 |

B. LDH leakage in MCF-7 treated with TM-V

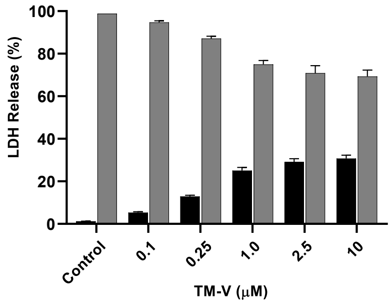


| | (%) | error (%) | (%) | error (%) |
|---|---|---|---|---|
| control | 1.25 | ±0.04 | 98.75 | ±0.05 |
| 0.1 μM | 5.32 | ±0.4 | 94.68 | ±0.8 |
| 0.25 μM | 12.85 | ±0.6 | 87.15 | ±1.1 |
| 1.0 μM | 25.01 | ±1.5 | 74.99 | ±1.8 |
| 2.5 μM | 29.15 | ±1.5 | 70.85 | ±3.5 |
| 10 μM | 30.70 | ±1.6 | 69.30 | ±3 |

**Figure S9.** Western blot analysis of cell-cycle regulatory proteins in MDA-MB-231 cells following treatment with TM-Cy-TBPA (**4**). β-Actin was used as the protein loading control.

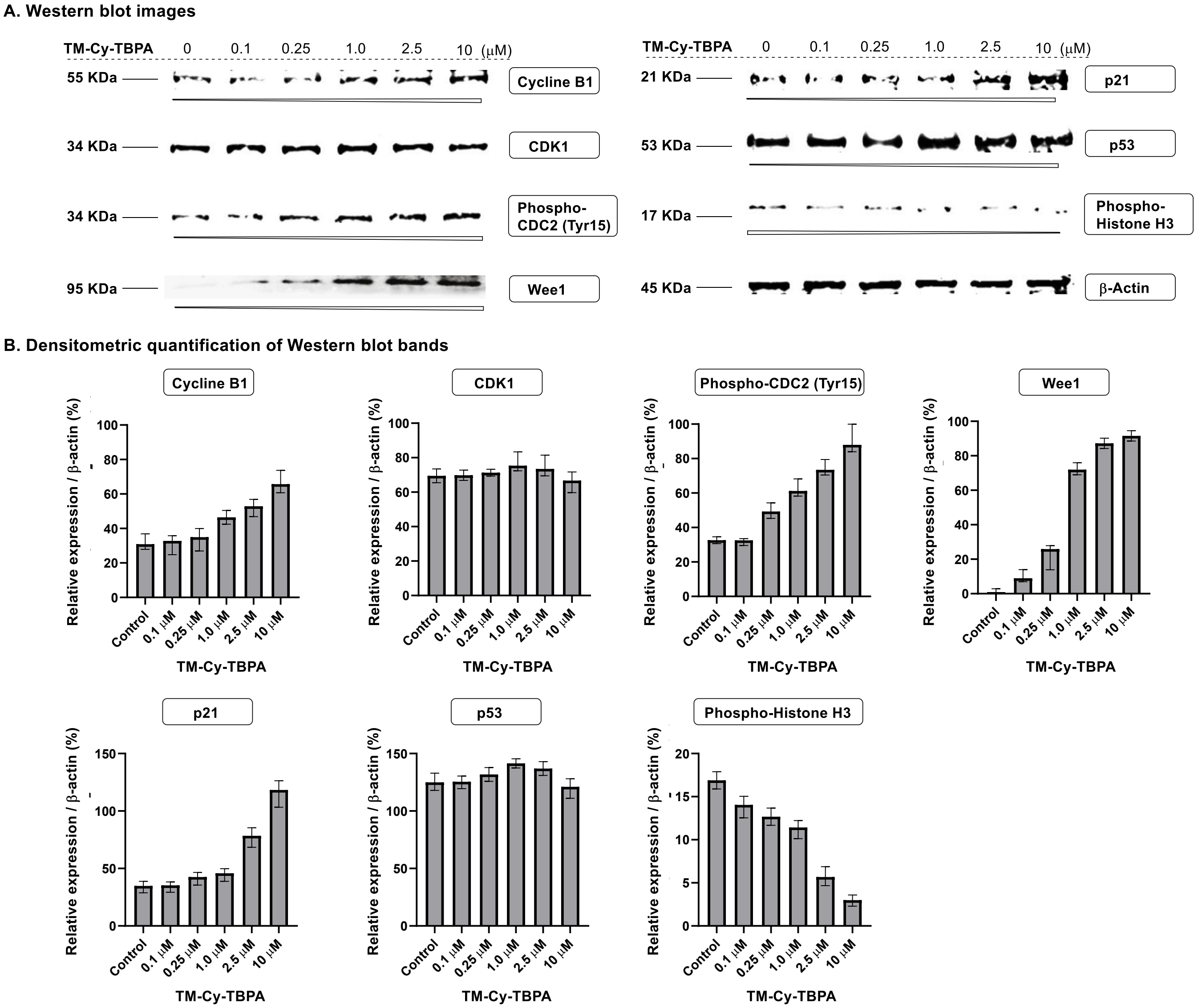


Wee1 (D10D2) Rabbit Monoclonal Antibody (Cell Signaling Technology, Cat. #13084), Cyclin B1 (D5C10) Rabbit Monoclonal Antibody (Cell Signaling Technology, Cat. #12231), p53 (7F5) Rabbit Monoclonal Antibody (Cell Signaling Technology, Cat. #2527), Anti-CDK1 antibody [1/Cdk1/Cdc2] (abcam, Cat. #ab280964), p21 Waf1/Cip1 (12D1) Rabbit Monoclonal Antibody (Cell Signaling Technology, Cat. #2947), Phospho-Histone H3 (Ser10) (D2C8) Rabbit Monoclonal Antibody (Cell Signaling Technology, Cat. #3377), Phospho-cdc2 (Tyr15) (10A11) Rabbit Monoclonal Antibody (Cell Signaling Technology, Cat. #4539), and beta-Actin (E4D9Z) Mouse Monoclonal Antibody (Cell Signaling Technology, Cat. #58169) were used as primary antibody. IRDye 680RD Goat anti-Rabbit IgG Secondary Antibody (LI-COR Biotech, Cat. #926-68071) and IRDye 800CW Goat anti-Mouse IgG Secondary Antibody (LI-COR Biotech, Cat. #926-32210) were used as secondary antibody. LI-COR Biotech ODYSSEY DLX was used to scan the probe signal. The data was quantified with ImageJ1.54d (NIH) processing software. ($n = 3$, $p<0.05$)

**Figure S10.** Western blot analysis of apoptosis regulatory proteins in MDA-MB-231 cells following treatment with TM-Cy-TBPA (**4**). GAPDH was used as the protein loading control.

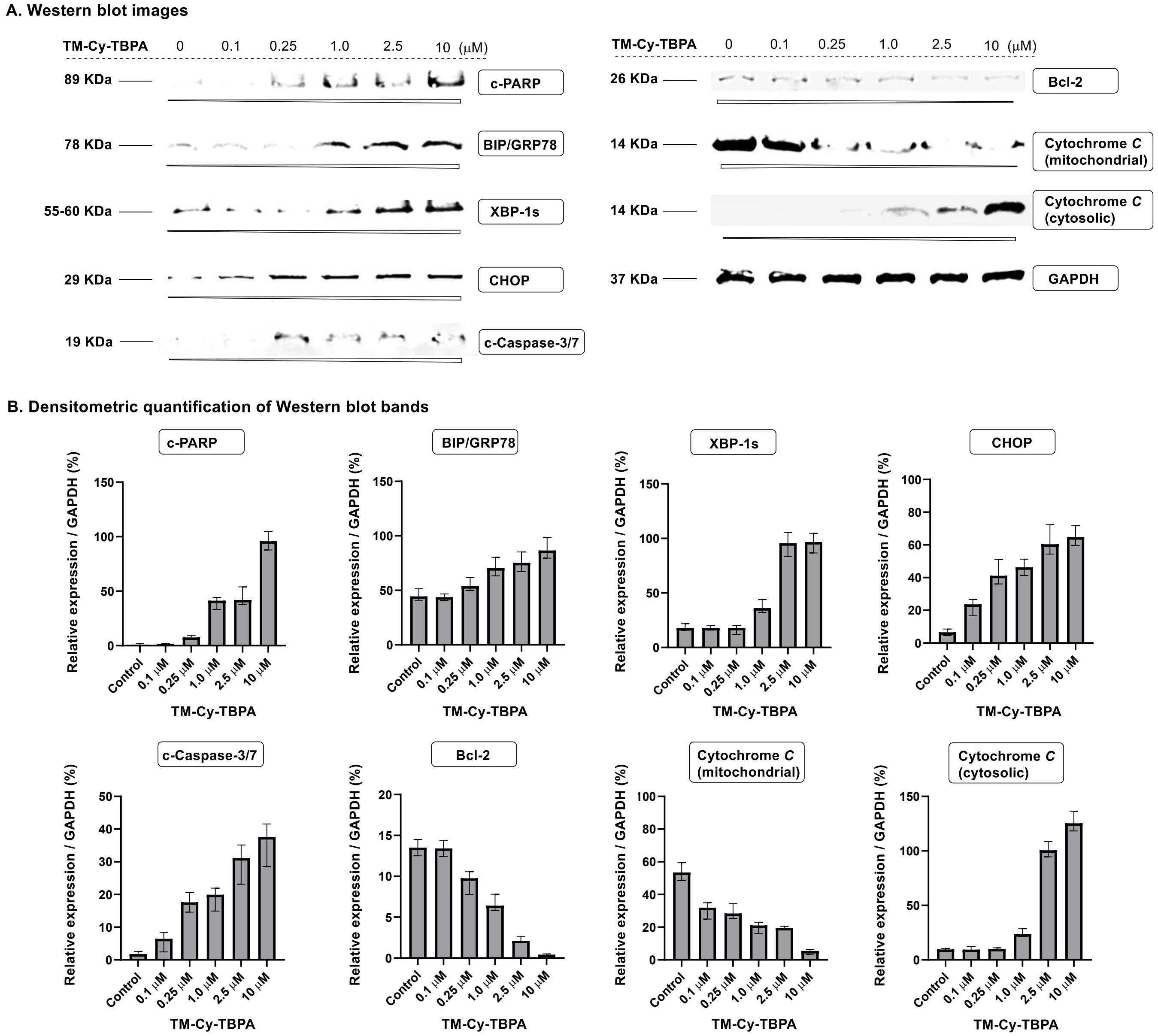


*To improve detection of treatment effects at lower concentrations of compound **4**, the amount of total protein loaded for Western blot analysis was increased four-fold. Cleaved PARP (Asp214) (D64E10) Rabbit Monoclonal Antibody (Cell Signaling Technology, Cat. #5625), CHOP (L63F7) Mouse Monoclonal Antibody (Cell Signaling Technoloty, Cat. #2895), BiP (C50B12) Rabbit Monoclonal Antibody (Cell Signaling Technoloty, Cat. #3177), XBP-1s (E9V3E) Rabbit Monoclonal Antibody (Cell Signaling Technoloty, Cat. #40435), Bcl-2 Recombinant Rabbit Monoclonal Antibody (JE10-17) (Invitrogen, Cat. #MA5-36172), Cleaved Caspase-3 (Asp175) Antibody (Cell Signaling Technoloty, Cat. #9661), Anti-Cytochrome c mouse mAb (Cytochrome c Releasing Apoptosis Assay Kit, BioVision, Cat. #K257-100), and GAPDH (D4C6R) Mouse Monoclonal Antibody (Cell Signaling Technoloty, Cat. #97166) were used as primary antibody. IRDye 680RD Goat anti-Rabbit IgG Secondary Antibody (LI-COR Biotech, Cat. #926-68071) and IRDye 800CW Goat anti-Mouse IgG Secondary Antibody (LI-COR Biotech, Cat. #926-32210) were used as secondary antibody. LI-COR Biotech ODYSSEY DLX was used to scan the probe signal. The data was quantified with ImageJ1.54d (NIH) processing software. ($n = 3$, $p<0.05$)

**Figure S11.** The contrast of the Bcl-2 Western blot image for SK-BR-3 cells was adjusted to improve band visibility (**Figure 11**).[a]

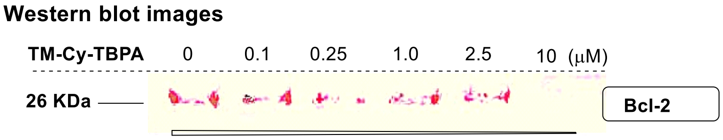


[a] Highlighted by contrast enhancement of the image in Figure 11.

**Figure S12.** The contrast of the Bcl-2 Western blot image for MDA-MB-231 cells was adjusted to improve band visibility (**Figure S11**).[a]

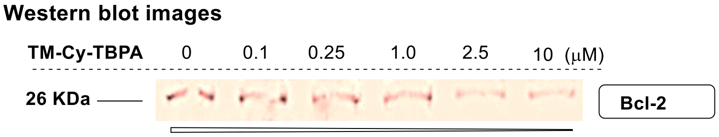


[a] Highlighted by contrast enhancement of the image in Figure S11.

**Table S5.** Caco-2 permeability determination.

| Compound | $P_{app}$ ($10^{-6}$ cm/s) | | Efflux ratio | Recovery (%) | |
|---|---|---|---|---|---|
| | A→B | B→A | | A→B | B→A |
| **TM-TBPA (3)** | 0.8 | 0.5 | **0.6** | 90.2 | 109.3 |
| **TM-Cy-TBPA (4)** | 0.13 | 0.14 | **1.1** | 100.0 | 121.0 |
| Nadolol | 0.3 | 0.7 | 2.3 | 102.9 | 103.6 |
| Propranolol | 35.4 | 27.1 | 0.8 | 99.0 | 101.5 |
| Digoxin | 0.4 | 11.9 | 29.4 | 103.5 | 102.1 |

**Permeability classification:**

| Permeability | $P_{app}$ ($10^{-6}$ cm/s) |
|---|---|
| High | > 10 |
| Moderate | 1 ~ 10 |
| Low | ≤ 1 |

Propranolol has a relatively variable but generally moderate oral bioavailability (range 15–35%). Nadolol has a moderate oral bioavailability of approximately 30–40%. Digoxin has a comparatively high oral bioavailability (range 70–80%).

**Table S6.** Plasma protein binding determination.

| Compound | Mouse | | Dog | | Human | |
|---|---|---|---|---|---|---|
| | Bound (%) | Recovery (%) | Bound (%) | Recovery (%) | Bound (%) | Recovery (%) |
| **TM-TBPA (3)** | 72.3 | 91.4 | 61.7 | 93.3 | 74.3 | 100.6 |
| **TM-Cy-TBPA (4)** | 87.3 | 96.8 | 65.5 | 95.5 | 68.5 | 98.9 |
| Warfarin | 99.1 | 96.6 | 95.0 | 91.1 | 98.9 | 91.5 |

Warfarin exhibits both very high oral bioavailability and extensive plasma protein binding, Oral bioavailability (*F*): approximately 95–100%, Protein binding: approximately 99%.

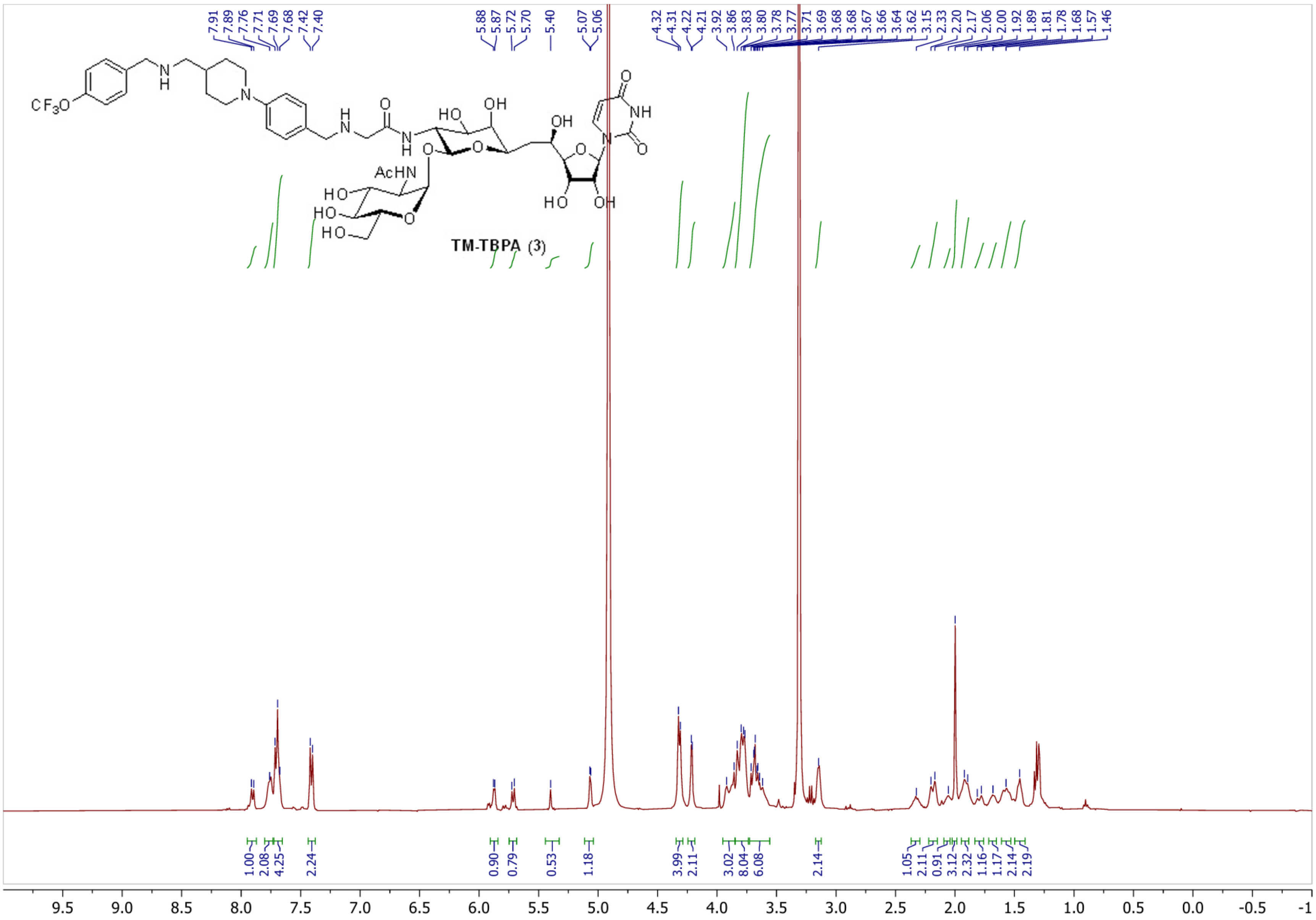

7.91 7.89 7.76 7.71 7.69 7.68 7.42 7.40
5.88 5.87 5.72 5.70 5.40 5.07 5.06
4.32 4.31 4.22 4.21 3.92 3.86 3.83 3.80 3.78 3.77 3.71 3.69 3.68 3.68 3.67 3.66 3.64 3.62 3.15 2.33 2.20 2.17 2.06 2.00 1.92 1.89 1.81 1.78 1.68 1.57 1.46
CF3O
N
H
N
N
H
O
N
H
HO
OH
OH
O
O
O
O
NH
N
O
AcHN
HO
HO
HO
O
HO
OH
TM-TBPA (3)
1.00 2.08 4.25 2.24 0.90 0.79 0.53 1.18 3.99 2.11 3.02 8.04 6.08 2.14 1.05 2.11 0.91 3.12 2.32 1.16 1.17 2.14 2.19
9.5 9.0 8.5 8.0 7.5 7.0 6.5 6.0 5.5 5.0 4.5 4.0 3.5 3.0 2.5 2.0 1.5 1.0 0.5 0.0 -0.5 -1

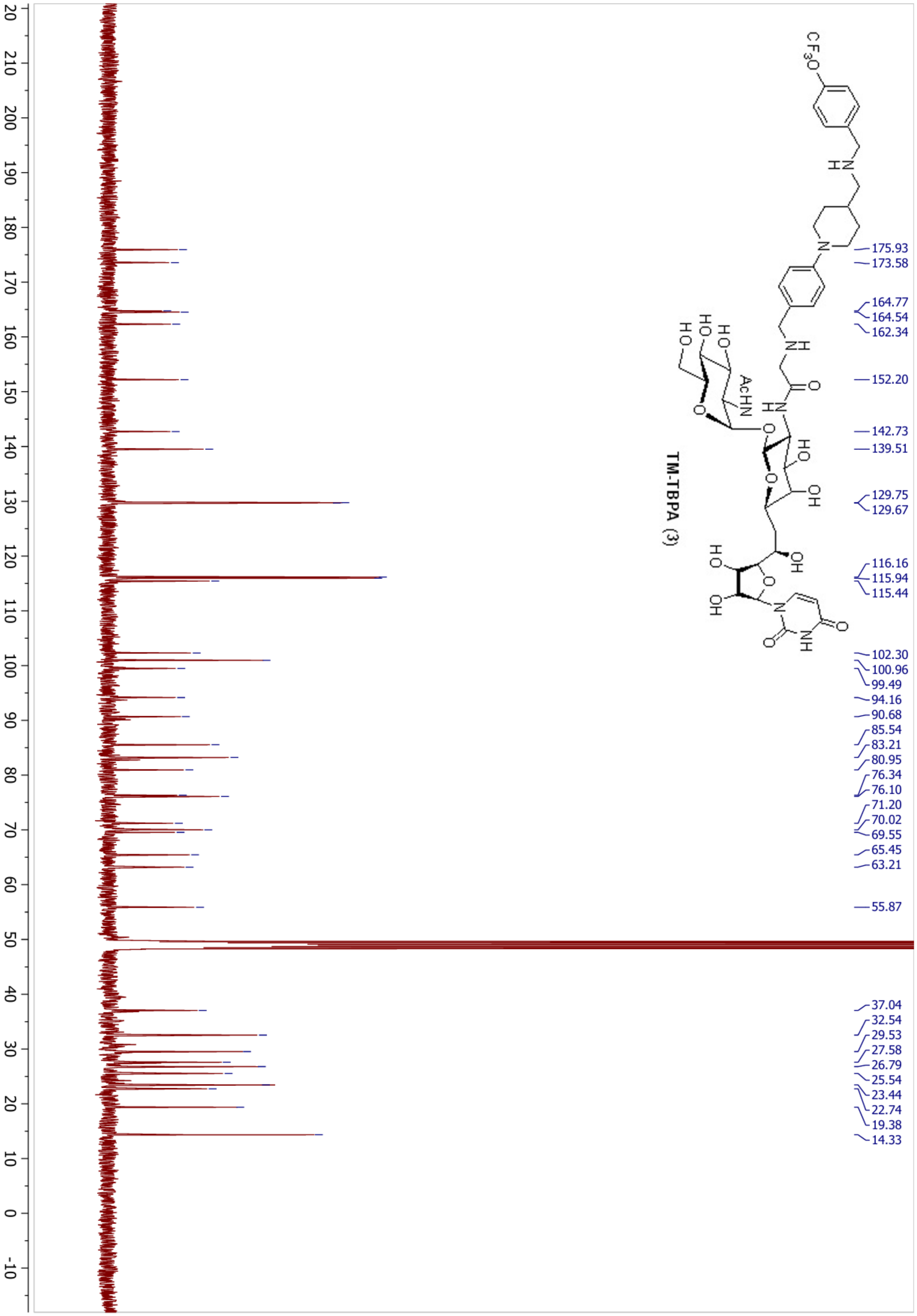

TM-TBPA (3)
175.93
173.58
164.77
164.54
162.34
152.20
142.73
139.51
129.75
129.67
116.16
115.94
115.44
102.30
100.96
99.49
94.16
90.68
85.54
83.21
80.95
76.34
76.10
71.20
70.02
69.55
65.45
63.21
55.87
37.04
32.54
29.53
27.58
26.79
25.54
23.44
22.74
19.38
14.33

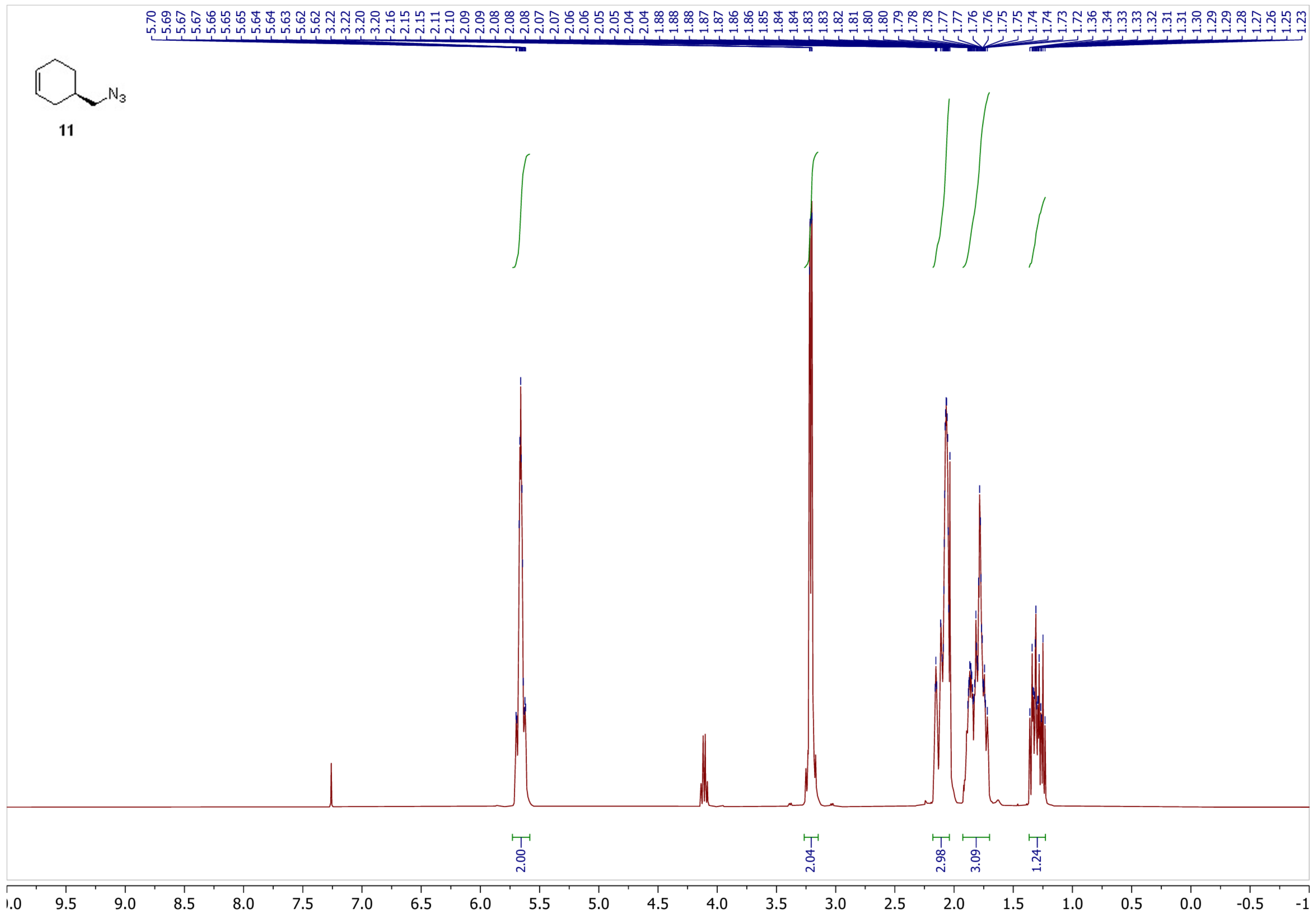

11
N3
5.70
5.69
5.67
5.67
5.66
5.65
5.65
5.64
5.64
5.63
5.62
5.62
3.22
3.22
3.20
3.20
2.16
2.15
2.15
2.11
2.10
2.09
2.09
2.08
2.08
2.08
2.07
2.07
2.06
2.06
2.05
2.05
2.04
2.04
1.88
1.88
1.88
1.87
1.87
1.86
1.86
1.85
1.84
1.84
1.83
1.83
1.82
1.81
1.80
1.80
1.79
1.78
1.78
1.77
1.77
1.76
1.76
1.75
1.75
1.74
1.74
1.73
1.72
1.36
1.34
1.33
1.33
1.32
1.31
1.31
1.30
1.29
1.29
1.28
1.27
1.26
1.25
1.23
2.00
2.04
2.98
3.09
1.24

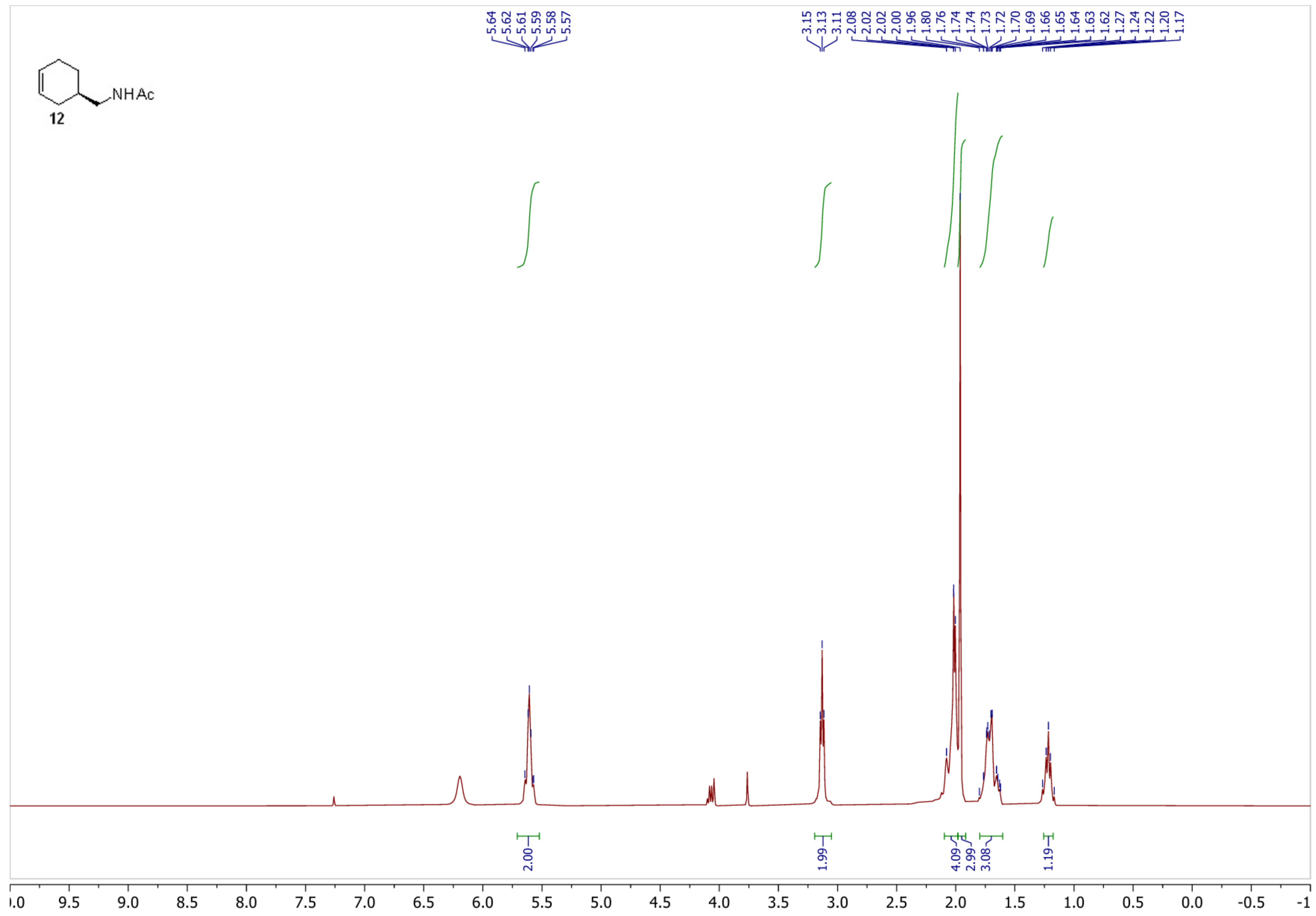
NHAc
12
5.64
5.62
5.61
5.59
5.58
5.57
3.15
3.13
3.11
2.08
2.02
2.02
2.00
1.96
1.80
1.76
1.74
1.74
1.73
1.72
1.70
1.69
1.66
1.65
1.64
1.63
1.62
1.27
1.24
1.22
1.20
1.17
2.00
1.99
4.09
2.99
3.08
1.19
9.5
9.0
8.5
8.0
7.5
7.0
6.5
6.0
5.5
5.0
4.5
4.0
3.5
3.0
2.5
2.0
1.5
1.0
0.5
0.0
-0.5

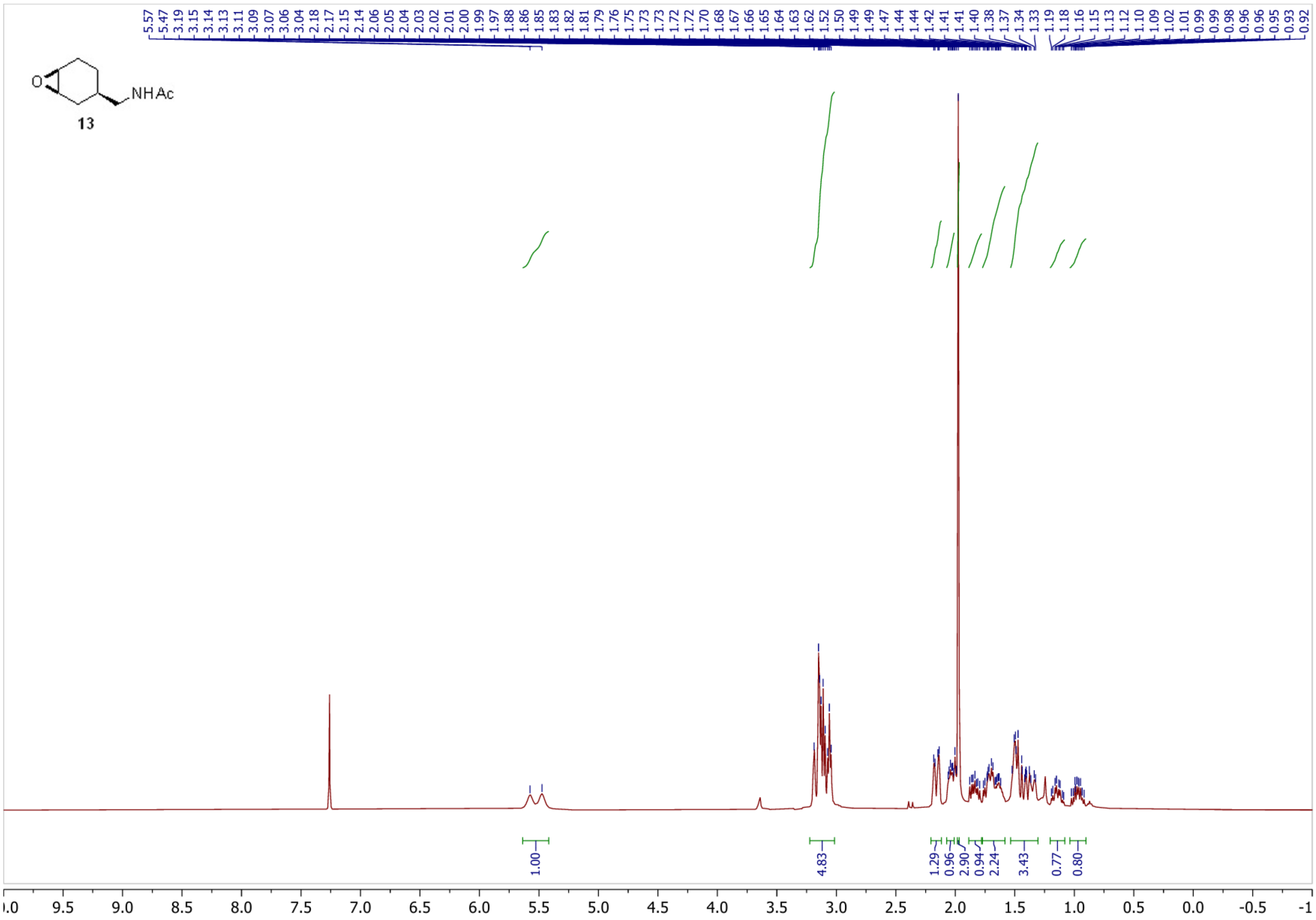
NHAc
13
5.57 5.47 3.19 3.15 3.14 3.13 3.11 3.09 3.07 3.06 3.04 2.18 2.17 2.15 2.14 2.06 2.05 2.04 2.03 2.02 2.01 2.00 1.99 1.97 1.88 1.86 1.85 1.83 1.82 1.81 1.79 1.76 1.75 1.73 1.73 1.72 1.72 1.70 1.68 1.67 1.66 1.65 1.64 1.63 1.62 1.52 1.50 1.49 1.49 1.47 1.44 1.44 1.42 1.41 1.41 1.40 1.38 1.37 1.34 1.33 1.19 1.18 1.16 1.15 1.13 1.12 1.10 1.09 1.02 1.01 0.99 0.99 0.98 0.96 0.96 0.95 0.93 0.92
1.00
4.83
1.29
0.96
2.90
0.94
2.24
3.43
0.77
0.80
9.5 9.0 8.5 8.0 7.5 7.0 6.5 6.0 5.5 5.0 4.5 4.0 3.5 3.0 2.5 2.0 1.5 1.0 0.5 0.0 -0.5 -1

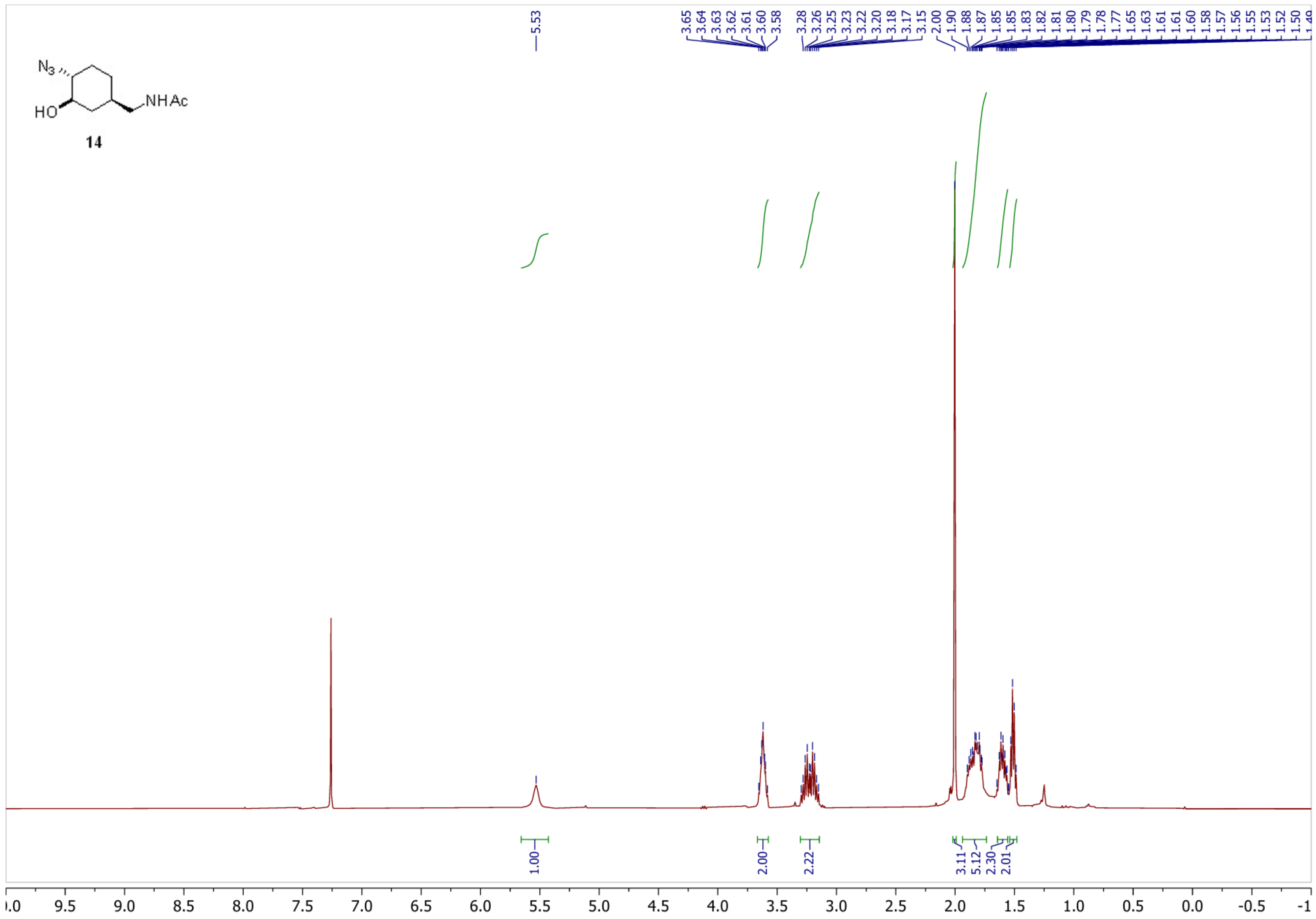

N3
HO
NHAc
14
5.53
3.65
3.64
3.63
3.62
3.61
3.60
3.58
3.28
3.26
3.25
3.23
3.22
3.20
3.18
3.17
3.15
2.00
1.90
1.88
1.87
1.85
1.85
1.83
1.82
1.81
1.80
1.79
1.78
1.77
1.65
1.63
1.61
1.61
1.60
1.58
1.57
1.56
1.55
1.53
1.52
1.50
1.00
2.00
2.22
3.11
5.12
2.30
2.01
9.5
9.0
8.5
8.0
7.5
7.0
6.5
6.0
5.5
5.0
4.5
4.0
3.5
3.0
2.5
2.0
1.5
1.0
0.5
0.0
-0.5

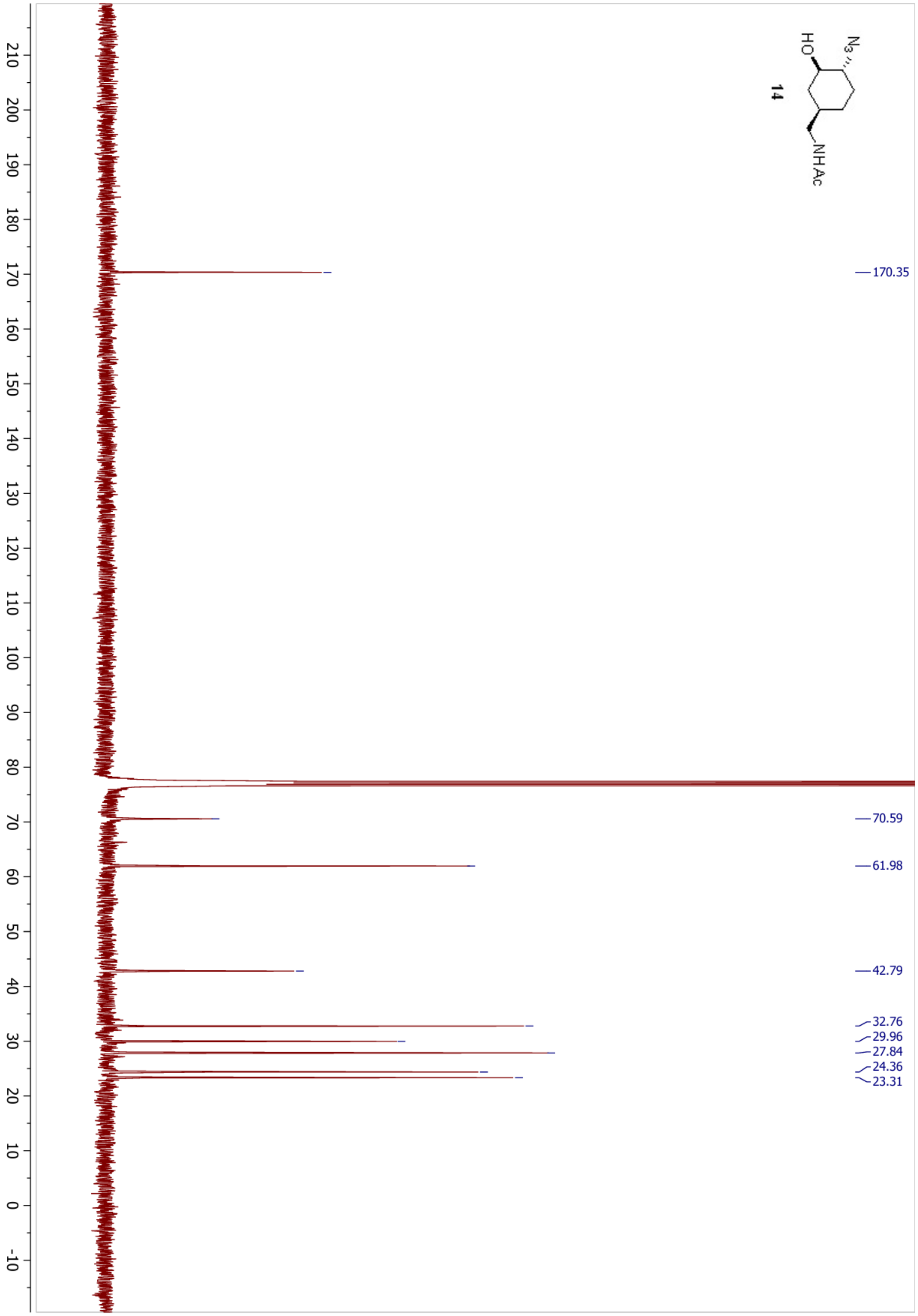

N3
HO
NHAc
14
170.35
70.59
61.98
42.79
32.76
29.96
27.84
24.36
23.31

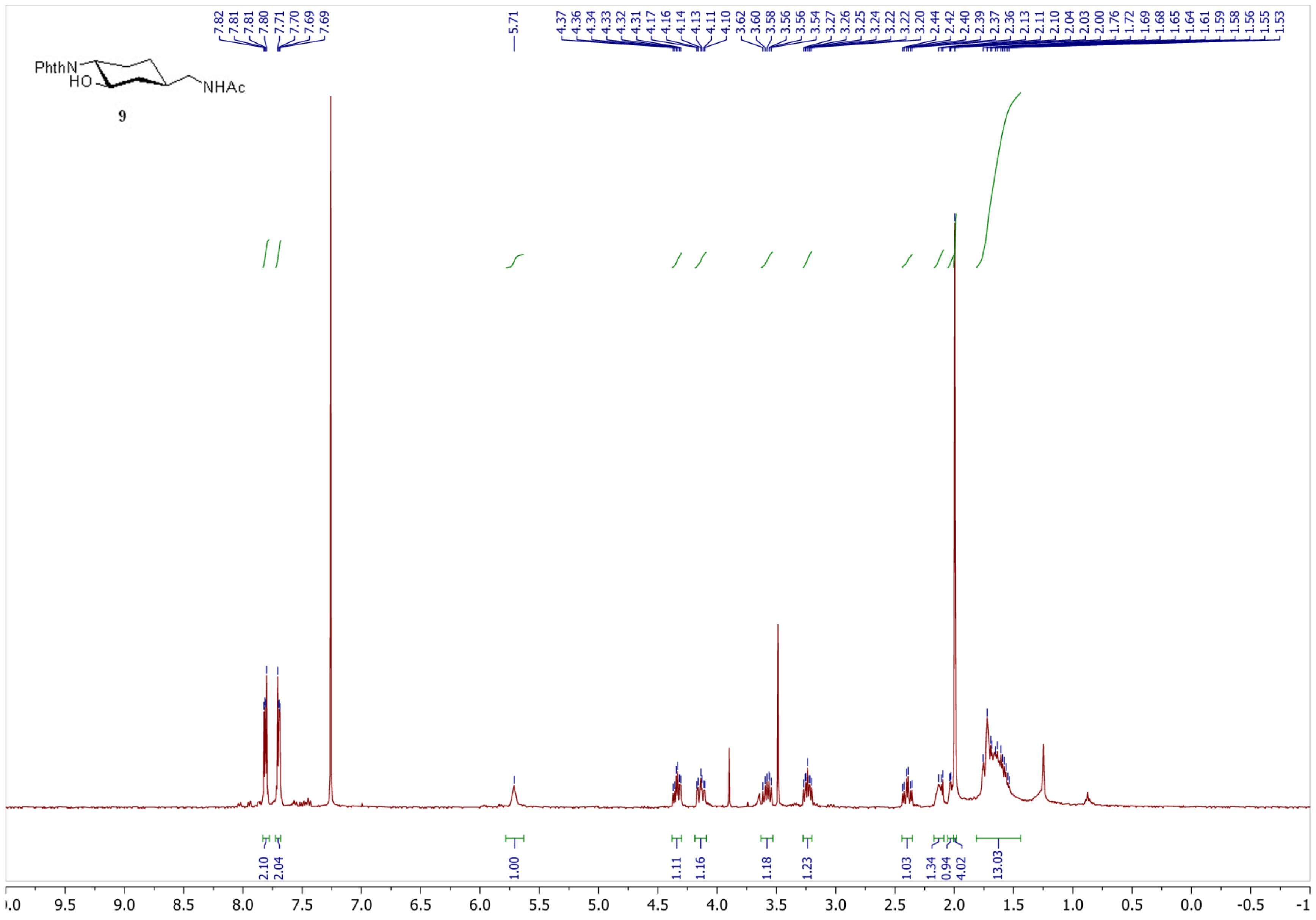
PhthN
HO
NHAc
9
7.82
7.81
7.81
7.80
7.71
7.70
7.69
7.69
5.71
4.37
4.36
4.34
4.33
4.32
4.31
4.17
4.16
4.14
4.13
4.11
4.10
3.62
3.60
3.58
3.56
3.56
3.54
3.27
3.26
3.25
3.24
3.22
3.22
3.20
2.44
2.42
2.40
2.39
2.37
2.36
2.13
2.11
2.10
2.04
2.03
2.00
1.76
1.72
1.69
1.68
1.65
1.64
1.61
1.59
1.58
1.56
1.55
1.53
2.10
2.04
1.00
1.11
1.16
1.18
1.23
1.03
1.34
0.94
4.02
13.03
10.0 9.5 9.0 8.5 8.0 7.5 7.0 6.5 6.0 5.5 5.0 4.5 4.0 3.5 3.0 2.5 2.0 1.5 1.0 0.5 0.0 -0.5 -1

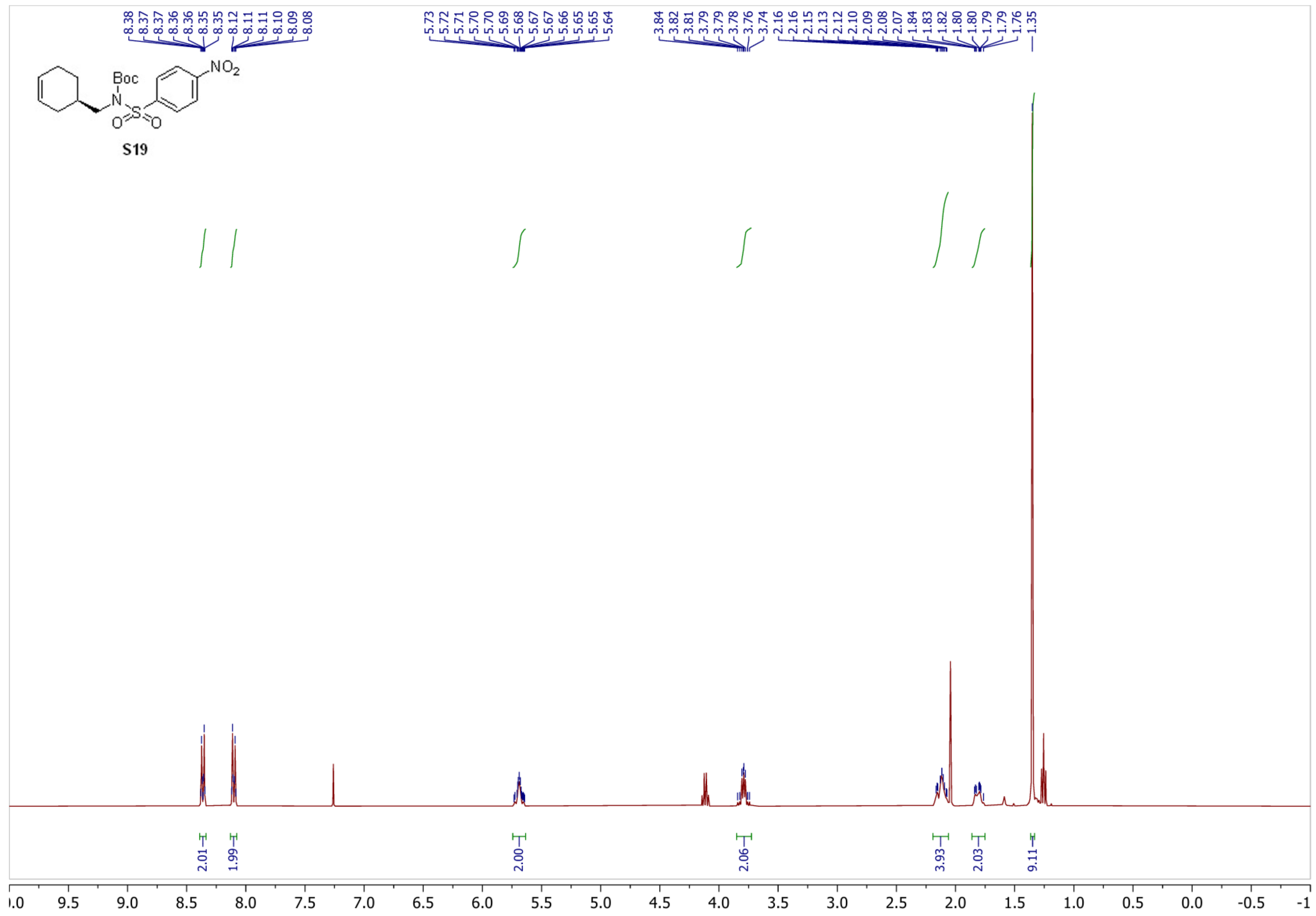

8.38 8.37 8.37 8.36 8.36 8.35 8.35 8.12 8.11 8.11 8.10 8.09 8.08
5.73 5.72 5.71 5.70 5.70 5.69 5.68 5.67 5.67 5.66 5.65 5.65 5.64
3.84 3.82 3.81 3.79 3.79 3.78 3.76 3.74 2.16 2.16 2.15 2.13 2.12 2.10 2.09 2.08 2.07 1.84 1.83 1.82 1.80 1.80 1.79 1.79 1.76 1.35
Boc
N
S
O
O
$NO_2$
S19
2.01
1.99
2.00
2.06
3.93
2.03
9.11
.0 9.5 9.0 8.5 8.0 7.5 7.0 6.5 6.0 5.5 5.0 4.5 4.0 3.5 3.0 2.5 2.0 1.5 1.0 0.5 0.0 -0.5 -1

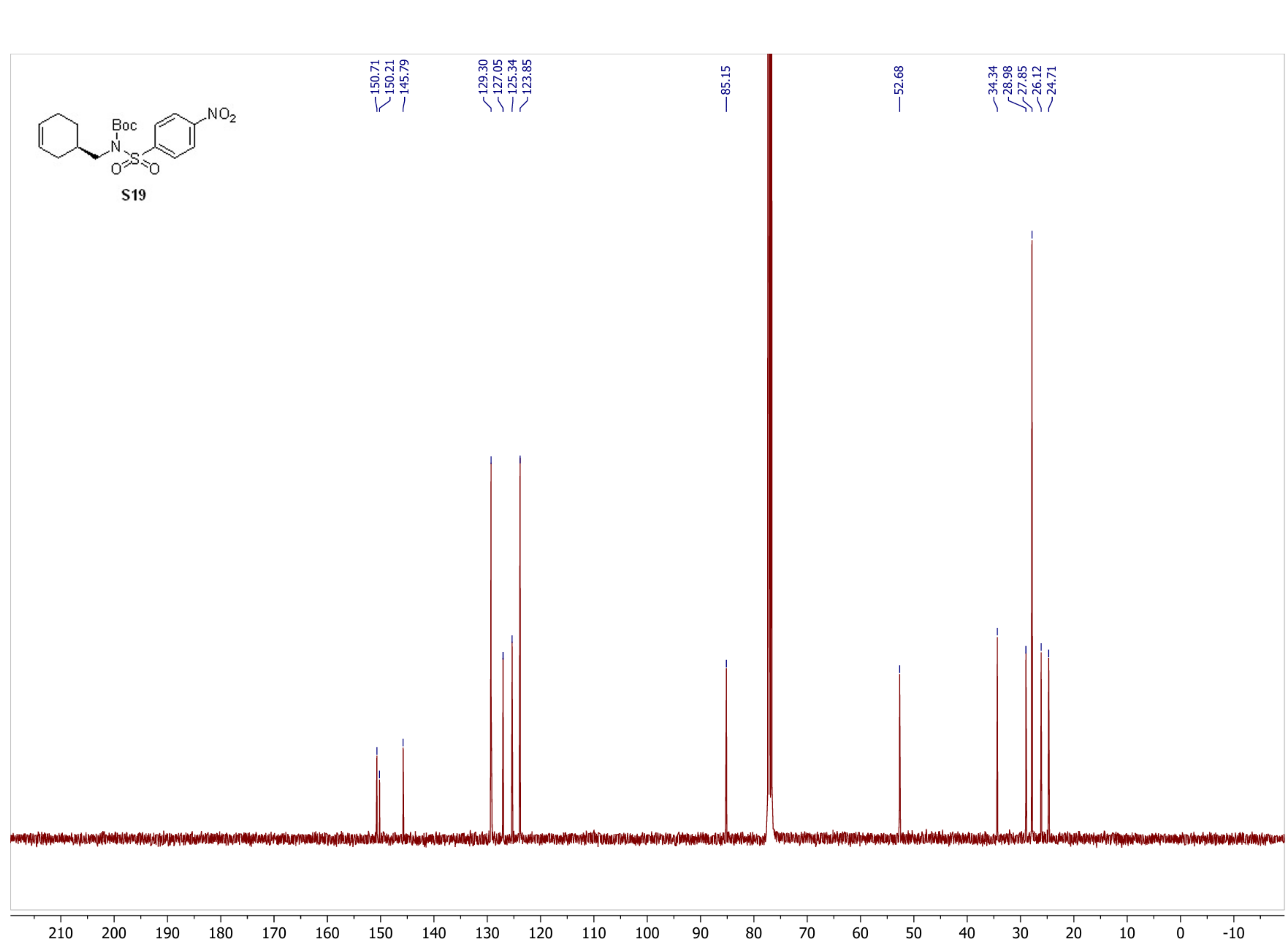

150.71
150.21
145.79
129.30
127.05
125.34
123.85
85.15
52.68
34.34
28.98
27.85
26.12
24.71
Boc
N
S
O
O
NO2
S19
210 200 190 180 170 160 150 140 130 120 110 100 90 80 70 60 50 40 30 20 10 0 -10

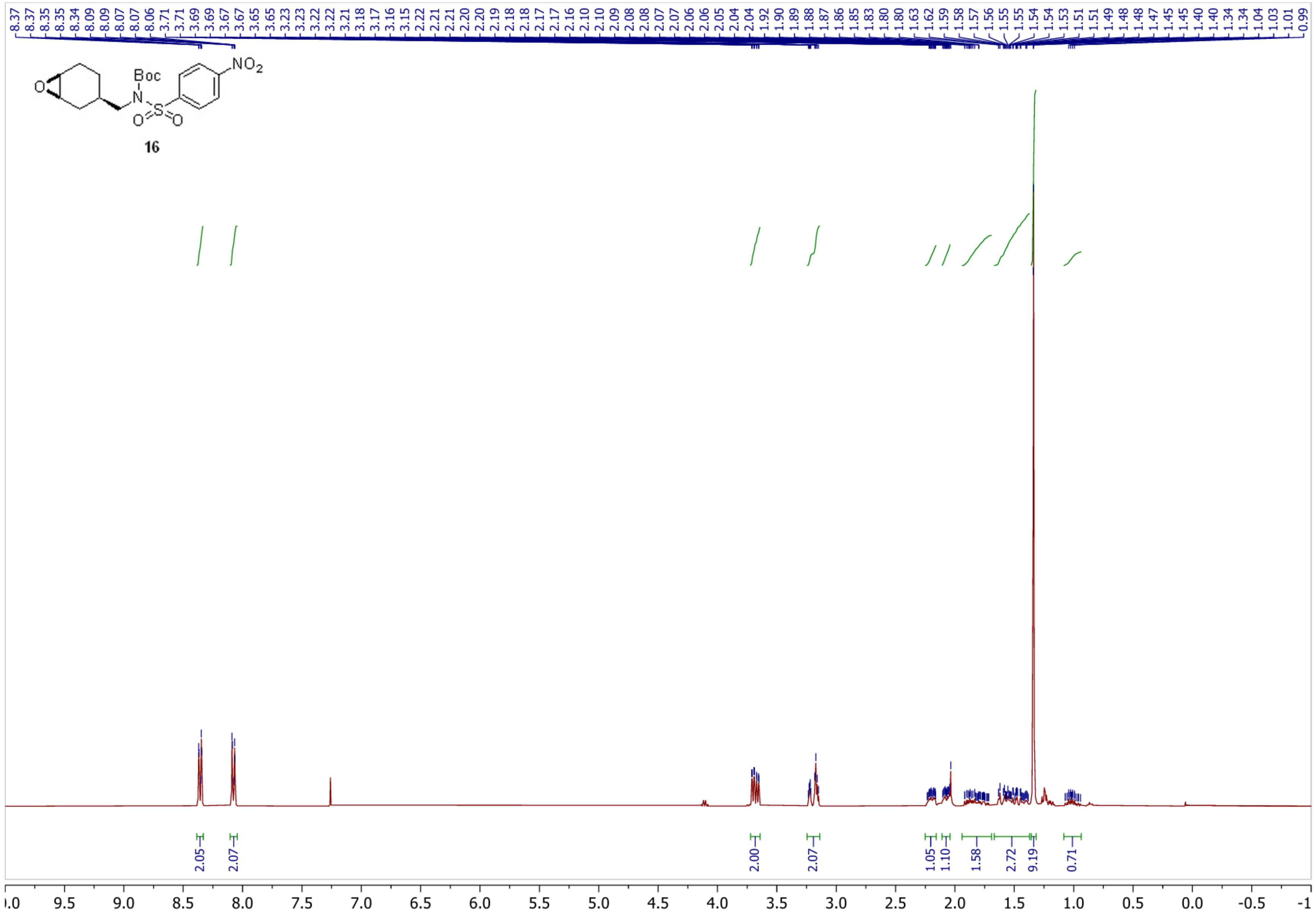

8.37
8.37
8.35
8.35
8.34
8.09
8.09
8.07
8.07
8.06
3.71
3.71
3.69
3.69
3.67
3.67
3.65
3.65
3.23
3.23
3.22
3.22
3.21
3.18
3.17
3.16
3.15
2.22
2.21
2.21
2.20
2.20
2.19
2.18
2.18
2.17
2.17
2.16
2.10
2.10
2.09
2.08
2.08
2.07
2.07
2.06
2.06
2.05
2.04
2.04
1.92
1.90
1.89
1.88
1.87
1.86
1.85
1.83
1.80
1.80
1.63
1.62
1.59
1.58
1.57
1.56
1.55
1.55
1.54
1.54
1.53
1.51
1.51
1.49
1.48
1.48
1.47
1.45
1.45
1.40
1.40
1.34
1.34
1.04
1.03
1.01
0.99
Boc
N
O
S
O
O
$NO_2$
16
2.05
2.07
2.00
2.07
1.05
1.10
1.58
2.72
9.19
0.71
.0
9.5
9.0
8.5
8.0
7.5
7.0
6.5
6.0
5.5
5.0
4.5
4.0
3.5
3.0
2.5
2.0
1.5
1.0
0.5
0.0
-0.5
-1

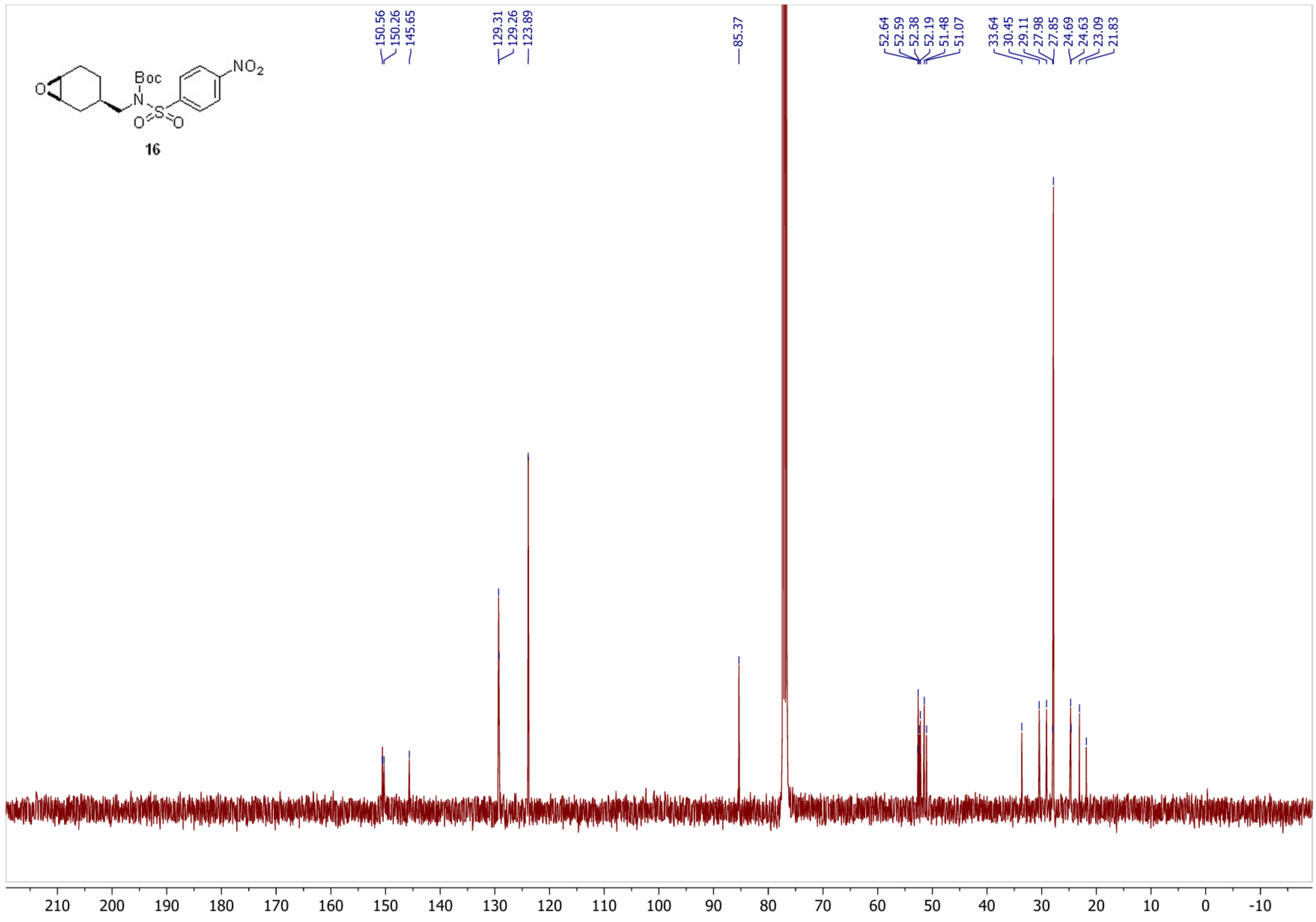
Boc
N
O=S=O
NO2
16
150.56
150.26
145.65
129.31
129.26
123.89
85.37
52.64
52.59
52.38
52.19
51.48
51.07
33.64
30.45
29.11
27.98
27.85
24.69
24.63
23.09
21.83
210
200
190
180
170
160
150
140
130
120
110
100
90
80
70
60
50
40
30
20
10
0
-10

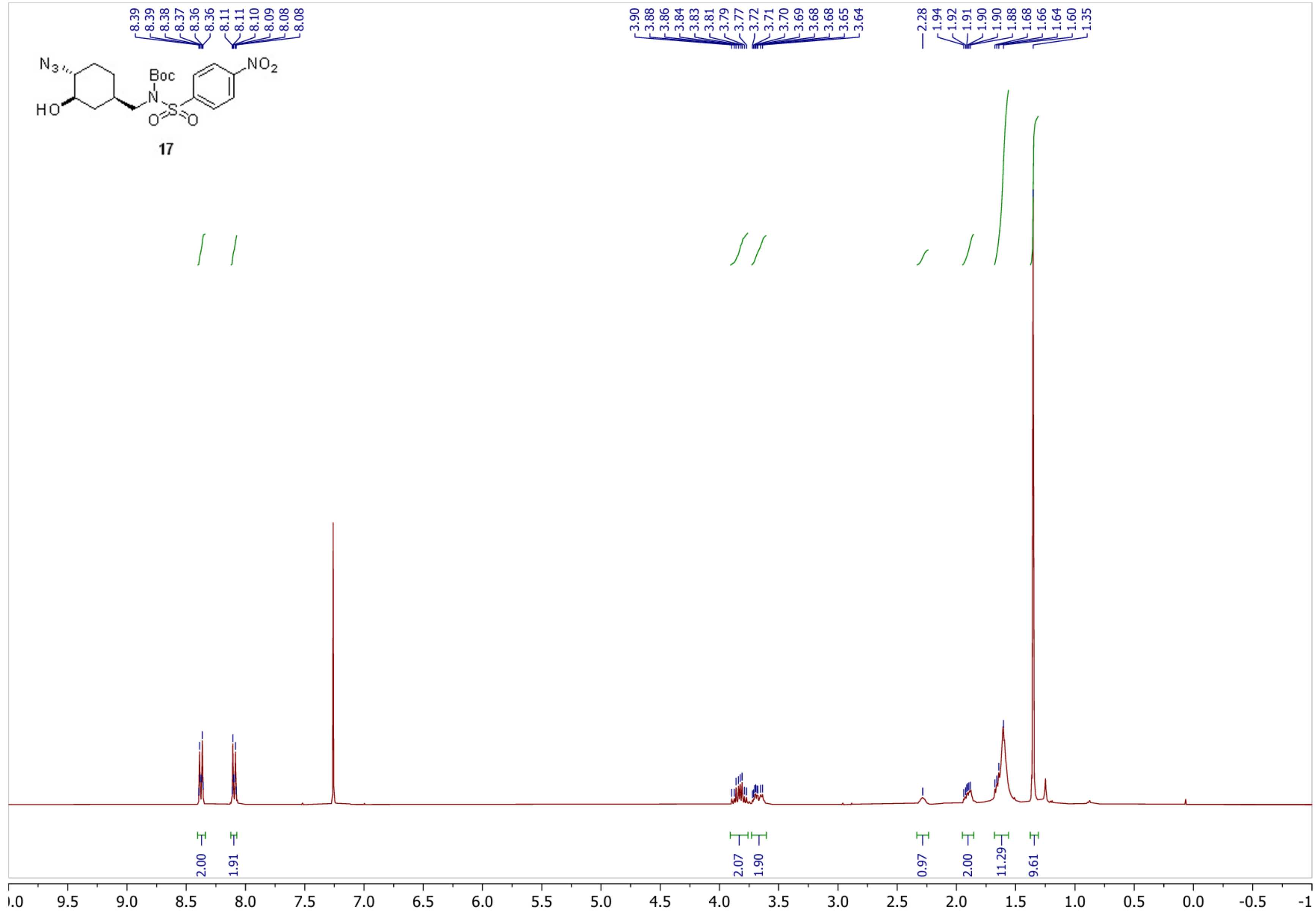
N3
HO
Boc
N
S
O
O
NO2
17
8.39
8.39
8.38
8.37
8.36
8.36
8.11
8.11
8.10
8.09
8.08
8.08
3.90
3.88
3.86
3.84
3.83
3.81
3.79
3.77
3.72
3.71
3.70
3.69
3.68
3.68
3.65
3.64
2.28
1.94
1.92
1.91
1.90
1.90
1.88
1.68
1.66
1.64
1.60
1.35
2.00
1.91
2.07
1.90
0.97
2.00
11.29
9.61
9.5
9.0
8.5
8.0
7.5
7.0
6.5
6.0
5.5
5.0
4.5
4.0
3.5
3.0
2.5
2.0
1.5
1.0
0.5
0.0
-0.5

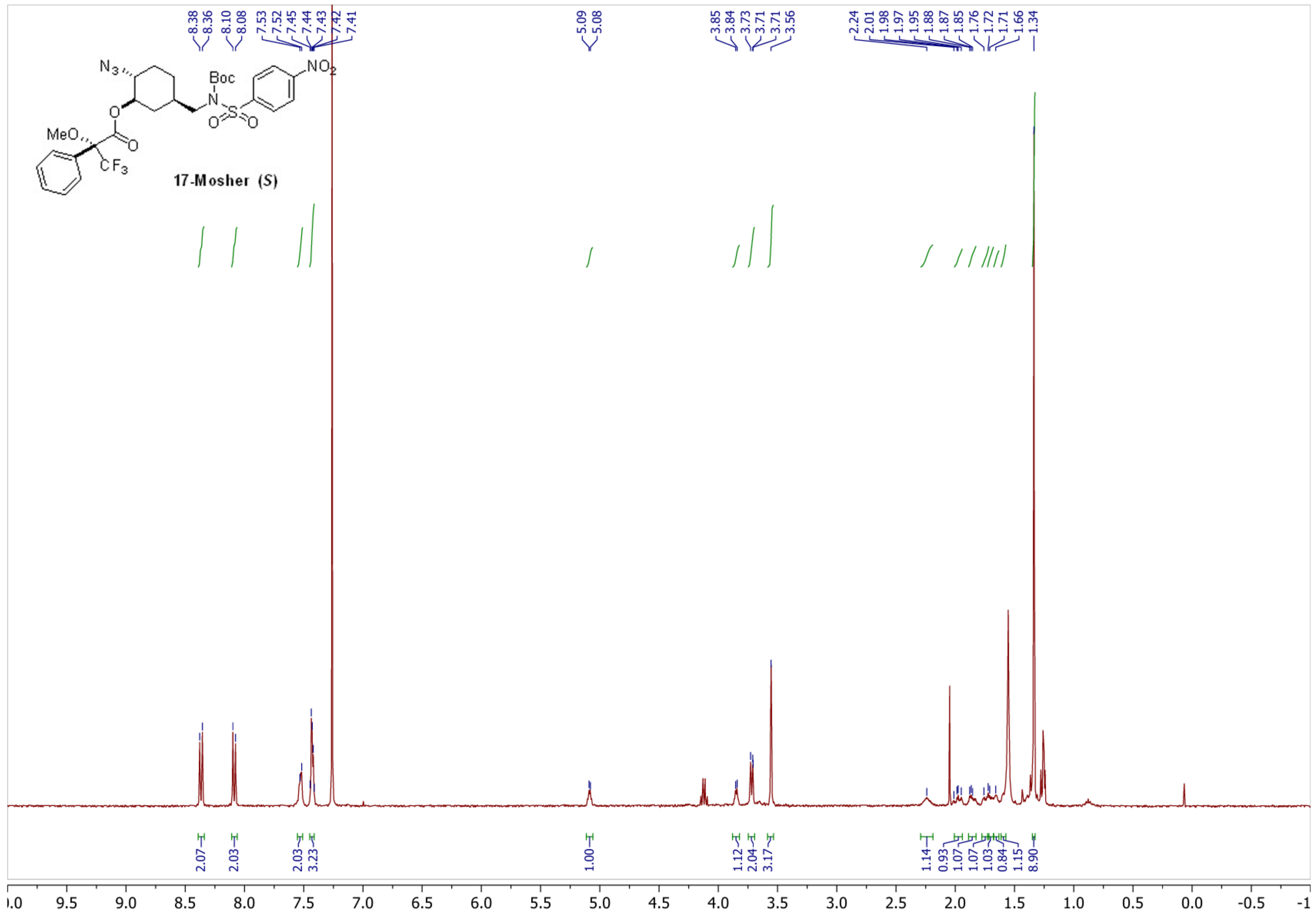

17-Mosher (S)
8.38
8.36
8.10
8.08
7.53
7.52
7.45
7.44
7.43
7.42
7.41
5.09
5.08
3.85
3.84
3.73
3.71
3.71
3.56
2.24
2.01
1.98
1.97
1.95
1.88
1.87
1.85
1.76
1.72
1.71
1.66
1.34
2.07
2.03
2.03
3.23
1.00
1.12
2.04
3.17
1.14
0.93
1.07
1.07
1.03
0.84
1.15
8.90

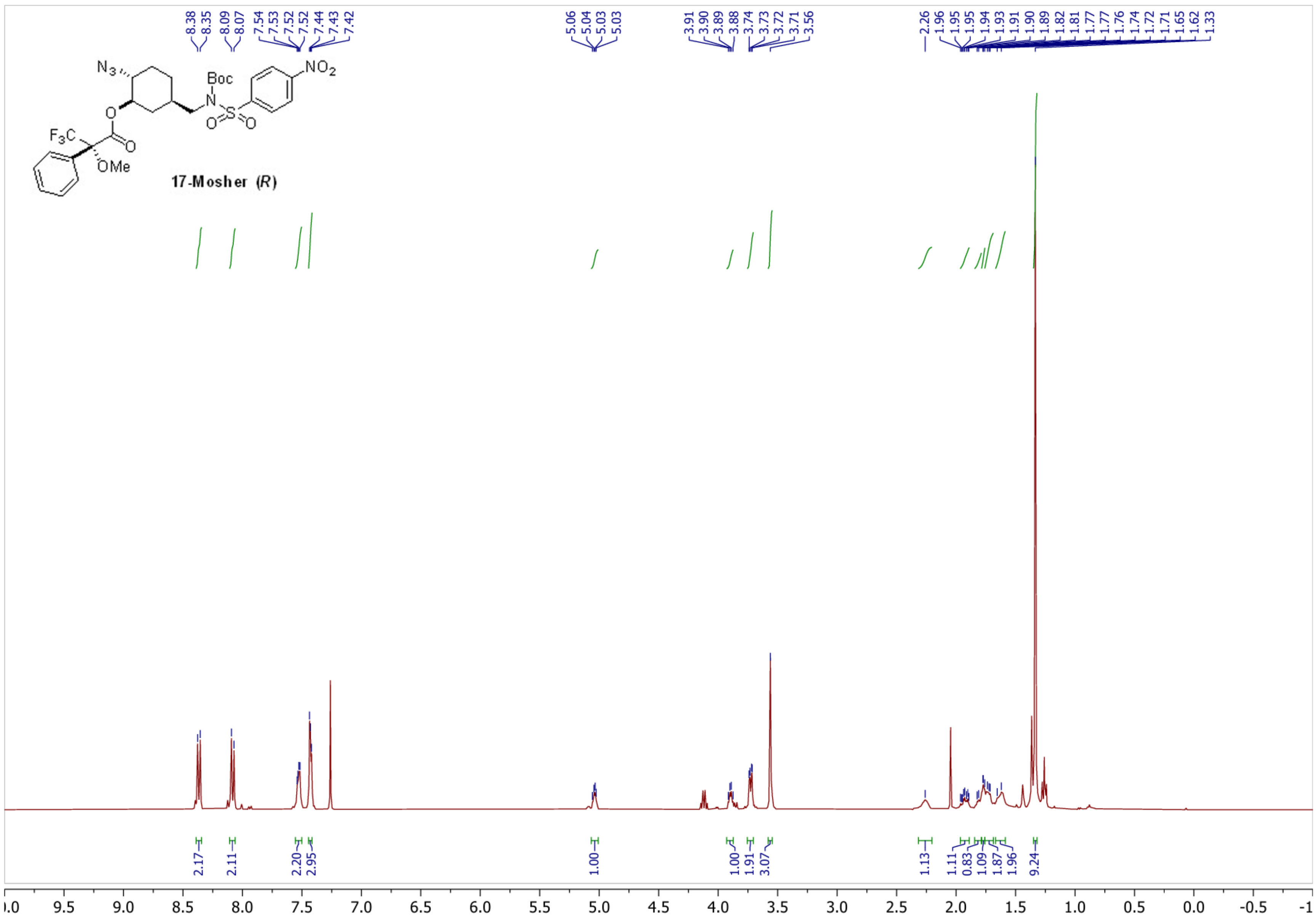
17-Mosher (R)
N3
Boc
NO2
F3C
OMe
8.38
8.35
8.09
8.07
7.54
7.53
7.52
7.52
7.44
7.43
7.42
5.06
5.04
5.03
5.03
3.91
3.90
3.89
3.88
3.74
3.73
3.72
3.71
3.56
2.26
1.96
1.95
1.95
1.94
1.93
1.91
1.90
1.89
1.82
1.81
1.77
1.77
1.76
1.74
1.72
1.71
1.65
1.62
1.33
2.17
2.11
2.20
2.95
1.00
1.00
1.91
3.07
1.13
1.11
0.83
1.09
1.87
1.96
9.24
.0
9.5
9.0
8.5
8.0
7.5
7.0
6.5
6.0
5.5
5.0
4.5
4.0
3.5
3.0
2.5
2.0
1.5
1.0
0.5
0.0
-0.5
-1

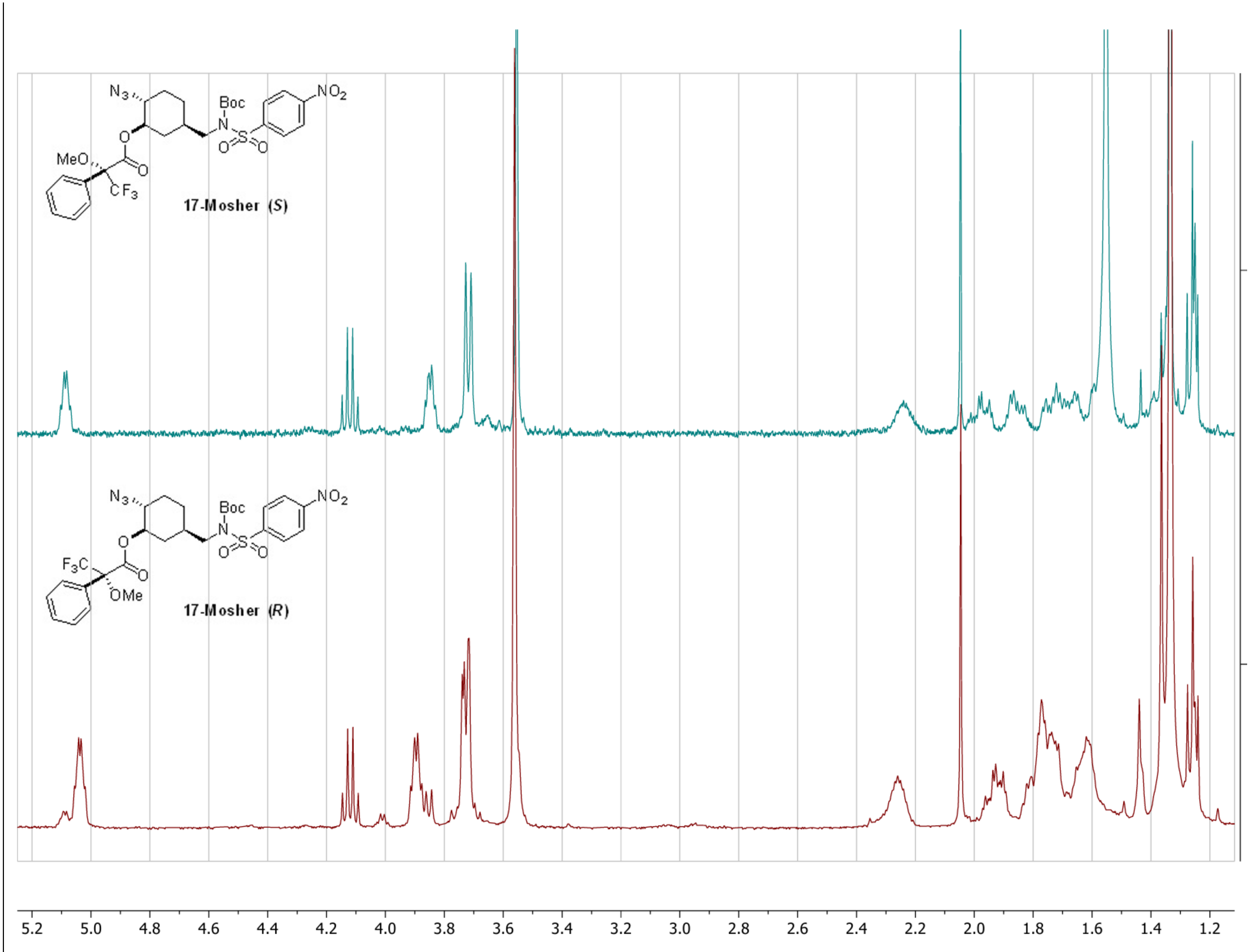
N3
Boc
N
S
NO2
MeO
CF3
17-Mosher (S)
F3C
OMe
17-Mosher (R)
5.2
5.0
4.8
4.6
4.4
4.2
4.0
3.8
3.6
3.4
3.2
3.0
2.8
2.6
2.4
2.2
2.0
1.8
1.6
1.4
1.2

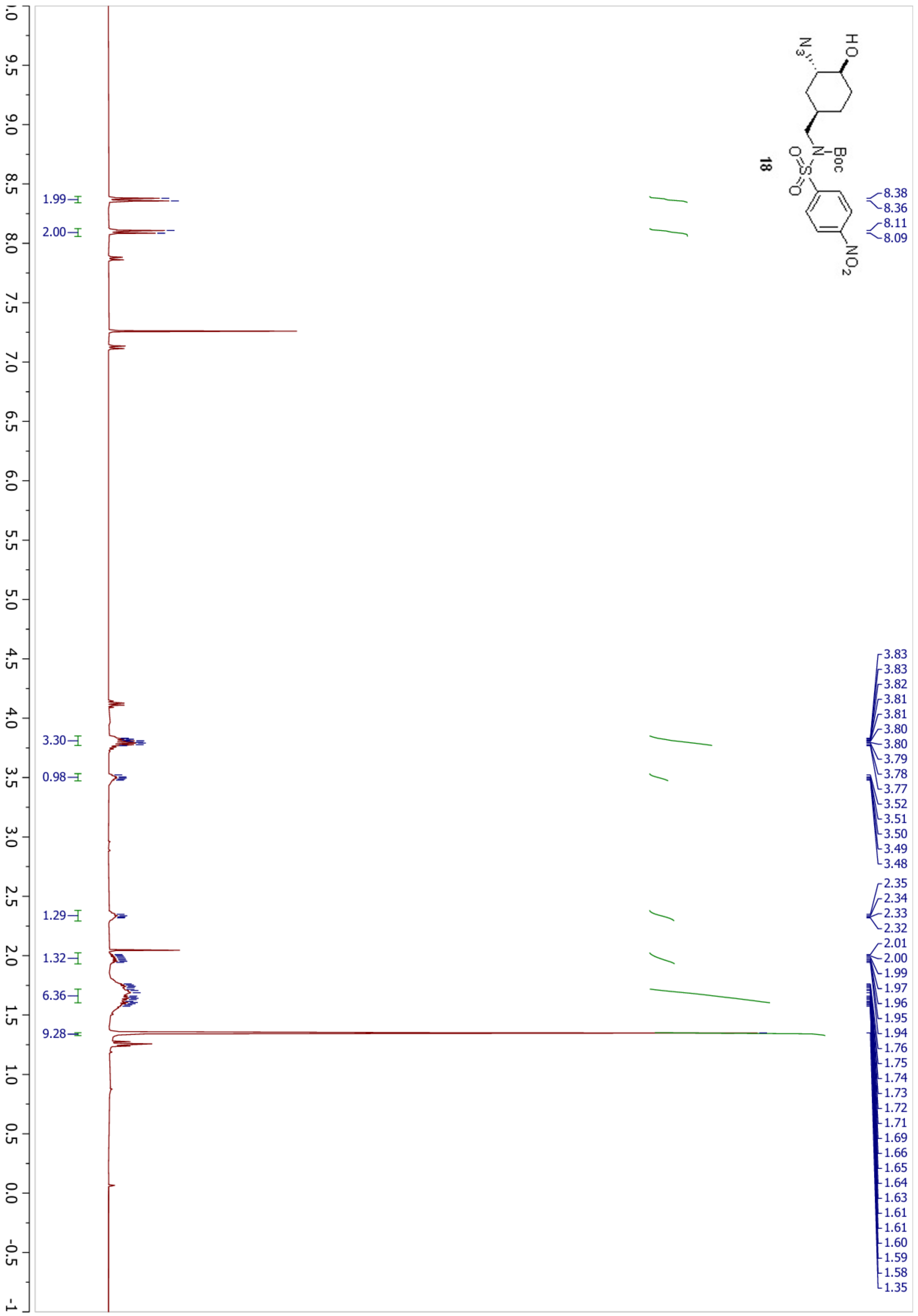

18
HO
N3
Boc
N
O=S=O
NO2
8.38
8.36
8.11
8.09
3.83
3.83
3.82
3.81
3.81
3.80
3.80
3.79
3.78
3.77
3.52
3.51
3.50
3.49
3.48
2.35
2.34
2.33
2.32
2.01
2.00
1.99
1.97
1.96
1.95
1.94
1.76
1.75
1.74
1.73
1.72
1.71
1.69
1.66
1.65
1.64
1.63
1.61
1.61
1.60
1.59
1.58
1.35
1.99
2.00
3.30
0.98
1.29
1.32
6.36
9.28
9.5
9.0
8.5
8.0
7.5
7.0
6.5
6.0
5.5
5.0
4.5
4.0
3.5
3.0
2.5
2.0
1.5
1.0
0.5
0.0
-0.5
-1

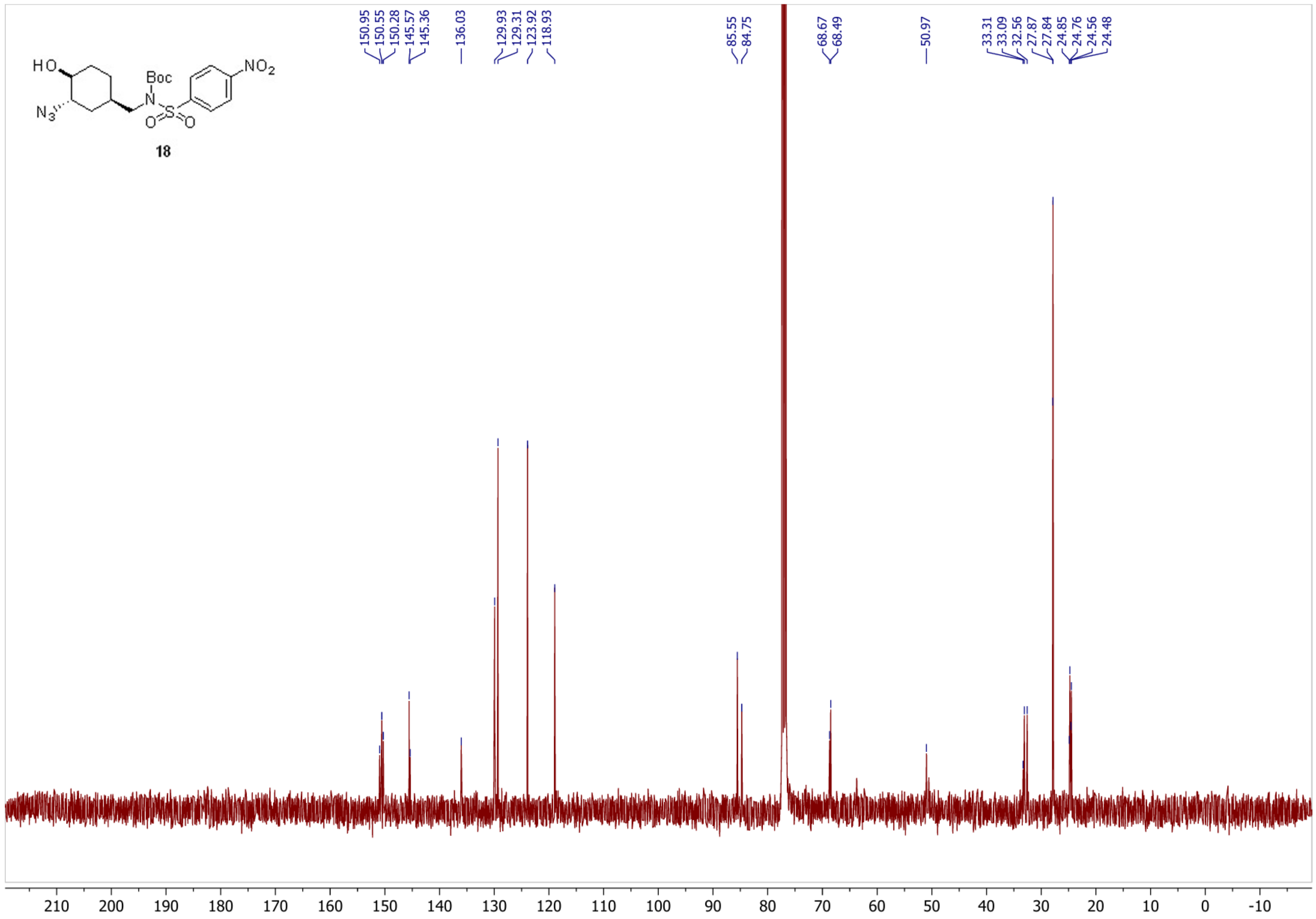
HO
N3
Boc
N
O
S
O
NO2
18
150.95
150.55
150.28
145.57
145.36
136.03
129.93
129.31
123.92
118.93
85.55
84.75
68.67
68.49
50.97
33.31
33.09
32.56
27.87
27.84
24.85
24.76
24.56
24.48
210
200
190
180
170
160
150
140
130
120
110
100
90
80
70
60
50
40
30
20
10
0
-10

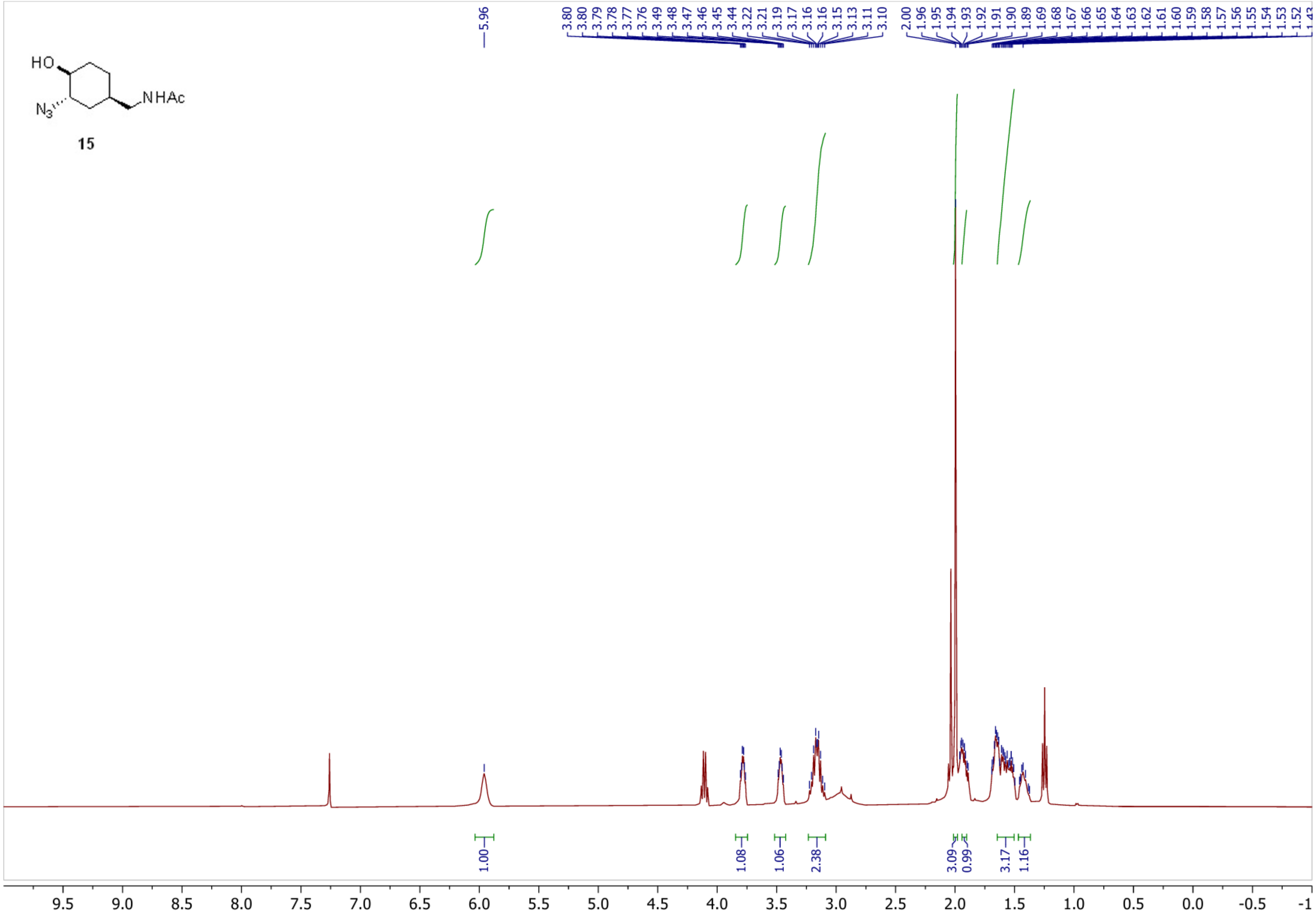
HO
N3
NHAc
15
5.96
3.80
3.80
3.79
3.78
3.77
3.76
3.49
3.48
3.47
3.46
3.45
3.44
3.22
3.21
3.19
3.17
3.16
3.16
3.15
3.13
3.11
3.10
2.00
1.96
1.95
1.94
1.93
1.92
1.91
1.90
1.89
1.69
1.68
1.67
1.66
1.65
1.64
1.63
1.62
1.61
1.60
1.59
1.58
1.57
1.56
1.55
1.54
1.53
1.52
1.00
1.08
1.06
2.38
3.09
0.99
3.17
1.16
9.5
9.0
8.5
8.0
7.5
7.0
6.5
6.0
5.5
5.0
4.5
4.0
3.5
3.0
2.5
2.0
1.5
1.0
0.5
0.0
-0.5
-1

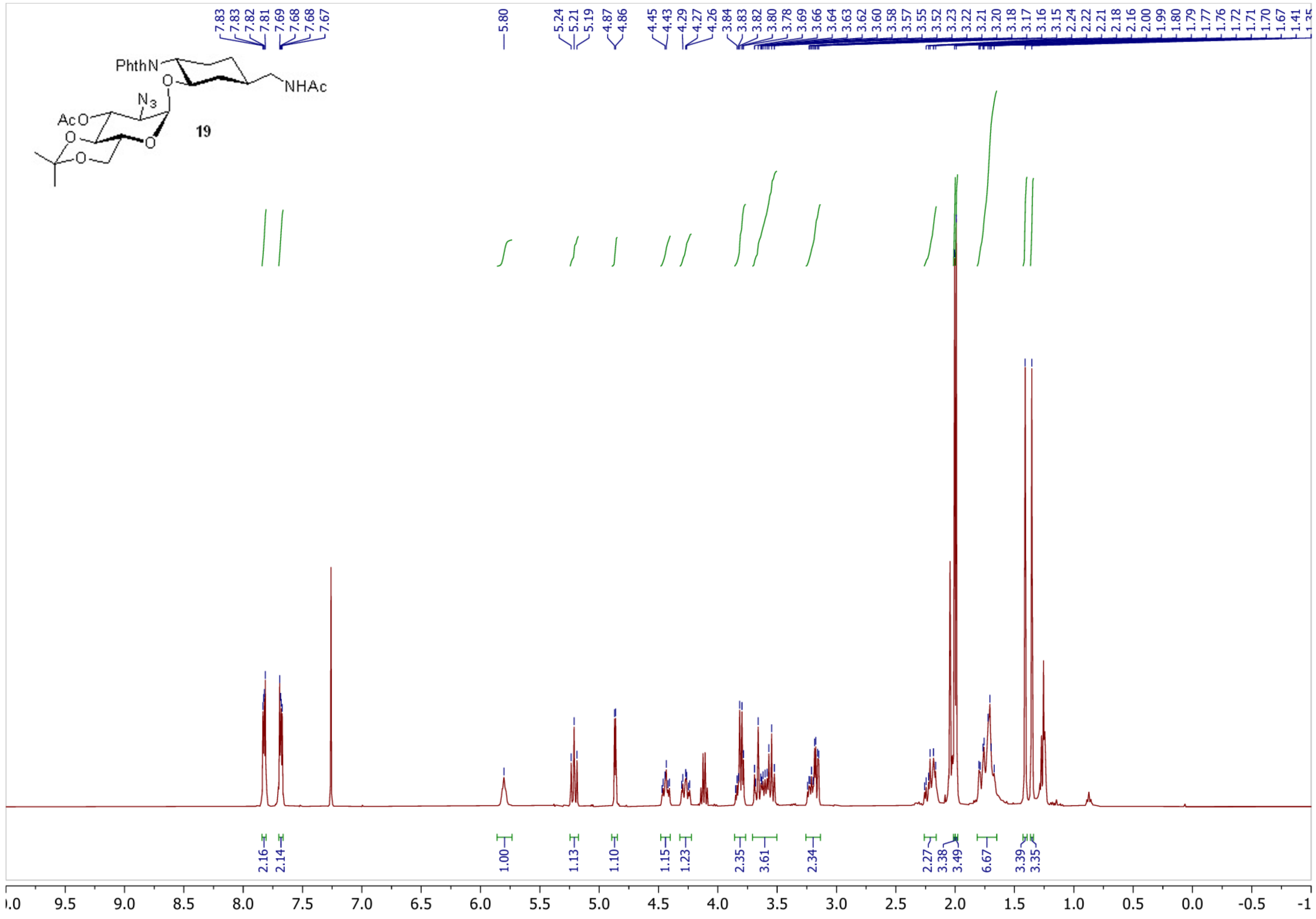

PhthN
NHAc
$N_3$
AcO
O
19
7.83
7.83
7.82
7.81
7.69
7.68
7.68
7.67
5.80
5.24
5.21
5.19
4.87
4.86
4.45
4.43
4.29
4.27
4.26
3.84
3.83
3.82
3.80
3.78
3.69
3.66
3.64
3.63
3.62
3.60
3.58
3.57
3.55
3.52
3.23
3.22
3.21
3.20
3.18
3.17
3.16
3.15
2.24
2.22
2.21
2.18
2.16
2.00
1.99
1.80
1.79
1.77
1.76
1.72
1.71
1.70
1.67
1.41
2.16
2.14
1.00
1.13
1.10
1.15
1.23
2.35
3.61
2.34
2.27
3.38
3.49
6.67
3.39
3.35
9.5
9.0
8.5
8.0
7.5
7.0
6.5
6.0
5.5
5.0
4.5
4.0
3.5
3.0
2.5
2.0
1.5
1.0
0.5
0.0
-0.5

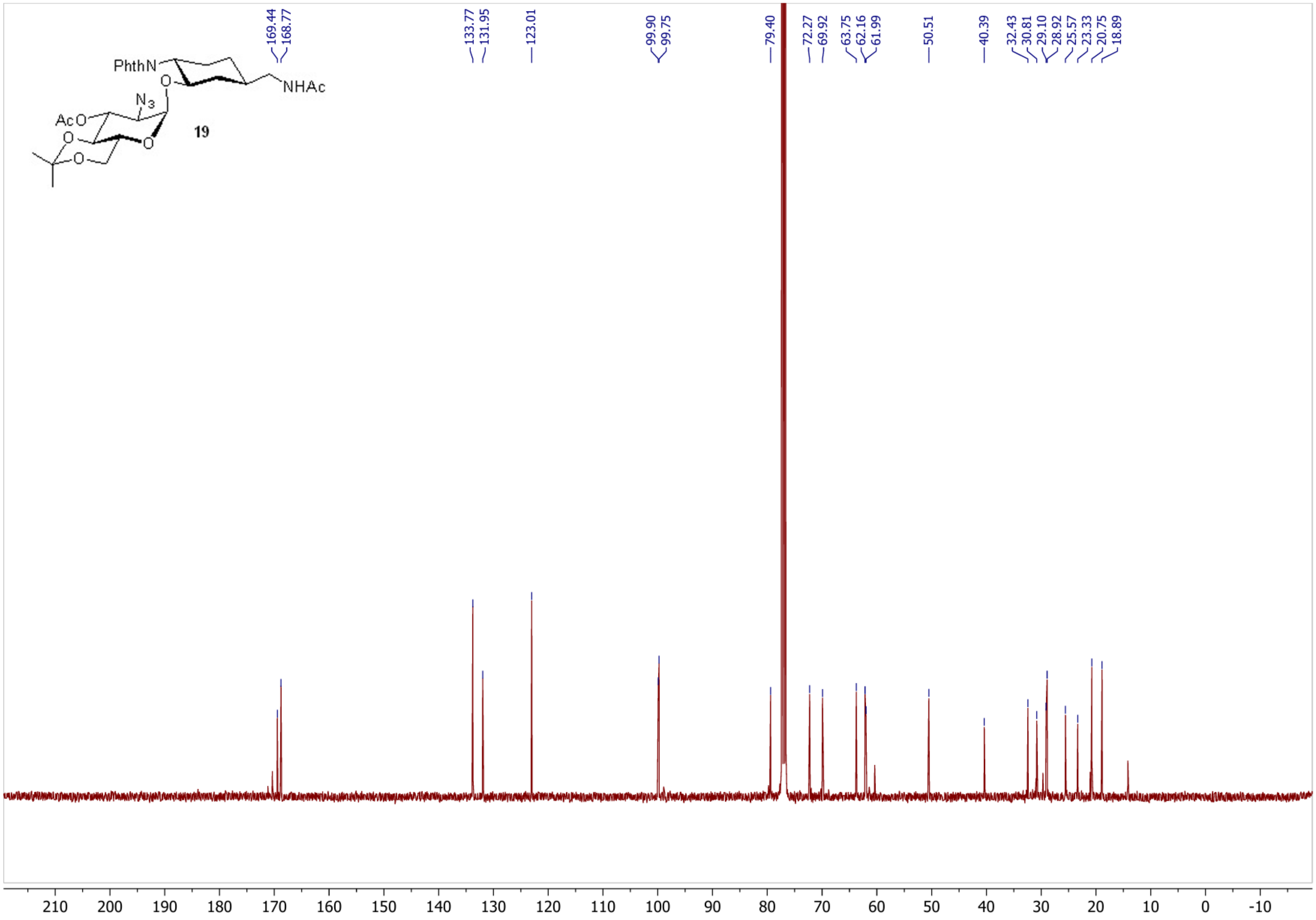

PhthN
NHAc
$N_3$
AcO
O
19
169.44
168.77
133.77
131.95
123.01
99.90
99.75
79.40
72.27
69.92
63.75
62.16
61.99
50.51
40.39
32.43
30.81
29.10
28.92
25.57
23.33
20.75
18.89
210
200
190
180
170
160
150
140
130
120
110
100
90
80
70
60
50
40
30
20
10
0
-10

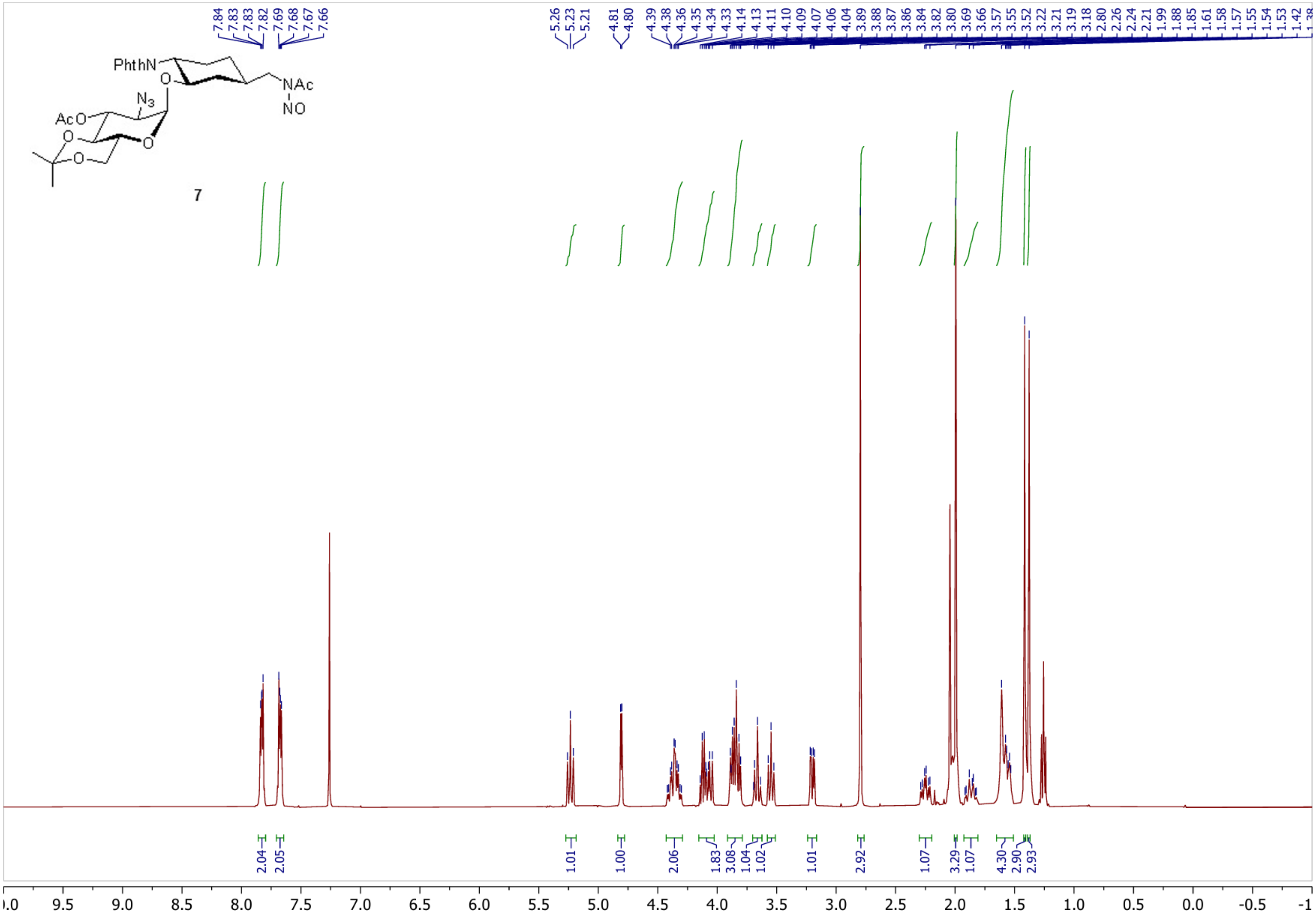
PhthN
O
N3
AcO
O
O
O
NAc
NO
7
7.84
7.83
7.83
7.82
7.69
7.68
7.67
7.66
5.26
5.23
5.21
4.81
4.80
4.39
4.38
4.36
4.35
4.34
4.33
4.14
4.13
4.11
4.10
4.09
4.07
4.06
4.04
3.89
3.88
3.87
3.86
3.84
3.82
3.80
3.69
3.66
3.57
3.55
3.52
3.22
3.21
3.19
3.18
2.80
2.26
2.24
2.21
1.99
1.88
1.85
1.61
1.58
1.57
1.55
1.54
1.53
1.42
2.04
2.05
1.01
1.00
2.06
1.83
3.08
1.04
1.02
1.01
2.92
1.07
3.29
1.07
4.30
2.90
2.93
9.5
9.0
8.5
8.0
7.5
7.0
6.5
6.0
5.5
5.0
4.5
4.0
3.5
3.0
2.5
2.0
1.5
1.0
0.5
0.0
-0.5

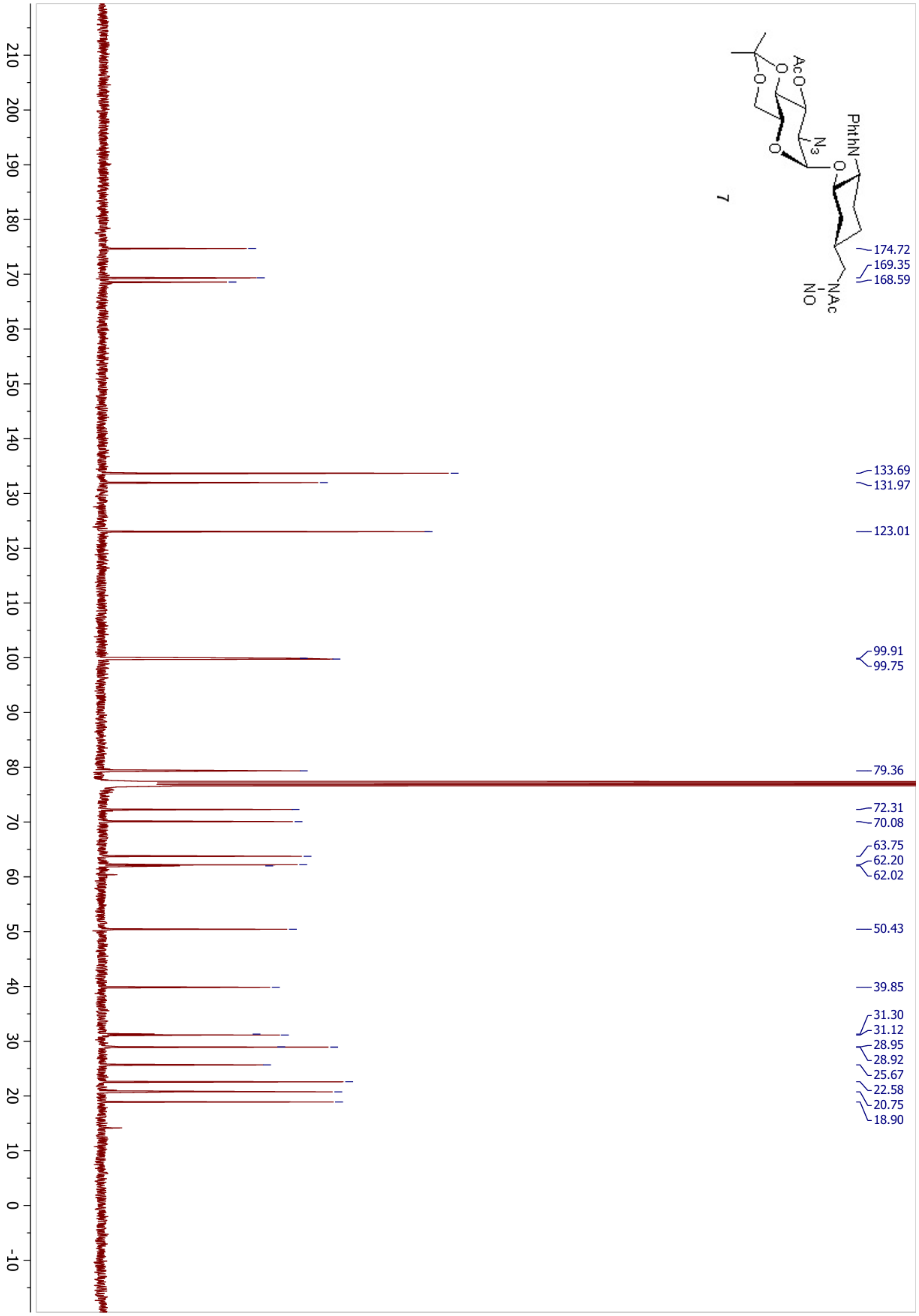

PhthN
AcO
N3
NAc
NO
7
174.72
169.35
168.59
133.69
131.97
123.01
99.91
99.75
79.36
72.31
70.08
63.75
62.20
62.02
50.43
39.85
31.30
31.12
28.95
28.92
25.67
22.58
20.75
18.90
210
200
190
180
170
160
150
140
130
120
110
100
90
80
70
60
50
40
30
20
10
0
-10

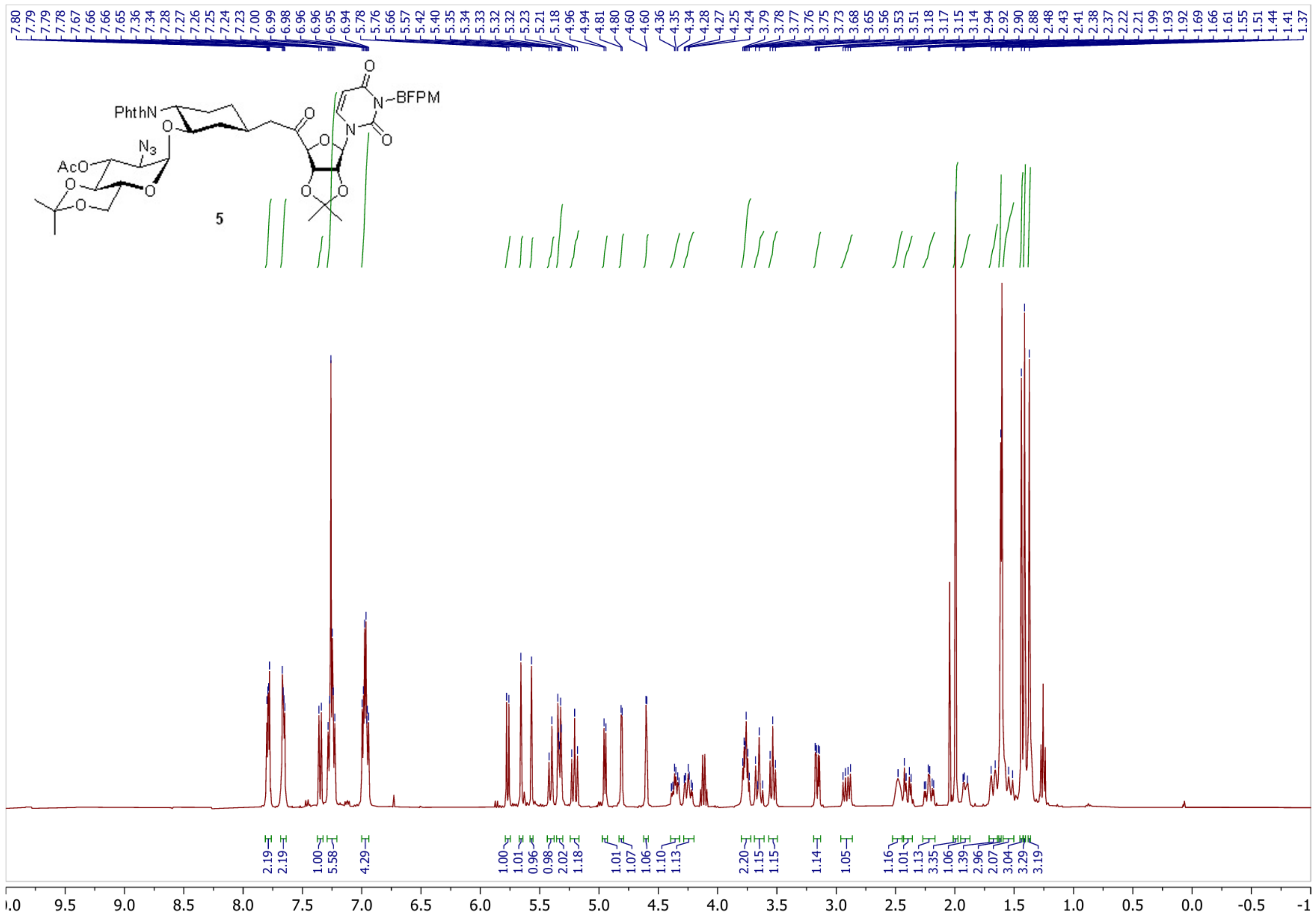
7.80 7.79 7.79 7.78 7.67 7.66 7.66 7.65 7.36 7.34 7.28 7.27 7.26 7.25 7.24 7.23 7.00 6.99 6.98 6.96 6.96 6.95 6.94 5.78 5.76 5.66 5.57 5.42 5.40 5.35 5.34 5.33 5.32 5.32 5.23 5.21 5.18 4.96 4.94 4.81 4.80 4.60 4.60 4.36 4.35 4.34 4.28 4.27 4.25 4.24 3.79 3.78 3.77 3.76 3.75 3.73 3.68 3.65 3.56 3.53 3.51 3.18 3.17 3.15 3.14 2.94 2.92 2.90 2.88 2.48 2.43 2.41 2.38 2.37 2.22 2.21 1.99 1.93 1.92 1.69 1.66 1.61 1.55 1.51 1.44 1.41 1.37
PhthN
N3
AcO
O
N
BFPM
5
2.19 2.19 1.00 5.58 4.29 1.00 1.01 0.96 0.98 2.02 1.18 1.01 1.07 1.06 1.10 1.13 2.20 1.15 1.15 1.14 1.05 1.16 1.01 1.13 3.35 1.06 1.39 2.96 2.07 3.04 3.29 3.19
10.0 9.5 9.0 8.5 8.0 7.5 7.0 6.5 6.0 5.5 5.0 4.5 4.0 3.5 3.0 2.5 2.0 1.5 1.0 0.5 0.0 -0.5 -1

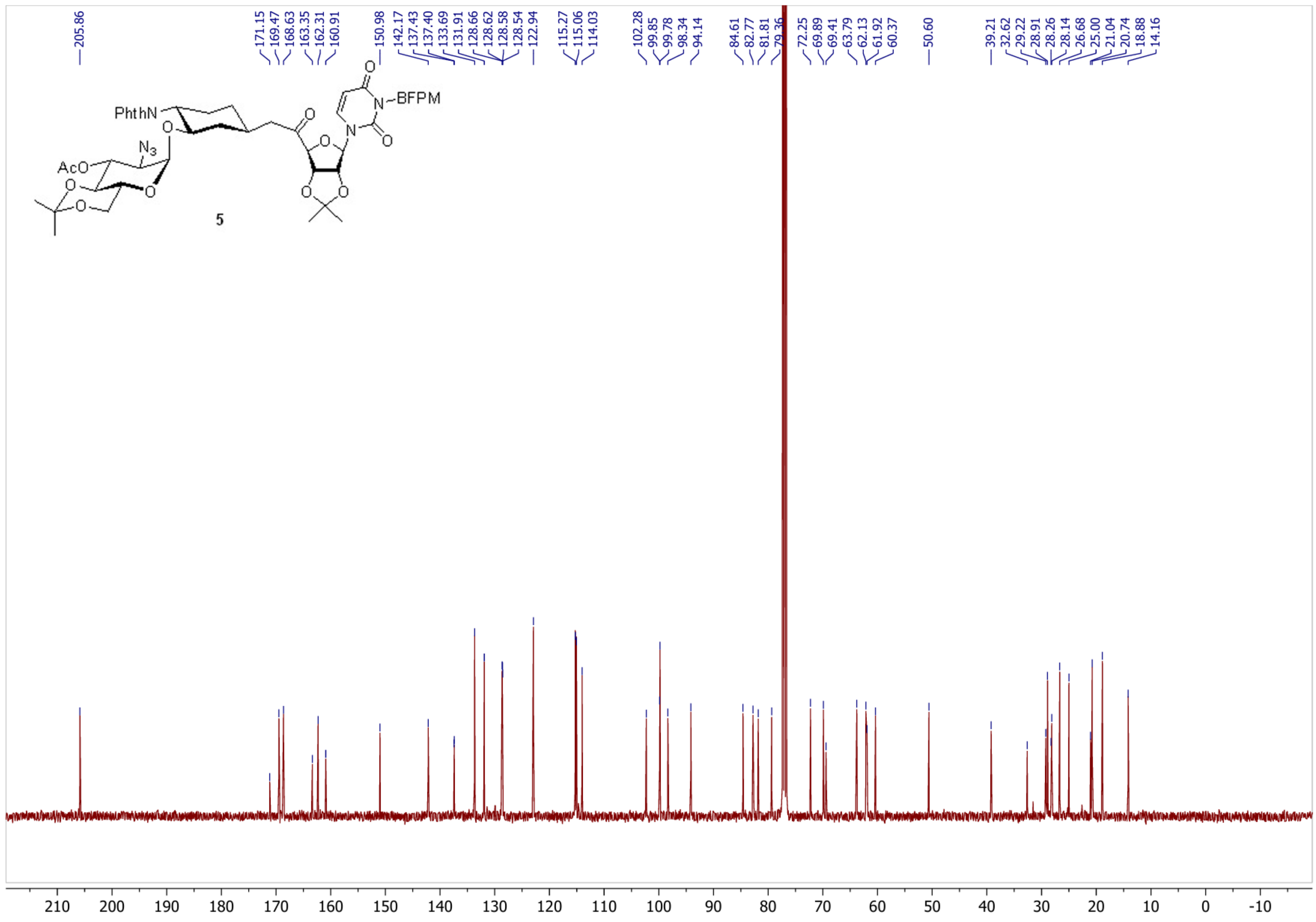
205.86
171.15
169.47
168.63
163.35
162.31
160.91
150.98
142.17
137.43
137.40
133.69
131.91
128.66
128.62
128.58
128.54
122.94
115.27
115.06
114.03
102.28
99.85
99.78
98.34
94.14
84.61
82.77
81.81
79.36
72.25
69.89
69.41
63.79
62.13
61.92
60.37
50.60
39.21
32.62
29.22
28.91
28.26
28.14
26.68
25.00
21.04
20.74
18.88
14.16
PhthN
N3
AcO
N-BFPM
5
210 200 190 180 170 160 150 140 130 120 110 100 90 80 70 60 50 40 30 20 10 0 -10

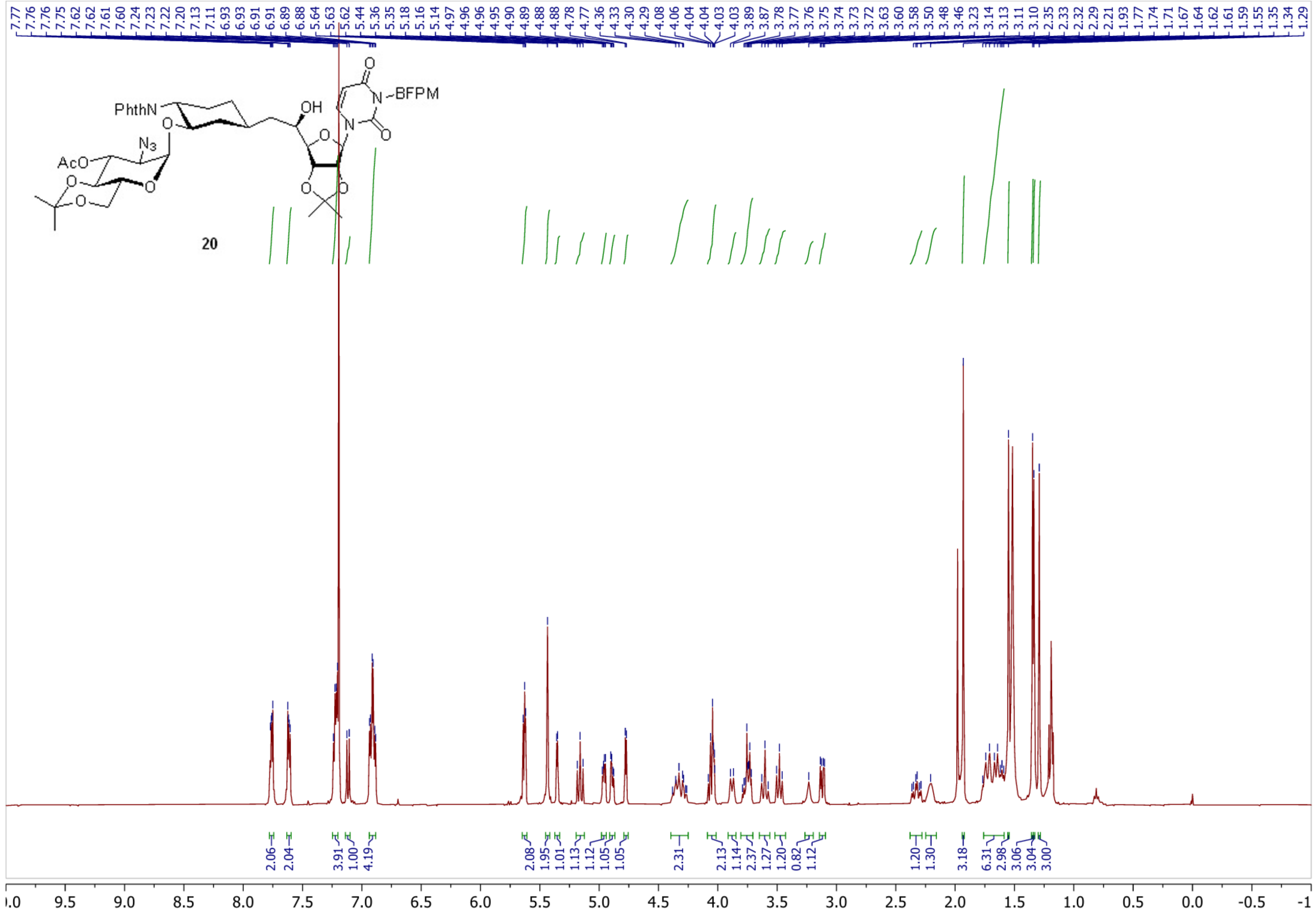

PhthN
N3
AcO
OH
N~BFPM
20

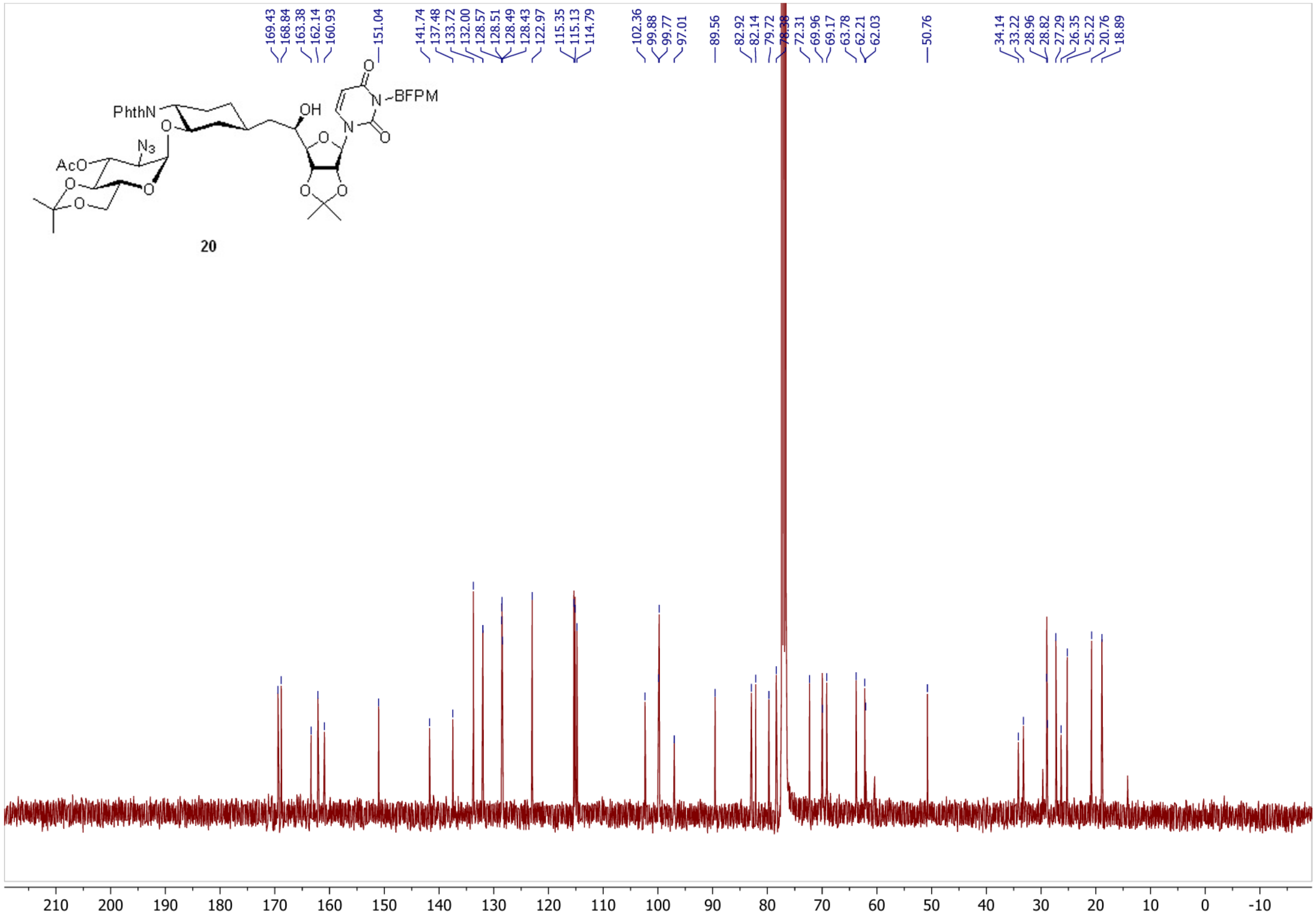
PhthN
AcO
$N_3$
OH
N–BFPM
20
169.43
168.84
163.38
162.14
160.93
151.04
141.74
137.48
133.72
132.00
128.57
128.51
128.49
128.43
122.97
115.35
115.13
114.79
102.36
99.88
99.77
97.01
89.56
82.92
82.14
79.72
78.38
72.31
69.96
69.17
63.78
62.21
62.03
50.76
34.14
33.22
28.96
28.82
27.29
26.35
25.22
20.76
18.89
210
200
190
180
170
160
150
140
130
120
110
100
90
80
70
60
50
40
30
20
10
0
-10

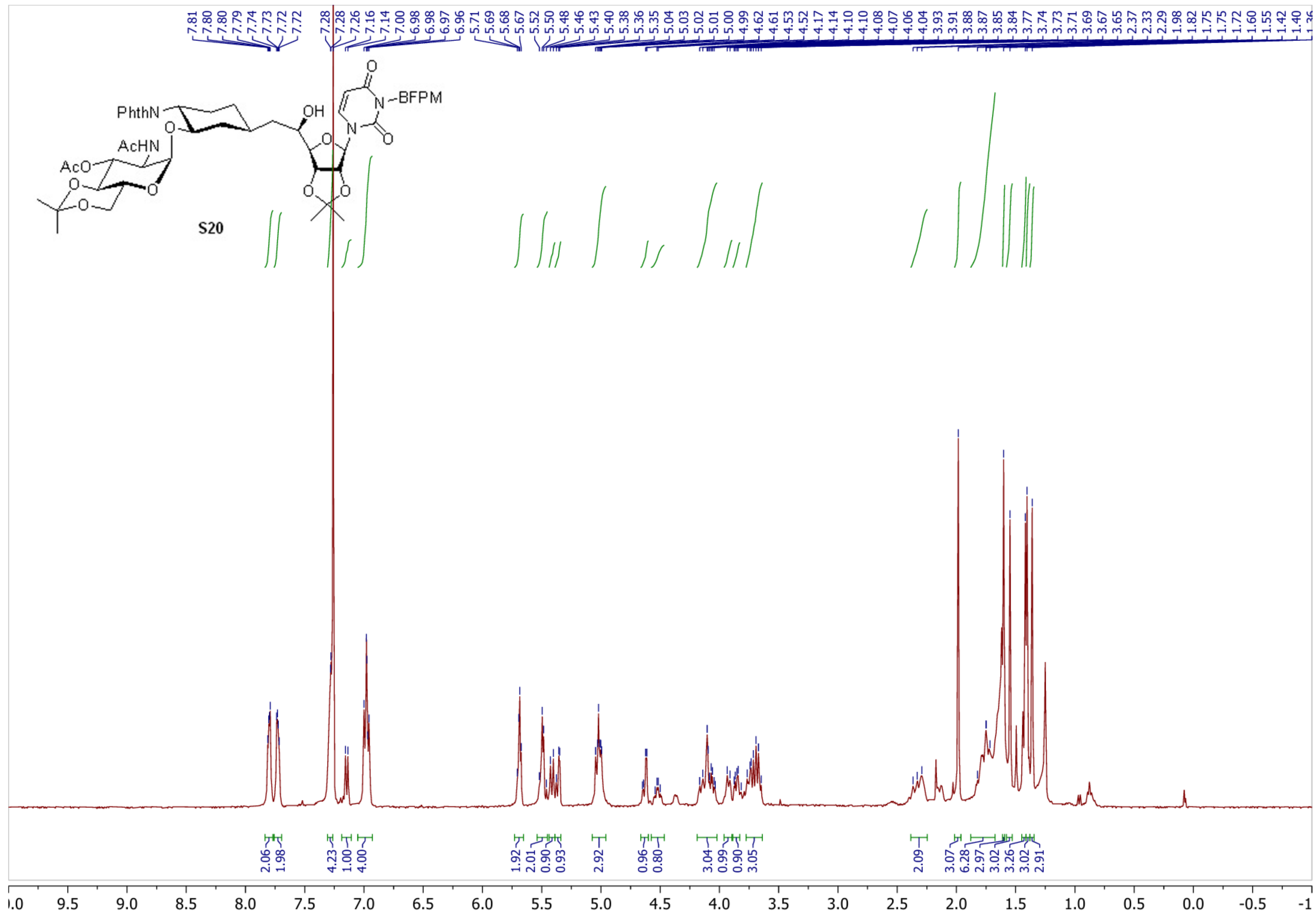

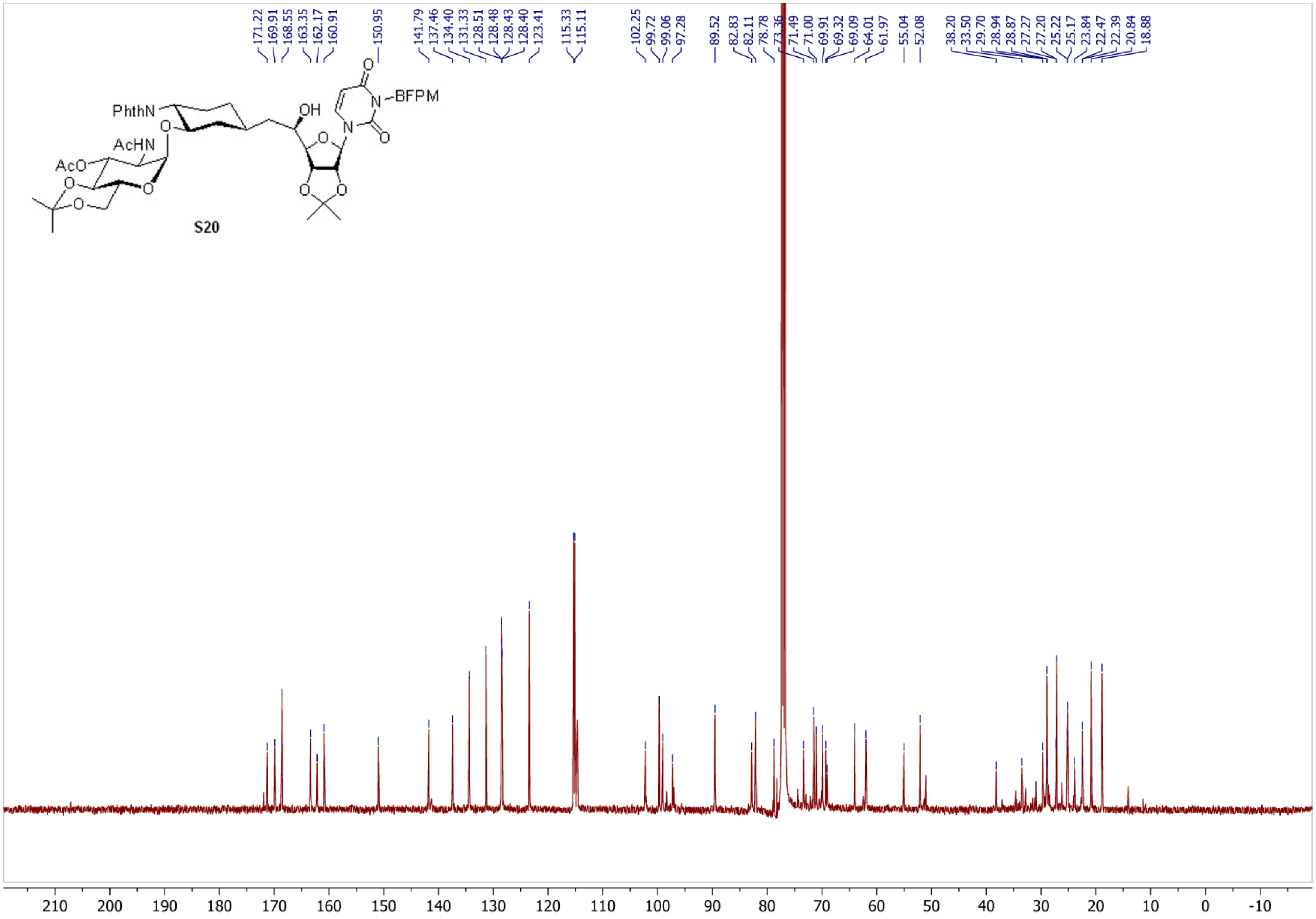

PhthN
AcHN
AcO
OH
BFPM
S20
171.22
169.91
168.55
163.35
162.17
160.91
150.95
141.79
137.46
134.40
131.33
128.51
128.48
128.43
128.40
123.41
115.33
115.11
102.25
99.72
99.06
97.28
89.52
82.83
82.11
78.78
73.36
71.49
71.00
69.91
69.32
69.09
64.01
61.97
55.04
52.08
38.20
33.50
29.70
28.94
28.87
27.27
27.20
25.22
25.17
23.84
22.47
22.39
20.84
18.88
210
200
190
180
170
160
150
140
130
120
110
100
90
80
70
60
50
40
30
20
10
0
-10

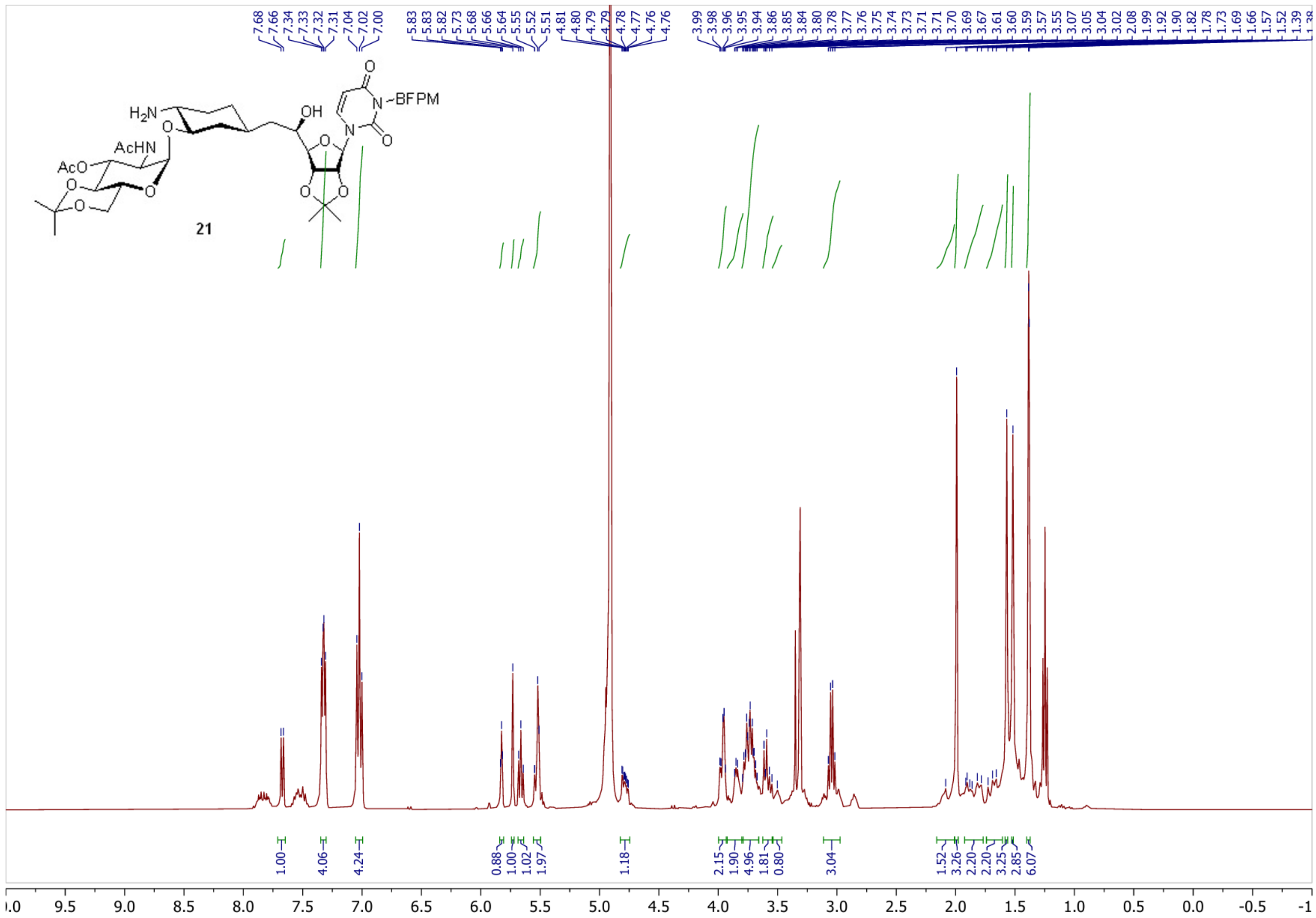

H2N
AcHN
AcO
OH
N-BFPM
21
7.68 7.66 7.34 7.33 7.32 7.31 7.04 7.02 7.00
5.83 5.83 5.82 5.73 5.68 5.66 5.64 5.55 5.52 5.51
4.81 4.80 4.79 4.79 4.78 4.77 4.76 4.76
3.99 3.98 3.96 3.95 3.94 3.86 3.85 3.84 3.80 3.78 3.77 3.76 3.75 3.74 3.73 3.71 3.71 3.70 3.69 3.67 3.61 3.60 3.59 3.57 3.55 3.07 3.05 3.04 3.02 2.08 1.99 1.92 1.90 1.82 1.78 1.73 1.69 1.66 1.57 1.52 1.39
1.00 4.06 4.24 0.88 1.00 1.02 1.97 1.18 2.15 1.90 4.96 1.81 0.80 3.04 1.52 3.26 2.20 2.20 3.25 2.85 6.07
9.5 9.0 8.5 8.0 7.5 7.0 6.5 6.0 5.5 5.0 4.5 4.0 3.5 3.0 2.5 2.0 1.5 1.0 0.5 0.0 -0.5

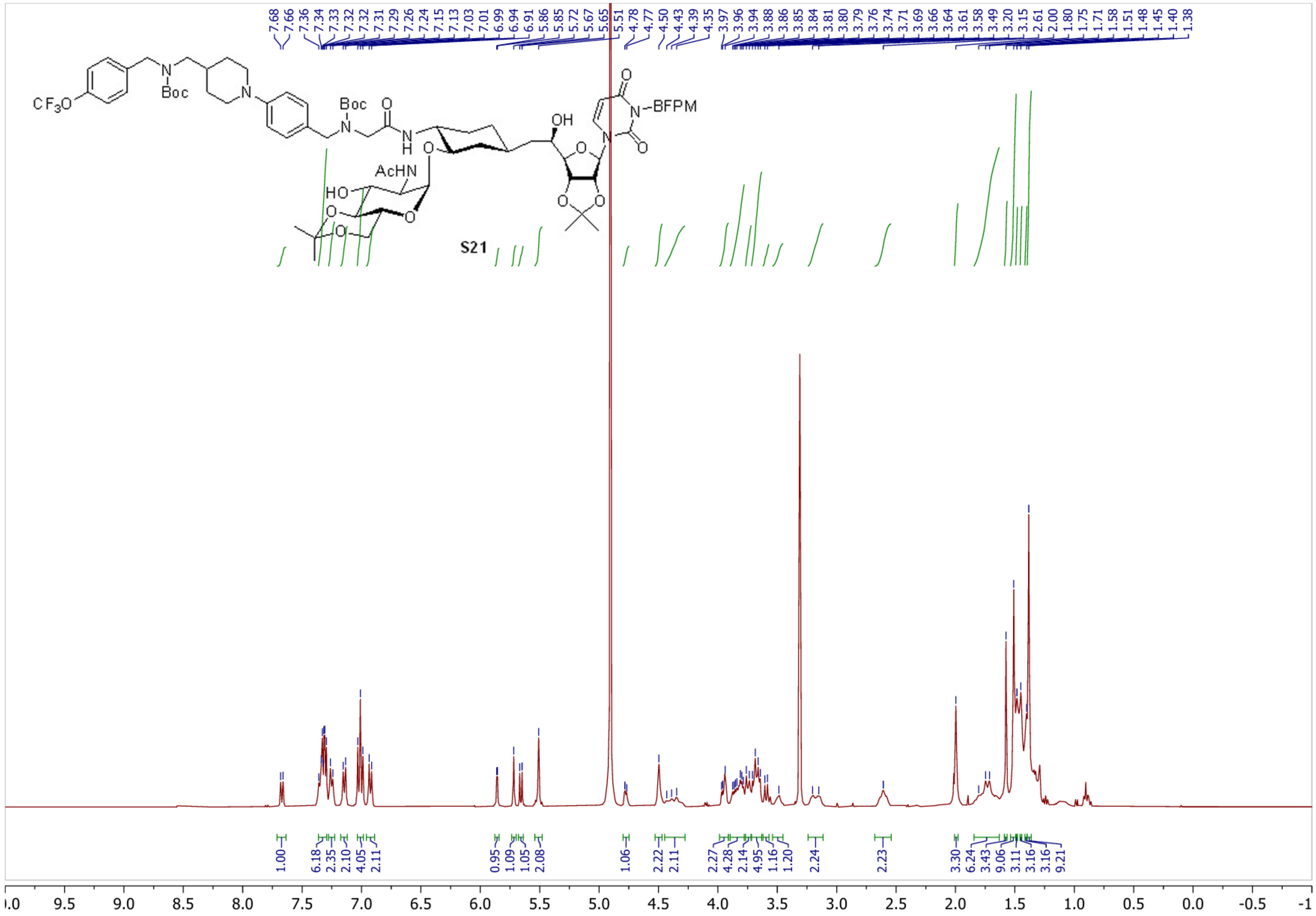

7.68 7.66 7.36 7.34 7.33 7.32 7.32 7.31 7.29 7.26 7.24 7.15 7.13 7.03 7.01 6.99 6.94 6.91 5.86 5.85 5.72 5.67 5.65 5.51 4.78 4.77 4.50 4.43 4.39 4.35 3.97 3.96 3.94 3.88 3.86 3.85 3.84 3.81 3.80 3.79 3.76 3.74 3.71 3.69 3.66 3.64 3.61 3.58 3.49 3.20 3.15 2.61 2.00 1.80 1.75 1.71 1.58 1.51 1.48 1.45 1.40 1.38
CF3O
Boc
N
Boc
O
N
H
AcHN
HO
O
O
O
O
OH
O
O
O
N
N–BFPM
O
O
S21
1.00 6.18 2.35 2.10 4.05 2.11 0.95 1.09 1.05 2.08 1.06 2.22 2.11 2.27 4.28 2.14 4.95 1.16 1.20 2.24 2.23 3.30 6.24 3.43 9.06 3.11 3.16 3.16 9.21
.0 9.5 9.0 8.5 8.0 7.5 7.0 6.5 6.0 5.5 5.0 4.5 4.0 3.5 3.0 2.5 2.0 1.5 1.0 0.5 0.0 -0.5 -1

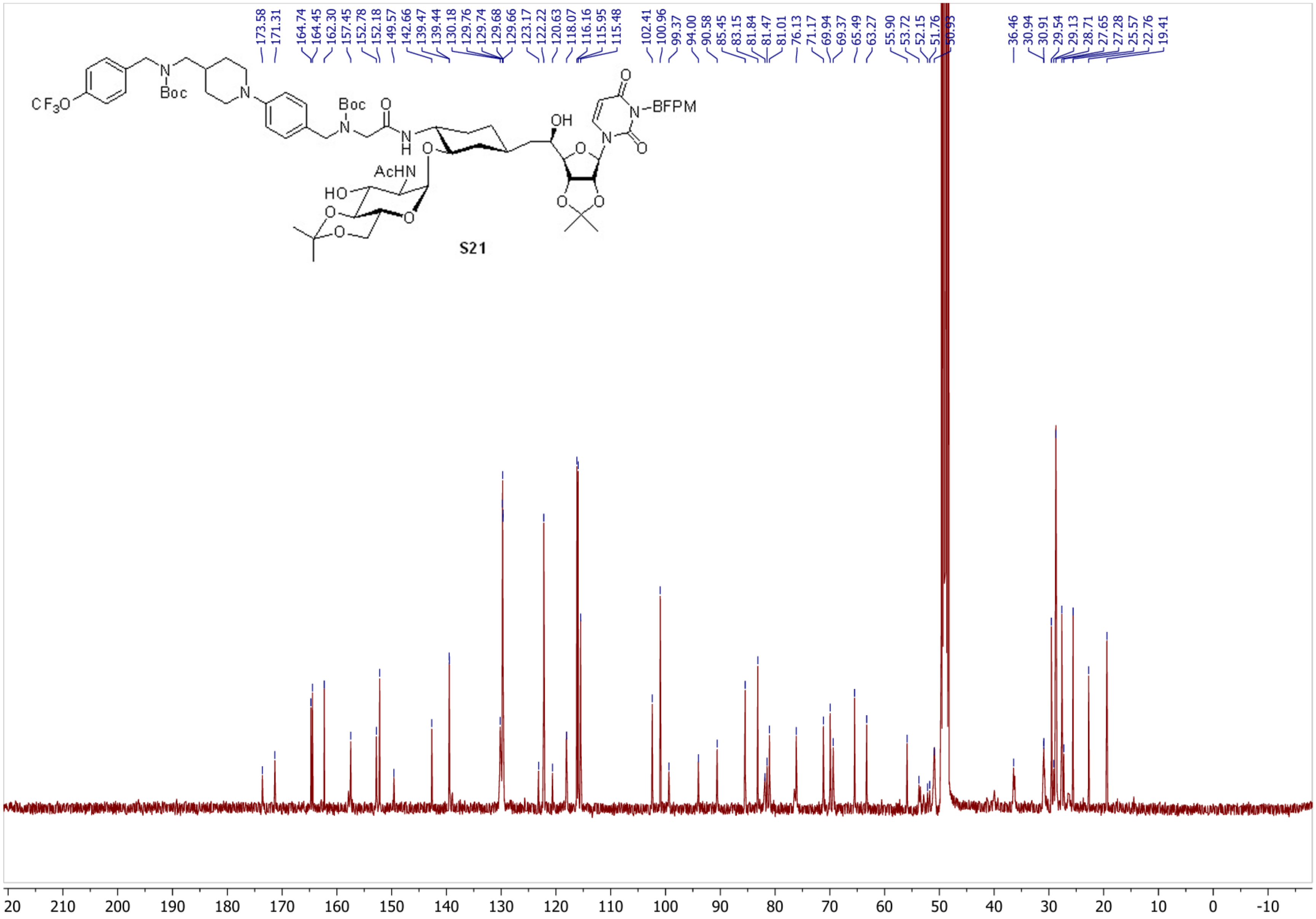

173.58
171.31
164.74
164.45
162.30
157.45
152.78
152.18
149.57
142.66
139.47
139.44
130.18
129.76
129.74
129.68
129.66
123.17
122.22
120.63
118.07
116.16
115.95
115.48
102.41
100.96
99.37
94.00
90.58
85.45
83.15
81.84
81.47
81.01
76.13
71.17
69.94
69.37
65.49
63.27
55.90
53.72
52.15
51.76
50.93
36.46
30.94
30.91
29.54
29.13
28.71
27.65
27.28
25.57
22.76
19.41
CF3O
Boc
N
Boc
O
N
H
AcHN
HO
O
O
O
OH
O
N-BFPM
S21
20 210 200 190 180 170 160 150 140 130 120 110 100 90 80 70 60 50 40 30 20 10 0 -10

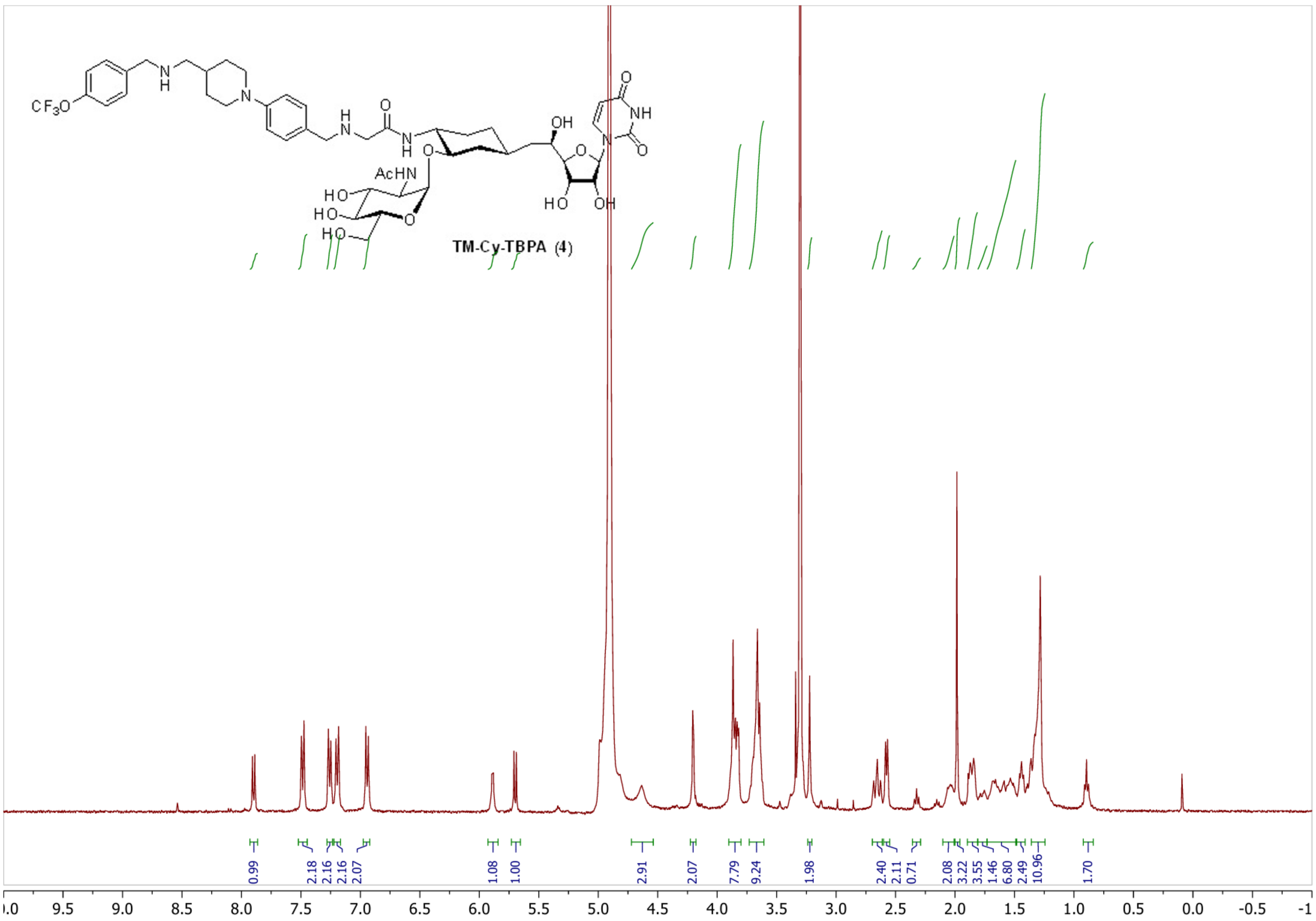
CF3O
N H
N
H N
O
N H
AcHN
HO
HO
HO
O
O
OH
O
N
NH
O
O
HO
OH
TM-Cy-TBPA (4)
0.99
2.18
2.16
2.16
2.07
1.08
1.00
2.91
2.07
7.79
9.24
1.98
2.40
2.11
0.71
2.08
3.22
3.55
1.46
6.80
2.49
10.96
1.70
9.5 9.0 8.5 8.0 7.5 7.0 6.5 6.0 5.5 5.0 4.5 4.0 3.5 3.0 2.5 2.0 1.5 1.0 0.5 0.0 -0.5 -1

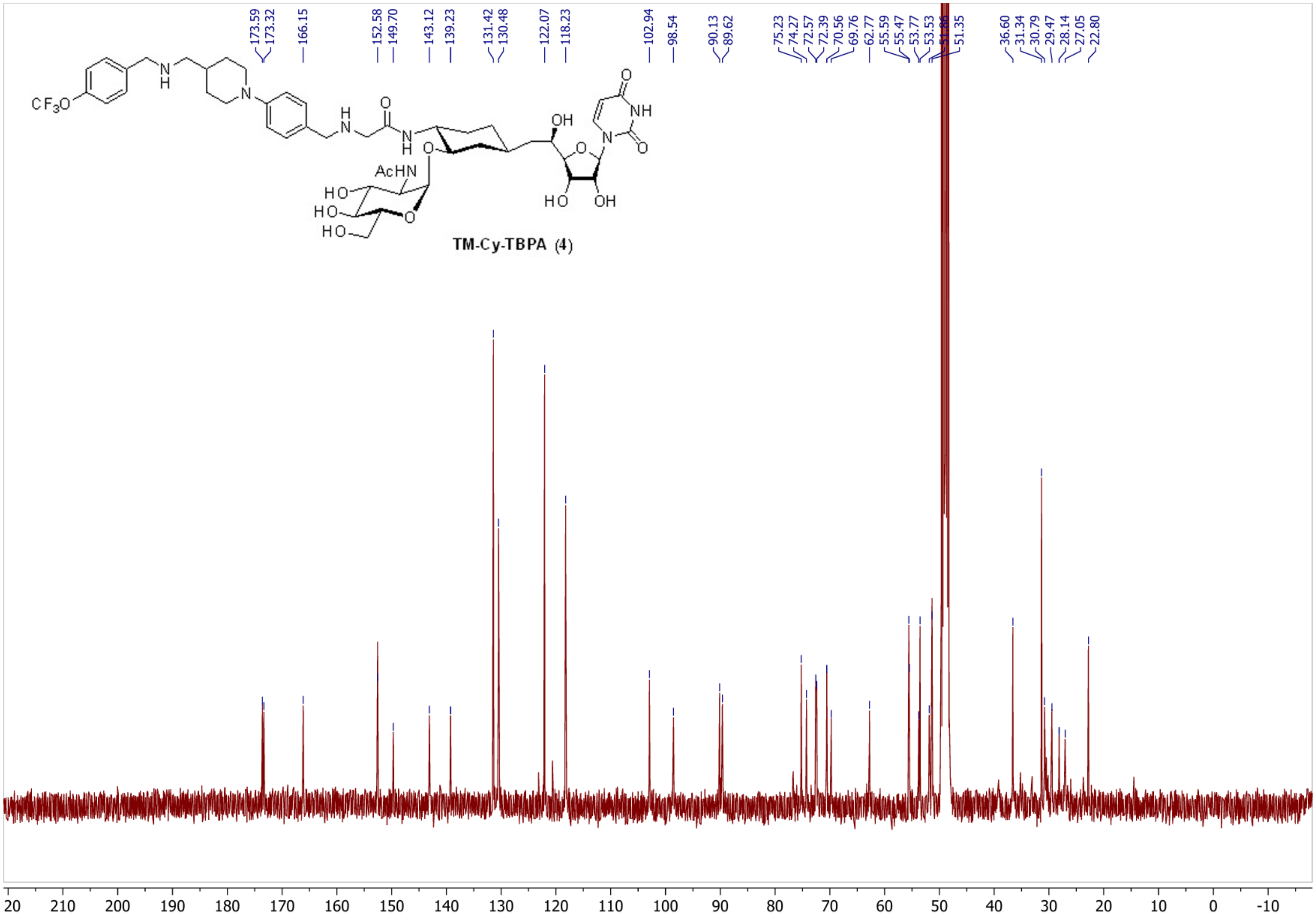

173.59
173.32
166.15
152.58
149.70
143.12
139.23
131.42
130.48
122.07
118.23
102.94
98.54
90.13
89.62
75.23
74.27
72.57
72.39
70.56
69.76
62.77
55.59
55.47
53.77
53.53
[illegible]
51.35
36.60
31.34
30.79
29.47
28.14
27.05
22.80
CF3O
N
H
N
N
H
O
N
H
O
AcHN
HO
HO
HO
O
OH
O
N
NH
O
O
HO
OH
20 210 200 190 180 170 160 150 140 130 120 110 100 90 80 70 60 50 40 30 20 10 0 -10


**TM-Cy-TBPA (4)**
